# Selective enrichment of *Methylococcaceae* versus *Methylocystaceae* methanotrophs via control of methane feeding schemes

**DOI:** 10.1101/2024.03.18.585448

**Authors:** Ju Yong Lee, Munjeong Choi, Min Joon Song, Daehyun Daniel Kim, Taeho Yun, Jin Chang, Adrian Ho, Jaewook Myung, Sukhwan Yoon

## Abstract

Methanotrophs are crucial in keeping environmental CH_4_ emissions in check. However, how different groups of methanotrophs contribute to this important role in different environmental settings remain ambiguous. Here, in a simplified laboratory setting of well-mixed batch reactors fed continuous flow of CH_4_-containing gas, methanotrophic microbiomes were enriched from paddy soils under six different incubation conditions prepared as combinations of two different CH_4_ mixing ratios (0.5% and 10% v/v) and three supplemented Cu^2+^ concentrations (0, 2, and 10 μM). Monitoring of the temporal community shifts in the reactor microbiomes observed domination of *Methylocystis* spp. in all three reactors fed 0.5% v/v, as further supported by the analyses of *pmoCAB* genes in the shotgun metagenomes of the single-point samples from the same reactors. Copper deficiency did not select for *mmoXYZ*-possessing methanotrophs. Instead, a cluster of *mbn* genes with an abundance accounting for approximately 5% of *Methylocystis* population was identified, suggesting a comparative ecological importance of methanobactin in Cu-deficient methanotrophy over soluble methane monooxygenases. These findings highlight the importance of *Methylocystis* spp. in mitigating emissions from CH_4_ hotspots, e.g., landfills and rice paddies, and suggest the feasibility of directed enrichment/isolation of *Methylocystis* spp. for utilization in, for example, methanobactin and polyhydroxybutyrate production.

**Synopsis:** This study reports enrichment of a complex soil microbiota with 0.5% methane resulting in dominance of a specific group of methane-consuming bacteria *Methylocystis*, highlighting their ecological significance as CH_4_ sink.

## INTRODUCTION

Methane (CH_4_) is a potent greenhouse gas estimated to be ∼30 times more effective in causing radiative forcing than CO_2_ on a 100-year scale, with an estimated 16% contribution to the global warming effect in accordance to the IPCC’s fifth assessment report, and due to its short lifespan mitigation of CH_4_ emissions would have an additional merit of immediacy.^1,2^ Non-energy sector emissions, accounting for >60% of the total CH_4_ emissions, being predominantly of microbial origin underlines the environmental significance of CH_4_-related microbiology and biogeochemistry.^1,3^ At its major environmental hotspots, e.g., wetlands, rice paddy soils, and landfills, CH_4_ is anaerobically produced via methanogenesis in depth and diffuses through gradually more oxidizing environments, where a substantial portion of the produced CH_4_ is oxidized via methanotrophy.^4,5^ Diverse groups of microorganisms participating in anaerobic and aerobic methanotrophy have been recently discovered; however, the aerobic methanotrophs of the *Proteobacteria* phylum are still considered the primary environmental CH_4_ sink at these hotspots.^4,6,7^ Atmospheric CH_4_ (∼1.9 ppmv) oxidation has also been attributed to the methanotrophs belonging to this phylogenetic group.^8–10^

These conventional methanotrophs include two phylogenetically distinct groups, one belonging to the *Methylocystaceae* and *Beijerinckiaceae* families (*Alphaproteobacteria*) and the other belonging to the *Methylococcaceae* family (*Gammaproteobacteria*).^11^ Despite this phylogenetic bifurcation, CH_4_-to- CH_3_OH oxidation is invariably catalyzed by either soluble or particulate methane monooxygenase (sMMO or pMMO, respectively).^11^ With basis on the physiological observations from *Methylosinus trichosporium* OB3b and *Methylococcus capsulatus* Bath, it is now widely perceived that expressions and activities of pMMO and sMMO are reciprocally regulated by copper, although ambiguities remain, however, as to the precise biochemical role of copper in pMMO has not yet been elucidated.^11–13^ Somewhat contrary to this perception, a predominantly large proportion of isolated proteobacterial methanotrophs possess only *pmo* operons, and *mmoX* genes, let alone transcripts, have rarely been observed in the environments.^14, 15^ Apart from the common reliance on MMOs, the two groups of methanotrophs differ substantially in their genomic and physiological properties.^11,16^ The methanotroph isolates with the highest CH_4_ turnover rates and the fastest growth rates (specific growth rate higher than 0.2 h^-1^) upon CH_4_-replete incubation invariably belonged to *Gammaproteobacteria*, while all isolates capable of utilizing <100 ppmv CH_4_ belonged to *Alphaproteobacteria*, suggesting possible niche specialization.^8,17–19^ Further, the ability to synthesize and utilize methanobactin, i.e., a potent copper chelator putatively utilized for copper scavenging, and polyhydroxybutyrate (PHB), an intracellular energy-storage compound and a promising building block for biodegradable plastics, have been found only among *Methylocystaceae* isolates.^20,21^

In the previous studies that examined methanotroph communities at CH_4_ hotspots with molecular analysis and sequencing techniques, selection of specific groups of methanotrophs have been witnessed in certain highly specialized environmental settings. Acidic peat bogs, typically with pH ranging between 4.5 and 5.5, were often observed as favoring *Methylocystaceae* and *Beijerinckiaceae*, while alkalic and halophilic environments, e.g., Mono Lake sediments, have tendency to be enriched with *Methylococcaceae* methanotrophs.^22–24^ Nevertheless, the methanotroph populations in paddy soils or landfill cover soils, i.e., the terrestrial CH_4_ hotspots with far larger importance to global CH_4_ emissions, have evaded generalized characterization.^5,25,26^ In such environments, characterized by high subsurface CH_4_ concentrations and circumneutral pH, methanotroph compositions vary greatly from one location to another and along the soil depth, without sufficiently explicable correlations with environmental parameters. To address this knowledge gap and improve understanding of competition between different groups of methanotrophs, the current study examined time-dependent microbial community development in batch reactors fed a continuous stream of gas containing two different concentrations of CH_4_. The three reactor runs at each CH_4_ concentration were performed with different Cu^2+^ concentrations, considering the importance of Cu^2+^ in methanotrophy. Additionally, shotgun metagenome analyses, performed with single time point samples from the reactors fed 0.5% CH_4_, semi-quantitatively investigated the functional genes involved with *Methylocystaceae*-dominated methanotrophy. Bridging culture-based and culture-independent analysis techniques, this study illustrates how methanotrophic subpopulation develops as a response to sustained CH_4_ influx within complex microbial communities under varying environmental conditions, and suggests a practical methodological approach to expedite enrichment and/or isolation of *Methylocystaceae* methanotrophs for production of methanobactin or PHB.

## MATERIALS AND METHODS

### Soil sampling and characterization

The paddy soil used as the inoculum was sampled in August 2017 from an experimental field site in Hwaseong, South Korea (37°13’21"N, 127°02’35"E) operated by Gyeonggido Agricultural Research Extension Services.^27^ The soil samples were collected from the top layer (0-20 cm depth) of submerged soil. After removing visible plant materials, the soil slurry was collected in pre-sterilized glass jars, transported to the laboratory, and stored at 4°C until use. The sample for microbial community analysis was separately stored at -80°C.

### Media preparation and general culture conditions

Unless otherwise mentioned, all experiments used nitrate mineral salts (NMS) medium, which contained per liter, 1 g MgSO_4_·7H_2_O, 1 g KNO_3_, 0.2 g CaCl_2_·2H_2_O, 0.1 mL 3.8% (w/v) Fe(III)-EDTA solution (Sigma-Aldrich, St. Louis, MO), 0.5 mL 0.1% (w/v) Na_2_MoO_4_·2H_2_O solution, and 0.1 mL 10,000X trace element solution.^28^ After autoclaving, separately prepared and sterilized 1 M pH 6.8 phosphate solution was added to a concentration of 5 mM, and filter-sterilized 1,000X vitamin stock solution was added. Glassware was washed in a 5 N HNO_3_ acid bath overnight before use, and Cu^2+^ was added as 5 mM CuCl_2_ stock solution.

### Enrichment of methanotrophic consortia under varying CH_4_ and Cu^2+^ conditions

Each enrichment process started with batch incubation in a closed vessel. The medium was distributed in a 50-mL aliquot to a 250-mL serum bottle, and after suspending 10 g wet paddy soil, the bottles were sealed with butyl-rubber stoppers and aluminum crimps. CH_4_ (>99.95%; Special Gas Co., Daejeon, South Korea) was added through a 0.2-µm syringe filter to the targeted concentration (0.5 or 10% v/v CH_4_ in the headspace). CuCl_2_ was added to the targeted concentration; however, as the soil carried 8.1±0.3 mg soluble Cu/kg dry wt, any visible Cu effect would have been highly unlikely at this stage.^27^ The culture bottles were incubated in dark at 30°C with shaking at 150 rpm. CH_4_ concentration was monitored with a GCMS-QP2010 gas chromatograph-mass spectrometer (Shimadzu Cooperation, Kyoto, Japan). Immediately after confirming CH_4_ depletion, soil particles were settled for 30 minutes, and 40 mL of the supernatant was added as the inoculum to 5 L medium prepared in a BioFlo^®^ 120 bioreactor (Eppendorf, Hamburg, Germany). Air containing 0.5% or 10% (v/v) CH_4_ was introduced into the reactor through a microsparger at a constant flowrate of 200 mL min^-1^. The temperature and pH were automatically controlled at 30°C and pH 6.8 (with 0.1 M HCl and NaOH), respectively, and the medium was agitated at 500 rpm. For each CH_4_ concentration, methanotrophic consortia were enriched in the medium prepared with three supplemented Cu^2+^ concentrations (0, 2, and 10 μM). The Cu^2+^ carryover from the soil into the reactor culture was unlikely to exceed 0.1 μmoles L^-1^. Each enrichment was performed independently, starting with the soil stored at 4°C. Cell growth was monitored using a GENESYS 30 visible spectrometer (Thermo Fisher Scientific, Waltham, MA), and at each sampling point, a pellet from 2-mL suspension was stored at -20°C. At the end of each enrichment, a 100-mL suspension was stored at 4°C for isolation of methanotrophs.

### Microbial community analyses via 16S rRNA amplicon sequencing

The microbial community composition shifts during enrichment were monitored with 16S rRNA gene amplicon sequencing. Genomic DNA was extracted from the soil or the suspensions from the reactors using DNeasy PowerSoil Pro Kit (Qiagen, Hilden, Germany) according to the protocol provided by the manufacturer. The V6-V8 hypervariable region was amplified using the 926F (5’-AAACTYAAAKGAATTGRCGG- 3′)/1392R (5’-ACGGGCGGTGTGTRC-3′) primer pair, and the amplicons were sequenced using an Illumina MiSeq sequencing platform (San Diego, CA). The sequence data were processed using the QIIME2 v2021.11 pipeline as described in detail in the supporting information.^29^

### Sequencing and analyses of the metagenomes

Three samples from the reactor cultures incubated with 0.5% CH_4_, one per each copper concentration selected based on the growth curve and the community profiles, were subjected to shotgun metagenome sequencing for further examination of these *Methylocystis*-dominated microbiomes. Shotgun sequencing was performed on an Illumina NovaSeq 6000 platform, generating ∼5 Gb of paired-end reads (2×150 bp) per sample. The raw sequences from both the 16S rRNA amplicon sequencing and shotgun metagenome sequencing were deposited in the NCBI SRA database under the BioProject accession number PRJNA940290. Analyses of the metagenomes were performed with focus on the functional genes relevant to aerobic methanotrophy, i.e., the *pmo*, *mmo*, and *mbn* genes, and the extracted metagenome-assembled genomes (MAG). For functional gene analyses, both the *de novo* assembly approach and the HMMER screening approach were used, as described in detail in the supporting information.^30,31^

### Isolation of methanotrophs from the fed-batch enrichments

After the end of each fed-batch incubation, 10 µL of the enrichment culture was spread onto fresh NMS agar plates, which were then incubated in a BBR^®^ GasPak^®^ 150 anaerobic jar (Becton Dickinson, Franklin Lakes, NJ). The CH_4_ mixing ratio and Cu^2+^ concentration were the same as those used for the reactor incubation. The jar was incubated in dark at 30°C, with the gas exchanged every day. Single colonies were transferred to fresh agar plates and incubated under the same condition.

After three transfers, the single colonies were used as the inoculums for batch incubation in aqueous NMS medium. The isolates were taxonomically classified with Sanger sequencing of 27F/1492R amplicons. The absence of secondary peak in the electropherogram was taken as a line of evidence for culture purity.

## RESULTS

### Enrichment of methanotrophic microbial consortia under varying CH_4_ and Cu^2+^ conditions

Microbial communities distinct from that of the inoculum developed in the reactor cultures enriched under six different conditions, as combinations of two CH_4_ feeding schemes (0.5% v/v and 10% v/v) and three supplemented Cu^2+^ concentrations (0, 2, and 10 µM). The CH_4_ concentration in the feed gas had a conspicuous effect on the growth of the overall microbial population. The exponential growth rates of the reactor cultures incubated with 10% v/v CH_4_ were significantly higher (0.019±0.008 h^-1^ as averaged across the incubations with three different Cu^2+^ concentrations) than those incubated with 0.5% v/v CH_4_ (0.005±0.002 h^-1^; *p*<0.05, Kruskal-Wallis test; Figure 1). Throughout incubation, none of the enrichment cultures showed a significantly lowered CH_4_ concentration in the gas effluent (data not shown), indicating that only an insignificant proportion of CH_4_ in the gas stream was utilized.

**Figure 1.**
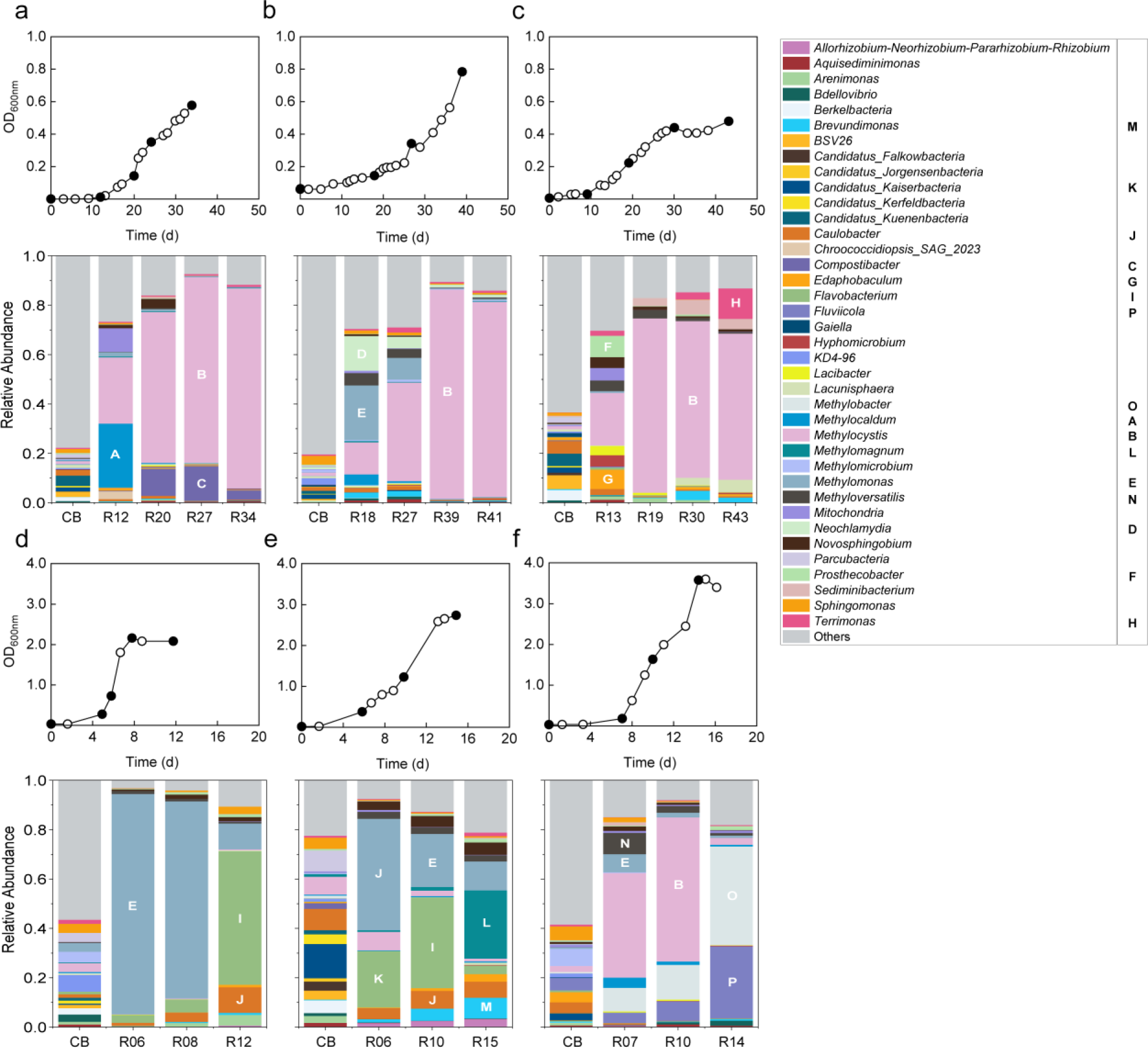
Growth curves and time series changes of microbial compositions of the reactor cultures fed air with 0.5% (a, b, c) and 10% (d, e, f) v/v CH_4_. Each CH_4_ feeding scheme was examined with three different concentrations of Cu^2+^ supplemented to the aqueous medium: 0 (a, d), 2 (b, e), and 10 µM (c, f). The time points at which samples were collected for microbial community analyses are indicated with filled circles. The genus level taxa with >3% relative abundance in any of the samples are represented in the bar charts. For convenience in cross-matching, the taxa specifically mentioned in the main text are denoted with alphabets A-P both in the bar charts and in the legend. CB: closed batch enrichment; R#: reactor enrichment at # hours after start of incubation.

The shifts in the overall microbial communities from the initial inoculum through the enrichment processes showed evident bifurcation between those incubated with two different CH_4_ mixing ratios (Figure 1, 2). The batch enrichments in serum bottles (i.e., the inoculum for the reactor culture with the same CH_4_ and Cu conditions), even though having been enriched with 826 µmoles (10%-CH_4_ cultures) and 41 µmoles of CH_4_ (0.5%-CH_4_ cultures), did not exhibit any particularly enriched methanotrophic taxon or clustering tendency (Figure 1, 2). The lack of clustering pattern at this stage could not have been due to the effect of the copper concentration, as the soil particles carried 0.58 μmoles of soluble copper into the 50-mL enrichments (Figure 2b). Invariable across the incubation conditions, reactor incubation decreased the microbial diversity substantially from the initial paddy soil community (PS) and the batch enrichment (CB), as indicated by the Shannon-Wiener diversity indices (Figure 1, S2). The community compositions of the enrichment cultures collected at different time points from the same reactor generally clustered together in both the hierarchically-clustered heatmap and the NMDS plot, suggesting that the community composition was determined early and largely maintained during the exponential growth (Figure 1, 2). For example, the biomass density increased 2.2-fold between 19-43 d in the 0.5%-CH_4_ 10-μM-Cu^2+^ reactor culture, but with little change in its taxonomic composition.

**Figure 2.**
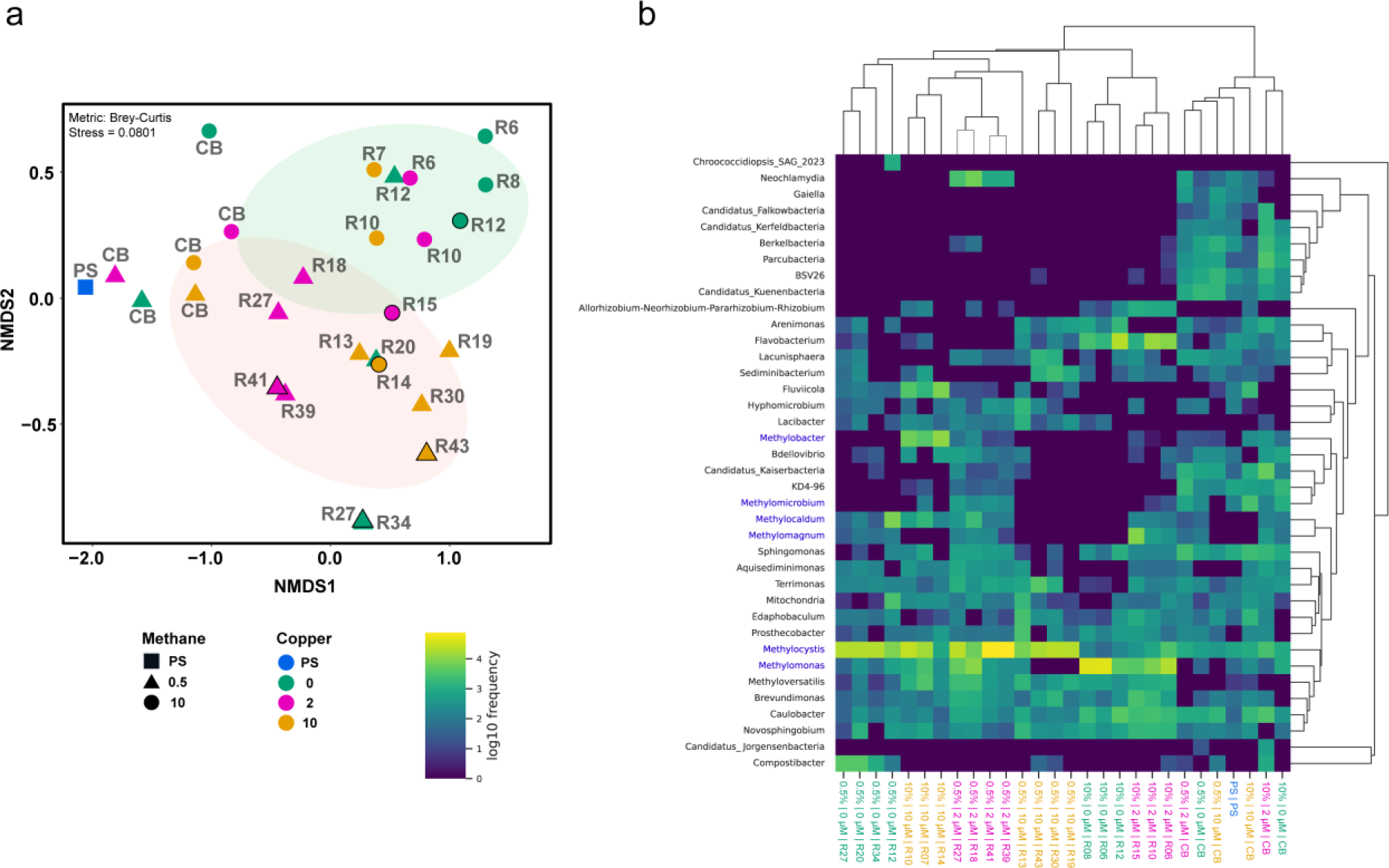
(a) Non-metric multidimensional scaling (NMDS) plot illustrating the beta-diversity of the enrichment cultures. Counts of the individual ASVs were used as inputs, and distances between the communities were calculated using Bray-Curtis distance metric. (b) Hierarchically-clustered heatmap visualizing relative abundances of the genus-level taxa in the culture samples. Only the taxa with >3% abundance in any of the samples were presented. Weighted Bray-Curtis metric was used for clustering of taxa (vertical) and samples (horizontal).

The rice paddy soil used as the inoculum contained diverse methanotrophic genera from both the *Alphaproteobacteria* (*Methylocystis* and *Methylosinus*) and *Gammaproteobacteria* (*Methylobacter*, *Methylocaldum*, *Methylomagnum*, *Methylomicrobium*, and *Methylomonas*) classes with the cumulative relative abundances of 0.9% and 0.4%, respectively (Table S1). Initial batch incubation in the serum bottles did not substantially enrich either group. The cumulative relative abundance of the methanotrophic genera belonging to *Alphaproteobacteria* and those belonging to *Gammaproteobacteria* amounted to only 3.3±2.5% and 7.3±2.9%, respectively, of the total microbial population in the 10%-CH_4_ cultures. The methanotroph relative abundances in the 0.5%-CH_4_ enrichments were statistically undistinguishable from that of the rice paddy community (*p*>0.05).

Further enrichments in the fed-batch reactors with continuous flow-throughs of CH_4_-containing air invariably resulted in aerobic methanotrophs dominating the microbial community, regardless of the CH_4_ mixing ratio in the gas stream or the concentration of supplemented Cu^2+^ in the aqueous medium (Figure 1). As to which aerobic methanotroph taxa were enriched appeared largely dependent on the CH_4_-feeding scheme (Figure S1). The methanotrophs belonging to *Alphaproteobacteria* outcompeted those belonging to the *Gammaproteobacteria* in all three 0.5%-CH_4_ enrichments. *Methylocystis* spp. were consistently the most abundant methanotrophic taxon in these cultures regardless of the Cu^2+^ concentration. Although several methanotrophic taxa affiliated to the *Gammaproteobacteria* phylum were transiently enriched (e.g., up to 27.0% in the 2-µM-Cu^2+^ culture), their total relative abundance was <1.5% at the end of incubation. The growth curves and the time series of the community composition profiles together infer that the overall biomass increase in the 0.5%-CH_4_ cultures could be explained mostly with the growth of *Methylocystis* spp.

Contrastingly, in the 10%-CH_4_ cultures, the methanotrophs affiliated to *Gammaproteobacteria* eventually became more abundant than those affiliated to *Alphaproteobacteria*, although methanotroph communities were not dominated by a single genus (Figure 1). In the Cu^2+^-free culture, *Methylomonas* spp. became the predominant constituents of the microbial community after 6 days, and although its relative abundance eventually decreased to 10.4% at 12 d (i.e., 4 days after the cell density growth had ceased), *Methylomonas* spp. was still the most abundant among the methanotrophic taxa. In the 2 µM-Cu^2+^ enrichment, *Methylomonas* was the most abundant taxon 6 days after start of incubation, but *Methylomagnum*, also affiliated to the *Gammaproteobacteria* class, overtook as the most abundant methanotroph taxon at the end of incubation (27.7% at 15 d). Interestingly, this particular taxon was not enriched at any other condition. In the 10 µM-Cu^2+^ reactor culture, *Methylocystis* spp. was the predominant taxon earlier during the incubation with its relative abundance reaching 57.2% at 10 d. However, *Methylobacter* spp. eventually became the predominant taxonomic group when the growth came to a halt after 14 days (39.4%).

### Core microbiome analysis of the methanotrophic enrichments

The core microbiome, defined here as the genus-level taxa that were present at >0.5% relative abundance in all 18 microbiomes from the six reactors, comprised of 28 genus-level taxa, which includes well-known methanotrophic genera *Methylomonas*, *Methylocaldum*, *Methylobacter*, *Methylomagnum*, *Methylosinus* and *Methylocystis* (Figure 3). The most interesting was the strong negative correlation between *Methylocystis* and *Methylomonas*, both among the most abundant taxa in the 0.5%-CH_4_ and 10%-CH_4_ reactor microbiomes, respectively (*r*=-0.84, *p*=1.5_E-5, Spearman’s rank correlation_; Figure 3). This correlation implies a competitive relationship between these two taxa, which may have been dictated by CH_4_ concentration. Interestingly, several significant positive correlations, but no negative correlation, were identified among the methanotrophic taxa belonging to *Gammaproteobacteria*, suggesting that these methanotrophs may favor similar incubation conditions, probably plentitude of dissolved CH_4_, but are not under a mutually exclusive competition. The strongly positive correlation between *Methylomonas* and *Flavobacterium* (*r*=0.77, *p*=1.6_E-4_) is also of note, as *Flavobacterium* was among the most abundant taxa in the 10%-CH_4_ Cu^2+^-free and 2-μM-Cu^2+^ reactor cultures at multiple time points. This correlation implies that *Methylomonas* may have some form of metabolic association with *Flavobacterium*, which is absent with *Methylocystaceae* methanotrophs.

**Figure 3.**
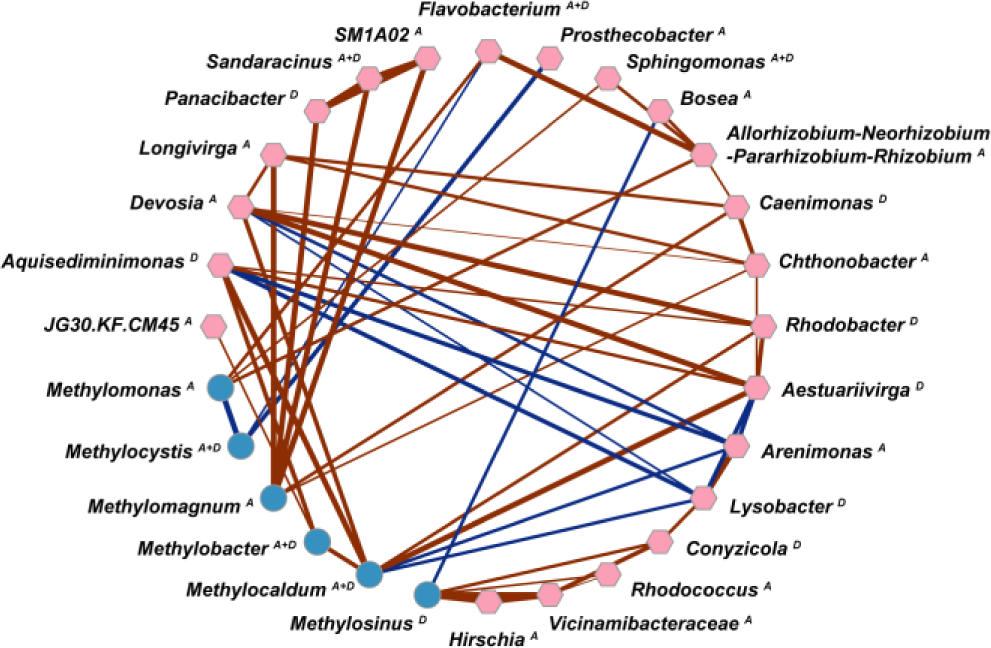
Co-occurrence network of the core microbiome drawn from the 18 reactor culture communities consisting of the 28 genus-level taxa found in all samples with relative abundances higher than 0.5%. Pairwise Spearman’s rank correlation was used to determine the strength and statistical significance of the correlation. Only strong (|r|>0.7; *p*<0.05) and statistically significant (*p*<0.05) correlations are shown with red (positive) and blue (negative) edges, thickness drawn to be proportional to |r|. Nodes with the denotation A are aggregates of ASVs assigned the same genus-level taxa with Naive Bayes classifier, and those with denotation D are OTUs clustered *de novo* from the pool of ASVs lacking genus-level assignments. The names of the OTUs are the best BLAST hits of the consensus sequence and thus, may not reflect accurate denomination. Visualization of the network used the Cytoscape v. 3.9.1 software.

### Analyses of methanotroph functional genes in the 0.5%-CH_4_ reactor microbiomes

The microbial samples collected from the three 0.5%-CH_4_ reactors, dominated by *Methylocystis* spp., were subjected to shotgun metagenome sequencing (Figure 4, 5). Searches for the *pmoCAB* genes using HMMER returned *pyrG*-normalized coverage values (RPKM of the target gene divided by RPKM of *pyrG*) of 1.70 (*pmoC*), 1.28 (*pmoA*), and 1.06 (*pmoB*) in the Cu-free enrichment. These numbers were in line with the relative abundance of putative methanotrophic taxa deduced from the 16S rRNA amplicon sequencing data. The *pmoCAB* coverages were lower in the metagenomes of the 2 μM-Cu^2+^ and 10 μM-Cu^2+^ reactor enrichments despite the similar *Methylocystis* relative abundances in these samples to that of the Cu-free enrichments. The coverage of *pmoC* (1.00), *pmoA* (0.23), and *pmoB* (0.47) in the 2 μM-Cu^2+^ enrichments and those in the 10 μM-Cu^2+^ reactor enrichment (1.08, 0.76, and 0.45, respectively) were nevertheless in the same order of magnitude as expected from the 16S rRNA-based microbial community profiles. The phylogenetic breakdown of the *pmoCAB* genes in all three metagenomes exhibit a strong bias to the cluster affiliated to *Methylocystis* sp. OBBP, with >50.4%, >98.6%, and >92.6% of mapped *pmoC*, *pmoA*, and *pmoB* reads, respectively, belonging to this taxon. No *pmoCAB* gene affiliated to *Gammaproteobacteria* was recovered in Cu-free and 10 μM-Cu^2+^ enrichments. The 2 μM-Cu^2+^ enrichment metagenome contained *Gammaproteobacteria pmoCAB* genes, albeit at low coverage (0.47, 0.56, and 6.13% of all *pmoC*, *pmoA*, and *pmoB* reads, respectively, were mapped to the clusters affiliated to *Gammaproteobacteria*).

**Figure 4.**
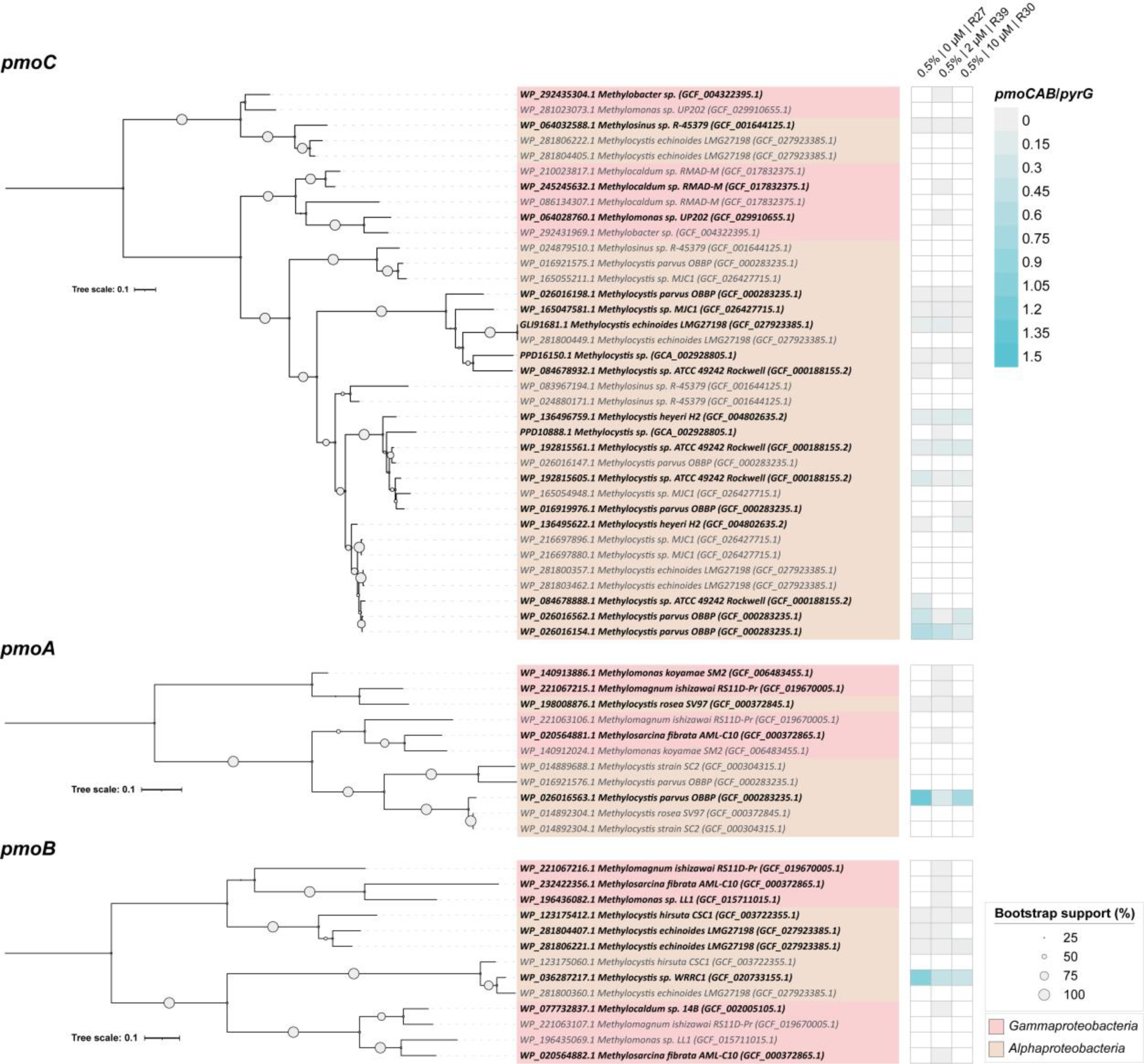
(a) Maximum-likelihood trees illustrating the phylogenetic placements of the clusters of *pmoCAB* genes recovered from the 0.5%-CH_4_ metagenomes. Sizes of the circles at the junctures indicate the bootstrap values. Taxa in bold represent the clusters of the metagenomic contigs and those in plain italics are reference sequences from the nr database. (b) Heatmaps showing the relative abundance of the *pmoCAB* clusters in the metagenomes from the reactor cultures with three different Cu^2+^ concentrations in the medium (Cu-free, 2 µM Cu^2+^, and 10 µM Cu^2+^). RPKM values of the *pmo* genes normalized by that of *pyrG* in the metagenome was used as the measure of relative abundance.

**Figure 5.**
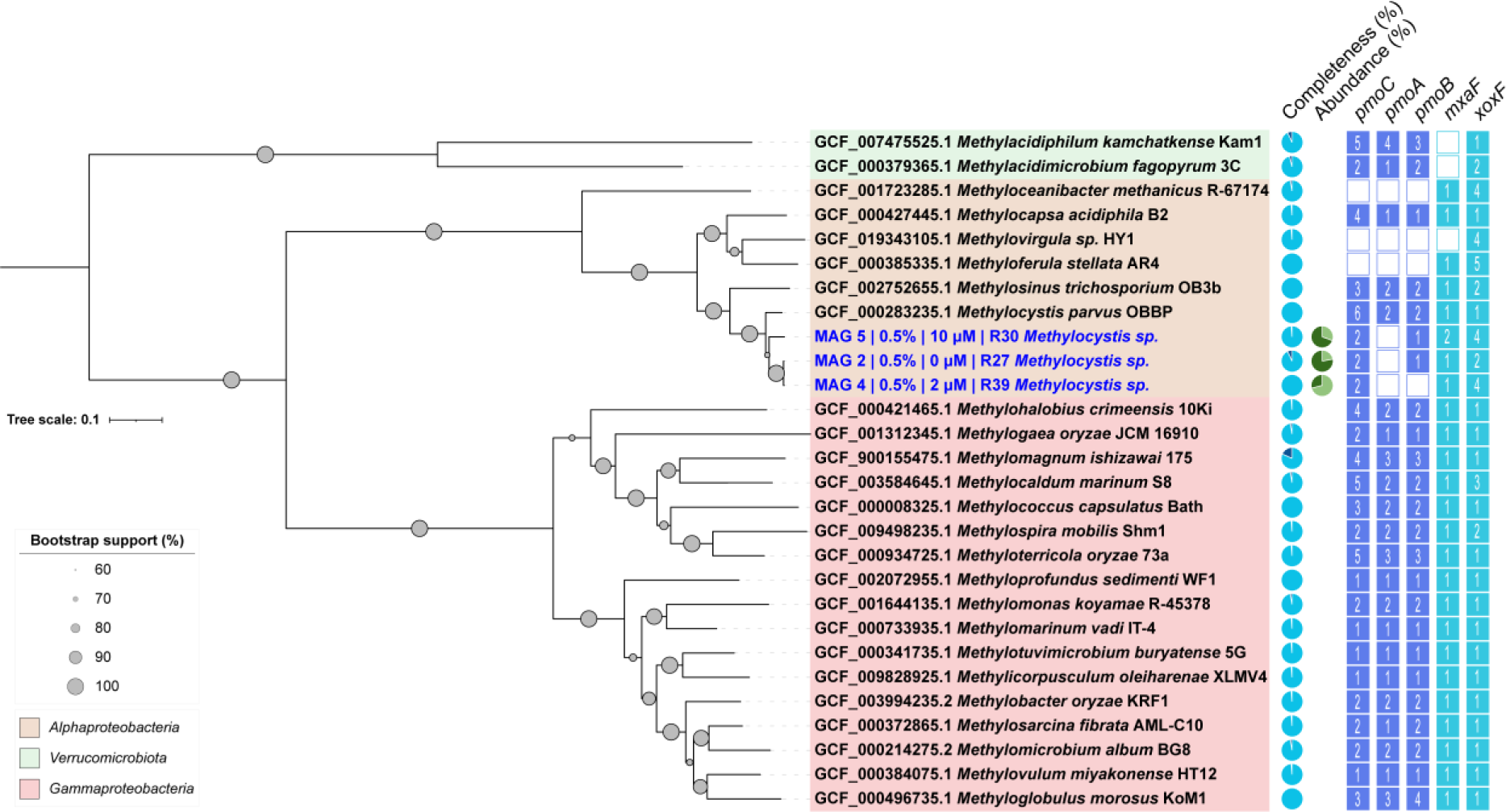
Genomic phylogenetic tree constructed with Anvi’o workflow illustrating the evolutionary placement of the three putatively methanotrophic MAGs reconstructed from the three 0.5%-CH_4_ metagenomes. Concatenated amino acid sequences of 15 ribosomal proteins were constructed from these three MAGs and 25 complete/draft methanotroph genomes. Visual output of the phylogenetic tree was generated using the iTOL v. 6.6 software. Sizes of the circles at the junctures indicate the bootstrap values. The presence or absence of the genes known to be essential for aerobic methanotrophy, i.e., *pmo* and *mxa*/*xox* genes, in the MAGs/genomes is shown next to the phylogenetic tree. The numbers indicate the number of the specified functional gene in the MAGs/genomes.

With the *de novo* assembly-first pipeline more frequently employed in target-gene-focused metagenomic analyses, the coverage values of *pmoA* were estimated to be only 0.009 and 0.019 in Cu-free and 10 μM-Cu^2+^ reactor cultures, respectively, orders of magnitudes lower than those expected from the *Methylocystis* abundance (Figure S3). This discrepancy suggested that the majority of the *pmoA* raw reads had evaded *de novo* assembly, and thus, questions the validity of the methodology when analyzing complex methanotrophic cultures. Initial screening with HMMER, albeit still imperfect, substantially improves the diversity of *pmoA* sequences that can be reconstructed from metagenomes, and thus, reliability of quantification (Figure 4; Table S2).

Despite the low availability of copper in the Cu-free enrichment, unlikely to exceed 92 nM even accounting for the carry-over from the soils, no *mmoXYZ* was detected via either gene detection approach. On the other hand, *de novo* assembly identified *mbnABC* as a part of a 4.2-kb operon that includes accessory genes putatively involved in regulation, maturation, and transport of methanobactin. This cluster of *mbn* genes was present only in this Cu-free culture metagenome, with the *pyrG*-normalized coverage of 0.03, suggesting a minor but substantial potential for methanobactin synthesis (Figure S3b). The translated amino acid sequences of all the genes in this putative methanobactin operon were nearly identical (>98.2%) to those previously found in the genomes of *Methylocystaceae* methanotrophs (Table S3). The microorganism possessing this operon, putatively a *Methylocystis* sp. and the sole producer of methanobactin, probably constituted at least 5% of the total *Methylocystaceae* population in the Cu-free reactor microbiome, as inferred from the coverage.

### Analyses of the MAGs recovered from the 0.5%-CH_4_ reactor metagenomes

A total of 10 MAGs (2, 2, and 6 from the Cu-free, 2-μM-Cu^2+^, and 10-μM-Cu^2+^ enrichments, respectively) passed the >90% completeness and <10% contamination cutoffs (Table S4). A MAG affiliated to *Methylocystis* was obtained from each of the three analyzed metagenomes. MAG2 reconstructed from the Cu-free enrichment was essentially identical to MAG4 from the 2-μM-Cu^2+^ enrichment (99.9% ANI) and shared 80.4% ANI with the MAG5 from the 10-μM-Cu^2+^ enrichment. All three MAGs, according to the GTDB-tk classification, were most closely affiliated to the genome of *Methylocystis* sp. OBBP, but still with substantial genome-wide difference, as indicated by the ANI values around 80%. The *Methylocystis* spp. represented by these MAGs had sizeable population in all three enrichments, with 20.7%, 70.5%, and 34.1% of the total quality-trimmed raw reads mapped onto them in the Cu-free, 2-μM-Cu^2+^ and 10-μM-Cu^2+^ enrichments, respectively. As expected from the tendency of *pmoA* to evade assembly and binning, *pmoA* was found in none of the MAGs. Stand- alone *pmoC*’s were found in all three MAGs. Additionally, a 3’-truncated *pmoC* was found in MAG2 and MAG4 and a 5’-truncated *pmoB* in MAG2 and MAG5. All these *pmoC* and *pmoB* genes were most similar to those of *Methylocystis* sp. OBBP, congruent with the genome-wide GTDB-tk classification. All three MAGs possess *mxaF* and/or *xoxF* encoding two distinct types of methanol dehydrogenases all closely affiliated to those of *Methylocystis* sp. OBBP, corroborating that the microorganisms represented by these MAGs are methanotrophs. Neither *mmo* nor *mbn* operon was found in any of the MAGs. Considering the complete absence of HMMER-screened *mmo* reads and the low coverage of the reconstructed *mbn* operon, the absence of either operon was very unlikely due to assembly and/or binning errors. Another noteworthy genomic observation was the presence of the *phaC* gene encoding a polyhydroxyalkanoate synthase in all three MAGs.

A gene annotated as *pmoA* found in MAG6 affiliated to *Lacunisphaera* of the phylum *Verrucomicrobiota* demands scrutiny. Despite its annotation, this gene is marginally similar to the typical *pmoA* genes of the confirmed methanotrophs (31.6%, 30.6%, and 31.4% amino-acid-level similarity to *Methylosinus trichosporium* OB3b, *Methylomonas methanica*, and *Methylacidiphilum fumariolicum* SolV; the *pmoA*-annotated gene was not captured by the HMMER search). As the *pmoA*-like gene was a standalone gene and MAG6 lacked either methanol hydrogenase, the microorganism represented by this MAG is probably not methanotrophic.

Including MAG6, seven of the ten extracted MAGs were non-methanotroph genomes. The functional gene inventories of several of the MAGs hint the sources of electrons for some of these organisms, probably unable to utilize the only provided electron donor CH_4_. MAG3, recovered from the 2-μM-Cu^2+^ culture and closely affiliated with a confirmed methylotrophic species *Methyloversatilis discipulorum* (ANI of 94.0%, AAI of 96.2%), possesses *mxa*, *fae*, *mtd*, *fhc*, *fwd*, and *fdo* genes/operons encoding the enzymes catalyzing methanol oxidation pathway terminally resulting in CO_2_. This putative methylotroph probably subsisted on leakage of one-carbon intermediates from the methanotrophs (Table S5). MAG7, MAG8, and MAG9 contained the *fdoG*, *fdhF*, and/or *fdwA* encoding the catalytic subunits of formate dehydrogenases, suggesting the possibility of formate leakage from methanotrophs and subsequent utilization by these heterotrophs. Although experimental evidence of extracellular release of sugars and organic acids (e.g., pyruvate, succinate, and citrate) has not yet been reported, the presence of complete glycolysis and tricarboxylic acid cycle in MAG1, MAG6, MAG7, and MAG8 suggest that these putative heterotrophs may subsist on the simple organic substrates exuded from the methanotrophs or, also possibly, from degradation of dead methanotroph biomass. The genes that encode high-affinity hydrogenases, i.e., *hucSL* and *hucLS*, were absent in any of the MAGs, precluding the possibility that atmospheric H_2_ was utilized as the electron donor for these organisms.

### Directed isolation of methanotrophs from the reactor cultures

At least one methanotrophic isolate was obtained from each reactor (Table S6, Figure S4). As anticipated from the community analyses, all of the isolates obtained from the 0.5%-CH_4_ enrichments were affiliated to *Methylocystis*, according to the 1465-bp partial 16S rRNA gene sequences (Figure S4). The 16S rRNA gene sequences of *Methylocystis* strains isolated from the 0.5% CH_4_ cultures, one from the Cu^2+^-free condition and the other from 10-μM-Cu^2+^ condition, were >99.8% identical to each other and to *Methylocystis* strain La-a12, once again supporting that copper deficiency is unlikely to be a determinant selective pressure. The isolates obtained from the 10% CH_4_ culture were taxonomically more diversified, and include those classified as *Methylomonas*, *Methylomagnum*, and *Methylocystis*. All three isolates belonging to the *Methylomonas* genus, one from each different copper condition, were nearly identical to each other and to *Methylomonas* sp. R-45378 (100%, 100%, and 99.81% for percent identities). The *Methylomagnum* sp. isolated from the 2-μM-Cu^2+^ enrichment had 16S rRNA sequence substantially differing from those of *Methylomonas* spp. (<88.0% identity). The *Methylocystis* sp. isolated from the 2-μM-Cu^2+^ enrichment was closely affiliated to *Methylocystis* sp. PRM2 (99.63% 16S rRNA sequence identity) but was distinct from the *Methylocystis* isolates obtained from the 0.5%-CH_4_ enrichments (<95.3%).

## DISCUSSION

Methane concentration is one of the most important factors to consider when investigating the biogeochemical role of methanotrophs as an environmental CH_4_ sink or developing a biotechnology for CH_4_ mitigation.^4,6,7,32,33^ Recent studies have verified the existence of microorganisms with an extraordinary capability to oxidize and grow on atmospheric CH_4_ and shown that even several familiar laboratory strains of methanotrophs, including *Methylotuvimicrobium buryatense* 5GB1C, can consume CH_4_ at concentrations as low as 200 ppmv.^8,33^ The current study adds onto these recent advances that CH_4_ concentration, even at a much higher range than that considered atmospheric, can be a major pressure towards selection of distinct groups of methanotrophs. Both the 16S rRNA-based community analyses and the metagenomic functional gene analyses have shown that *Methylocystis* spp. eventually dominated all three 0.5%-CH_4_ reactor cultures, regardless of the Cu^2+^ concentration. In a stark contrast, 10%-CH_4_ enrichments, performed in parallel as high-CH_4_-concentration controls, lacked such consistency, although the methanotrophic taxa belonging to *Gammaproteobacteria* eventually outcompeted those belonging to *Methylocystis* spp. at all three Cu^2+^ conditions. *Methylocystis* spp. and methanotrophs belonging to *Alphaproteobacteria* have previously been suggested as more competitive under low-CH_4_ environments (e.g., <1% v/v); however, direct experimental evidence had been lacking.^34–36^ The current research presents clear evidence such selection may take place at environmental CH_4_ hotspots.

The predominance of *Methylocystis* spp., in the 0.5%-CH_4_ microbiomes is due to reasons apart from microbial kinetics. The vast majority of *pmoA* genes (>99.4% in all three reactors) recovered from the metagenomes were very similar to those of *Methylocystaceae* methanotrophs, including those with kinetically characterized strains, e.g., *Methylocystis* sp. LR1, which exhibited no particularly high affinity to CH_4_ as compared to the methanotrophs belonging to *Gammaproteobacteria*.^35^ The *Methylocystis*-specific *pmoA*, previously termed as *pmoA2*, was absent in any of the analyzed metagenomes, and no *Methylocapsa* or upland soil cluster alpha (USCα) sequence was identified from either 16S rRNA amplicon sequencing or metagenomic analyses.^8,19^ The actual dissolved CH_4_ concentration could have been substantially lower than that calculated assuming an equilibrium between the gaseous and aqueous phases; however, even so, the absence of the aforementioned sequences precludes a possible involvement of the methanotrophs or pMMO previously characterized with high affinities to CH_4_. A more plausible explanation is that the dominant *Methylocystis* spp. may harbor a distinguishably efficient means to synthesize biomass with energy and carbon from a limited source.^33,35^ The ability to accumulate and utilize polyhydroxybutyrate (PHB) could also have been a crucial factor to the selection of *Methylocystis* spp over the methanotrophs affiliated to *Gammaproteobacteria*. The genes encoding the PHB synthesis pathway are found in all sequenced *Methylocystis* genomes, as was the case with the three *Methylocystis* MAGs in this study.^37–39^ The spatiotemporal variations in CH_4_ availability within the reactor would have been more probable in the 0.5%-CH_4_ cultures than the 10%-CH_4_ cultures. With occasional starvation spells, the capability to store PHB and utilize it along with CH_4_ as a reducing equivalent would endow a comparative advantage.^38^ It would be interesting to observe whether and how the expression of the PHB synthase genes, at both transcription and translation level, respond to CH_4_ feeding schemes in the ensuing investigations.

The current study suggests that the ecological role that sMMO may play in methanotrophy under Cu^2+^ deficiency may have been over-emphasized.^11^ The difficulty of detecting *mmoX* genes and transcripts in environmental microbiomes apart from acidic wetlands where *Methylocella* spp. and *Methyloferula* spp. thrive has casted doubts on this decades-old ecological rationalization of sMMO possession.^15,23,41,42^ Consistent with these previous observations, the Cu-free incubation failed to enrich, let alone select for, *mmoX*-possessing methanotrophs. Rather, the metagenomic investigations suggested the importance of methanobactin-producing methanotrophs in Cu-deficient environments, as the cluster of *mbn* genes was found only in the culture without Cu^2+^ supplementation. It is highly probable that the *mbn*-possessing *Methylocystis* sp. was selected by Cu deficiency. The estimated relative abundance of *mbnA* was <10% of the *Methylocystis* population; however, ‘piracy’ of extracellular Cu-bound methanobactin, i.e., uptake and utilization of Cu bound to methanobactin synthesized by cohabiting methanotrophs, has been repeatedly observed.^43–45^ Possibly, methanobactin produced by this minor but still sizeable subgroup may have aided communal expression and synthesis of functional pMMO among *Methylocystis*. In support of this hypothesis, the putatively methanotrophic constituents in the Cu-free and 10-μM-Cu^2+^ enrichments, were virtually indistinguishable apart from this single cluster of *mbn* genes, as inferred from the compositions of the metagenomic *pmoA* pool.

Metagenomic or metatranscriptomic investigations of environmental *pmo* genes are rare, even in this era of excessive sequencing.^46,47^ Although not having been explicitly stated in literature, *pmoCAB* operon, and *pmoA* in particular, has tendency to evade *de novo* metagenome assembly, due to its presence in most known proteobacterial methanotroph genomes as nearly identical duplicates.^48–50^ *De novo* assembly failed to produce quantitatively sensible *pmoA* constructs from the Cu-free and 10-μM-Cu^2+^ enrichment metagenomes (0.009 and 0.019, respectively). Assembly was probably further compounded by microdiversity among *pmoA* of the *Methylocystis* spp. in these enrichments, as the more uniform 2-μM-Cu^2+^ enrichment, with 70.5% of the reads mapped onto a single *Methylocystis* MAG, yielded a much more credible coverage value of 1.02. The *pmoA* genes reconstructed from the HMMER-screened reads were much more agreeable with the amplicon-sequencing-based *Methylocystis* abundance, despite the moderate discrepancy. These findings show that mechanical analyses of *pmoA* abundance and composition in complex microbiomes using pipelines starting with *de novo* sequencing, as have quite often been performed recently, may lead to severely erroneous results.

The importance of CH_4_ as a potent greenhouse gas and the importance of methanotrophs as its sink have been emphasized for decades.^1,3,4^ Most of the previous laboratory investigations looking into this aspect of methanotrophy were performed either at the initial CH_4_:air mixing ratios considered to be near-optimal for accommodation of the highest CH_4_ oxidation rates, i.e., 10-20% v/v CH_4_ in air, or those considered to be relevant to (circum-)atmospheric CH_4_ uptake (<500 ppmv CH_4_).^8,33,34,36,51–53^ The range of mixing ratios in between attracted little attention, although much of microbial CH_4_ oxidation occurs at this concentration range at CH_4_ hotspots such as landfills and paddy soils.^54^ The near exclusive selection of *Methylocystis* upon incubation under 0.5% CH_4_ gas demonstrated the importance of this tight group of organisms in keeping CH_4_ emissions from the hotspots in check, which had been shrouded by inconspicuous kinetic characteristics and functional gene inventories. This study also verified the unlikelihood of sMMO involvement in environmental CH_4_ oxidation apart from those of *Methylocella* spp. and *Methyloferulla* spp. lacking pMMO.^15,55,56^ From an application perspective, the feasibility to selectively isolate *Methylocystis* spp. would provide interesting opportunities for metabolic engineering and PHB production.^37–38,57,58^ Selective enrichment without supplemented Cu may also be implemented as a preparation step for acquisition of new *Methylocystis* isolates wielding methanobactin, with great clinical potential as medication for Wilson’s disease.^59^

## ASSOCIATED CONTENT

### Supporting Information

The following files are available free of charge.

- Supplemental information text; four figures: schematic depiction of the reactor setup; Shannon-Weiner indices of the enrichments; relative abundances of *pmo*, *mmo,* and *mbn* genes identified in the contigs constructed via *de novo* assembly and the orientation of the genes in the sole *mbn* cluster; 16S rRNA gene phylogenetic tree showing the placement of 8 new methanotroph isolates among known methanotrophs (DOCX)
- Nine tables: microbial community compositions organized at genus level; validation of HMMs for *pmoCAB* and *mmoXYZ* genes; genes constituting the *mbn* cluster in the Cu-free culture metagenome; list of MAGs acquired from the metagenomes; inventory of key functional genes in the MAGs; methanotroph isolates obtained from the reactor enrichments; curated list of *pmoCAB* and *mmoXYZ* gene sequences used for HMM construction; list of reference *pmoCAB* and 16S rRNA sequences; inventory of 15 ribosomal protein subunits in the reference methanotroph sequences used for construction of the whole-genome phylogenetic tree (XLSX)

## AUTHOR INFORMATION

### Corresponding Author

**Sukhwan Yoon** − Department of Civil and Environmental Engineering, Korea Advanced Institute of Science and Technology (KAIST), Daejeon 34141, South Korea; orcid.org/0000-0002-9933-7054; Phone: +82 10-3461-2024; Fax: +82 42-350-3610

### Authors

**Ju Yong Lee −** Department of Civil and Environmental Engineering, Korea Advanced Institute of Science and Technology (KAIST), Daejeon 34141, South Korea; orcid.org/0000-0001-7243-9483

**Munjeong Choi −** Department of Civil and Environmental Engineering, Korea Advanced Institute of Science and Technology (KAIST), Daejeon 34141, South Korea

**Min Joon Song −** Department of Civil and Environmental Engineering, Korea Advanced Institute of Science and Technology (KAIST), Daejeon 34141, South Korea

**Daehyun D. Kim −** Department of Civil and Environmental Engineering, University of California, Berkeley, CA, USA; orcid.org/0000-0002-7234-4721

**Taeho Yun −** Department of Civil and Environmental Engineering, Korea Advanced Institute of Science and Technology (KAIST), Daejeon 34141, South Korea

**Jin Chang −** Department of Civil and Environmental Engineering, Korea Advanced Institute of Science and Technology (KAIST), Daejeon 34141, South Korea

**Adrian Ho −** Nestlè Research, CH 1000 Lausanne 26, Switzerland

**Jaewook Myung −** Department of Civil and Environmental Engineering, Korea Advanced Institute of Science and Technology (KAIST), Daejeon 34141, South Korea

### Author Contributions

SY conceived and supervised the study. JYL, MJC, and SY designed experiments. JYL and MJC performed experiments. JYL, MJS, DDK, TY, and JC analyzed experimental and sequencing data. JYL and SY wrote manuscript with contributions from JC, AH, and JM. All authors critically reviewed the manuscript.

### Notes

The authors declare that they have no conflict of interest.

## ACKNOWLEDGEMENTS

This work was financially supported by the National Research Foundation of Korea (2015M3D3A1A01064881, 2020R1C1C1007970, 2022R1A4A5031447). MS was supported by NRF (2021R1I1A1A01054770).

## For Table of Contents Only

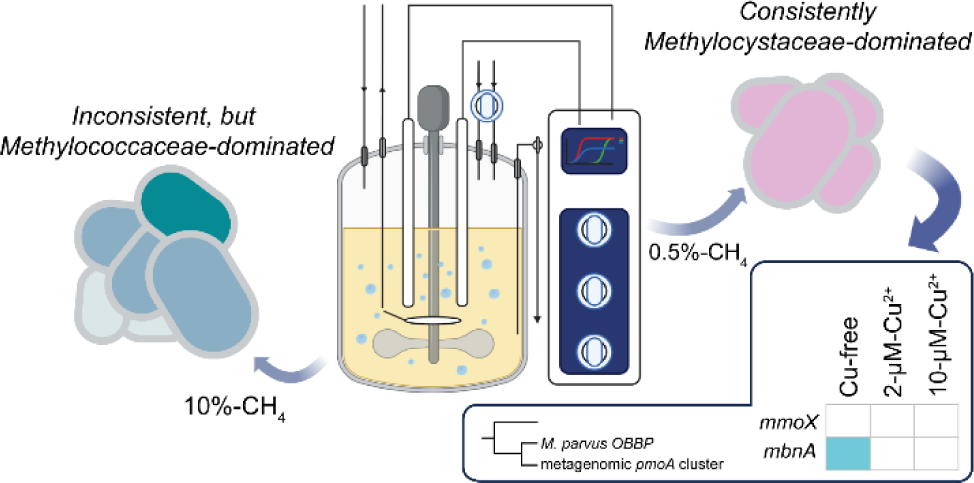

## REFERENCE

(1) Myhre, G.; Shindell, D.; Bréon, F. M.; Collins, W.; Fuglestvedt, J.; Huang, J.; Koch, D.; Lamarque, J. F.; Lee, D.; Mendoza, B.; Nakajima, T.; Robock, A.; Stephens, G.; Takemura, T.; Zhang, H. Anthropogenic and natural radiative forcing. In Climate Change 2013: *The Physical Science Basis. Contribution of Working Group I to the Fifth Assessment Report of the Intergovernmental Panel on Climate Change*, Stocker, T. F., Qin, D., Plattner, G. K., Tignor, M., Allen, S. K., Doschung, J., Nauels, A., Xia, Y., Bex, V., Midgley, P. M. Eds.; Cambridge University Press, 2013; pp 659–740.

(2) Allen, M. R.; Shine, K. P.; Fuglestvedt, J. S.; Millar, R. J.; Cain, M.; Frame, D. J.; Macey, A. H. A solution to the misrepresentations of CO_2_-equivalent emissions of short-lived climate pollutants under ambitious mitigation. *npj Clim*. Atmos. Sci. 2018, 1 (1), 16.

(3) IEA(2023). *Global Methane Tracker* 2023; IEA, Paris, https://www.iea.org/reports/global-methane-tracker-2023.

(4) Bridgham, S. D.; Cadillo-Quiroz, H.; Keller, J. K.; Zhuang, Q. Methane emissions from wetlands: biogeochemical, microbial, and modeling perspectives from local to global scales. Glob. Change Biol. 2013, 19 (5), 1325–1346.

(5) Lee, H. J.; Jeong, S. E.; Kim, P. J.; Madsen, E.; Jeon, C. O. High resolution depth distribution of Bacteria, Archaea, methanotrophs, and methanogens in the bulk and rhizosphere soils of a flooded rice paddy. Front. Microbiol. 2015, 6, 639.

(6) Ho, A.; Lee, H. J.; Reumer, M.; Meima-Franke, M.; Raaijmakers, C.; Zweers, H.; de Boer, W.; van der Putten, W. H.; Bodelier, P. L. E. Unexpected role of canonical aerobic methanotrophs in upland agricultural soils. Soil Biol. Biochem. 2019, 131, 1–8.

(7) Täumer, J.; Kolb, S.; Boeddinghaus, R. S.; Wang, H.; Schöning, I.; Schrumpf, M.; Urich, T.; Marhan, S. Divergent drivers of the microbial methane sink in temperate forest and grassland soils. Glob. Change Biol. 2021, 27 (4), 929–940.

(8) Tveit, A. T.; Hestnes, A. G.; Robinson, S. L.; Schintlmeister, A.; Dedysh, S. N.; Jehmlich, N.; von Bergen, M.; Herbold, C.; Wagner, M.; Richter, A.; Svenning, M. M. Widespread soil bacterium that oxidizes atmospheric methane. Proc. Natl. Acad. Sci. 2019, 116 (17), 8515–8524.

(9) Kolb, S. The quest for atmospheric methane oxidizers in forest soils. Environ. Microbiol. Rep. 2009, 1 (5), 336–346.

(10) Cai, Y.; Zheng, Y.; Bodelier, P. L. E.; Conrad, R.; Jia, Z. Conventional methanotrophs are responsible for atmospheric methane oxidation in paddy soils. Nat. Commun. 2016, 7 (1), 11728.

(11) Semrau, J. D.; DiSpirito, A. A.; Yoon, S. Methanotrophs and copper. FEMS Microbiol. Rev. 2010, 34 (4), 496–531.

(12) Ross, M. O.; MacMillan, F.; Wang, J.; Nisthal, A.; Lawton, T. J.; Olafson, B. D.; Mayo, S. L.; Rosenzweig, A. C.; Hoffman, B. M. Particulate methane monooxygenase contains only mononuclear copper centers. Science 2019, 364 (6440), 566–570.

(13) Nielsen, A. K.; Gerdes, K.; Murrell, J. C. Copper-dependent reciprocal transcriptional regulation of methane monooxygenase genes in *Methylococcus capsulatus* and *Methylosinus trichosporium*. Mol. Microbiol. 1997, 25 (2), 399–409.

(14) Reim, A.; Lüke, C.; Krause, S.; Pratscher, J.; Frenzel, P. One millimetre makes the difference: high-resolution analysis of methane-oxidizing bacteria and their specific activity at the oxic–anoxic interface in a flooded paddy soil. ISME J. 2012, 6 (11), 2128–2139.

(15) Liebner, S.; Svenning, M. M. Environmental transcription of *mmoX* by methane-oxidizing *Proteobacteria* in a Subarctic Palsa peatland. Appl. Environ. Microbiol. 2013, 79 (2), 701–706.

(16) Kwon, M.; Ho, A.; Yoon, S. Novel approaches and reasons to isolate methanotrophic bacteria with biotechnological potentials: recent achievements and perspectives. Appl. Microbiol. Biotechnol. 2019, 103 (1), 1–8.

(17) Kim, J.; Kim, D. D.; Yoon, S. Rapid isolation of fast-growing methanotrophs from environmental samples using continuous cultivation with gradually increased dilution rates. Appl. Microbiol. Biotechnol. 2018, 102 (13), 5707–5715.

(18) de la Torre, A.; Metivier, A.; Chu, F.; Laurens, L. M. L.; Beck, D. A. C.; Pienkos, P. T.; Lidstrom, M. E.; Kalyuzhnaya, M. G. Genome-scale metabolic reconstructions and theoretical investigation of methane conversion in *Methylomicrobium buryatense* strain 5G(B1). Microb. Cell Factories. 2015, 14 (1), 188.

(19) Baani, M.; Liesack, W. Two isozymes of particulate methane monooxygenase with different methane oxidation kinetics are found in *Methylocystis* sp. strain SC2. Proc. Natl. Acad. Sci. 2008, 105 (29), 10203–10208.

(20) Pieja, A. J.; Rostkowski, K. H.; Criddle, C. S. Distribution and selection of poly-3-hydroxybutyrate production capacity in methanotrophic proteobacteria. Microb. Ecol. 2011, 62 (3), 564–573.

(21) DiSpirito, A. A.; Semrau, J. D.; Murrell, J. C.; Gallagher, W. H.; Dennison, C.; Vuilleumier, S. Methanobactin and the link between copper and bacterial methane oxidation. Microbiol. Mol. Biol. Rev. 2016, 80 (2), 387–409.

(22) Dedysh, S. N.; Panikov, N. S.; Tiedje, J. M. Acidophilic methanotrophic communities from *Sphagnum* peat bogs. Appl. Environ. Microbiol. 1998, 64 (3), 922–929.

(23) Lin, J.-L.; Joye, S. B.; Scholten, J. C. M.; Schäfer, H.; McDonald, I. R.; Murrell, J. C. Analysis of methane monooxygenase genes in Mono Lake suggests that increased methane oxidation activity may correlate with a change in methanotroph community structure. Appl. Environ. Microbiol. 2005, 71 (10), 6458–6462.

(24) Dedysh, S. N.; Derakshani, M.; Liesack, W. Detection and enumeration of methanotrophs in acidic *Sphagnum* peat by 16S rRNA fluorescence *In situ* hybridization, including the use of newly developed oligonucleotide probes for *Methylocella palustris*. Appl. Environ. Microbiol. 2001, 67 (10), 4850–4857.

(25) Stralis-Pavese, N.; Sessitsch, A.; Weilharter, A.; Reichenauer, T.; Riesing, J.; Csontos, J.; Murrell, J. C.; Bodrossy, L. Optimization of diagnostic microarray for application in analysing landfill methanotroph communities under different plant covers. Environ. Microbiol. 2004, 6 (4), 347–363.

(26) Kumaresan, D.; Abell, G. C. J.; Bodrossy, L.; Stralis-Pavese, N.; Murrell, J. C. Spatial and temporal diversity of methanotrophs in a landfill cover soil are differentially related to soil abiotic factors. Environ. Microbiol. Rep. 2009, 1 (5), 398–407.

(27) Chang, J.; Kim, D. D.; Semrau, J. D.; Lee, J. Y.; Heo, H.; Gu, W.; Yoon, S. Enhancement of nitrous oxide emissions in soil microbial consortia via copper competition between proteobacterial methanotrophs and denitrifiers. Appl. Environ. Microbiol. 2021, 87 (5), e02301–02320.

(28) Whittenbury, R.; Phillips, K. C.; Wilkinson, J. F. Enrichment, isolation and some properties of methane-utilizing bacteria. Microbiology 1970, 61 (2), 205–218.

(29) Bolyen, E.; Rideout, J. R.; Dillon, M. R.; Bokulich, N. A.; Abnet, C. C.; Al-Ghalith, G. A.; Alexander, H.; Alm, E. J.; Arumugam, M.; Asnicar, F.; Bai, Y.; Bisanz, J. E.; Bittinger, K.; Brejnrod, A.; Brislawn, C. J.; Brown, C. T.; Callahan, B. J.; Caraballo-Rodríguez, A. M.; Chase, J.; Cope, E. K.; Da Silva, R.; Diener, C.; Dorrestein, P. C.; Douglas, G. M.; Durall, D. M.; Duvallet, C.; Edwardson, C. F.; Ernst, M.; Estaki, M.; Fouquier, J.; Gauglitz, J. M.; Gibbons, S. M.; Gibson, D. L.; Gonzalez, A.; Gorlick, K.; Guo, J.; Hillmann, B.; Holmes, S.; Holste, H.; Huttenhower, C.; Huttley, G. A.; Janssen, S.; Jarmusch, A. K.; Jiang, L.; Kaehler, B. D.; Kang, K. B.; Keefe, C. R.; Keim, P.; Kelley, S. T.; Knights, D.; Koester, I.; Kosciolek, T.; Kreps, J.; Langille, M. G. I.; Lee, J.; Ley, R.; Liu, Y.-X.; Loftfield, E.; Lozupone, C.; Maher, M.; Marotz, C.; Martin, B. D.; McDonald, D.; McIver, L. J.; Melnik, A. V.; Metcalf, J. L.; Morgan, S. C.; Morton, J. T.; Naimey, A. T.; Navas-Molina, J. A.; Nothias, L. F.; Orchanian, S. B.; Pearson, T.; Peoples, S. L.; Petras, D.; Preuss, M. L.; Pruesse, E.; Rasmussen, L. B.; Rivers, A.; Robeson, M. S.; Rosenthal, P.; Segata, N.; Shaffer, M.; Shiffer, A.; Sinha, R.; Song, S. J.; Spear, J. R.; Swafford, A. D.; Thompson, L. R.; Torres, P. J.; Trinh, P.; Tripathi, A.; Turnbaugh, P. J.; Ul-Hasan, S.; van der Hooft, J. J. J.; Vargas, F.; Vázquez-Baeza, Y.; Vogtmann, E.; von Hippel, M.; Walters, W.; Wan, Y.; Wang, M.; Warren, J.; Weber, K. C.; Williamson, C. H. D.; Willis, A. D.; Xu, Z. Z.; Zaneveld, J. R.; Zhang, Y.; Zhu, Q.; Knight, R.; Caporaso, J. G. Reproducible, interactive, scalable and extensible microbiome data science using QIIME 2. Nat. Biotechnol. 2019, 37 (8), 852–857.

(30) Kim, D. D.; Han, H.; Yun, T.; Song, M. J.; Terada, A.; Laureni, M.; Yoon, S. Identification of *nosZ*-expressing microorganisms consuming trace N_2_O in microaerobic chemostat consortia dominated by an uncultured *Burkholderiales*. ISME J. 2022, 16 (9), 2087–2098.

(31) Johnson, L. S.; Eddy, S. R.; Portugaly, E. Hidden Markov model speed heuristic and iterative HMM search procedure. BMC Bioinformatics 2010, 11 (1), 431.

(32) Yoon, S.; Carey, J. N.; Semrau, J. D. Feasibility of atmospheric methane removal using methanotrophic biotrickling filters. Appl. Microbiol. Biotechnol. 2009, 83 (5), 949–956.

(33) He, L.; Groom, J. D.; Wilson, E. H.; Fernandez, J.; Konopka, M. C.; Beck, D. A. C.; Lidstrom, M. E. A methanotrophic bacterium to enable methane removal for climate mitigation. Proc. Natl. Acad. Sci. 2023, 120 (35), e2310046120.

(34) Knief, C.; Lipski, A.; Dunfield, P. F. Diversity and activity of methanotrophic bacteria in different upland soils. Appl. Environ. Microbiol. 2003, 69 (11), 6703–6714.

(35) Knief, C.; Dunfield, P. F. Response and adaptation of different methanotrophic bacteria to low methane mixing ratios. Environ. Microbiol. 2005, 7 (9), 1307–1317.

(36) Dunfield, P. F.; Liesack, W.; Henckel, T.; Knowles, R.; Conrad, R. High-affinity methane oxidation by a soil enrichment culture containing a Type II methanotroph. Appl. Environ. Microbiol. 1999, 65 (3), 1009–1014.

(37) Sundstrom, E. R.; Criddle, C. S. Optimization of methanotrophic growth and production of poly(3-hydroxybutyrate) in a high-throughput microbioreactor system. Appl. Environ. Microbiol. 2015, 81 (14), 4767–4773.

(38) Pieja, A. J.; Sundstrom, E. R.; Criddle, C. S. Poly-3-hydroxybutyrate metabolism in the Type II methanotroph *Methylocystis parvus* OBBP. Appl. Environ. Microbiol. 2011, 77 (17), 6012–6019.

(39) Bordel, S.; Rojas, A.; Muñoz, R. Reconstruction of a genome scale metabolic model of the polyhydroxybutyrate producing methanotroph *Methylocystis parvus* OBBP. Microb. Cell Factories 2019, 18 (1), 104.

(40) Chen, Y.; Dumont, M. G.; Cébron, A.; Murrell, J. C. Identification of active methanotrophs in a landfill cover soil through detection of expression of 16S rRNA and functional genes. Environ. Microbiol. 2007, 9 (11), 2855–2869.

(41) Lee, S.-W.; Im, J.; DiSpirito, A. A.; Bodrossy, L.; Barcelona, M. J.; Semrau, J. D. Effect of nutrient and selective inhibitor amendments on methane oxidation, nitrous oxide production, and key gene presence and expression in landfill cover soils: characterization of the role of methanotrophs, nitrifiers, and denitrifiers. Appl. Microbiol. Biotechnol. 2009, 85 (2), 389–403.

(42) Inagaki, F.; Tsunogai, U.; Suzuki, M.; Kosaka, A.; Machiyama, H.; Takai, K.; Nunoura, T.; Nealson, K. H.; Horikoshi, K. Characterization of C1-metabolizing prokaryotic communities in methane seep habitats at the Kuroshima Knoll, southern Ryukyu Arc, by analyzing *pmoA*, *mmoX*, *mxaF*, *mcrA*, and 16S rRNA Genes. Appl. Environ. Microbiol. 2004, 70 (12), 7445–7455.

(43) Ul-Haque, M. F.; Kalidass, B.; Vorobev, A.; Baral, B. S.; DiSpirito, A. A.; Semrau, J. D. Methanobactin from *Methylocystis* sp. strain SB2 affects gene expression and methane monooxygenase activity in *Methylosinus trichosporium* OB3b. Appl. Environ. Microbiol. 2015, 81 (7), 2466–2473.

(44) Peng, P.; Gu, W.; DiSpirito, A. A.; Semrau, J. D. Multiple mechanisms for copper uptake by *Methylosinus trichosporium* OB3b in the presence of heterologous methanobactin. mBio 2022, 13 (5), e02239–02222.

(45) Kang-Yun, C. S.; Liang, X.; Dershwitz, P.; Gu, W.; Schepers, A.; Flatley, A.; Lichtmannegger, J.; Zischka, H.; Zhang, L.; Lu, X.; Gu, B.; Ledesma, J. C.; Pelger, D. J.; DiSpirito, A. A.; Semrau, J. D. Evidence for methanobactin “Theft” and novel chalkophore production in methanotrophs: impact on methanotrophic-mediated methylmercury degradation. ISME J. 2021, 16 (1), 211–220.

(46) Yun, J.; Crombie, A. T.; Ul Haque, M. F.; Cai, Y.; Zheng, X.; Wang, J.; Jia, Z.; Murrell, J. C.; Wang, Y.; Du, W. Revealing the community and metabolic potential of active methanotrophs by targeted metagenomics in the Zoige wetland of the Tibetan Plateau. Environ. Microbiol. 2021, 23 (11), 6520–6535.

(47) Venetz, J.; Żygadłowska, O. M.; Lenstra, W. K.; van Helmond, N. A. G. M.; Nuijten, G. H. L.; Wallenius, A. J.; Dalcin Martins, P.; Slomp, C. P.; Jetten, M. S. M.; Veraart, A. J. Versatile methanotrophs form an active methane biofilter in the oxycline of a seasonally stratified coastal basin. Environ. Microbiol. 2023, 25 (11), 2277–2288.

(48) Stolyar, S.; Costello, A. M.; Peeples, T. L.; Lidstrom, M. E. Role of multiple gene copies in particulate methane monooxygenase activity in the methane-oxidizing bacterium *Methylococcus capsulatus* Bath. Microbiology 1999, 145 (5), 1235–1244.

(49) Stein, L. Y.; Yoon, S.; Semrau, J. D.; DiSpirito, A. A.; Crombie, A.; Murrell, J. C.; Vuilleumier, S.; Kalyuzhnaya, M. G.; Camp, H. J. M. O. d.; Bringel, F.; Bruce, D.; Cheng, J.-F.; Copeland, A.; Goodwin, L.; Han, S.; Hauser, L.; Jetten, M. S. M.; Lajus, A.; Land, M. L.; Lapidus, A.; Lucas, S.; Médigue, C.; Pitluck, S.; Woyke, T.; Zeytun, A.; Klotz, M. G. Genome sequence of the obligate methanotroph *Methylosinus trichosporium* Strain OB3b. J. Bacteriol. 2010, 192 (24), 6497–6498.

(50) Gilbert, B.; McDonald, I. R.; Finch, R.; Stafford, G. P.; Nielsen, A. K.; Murrell, J. C. Molecular analysis of the *pmo* (particulate methane monooxygenase) operons from two Type II methanotrophs. Appl. Environ. Microbiol. 2000, 66 (3), 966–975.

(51) Dedysh, S. N.; Dunfield, P. F. Cultivation of Methanotrophs. In Hydrocarbon and Lipid Microbiology Protocols: Isolation and Cultivation, McGenity, T. J., Timmis, K. N., Nogales, B. Eds.; Springer Berlin Heidelberg, 2017; pp 231–247.

(52) He, R.; Wooller, M. J.; Pohlman, J. W.; Quensen, J.; Tiedje, J. M.; Leigh, M. B. Shifts in identity and activity of methanotrophs in arctic lake sediments in response to temperature changes. Appl. Environ. Microbiol. 2012, 78 (13), 4715–4723.

(53) Morris, S. A.; Radajewski, S.; Willison, T. W.; Murrell, J. C. Identification of the functionally active methanotroph population in a peat soil microcosm by stable-isotope probing. Appl. Environ. Microbiol. 2002, 68 (3), 1446–1453.

(54) Cébron, A.; Bodrossy, L.; Chen, Y.; Singer, A. C.; Thompson, I. P.; Prosser, J. I.; Murrell, J. C. Identity of active methanotrophs in landfill cover soil as revealed by DNA-stable isotope probing. FEMS Microbiol. Ecol. 2007, 62 (1), 12–23.

(55) Singleton, C. M.; McCalley, C. K.; Woodcroft, B. J.; Boyd, J. A.; Evans, P. N.; Hodgkins, S. B.; Chanton, J. P.; Frolking, S.; Crill, P. M.; Saleska, S. R.; Rich, V. I.; Tyson, G. W. Methanotrophy across a natural permafrost thaw environment. ISME J. 2018, 12 (10), 2544–2558.

(56) Farhan Ul Haque, M.; Crombie, A. T.; Murrell, J. C. Novel facultative *Methylocella* strains are active methane consumers at terrestrial natural gas seeps. Microbiome 2019, 7 (1), 134.

(57) Bordel, S.; Rodríguez, Y.; Hakobyan, A.; Rodríguez, E.; Lebrero, R.; Muñoz, R. Genome scale metabolic modeling reveals the metabolic potential of three Type II methanotrophs of the genus *Methylocystis*. Metab. Eng. 2019, 54, 191–199.

(58) Nguyen, A. D.; Lee, E. Y. Engineered methanotrophy: a sustainable solution for methane-based industrial biomanufacturing. Trends in Biotechnology 2021, 39 (4), 381–396.

(59) Einer, C.; Munk, D. E.; Park, E.; Akdogan, B.; Nagel, J.; Lichtmannegger, J.; Eberhagen, C.; Rieder, T.; Vendelbo, M. H.; Michalke, B.; Wimmer, R.; Blutke, A.; Feuchtinger, A.; Dershwitz, P.; DiSpirito, A. M.; Islam, T.; Castro, R. E.; Min, B.-K.; Kim, T.; Choi, S.; Kim, D.; Jung, C.; Lee, H.; Park, D.; Im, W.; Eun, S.-Y.; Cho, Y.-H.; Semrau, J. D.; Rodrigues, C. M. P.; Hohenester, S.; Damgaard Sandahl, T.; DiSpirito, A. A.; Zischka, H. ARBM101 (Methanobactin SB2) drains excess liver copper via biliary excretion in Wilson’s disease rats. Gastroenterology 2023, 165 (1), 187–200.e187.

