## Supplementray Information and Tables for "Selective enrichment of *Methylococcaceae* versus *Methylocystaceae* methanotrophs via control of methane feeding schemes": Supplementary table.pdf

### Table S1. Microbial community compositions organized at genus level

| Taxonomy | | | | | | Paddy Soi | 0.5%-$CH_4$ | Cu-free |
|---|---|---|---|---|---|---|---|---|
| Domain | Phylum | Class | Order | Family | Genus | PS | CB | R12 |
| Bacteria | Proteobact | Alphaprote | Rhizobiale | Beijerincki | Methylocy | 642 | 508 | 4916 |
| Bacteria | Proteobact | Gammapro | Methyloco | Methylomo | Methylomo | 42 | 38 | 555 |
| Bacteria | Proteobact | Alphaprote | Rhizobiale | Beijerincki | __ | 92 | 63 | 3659 |
| Bacteria | Bacteroido | Bacteroidia | Flavobacte | Flavobacte | Flavobacte | 380 | 0 | 0 |
| Bacteria | Proteobact | Gammapro | Methyloco | Methylomo | Methyloba | 0 | 15 | 0 |
| Bacteria | Proteobact | Gammapro | Burkholde | Rhodocycl | Methylove | 6 | 0 | 44 |
| Bacteria | Proteobact | Gammapro | Methyloco | Methyloco | Methyloma | 0 | 104 | 112 |
| Bacteria | Bacteroido | Bacteroidia | Chitinopha | Chitinopha | __ | 88 | 43 | 99 |
| Bacteria | Bacteroido | Bacteroidia | Flavobacte | Crociniton | Fluviicola | 142 | 0 | 46 |
| Bacteria | Bacteroido | Bacteroidia | Flavobacte | Flavobacte | __ | 36 | 0 | 0 |
| Bacteria | Proteobact | Alphaprote | Caulobacte | Caulobacte | Brevundim | 68 | 30 | 105 |
| Bacteria | Verrucomi | Chlamydia | Chlamydia | Parachlam | Neochlamy | 234 | 20 | 0 |
| Bacteria | Proteobact | Alphaprote | Sphingomo | Sphingomo | Novosphin | 54 | 0 | 333 |
| Bacteria | Proteobact | Alphaprote | Caulobacte | Caulobacte | __ | 67 | 741 | 283 |
| Bacteria | Bacteroido | Bacteroidia | Chitinopha | Chitinopha | Sediminiba | 38 | 34 | 0 |
| Bacteria | Bacteroido | Bacteroidia | Chitinopha | Chitinopha | Terrimonas | 34 | 165 | 155 |
| Bacteria | Proteobact | Gammapro | __ | __ | __ | 343 | 1013 | 8219 |
| Bacteria | Proteobact | Alphaprote | Rhizobiale | Rhizobiace | __ | 138 | 57 | 12 |
| Bacteria | Proteobact | Gammapro | Methyloco | Methyloco | Methyloca | 0 | 45 | 0 |
| Bacteria | Proteobact | Alphaprote | Sphingomo | Sphingomo | __ | 291 | 685 | 283 |
| Bacteria | Proteobact | Gammapro | Xanthomo | Xanthomo | Arenimona | 27 | 23 | 67 |
| Bacteria | Patescibac | Berkelbact | Berkelbact | Berkelbact | Berkelbact | 362 | 679 | 0 |
| Bacteria | Proteobact | Alphaprote | Rhizobiale | Devosiacea | Devosia | 17 | 194 | 192 |
| Bacteria | Verrucomi | Verrucomi | Opitutales | Opitutacea | Lacunispha | 298 | 548 | 0 |
| Bacteria | Proteobact | Alphaprote | Rhizobiale | Rhizobiace | Allorhizob | 0 | 0 | 0 |
| Bacteria | Verrucomi | Verrucomi | Verrucomi | Verrucomi | Prosthecob | 0 | 184 | 73 |
| Bacteria | Patescibac | ABY1 | Candidatus | Candidatus | Candidatus | 111 | 1759 | 0 |
| Bacteria | Bacteroido | Bacteroidia | Chitinopha | Chitinopha | Edaphobac | 43 | 54 | 338 |
| Bacteria | Proteobact | Alphaprote | Sphingomo | Sphingomo | Sphingopy | 0 | 0 | 0 |
| Bacteria | Proteobact | Alphaprote | Caulobacte | Caulobacte | Caulobacte | 39 | 285 | 0 |
| Bacteria | Proteobact | Alphaprote | Sphingomo | Sphingomo | Sphingomo | 823 | 398 | 0 |
| Bacteria | Patescibac | ABY1 | Candidatus | Candidatus | Candidatus | 140 | 246 | 0 |
| Bacteria | Proteobact | Gammapro | Xanthomo | Xanthomo | __ | 24 | 16 | 216 |
| Bacteria | Actinobact | Actinobact | Micrococc | Microbacte | __ | 122 | 20 | 8 |
| Bacteria | Patescibac | Saccharim | Saccharim | LWQ8 | LWQ8 | 202 | 137 | 0 |
| Bacteria | Bacteroido | Kryptonia | Kryptonial | BSV26 | BSV26 | 440 | 890 | 0 |
| Bacteria | Bacteroido | Bacteroidia | Chitinopha | Saprospira | uncultured | 109 | 63 | 0 |
| Bacteria | Proteobact | Alphaprote | Rhizobiale | Hyphomic | __ | 17 | 40 | 0 |
| Bacteria | Proteobact | Gammapro | Methyloco | Methylomo | __ | 134 | 236 | 0 |
| Bacteria | Proteobact | Alphaprote | Rickettsial | Mitochond | Mitochond | 33 | 8 | 2963 |
| Bacteria | Actinobact | Actinobact | Frankiales | Sporichthy | Candidatus | 59 | 79 | 0 |
| Bacteria | Chloroflex | KD4-96 | KD4-96 | KD4-96 | KD4-96 | 702 | 223 | 0 |
| Bacteria | Proteobact | Gammapro | Burkholde | Methyloph | Methyloter | 0 | 0 | 0 |
| Bacteria | Verrucomi | Chlamydia | Chlamydia | cvE6 | cvE6 | 215 | 329 | 0 |
| Bacteria | Patescibac | Parcubacte | Candidatus | Candidatus | Candidatus | 40 | 703 | 0 |
| Bacteria | Proteobact | Alphaprote | Rhizobiale | Hyphomic | Hyphomic | 91 | 155 | 241 |

| | | | | | | | | |
|---|---|---|---|---|---|---|---|---|
| Bacteria | Actinobact | Actinobact | Corynebac | Nocardiace | Rhodococc | 1031 | 0 | 0 |
| Bacteria | Proteobact | Alphaprote | Rhizobiale | Hyphomicr | uncultured | 114 | 80 | 0 |
| Bacteria | Patescibac | ABY1 | Candidatus | Candidatus | Candidatus | 74 | 39 | 0 |
| Bacteria | Verrucomi | Chlamydia | Chlamydia | Parachlam | __ | 401 | 30 | 0 |
| Bacteria | Actinobact | Actinobact | Frankiales | Sporichthy | __ | 196 | 251 | 0 |
| Bacteria | Bacteroido | Bacteroidia | Chitinopha | Chitinopha | Lacibacter | 0 | 0 | 22 |
| Bacteria | Acidobacte | Vicinamiba | Vicinamiba | uncultured | uncultured | 584 | 1773 | 0 |
| Bacteria | Actinobact | Acidimicro | Microtrich | Iamiaceae | Iamia | 253 | 0 | 0 |
| Bacteria | Actinobact | Actinobact | Micrococc | Microbacte | Agromyces | 0 | 0 | 0 |
| Bacteria | Proteobact | Alphaprote | Rhizobiale | Pleomorph | Chthonoba | 26 | 0 | 120 |
| Bacteria | Actinobact | Actinobact | Propioniba | Nocardioid | __ | 378 | 11 | 52 |
| Bacteria | Proteobact | Alphaprote | Rhizobiale | Beijerincki | Bosea | 0 | 0 | 233 |
| Bacteria | Proteobact | Alphaprote | Reyranella | Reyranella | Reyranella | 215 | 402 | 131 |
| Bacteria | Planctomy | Planctomy | Planctomy | Schlesneria | Planctopiru | 43 | 122 | 0 |
| Bacteria | Bacteroido | Bacteroidia | Chitinopha | Chitinopha | Ferruginiba | 36 | 0 | 0 |
| Bacteria | Proteobact | Gammapro | Burkholde | Comamona | __ | 21 | 287 | 80 |
| Bacteria | Verrucomi | Verrucomi | Pedosphae | Pedosphae | Pedosphae | 15 | 52 | 0 |
| Bacteria | Myxococc | Polyangia | Polyangial | Sandaracin | Sandaracin | 0 | 0 | 0 |
| Bacteria | Cyanobact | Cyanobact | Cyanobact | Chroococc | Chroococc | 0 | 0 | 995 |
| Bacteria | Verrucomi | Verrucomi | Opitutales | Opitutacea | __ | 26 | 488 | 0 |
| Bacteria | Proteobact | Gammapro | Diplorické | Diplorické | Aquicella | 458 | 594 | 0 |
| Bacteria | Proteobact | Alphaprote | Rhizobiale | Devosiacea | __ | 0 | 87 | 157 |
| Bacteria | Patescibac | Parcubacte | Parcubacte | Parcubacte | Parcubacte | 24 | 478 | 0 |
| Bacteria | Proteobact | Alphaprote | Rhizobiale | Rhizobiale | uncultured | 214 | 258 | 299 |
| Bacteria | Patescibac | Parcubacte | __ | __ | __ | 0 | 241 | 0 |
| Bacteria | Proteobact | Alphaprote | Rhodobact | Rhodobact | __ | 65 | 104 | 668 |
| Bacteria | Proteobact | Alphaprote | Rhizobiale | Xanthobac | Pseudoxan | 0 | 0 | 0 |
| Bacteria | Proteobact | Gammapro | Pseudomon | Pseudohon | BIyi10 | 4 | 190 | 0 |
| Bacteria | Proteobact | Alphaprote | Azospirilla | Azospirilla | Azospirillu | 0 | 0 | 587 |
| Bacteria | Proteobact | Alphaprote | Rhizobiale | Xanthobac | __ | 451 | 347 | 0 |
| Bacteria | Proteobact | Alphaprote | Caulobacte | Caulobacte | Asticcacau | 0 | 0 | 0 |
| Bacteria | Cyanobact | Vampirivib | Obscuribac | Obscuribac | Candidatus | 30 | 0 | 0 |
| Bacteria | Patescibac | Saccharim | Saccharim | Saccharim | Saccharim | 62 | 365 | 0 |
| Bacteria | Proteobact | Alphaprote | Micavibrio | uncultured | uncultured | 0 | 4 | 0 |
| Bacteria | Bacteroido | Bacteroidia | __ | __ | __ | 12 | 22 | 106 |
| Bacteria | Acidobacte | Acidobacte | Bryobacter | Bryobacter | Bryobacter | 142 | 23 | 0 |
| Bacteria | Dependent | Babeliae | Babeliales | __ | __ | 206 | 36 | 0 |
| Bacteria | Bdellovibr | Bdellovibr | Bdellovibr | Bdellovibr | Bdellovibr | 77 | 256 | 0 |
| Bacteria | Actinobact | Actinobact | Propioniba | Propioniba | Cutibacteri | 616 | 0 | 588 |
| Bacteria | Bacteroido | Bacteroidia | Sphingoba | NS11-12_r | NS11-12_r | 75 | 126 | 264 |
| Bacteria | Bacteroido | Bacteroidia | Chitinopha | Chitinopha | Flavisoliba | 0 | 23 | 0 |
| Bacteria | Bacteroido | Bacteroidia | Sphingoba | KD3-93 | KD3-93 | 0 | 755 | 0 |
| Bacteria | __ | __ | __ | __ | __ | 1622 | 999 | 337 |
| Bacteria | Proteobact | Alphaprote | Reyranella | Reyranella | uncultured | 89 | 44 | 0 |
| Bacteria | Bacteroido | Bacteroidia | Cytophaga | Cytophaga | Cytophaga | 0 | 0 | 634 |
| Bacteria | Myxococc | Polyangia | Polyangial | Polyangiac | Pajaroellob | 162 | 0 | 0 |
| Bacteria | Gemmatim | Gemmatim | Gemmatim | Gemmatim | uncultured | 607 | 584 | 0 |
| Bacteria | Actinobact | Actinobact | Corynebac | Mycobacte | Mycobacte | 751 | 240 | 0 |
| Bacteria | MBNT15 | MBNT15 | MBNT15 | MBNT15 | MBNT15 | 630 | 89 | 0 |
| Bacteria | Actinobact | Actinobact | Micrococc | Micrococc | __ | 444 | 19 | 0 |

| | | | | | | | | |
|---|---|---|---|---|---|---|---|---|
| Bacteria | Proteobact | Alphaprote | Rhizobiale | Methylolig | __ | 399 | 75 | 0 |
| Bacteria | Actinobact | Acidimicro | Microtrich | Ilumatobac | __ | 90 | 89 | 0 |
| Bacteria | Bacteroido | Bacteroidia | Sphingoba | AKYH767 | AKYH767 | 23 | 34 | 0 |
| Bacteria | Actinobact | Thermoleo | Solirubrob | 67-14 | 67-14 | 928 | 117 | 0 |
| Bacteria | Proteobact | Alphaprote | Rhizobiale | Xanthobac | Pseudolabr | 668 | 659 | 0 |
| Bacteria | Proteobact | Alphaprote | Sphingomo | Sphingomo | Sphingosin | 0 | 0 | 0 |
| Bacteria | Bacteroido | Bacteroidia | Sphingoba | env.OPS_1 | env.OPS_1 | 229 | 386 | 23 |
| Bacteria | Proteobact | Alphaprote | Caulobacte | Hyphomon | SWB02 | 0 | 23 | 0 |
| Bacteria | Bacteroido | Bacteroidia | Chitinopha | Saprospira | __ | 0 | 0 | 0 |
| Bacteria | Acidobacte | Acidobacte | Acidobacte | Acidobacte | Paludibacu | 182 | 0 | 0 |
| Bacteria | Actinobact | Actinobact | Propioniba | Nocardioid | Nocardioid | 910 | 37 | 0 |
| Bacteria | Actinobact | Acidimicro | Microtrich | Ilumatobac | CL500-29_ | 19 | 25 | 10 |
| Bacteria | Gemmatim | Gemmatim | Gemmatim | Gemmatim | __ | 14 | 53 | 0 |
| Bacteria | Actinobact | Actinobact | Micrococc | Intrasporan | __ | 396 | 0 | 0 |
| Bacteria | Proteobact | Gammapro | Methyloco | Methylomo | Methylomi | 0 | 27 | 0 |
| Bacteria | Acidobacte | Vicinamiba | Vicinamiba | Vicinamiba | Vicinamiba | 297 | 329 | 0 |
| Bacteria | Acidobacte | Thermoana | Thermoana | Thermoana | Subgroup_ | 663 | 179 | 0 |
| Bacteria | Myxococc | Polyangia | Polyangial | Phaselicys | Phaselicys | 79 | 0 | 0 |
| Bacteria | Proteobact | Alphaprote | Sphingomo | Sphingomo | Parablaston | 76 | 0 | 0 |
| Bacteria | Actinobact | Actinobact | Frankiales | Sporichthy | Longivirga | 156 | 0 | 0 |
| Bacteria | Proteobact | Alphaprote | __ | __ | __ | 198 | 139 | 0 |
| Bacteria | Patescibac | Microgeno | Candidatus | Candidatus | Candidatus | 262 | 48 | 0 |
| Bacteria | Proteobact | Alphaprote | Rhizobiale | Xanthobac | Bradyrhizo | 62 | 54 | 51 |
| Bacteria | Bacteroido | Bacteroidia | Chitinopha | Chitinopha | Dinghuiba | 0 | 0 | 0 |
| Bacteria | Proteobact | Gammapro | Burkholde | Comamona | Aquabacte | 0 | 195 | 0 |
| Bacteria | Desulfobac | Desulfobac | Desulfobac | Desulfobac | Desulfobac | 1518 | 213 | 0 |
| Bacteria | Myxococc | Polyangia | Nannocyst | Nannocyst | Nannocyst | 50 | 0 | 0 |
| Bacteria | Patescibac | __ | __ | __ | __ | 246 | 478 | 0 |
| Bacteria | Proteobact | Alphaprote | Rhizobiale | Pleomorph | uncultured | 0 | 0 | 0 |
| Bacteria | Patescibac | Parcubacte | Candidatus | Candidatus | Candidatus | 25 | 395 | 0 |
| Bacteria | Proteobact | Gammapro | Burkholde | Nitrosomo | GOUTA6 | 26 | 356 | 0 |
| Bacteria | Actinobact | Thermoleo | Gaiellales | Gaiellacea | Gaiella | 684 | 25 | 0 |
| Bacteria | Proteobact | Gammapro | Pseudomon | Pseudomon | Pseudomon | 835 | 75 | 26 |
| Bacteria | Proteobact | Alphaprote | Caulobacte | Hyphomon | Hirschia | 0 | 0 | 0 |
| Bacteria | Proteobact | Gammapro | Burkholde | Comamona | Hydrogeno | 0 | 0 | 0 |
| Bacteria | Firmicutes | Bacilli | Bacillales | Bacillacea | __ | 140 | 7 | 0 |
| Bacteria | Patescibac | Microgeno | Candidatus | Candidatus | Candidatus | 51 | 121 | 0 |
| Bacteria | Chloroflex | Gitt-GS-13 | Gitt-GS-13 | Gitt-GS-13 | Gitt-GS-13 | 83 | 15 | 0 |
| Bacteria | Bacteroido | Kapabacte | Kapabacte | Kapabacte | Kapabacte | 137 | 519 | 0 |
| Bacteria | Proteobact | Gammapro | Xanthomo | Rhodanoba | __ | 0 | 0 | 0 |
| Bacteria | Bdellovibr | Bdellovibr | Bdellovibr | Bdellovibr | OM27_cla | 92 | 0 | 0 |
| Bacteria | Proteobact | Gammapro | Methyloco | Methylomo | Methylosa | 0 | 0 | 0 |
| Bacteria | Gemmatim | Gemmatim | Gemmatim | Gemmatim | Gemmatim | 88 | 143 | 0 |
| Unassigned | __ | __ | __ | __ | __ | 493 | 90 | 864 |
| Bacteria | Myxococc | Polyangia | Haliangial | Haliangiac | Haliangium | 550 | 71 | 0 |
| Bacteria | Actinobact | Thermoleo | Gaiellales | uncultured | uncultured | 1540 | 417 | 0 |
| Bacteria | Proteobact | Alphaprote | Caulobacte | Caulobacte | Phenyloba | 132 | 370 | 0 |
| Bacteria | Patescibac | Gracilibact | Candidatus | Candidatus | Candidatus | 0 | 126 | 0 |
| Bacteria | Firmicutes | Negativicu | uncultured | uncultured | uncultured | 239 | 53 | 0 |
| Bacteria | Actinobact | Actinobact | Propioniba | Nocardioid | Aeromicro | 0 | 306 | 0 |

| | | | | | | | | |
|---|---|---|---|---|---|---|---|---|
| Bacteria | Actinobact | Thermoleo | Gaiellales | __ | __ | 297 | 93 | 0 |
| Bacteria | Armatimor | Fimbriimo | Fimbriimo | Fimbriimo | Fimbriimo | 119 | 33 | 0 |
| Bacteria | Proteobact | Alphaprote | Rhizobiale | Xanthobac | uncultured | 189 | 241 | 0 |
| Bacteria | Patescibac | Microgeno | Candidatus | Candidatus | Candidatus | 253 | 112 | 0 |
| Bacteria | Proteobact | Gammapro | Burkholde | Comamona | Ramlibacte | 0 | 0 | 0 |
| Bacteria | Proteobact | Gammapro | Steroidoba | Steroidoba | uncultured | 209 | 62 | 0 |
| Bacteria | Actinobact | Acidimicro | Microtrich | __ | __ | 309 | 50 | 0 |
| Bacteria | Proteobact | Gammapro | Burkholde | Methyloph | __ | 0 | 12 | 117 |
| Bacteria | Myxococc | Polyangia | mle1-27 | mle1-27 | mle1-27 | 64 | 0 | 0 |
| Bacteria | Firmicutes | Bacilli | Staphyloco | Staphyloco | Staphyloco | 238 | 0 | 192 |
| Bacteria | Gemmatim | S0134_ter | S0134_ter | S0134_ter | S0134_ter | 6 | 327 | 0 |
| Bacteria | Bacteroido | Bacteroidi | Cytophaga | Cyclobacte | Algoriphag | 0 | 0 | 0 |
| Bacteria | Proteobact | Gammapro | Burkholde | Methyloph | Methyloph | 0 | 0 | 0 |
| Bacteria | Actinobact | Acidimicro | IMCC2625 | IMCC2625 | IMCC2625 | 613 | 73 | 0 |
| Bacteria | Proteobact | Alphaprote | Caulobacte | Caulobacte | uncultured | 106 | 68 | 0 |
| Bacteria | Patescibac | Microgeno | Candidatus | Candidatus | Candidatus | 34 | 0 | 0 |
| Bacteria | Patescibac | Microgeno | Candidatus | Candidatus | Candidatus | 211 | 183 | 0 |
| Bacteria | Actinobact | Actinobact | Micrococc | Intrasporar | Oryzihumu | 303 | 15 | 0 |
| Bacteria | Proteobact | Gammapro | Legionella | Legionella | Legionella | 665 | 269 | 0 |
| Bacteria | Firmicutes | Bacilli | Bacillales | Planococca | __ | 11 | 0 | 0 |
| Bacteria | Latescibac | Latescibac | Latescibac | Latescibac | Latescibac | 335 | 0 | 0 |
| Bacteria | Planctomy | Phycisphae | Phycisphae | Phycisphae | SM1A02 | 0 | 75 | 0 |
| Bacteria | Proteobact | Alphaprote | Rickettsial | __ | __ | 0 | 0 | 0 |
| Bacteria | Actinobact | Acidimicro | __ | __ | __ | 209 | 20 | 0 |
| Bacteria | Proteobact | Gammapro | Coxiellales | Coxiellace | Coxiella | 334 | 97 | 0 |
| Bacteria | Nitrospirot | Nitrospiria | Nitrospiral | Nitrospirac | Nitrospira | 408 | 16 | 0 |
| Bacteria | Actinobact | MB-A2-10 | MB-A2-10 | MB-A2-10 | MB-A2-10 | 831 | 69 | 0 |
| Bacteria | Bacteroido | Bacteroidi | Chitinopha | Chitinopha | Pseudoflav | 0 | 0 | 0 |
| Bacteria | Patescibac | Saccharim | Saccharim | __ | __ | 66 | 298 | 0 |
| Bacteria | Firmicutes | Bacilli | Thermoact | Thermoact | Thermoact | 95 | 11 | 0 |
| Bacteria | Chloroflex | OLB14 | OLB14 | OLB14 | OLB14 | 51 | 10 | 0 |
| Bacteria | Chloroflex | Anaeroline | SBR1031 | A4b | A4b | 25 | 0 | 0 |
| Bacteria | Proteobact | Gammapro | Enterobact | Enterobact | __ | 60 | 5 | 50 |
| Bacteria | Actinobact | Coriobacte | OPB41 | OPB41 | OPB41 | 1312 | 151 | 0 |
| Bacteria | Proteobact | Alphaprote | Rhizobiale | Beijerincki | alphaI_clu | 110 | 0 | 0 |
| Bacteria | Acidobacte | Holophaga | Holophaga | Holophaga | Geothrix | 0 | 26 | 0 |
| Bacteria | Proteobact | Gammapro | Methyloco | Methyloco | __ | 0 | 21 | 0 |
| Bacteria | Planctomy | Planctomy | Planctomy | Schlesneri | Schlesneri | 67 | 64 | 0 |
| Bacteria | Verrucomi | Chlamydia | Chlamydia | Parachlam | Candidatus | 203 | 45 | 0 |
| Bacteria | Patescibac | Microgeno | Microgeno | Microgeno | Microgeno | 119 | 7 | 0 |
| Bacteria | Planctomy | Planctomy | Gemmatale | Gemmatac | uncultured | 84 | 61 | 0 |
| Bacteria | Actinobact | Actinobact | PeM15 | PeM15 | PeM15 | 63 | 109 | 0 |
| Bacteria | Bacteroido | Bacteroidi | Chitinopha | Chitinopha | Flavihumib | 0 | 0 | 0 |
| Bacteria | Proteobact | Alphaprote | Rhodospiri | uncultured | uncultured | 14 | 85 | 0 |
| Bacteria | Acidobacte | Vicinamib | Subgroup_ | Subgroup_ | Subgroup_ | 440 | 525 | 0 |
| Bacteria | Verrucomi | Verrucomi | Pedosphae | Pedosphae | __ | 147 | 59 | 0 |
| Bacteria | Proteobact | Gammapro | KF-JG30-( | KF-JG30-( | KF-JG30-( | 0 | 43 | 0 |
| Bacteria | Actinobact | Actinobact | Frankiales | Frankiales | Frankiales | 176 | 0 | 0 |
| Bacteria | Nitrospirot | Thermodes | uncultured | uncultured | uncultured | 1074 | 730 | 0 |
| Bacteria | Chloroflex | Anaeroline | __ | __ | __ | 136 | 45 | 0 |

| | | | | | | | | |
|---|---|---|---|---|---|---|---|---|
| Bacteria | Bacteroido | Bacteroidia | Cytophaga | Microscilla | __ | 182 | 0 | 0 |
| Bacteria | Bacteroido | Bacteroidia | Sphingoba | LiUU-11-1 | LiUU-11-1 | 23 | 272 | 0 |
| Bacteria | Firmicutes | Bacilli | Bacillales | Bacillacea | Bacillus | 54 | 0 | 166 |
| Bacteria | Firmicutes | Clostridia | Clostridial | Clostridiac | Clostridium | 321 | 73 | 0 |
| Bacteria | Acidobacte | Blastocatel | Blastocatel | Blastocatel | __ | 119 | 26 | 0 |
| Bacteria | Firmicutes | Bacilli | Lactobacil | Lactobacil | Leuconosto | 0 | 0 | 0 |
| Bacteria | Actinobact | Actinobact | Propioniba | Nocardioid | Marmorico | 204 | 91 | 0 |
| Bacteria | Proteobact | Gammapro | Burkholder | Nitrosomo | Ellin6067 | 27 | 363 | 0 |
| Bacteria | Bacteroido | Bacteroidia | Chitinopha | Chitinopha | Flavitalea | 0 | 0 | 0 |
| Bacteria | Chloroflex | Anaeroline | Caldilineal | Caldilineac | uncultured | 16 | 0 | 0 |
| Bacteria | Proteobact | Alphaprote | Holosporal | Holospora | uncultured | 405 | 154 | 0 |
| Bacteria | Armatimor | uncultured | uncultured | uncultured | uncultured | 207 | 152 | 0 |
| Bacteria | Myxococc | Polyangia | Blfdi19 | Blfdi19 | Blfdi19 | 146 | 0 | 0 |
| Bacteria | Actinobact | Actinobact | Micrococc | Intrasporan | Intrasporan | 43 | 0 | 0 |
| Bacteria | Armatimor | Fimbriimo | Fimbriimo | Fimbriimo | __ | 0 | 0 | 0 |
| Bacteria | Planctomy | Planctomy | Planctomy | Rubinispha | SH-PL14 | 0 | 0 | 0 |
| Bacteria | Actinobact | Thermoleo | Solirubrob | Solirubrob | Conexibac | 287 | 0 | 0 |
| Bacteria | Myxococc | Polyangia | Polyangial | Polyangiac | __ | 196 | 0 | 0 |
| Bacteria | Cyanobact | Cyanobact | Chloroplas | Chloroplas | Chloroplas | 313 | 0 | 90 |
| Bacteria | Proteobact | Alphaprote | Rhodobact | Rhodobact | Paracoccus | 51 | 0 | 0 |
| Bacteria | Proteobact | Gammapro | Methyloco | Methylomo | Methyloglo | 0 | 217 | 0 |
| Bacteria | Desulfobac | Desulfobac | Desulfobac | Desulfosar | Desulfatirh | 0 | 17 | 0 |
| Bacteria | Acidobacte | Holophaga | Subgroup_ | Subgroup_ | Subgroup_ | 509 | 636 | 0 |
| Bacteria | Actinobact | Actinobact | Corynebac | Nocardiace | Skermania | 0 | 0 | 188 |
| Bacteria | Bacteroido | Bacteroidia | Cytophaga | __ | __ | 0 | 0 | 346 |
| Bacteria | Planctomy | Planctomy | Pirellulales | Pirellulace | Pirellula | 8 | 15 | 0 |
| Bacteria | Proteobact | Gammapro | Methyloco | Methylomo | Methylovu | 0 | 149 | 0 |
| Bacteria | Proteobact | Alphaprote | Rhizobiale | Xanthobac | Ancylobac | 0 | 0 | 0 |
| Bacteria | Bacteroido | Bacteroidia | Chitinopha | Chitinopha | Aurantisol | 88 | 6 | 0 |
| Bacteria | Proteobact | Gammapro | Methyloco | Methyloco | Methylopa | 0 | 0 | 0 |
| Bacteria | Proteobact | Alphaprote | Rickettsial | SM2D12 | SM2D12 | 2 | 0 | 0 |
| Bacteria | Chloroflex | Chloroflex | Thermomic | JG30-KF-C | JG30-KF-C | 38 | 0 | 0 |
| Bacteria | Actinobact | Actinobact | Frankiales | Frankiacea | Jatrophihal | 124 | 0 | 0 |
| Bacteria | Sva0485 | Sva0485 | Sva0485 | Sva0485 | Sva0485 | 418 | 0 | 0 |
| Bacteria | Proteobact | Gammapro | Burkholder | Comamona | Polaromon | 0 | 7 | 0 |
| Bacteria | Desulfobac | Syntrophia | Syntrophal | uncultured | uncultured | 259 | 17 | 0 |
| Bacteria | Chloroflex | Anaeroline | SJA-15 | SJA-15 | SJA-15 | 541 | 0 | 0 |
| Bacteria | Elusimicro | Lineage_II | Lineage_II | Lineage_II | Lineage_II | 0 | 7 | 0 |
| Bacteria | Proteobact | Alphaprote | Rhizobiale | Pleomorph | Prosthecon | 0 | 0 | 0 |
| Bacteria | Proteobact | Alphaprote | Micropeps | Micropeps | __ | 78 | 0 | 0 |
| Bacteria | Verrucomi | Chlamydia | Chlamydia | Simkaniac | __ | 114 | 0 | 0 |
| Bacteria | Proteobact | Gammapro | Burkholder | Hydrogeno | Thiobacillu | 75 | 33 | 0 |
| Bacteria | Proteobact | Alphaprote | Sphingomo | Sphingomo | Porphyroba | 0 | 0 | 0 |
| Bacteria | Proteobact | Alphaprote | Rhizobiale | Beijerincki | Methyloba | 90 | 0 | 0 |
| Bacteria | Planctomy | Planctomy | Isosphaera | Isosphaera | Paludispha | 0 | 0 | 0 |
| Bacteria | Acidobacte | Vicinamiba | Vicinamiba | __ | __ | 739 | 194 | 0 |
| Bacteria | Acidobacte | Aminicena | Aminicena | Aminicena | Aminicena | 94 | 157 | 0 |
| Bacteria | Desulfobac | Desulfuron | Geobactera | Geobactera | __ | 733 | 180 | 0 |
| Bacteria | Proteobact | Gammapro | Salinispha | Solimonad | __ | 105 | 0 | 0 |
| Bacteria | Desulfobac | Desulfomo | Desulfomo | Desulfomo | Desulfomo | 48 | 374 | 0 |

| | | | | | | | | |
|---|---|---|---|---|---|---|---|---|
| Bacteria | Proteobact | Gammapro | Burkholde | Oxalobacte | __ | 24 | 5 | 61 |
| Bacteria | Desulfobac | Desulfobul | Desulfobul | Desulfocap | __ | 17 | 9 | 0 |
| Bacteria | Myxococc | Myxococc | Myxococc | Myxococc | uncultured | 153 | 0 | 0 |
| Bacteria | Bdellovibr | Oligoflexia | 0319-6G20 | 0319-6G20 | 0319-6G20 | 138 | 140 | 0 |
| Bacteria | Firmicutes | Clostridia | Clostridial | Clostridiac | __ | 217 | 0 | 337 |
| Bacteria | Proteobact | Gammapro | CCM19a | CCM19a | CCM19a | 84 | 0 | 0 |
| Bacteria | Desulfobac | Desulfobac | Desulfobac | Desulfosar | Sva0081_s | 147 | 0 | 0 |
| Bacteria | Proteobact | Alphaprote | Rhizobiale | Rhizobiale | Bauldia | 13 | 43 | 0 |
| Bacteria | Myxococc | Myxococc | Myxococc | Myxococc | P3OB-42 | 40 | 0 | 0 |
| Bacteria | Actinobact | Thermoleo | Solirubrob | Solirubrob | __ | 603 | 0 | 0 |
| Bacteria | Proteobact | Gammapro | Acidiferrol | Acidiferrol | Sulfurifust | 85 | 0 | 0 |
| Bacteria | Proteobact | Alphaprote | Rhizobiale | Xanthobac | Pseudorhoo | 65 | 37 | 0 |
| Bacteria | Myxococc | Myxococc | Myxococc | __ | __ | 158 | 0 | 0 |
| Bacteria | Patescibac | Parcubacte | Candidatus | Candidatus | Candidatus | 104 | 478 | 0 |
| Bacteria | Proteobact | Alphaprote | Rhodospiri | Magnetosp | Magnetosp | 0 | 0 | 0 |
| Bacteria | Patescibac | Microgeno | Candidatus | Candidatus | Candidatus | 805 | 277 | 0 |
| Bacteria | Firmicutes | Bacilli | Bacillales | __ | __ | 137 | 0 | 0 |
| Bacteria | Myxococc | Polyangia | Polyangial | Sandaracin | uncultured | 367 | 153 | 0 |
| Bacteria | Proteobact | Gammapro | Pseudomon | Moraxella | Enhydroba | 19 | 0 | 108 |
| Bacteria | Patescibac | Parcubacte | Candidatus | Candidatus | Candidatus | 6 | 0 | 0 |
| Bacteria | Patescibac | Dojkabacte | Dojkabacte | Dojkabacte | Dojkabacte | 81 | 61 | 0 |
| Bacteria | Bdellovibr | __ | __ | __ | __ | 0 | 122 | 0 |
| Bacteria | Proteobact | Alphaprote | Rhizobiale | Xanthobac | Rhodopseu | 48 | 0 | 0 |
| Bacteria | Proteobact | Gammapro | Burkholde | Comamona | Piscinibact | 0 | 0 | 0 |
| Bacteria | Proteobact | Gammapro | Burkholde | Oxalobacte | Massilia | 0 | 0 | 0 |
| Bacteria | Myxococc | Myxococc | Myxococc | Anaeromyx | Anaeromyx | 372 | 15 | 0 |
| Bacteria | Actinobact | Actinobact | Corynebac | Corynebac | __ | 71 | 0 | 0 |
| Bacteria | Verrucomi | Omnitroph | Omnitroph | Omnitroph | Omnitroph | 82 | 4 | 0 |
| Bacteria | Patescibac | ABY1 | Candidatus | Candidatus | Candidatus | 144 | 198 | 0 |
| Bacteria | Planctomy | Planctomy | Planctomy | uncultured | uncultured | 152 | 141 | 0 |
| Bacteria | Firmicutes | Clostridia | Clostridial | Clostridiac | Clostridiur | 456 | 53 | 0 |
| Bacteria | Desulfobac | __ | __ | __ | __ | 112 | 0 | 0 |
| Bacteria | Proteobact | Alphaprote | Rhizobiale | __ | __ | 0 | 34 | 0 |
| Bacteria | Actinobact | Actinobact | Micrococc | Microbacte | Microbacte | 25 | 0 | 111 |
| Bacteria | Actinobact | Actinobact | Micrococc | __ | __ | 68 | 26 | 0 |
| Bacteria | Acidobacte | Blastocatel | Blastocatel | Blastocatel | uncultured | 0 | 0 | 110 |
| Bacteria | Bacteroido | Bacteroidia | Chitinopha | Chitinopha | Taibaiella | 0 | 0 | 0 |
| Bacteria | Proteobact | Gammapro | CCD24 | CCD24 | CCD24 | 50 | 0 | 0 |
| Bacteria | Firmicutes | Clostridia | Peptostrep | Anaerovor | [Eubacteri | 20 | 15 | 0 |
| Bacteria | Actinobact | Actinobact | Micrococc | Microbacte | Cryobacter | 152 | 0 | 0 |
| Bacteria | Proteobact | Alphaprote | Acetobacte | Acetobacte | Rhodovast | 132 | 0 | 0 |
| Bacteria | Proteobact | Gammapro | Burkholde | __ | __ | 32 | 152 | 0 |
| Bacteria | Verrucomi | Verrucomi | Opitutales | Opitutacea | Opitutus | 0 | 0 | 0 |
| Bacteria | Bacteroido | Bacteroidia | Sphingoba | Sphingoba | __ | 52 | 0 | 0 |
| Bacteria | Patescibac | Gracilibact | Candidatus | Candidatus | Candidatus | 12 | 136 | 0 |
| Bacteria | Actinobact | Actinobact | Micrococc | Intrasporar | Terrabacte | 103 | 0 | 0 |
| Bacteria | Chloroflex | Anaeroline | SBR1031 | __ | __ | 75 | 52 | 0 |
| Bacteria | Proteobact | Alphaprote | Rhodobact | Rhodobact | Rhodobact | 0 | 0 | 0 |
| Bacteria | Firmicutes | Clostridia | Clostridial | Caloramato | Fonticella | 29 | 0 | 0 |
| Bacteria | Bacteroido | Bacteroidia | Chitinopha | 37-13 | 37-13 | 0 | 74 | 0 |

| | | | | | | | | |
|---|---|---|---|---|---|---|---|---|
| Bacteria | Verrucomi | Chlamydia | Chlamydia | __ | __ | 149 | 92 | 0 |
| Bacteria | Methylomi | Methylomi | Methylomi | Methylomi | Sh765B-Tz | 252 | 0 | 0 |
| Bacteria | Actinobact | Coriobacte | FS118-23E | FS118-23E | FS118-23E | 103 | 38 | 0 |
| Bacteria | Hydrogene | Hydrogene | Hydrogene | Hydrogene | Hydrogene | 23 | 0 | 0 |
| Bacteria | Actinobact | Actinobact | Micrococc | Demequina | Demequina | 0 | 24 | 0 |
| Bacteria | Bacteroido | Bacteroidia | Flavobacte | Crociniton | __ | 0 | 38 | 0 |
| Bacteria | Bacteroido | Bacteroidia | Bacteroida | Prolixibact | Roseimarin | 0 | 146 | 0 |
| Bacteria | Bacteroido | Bacteroidia | Cytophaga | Hymenoba | Hymenoba | 76 | 0 | 96 |
| Bacteria | Bacteroido | Ignavibacte | Ignavibacte | SR-FBR-L | SR-FBR-L | 0 | 348 | 0 |
| Bacteria | Firmicutes | Desulfitoba | Desulfitoba | Desulfitoba | Desulfospo | 30 | 13 | 0 |
| Bacteria | Acidobacte | Blastocatel | Blastocatel | Blastocatel | JGI_00010 | 0 | 124 | 0 |
| Bacteria | Actinobact | Actinobact | Kineospori | Kineospori | __ | 76 | 0 | 0 |
| Bacteria | Firmicutes | Clostridia | Clostridial | Clostridiac | Clostridium | 335 | 35 | 0 |
| Bacteria | Bacteroido | Ignavibacte | Ignavibacte | PHOS-HE3 | PHOS-HE3 | 0 | 19 | 0 |
| Bacteria | Bacteroido | Bacteroidia | Cytophaga | Microscilla | uncultured | 27 | 0 | 0 |
| Bacteria | Patescibac | Microgeno | Candidatus | Candidatus | Candidatus | 202 | 71 | 0 |
| Bacteria | Actinobact | Acidimicro | Microtrich | uncultured | uncultured | 61 | 0 | 0 |
| Bacteria | Patescibac | ABY1 | __ | __ | __ | 149 | 221 | 0 |
| Bacteria | Proteobact | Gammapro | Diplorické | Diplorické | uncultured | 75 | 355 | 0 |
| Bacteria | Proteobact | Alphaprote | Micropeps | Micropeps | uncultured | 0 | 48 | 0 |
| Bacteria | Firmicutes | Bacilli | Thermoact | Thermoact | Planifilum | 149 | 0 | 0 |
| Bacteria | Firmicutes | Clostridia | Clostridial | Clostridiac | Clostridium | 144 | 0 | 0 |
| Bacteria | Desulfobac | Syntrophol | Syntrophol | Syntrophol | uncultured | 86 | 29 | 0 |
| Bacteria | Firmicutes | Bacilli | Lactobacil | Carnobacte | __ | 68 | 0 | 0 |
| Bacteria | Firmicutes | Bacilli | Lactobacil | Lactobacil | Lactiplanti | 0 | 0 | 102 |
| Bacteria | Bacteroido | Bacteroidia | Cytophaga | Microscilla | Ohtaekwan | 0 | 0 | 0 |
| Bacteria | Bdellovibr | Bdellovibr | Bacteriovo | Bacteriovo | Peredibacte | 0 | 0 | 0 |
| Bacteria | Bacteroido | Bacteroidia | Sphingoba | Sphingoba | Pedobacter | 227 | 0 | 0 |
| Bacteria | Actinobact | Actinobact | Frankiales | __ | __ | 73 | 5 | 0 |
| Bacteria | Chloroflex | Anaeroline | Anaeroline | Anaeroline | UTCFX1 | 123 | 30 | 0 |
| Bacteria | Acidobacte | Blastocatel | DS-100 | DS-100 | DS-100 | 24 | 25 | 0 |
| Bacteria | Proteobact | Alphaprote | Rhizobiale | Kaistiacea | Kaistia | 0 | 0 | 0 |
| Bacteria | Actinobact | Thermoleo | Solirubrob | Solirubrob | uncultured | 0 | 94 | 0 |
| Bacteria | Dependent | Babeliae | Babeliales | Vermiphila | Vermiphila | 137 | 36 | 0 |
| Bacteria | Verrucomi | Chlamydia | Chlamydia | Simkaniac | uncultured | 36 | 25 | 0 |
| Bacteria | Patescibac | Microgeno | Candidatus | Candidatus | Candidatus | 101 | 13 | 0 |
| Bacteria | Acidobacte | Holophaga | Holophaga | Holophaga | __ | 4 | 37 | 0 |
| Bacteria | Chloroflex | Anaeroline | RBG-13-54 | RBG-13-54 | RBG-13-54 | 585 | 21 | 0 |
| Bacteria | Patescibac | WWE3 | WWE3 | WWE3 | WWE3 | 227 | 115 | 0 |
| Bacteria | Actinobact | __ | __ | __ | __ | 298 | 5 | 0 |
| Bacteria | Patescibac | Parcubacte | GWA2-38- | GWA2-38- | GWA2-38- | 127 | 0 | 0 |
| Bacteria | Myxococc | Polyangia | Polyangial | Polyangiac | Aetherobac | 100 | 0 | 0 |
| Bacteria | Desulfobac | uncultured | uncultured | uncultured | uncultured | 105 | 0 | 0 |
| Bacteria | Proteobact | Alphaprote | Acetobacte | Acetobacte | __ | 122 | 28 | 0 |
| Bacteria | Bacteroido | Bacteroidia | Chitinopha | Chitinopha | Parasegetib | 0 | 0 | 0 |
| Eukaryota | Cnidaria | Myxozoa | Bivalvulida | Bivalvulida | Bivalvulida | 2 | 0 | 0 |
| Bacteria | Myxococc | Polyangia | Polyangial | BIrii41 | BIrii41 | 20 | 34 | 0 |
| Bacteria | Bacteroido | Ignavibacte | Ignavibacte | __ | __ | 81 | 98 | 0 |
| Bacteria | Proteobact | Alphaprote | uncultured | uncultured | uncultured | 22 | 0 | 0 |
| Bacteria | Proteobact | Gammapro | JG36-GS-5 | JG36-GS-5 | JG36-GS-5 | 21 | 0 | 0 |

| | | | | | | | | |
|---|---|---|---|---|---|---|---|---|
| Bacteria | Chloroflex | Anaeroline | Anaeroline | Anaeroline | __ | 522 | 86 | 0 |
| Bacteria | Bacteroido | Bacteroidia | Cytophaga | Microscilla | Hassallia | 58 | 0 | 0 |
| Bacteria | Actinobact | Actinobact | Micrococc | Intrasporar | Humibacill | 51 | 0 | 0 |
| Bacteria | Deinococc | Deinococc | Deinococc | Trueperace | Truepera | 0 | 0 | 0 |
| Bacteria | Bacteroido | Bacteroidia | Sphingoba | __ | __ | 100 | 156 | 0 |
| Bacteria | Proteobact | Gammapro | Pseudomo | Pseudomo | __ | 35 | 0 | 57 |
| Bacteria | Desulfoba | Desulfuror | Geobactera | Geobactera | uncultured | 101 | 0 | 0 |
| Bacteria | Desulfoba | Syntropho | Syntropho | Syntropho | Syntropho | 247 | 59 | 0 |
| Bacteria | Actinobact | Actinobact | Micromon | Micromon | __ | 36 | 0 | 0 |
| Bacteria | Planctomy | Planctomy | Isosphaera | Isosphaera | __ | 13 | 16 | 0 |
| Bacteria | Patescibac | Microgeno | Candidatus | Candidatus | Candidatus | 85 | 79 | 0 |
| Bacteria | Bacteroido | Bacteroidia | Cytophaga | Microscilla | OLB12 | 0 | 23 | 0 |
| Bacteria | Bacteroido | Kryptonia | Kryptonial | MSB-3C8 | MSB-3C8 | 0 | 0 | 0 |
| Bacteria | Actinobact | Actinobact | Frankiales | uncultured | uncultured | 243 | 0 | 0 |
| Bacteria | FCPU426 | FCPU426 | FCPU426 | FCPU426 | FCPU426 | 72 | 0 | 0 |
| Bacteria | Elusimicro | Elusimicro | Lineage_IV | Lineage_IV | Lineage_IV | 102 | 60 | 0 |
| Bacteria | Gemmatim | BD2-11_te | BD2-11_te | BD2-11_te | BD2-11_te | 0 | 103 | 0 |
| Bacteria | Chloroflex | Anaeroline | Anaeroline | Anaeroline | uncultured | 354 | 159 | 0 |
| Bacteria | Proteobact | Alphaprote | Rhizobiale | Pleomorph | Pleomorph | 0 | 0 | 0 |
| Bacteria | Firmicutes | Clostridia | Clostridia | Gracilibact | Lutispora | 103 | 0 | 0 |
| Bacteria | Acidobacte | Acidobacte | Acidobacte | uncultured | uncultured | 88 | 0 | 0 |
| Bacteria | Proteobact | Gammapro | Burkholde | Comamona | Leptothrix | 0 | 0 | 0 |
| Bacteria | Verrucomi | Verrucomi | Pedosphae | Pedosphae | Oikopleura | 10 | 0 | 0 |
| Bacteria | Planctomy | Phycisphae | Tepidispha | WD2101_s | WD2101_s | 209 | 10 | 0 |
| Bacteria | Actinobact | Actinobact | Corynebac | Corynebac | Corynebac | 32 | 0 | 69 |
| Bacteria | Nitrospino | P9X2b3D0 | P9X2b3D0 | P9X2b3D0 | P9X2b3D0 | 96 | 212 | 0 |
| Bacteria | Dependent | Babeliae | Babeliales | Babeliacea | Babeliacea | 0 | 0 | 0 |
| Bacteria | Acidobacte | Acidobacte | Solibactera | Solibactera | Candidatus | 132 | 0 | 0 |
| Bacteria | Actinobact | Actinobact | Streptomy | Streptomy | Streptomy | 25 | 55 | 0 |
| Bacteria | Verrucomi | Verrucomi | S-BQ2-57_ | S-BQ2-57_ | S-BQ2-57_ | 36 | 0 | 0 |
| Bacteria | Firmicutes | Clostridia | Clostridial | Clostridiac | Clostridiur | 43 | 0 | 0 |
| Bacteria | Patescibac | Microgeno | __ | __ | __ | 96 | 61 | 0 |
| Bacteria | Verrucomi | Chlamydia | Chlamydia | Parachlam | uncultured | 54 | 0 | 0 |
| Bacteria | Halanaerol | Halanaerol | Halanaerol | Halanaerol | Halocella | 25 | 0 | 0 |
| Archaea | Crenarcha | Nitrososph | SCGC_AB | SCGC_AB | SCGC_AB | 0 | 0 | 67 |
| Bacteria | Bacteroido | Bacteroidia | Flavobacte | Weeksella | Chryseoba | 0 | 0 | 7 |
| Bacteria | Acidobacte | Subgroup_ | Subgroup_ | Subgroup_ | Subgroup_ | 300 | 0 | 0 |
| Bacteria | Firmicutes | Clostridia | Clostridial | Clostridiac | Clostridiur | 95 | 0 | 0 |
| Bacteria | Chloroflex | TK10 | TK10 | TK10 | TK10 | 122 | 17 | 0 |
| Bacteria | Bacteroido | Bacteroidia | Sphingoba | Lentimicro | Lentimicro | 51 | 63 | 0 |
| Bacteria | Proteobact | Gammapro | Pseudomo | Moraxella | Cavicella | 0 | 66 | 0 |
| Bacteria | Actinobact | Actinobact | Micrococc | Microbacte | Yonghapar | 0 | 0 | 0 |
| Bacteria | Firmicutes | Bacilli | Bacillales | Bacillacea | Ureibacillu | 65 | 0 | 0 |
| Bacteria | Proteobact | Alphaprote | Azospirilla | Azospirilla | Skermanel | 65 | 0 | 0 |
| Bacteria | Actinobact | Actinobact | Frankiales | Nakamurel | Nakamurel | 65 | 0 | 0 |
| Bacteria | Actinobact | Actinobact | Frankiales | Sporichthy | uncultured | 20 | 0 | 0 |
| Bacteria | Proteobact | Alphaprote | Acetobacte | Acetobacte | Roseomon | 134 | 30 | 0 |
| Bacteria | Proteobact | Gammapro | Steroidoba | Steroidoba | __ | 25 | 39 | 0 |
| Bacteria | Proteobact | Gammapro | Burkholde | Nitrosomo | MND1 | 23 | 36 | 0 |
| Bacteria | Actinobact | Acidimicro | Microtrich | Microtrich | IMCC2620 | 0 | 0 | 0 |

| | | | | | | | | |
|---|---|---|---|---|---|---|---|---|
| Bacteria | Bacteroido | Bacteroidia | Cytophaga | Microscilla | Chryseolin | 0 | 0 | 0 |
| Bacteria | Proteobact | Alphaprote | Rhizobiale | Rhizobiale | Phreatobac | 0 | 0 | 0 |
| Bacteria | Firmicutes | Bacilli | Bacillales | Planococca | Lysinibacil | 0 | 0 | 0 |
| Bacteria | Proteobact | Gammapro | Burkholde | Comamona | Pelomonas | 0 | 0 | 0 |
| Bacteria | Firmicutes | Incertae_S | DTU014 | DTU014 | DTU014 | 78 | 0 | 0 |
| Bacteria | Planctomy | Phycisphae | Phycisphae | Phycisphae | uncultured | 105 | 0 | 0 |
| Bacteria | Proteobact | Alphaprote | Acetobacte | Acetobacte | Acidiphiliu | 0 | 0 | 63 |
| Bacteria | Actinobact | Actinobact | Micrococc | Intrasporan | Pedococcu | 0 | 0 | 0 |
| Bacteria | Planctomy | Planctomy | Pirellulales | Pirellulace | __ | 39 | 0 | 0 |
| Bacteria | Proteobact | Alphaprote | Rhizobiale | KF-JG30-B | KF-JG30-B | 51 | 0 | 0 |
| Bacteria | Patescibact | Microgeno | Candidatus | Candidatus | Candidatus | 212 | 99 | 0 |
| Bacteria | Desulfobac | Desulfobac | Desulfatigl | Desulfatigl | Desulfatigl | 102 | 27 | 0 |
| Bacteria | Campyloba | Campyloba | Campyloba | Sulfurimon | Sulfuricurv | 117 | 24 | 0 |
| Bacteria | Actinobact | Thermoleo | uncultured | uncultured | uncultured | 31 | 0 | 0 |
| Bacteria | Firmicutes | Desulfoton | Desulfoton | Pelotomac | Cryptanaer | 16 | 0 | 0 |
| Bacteria | Cyanobact | Sericytoch | Sericytoch | Sericytoch | Sericytoch | 109 | 22 | 0 |
| Bacteria | Chloroflex | Anaeroline | SBR1031 | SBR1031 | SBR1031 | 84 | 17 | 0 |
| Bacteria | Verrucomi | Verrucomi | Chthoniob | Terrimicro | Terrimicro | 0 | 0 | 0 |
| Bacteria | Verrucomi | Omnitroph | Omnitroph | Omnitroph | Candidatus | 141 | 31 | 0 |
| Bacteria | Actinobact | Actinobact | Frankiales | Geodermat | __ | 112 | 0 | 0 |
| Bacteria | Firmicutes | Bacilli | Alicycloba | Alicycloba | Tumebacil | 131 | 0 | 0 |
| Bacteria | Proteobact | Gammapro | Xanthomo | Rhodanoba | Dyella | 0 | 0 | 0 |
| Bacteria | Firmicutes | Clostridia | Peptostrep | Acidamino | Acidamino | 0 | 15 | 0 |
| Bacteria | Firmicutes | Bacilli | Paenibacil | Paenibacil | Thermobac | 58 | 0 | 0 |
| Bacteria | Proteobact | Gammapro | Pseudomo | Moraxella | Acinetobac | 25 | 0 | 58 |
| Bacteria | Bacteroido | Bacteroidia | Chitinopha | Chitinopha | Hydrotalea | 0 | 0 | 0 |
| Bacteria | Proteobact | Alphaprote | Sphingomo | Sphingomo | Qipengyua | 0 | 0 | 0 |
| Bacteria | Myxococc | Polyangia | Nannocyst | Nannocyst | Nannocyst | 57 | 0 | 0 |
| Bacteria | Actinobact | Acidimicro | Actinomar | uncultured | uncultured | 57 | 0 | 0 |
| Bacteria | Proteobact | Alphaprote | Rickettsial | Rickettsiac | uncultured | 0 | 0 | 0 |
| Bacteria | Firmicutes | Limnochor | Limnochor | Limnochor | Hydrogeni | 202 | 3 | 0 |
| Bacteria | Acidobacte | Blastocatel | Blastocatel | Blastocatel | Blastocatel | 0 | 0 | 56 |
| Bacteria | Patescibac | ABY1 | Candidatus | Candidatus | Candidatus | 0 | 9 | 0 |
| Bacteria | Deinococc | Deinococc | Deinococc | Deinococc | Deinococc | 18 | 0 | 10 |
| Bacteria | Myxococc | Polyangia | UASB-TL2 | UASB-TL2 | UASB-TL2 | 55 | 0 | 0 |
| Bacteria | WS2 | WS2 | WS2 | WS2 | WS2 | 0 | 100 | 0 |
| Bacteria | Gemmatim | Longimicro | Longimicro | Longimicro | Longimicro | 0 | 0 | 0 |
| Bacteria | Bacteroido | Ignavibacte | Ignavibacte | Ignavibacte | Ignavibacte | 55 | 0 | 0 |
| Bacteria | Myxococc | Polyangia | Polyangial | Polyangiac | Minicystis | 0 | 0 | 0 |
| Bacteria | Proteobact | Gammapro | Burkholde | Comamona | Caenimona | 0 | 0 | 0 |
| Bacteria | Chloroflex | Anaeroline | uncultured | uncultured | uncultured | 37 | 0 | 0 |
| Bacteria | Firmicutes | Clostridia | Christense | Christense | Christense | 19 | 165 | 0 |
| Bacteria | Bacteroido | Bacteroidia | Bacteroida | Bacteroida | Bacteroide | 0 | 0 | 0 |
| Bacteria | Desulfobac | Syntrophol | Syntrophol | uncultured | uncultured | 96 | 0 | 0 |
| Bacteria | Firmicutes | Clostridia | Clostridial | Clostridiac | Clostridiun | 143 | 0 | 0 |
| Bacteria | Verrucomi | Verrucomi | Pedosphae | Pedosphae | RS25G | 0 | 0 | 0 |
| Bacteria | Bacteroido | Bacteroidia | Sphingoba | Lentimicro | __ | 0 | 53 | 0 |
| Bacteria | Actinobact | Actinobact | Corynebac | Nocardiace | Smaragdic | 0 | 63 | 0 |
| Bacteria | Firmicutes | Clostridia | Christense | Christense | __ | 0 | 17 | 0 |
| Bacteria | Firmicutes | Bacilli | Bacillales | Bacillaceae | Anaerobac | 24 | 0 | 0 |

| | | | | | | | | |
|---|---|---|---|---|---|---|---|---|
| Bacteria | Desulfobac | Desulfobac | Desulfobac | Desulfosar | __ | 52 | 0 | 0 |
| Bacteria | Acidobacte | Acidobacte | Subgroup_ | Subgroup_ | Subgroup_ | 44 | 0 | 0 |
| Bacteria | Proteobact | Alphaprote | Puniceispi | Puniceispi | Puniceispi | 36 | 0 | 0 |
| Bacteria | Armatimor | Chthonom | Chthonom | Chthonom | Chthonom | 12 | 0 | 0 |
| Bacteria | Firmicutes | Bacilli | Thermoact | Thermoact | __ | 91 | 0 | 0 |
| Bacteria | Acidobacte | Vicinamib | Vicinamib | Vicinamib | Luteitalea | 163 | 0 | 0 |
| Bacteria | Firmicutes | Clostridia | Clostridial | Clostridiac | uncultured | 50 | 0 | 0 |
| Bacteria | Firmicutes | Bacilli | Thermoact | Thermoact | Laceyella | 50 | 0 | 0 |
| Bacteria | Nitrospirot | 4-29-1 | 4-29-1 | 4-29-1 | 4-29-1 | 125 | 12 | 0 |
| Bacteria | Proteobact | Alphaprote | Sphingom | Sphingom | Altererythr | 0 | 53 | 0 |
| Bacteria | Firmicutes | Bacilli | Lactobacil | Carnobacte | Dolosigran | 49 | 0 | 0 |
| Bacteria | Nitrospirot | Thermodes | __ | __ | __ | 49 | 0 | 0 |
| Bacteria | Proteobact | Alphaprote | Rhizobiale | Rhizobiace | Shinella | 49 | 0 | 0 |
| Bacteria | Firmicutes | Clostridia | Clostridial | Caloramat | Fervidicell | 49 | 0 | 0 |
| Bacteria | Planctomy | Pla4_linea | Pla4_linea | Pla4_linea | Pla4_linea | 49 | 11 | 0 |
| Bacteria | Verrucomi | Chlamydia | Chlamydia | Criblamyd | uncultured | 0 | 11 | 0 |
| Bacteria | Actinobact | Actinobact | Elev-16S-9 | Elev-16S-9 | Elev-16S-9 | 19 | 0 | 0 |
| Bacteria | Bacteroido | Bacteroidia | Bacteroida | Prolixibact | WCHB1-3 | 22 | 32 | 0 |
| Bacteria | Chloroflex | Anaeroline | Thermofle | Thermofle | Thermofle | 48 | 0 | 0 |
| Bacteria | Myxococc | Polyangia | Polyangial | Sandaracin | __ | 71 | 0 | 0 |
| Bacteria | Actinobact | Acidimicro | Microtrich | Ilumatobac | Ilumatobac | 48 | 0 | 0 |
| Bacteria | WPS-2 | WPS-2 | WPS-2 | WPS-2 | WPS-2 | 0 | 0 | 0 |
| Bacteria | Desulfobac | Syntrophia | Syntrophal | Syntrophac | Syntrophus | 55 | 32 | 0 |
| Bacteria | Verrucomi | Chlamydia | Chlamydia | Criblamyd | Estrella | 0 | 7 | 0 |
| Bacteria | Firmicutes | Clostridia | Lachnospi | Lachnospi | Stomatoba | 46 | 0 | 0 |
| Bacteria | Firmicutes | Clostridia | Oscillospi | Hungateicl | __ | 71 | 12 | 0 |
| Bacteria | Firmicutes | Clostridia | Clostridial | Clostridiac | Clostridiur | 46 | 0 | 0 |
| Bacteria | Firmicutes | Clostridia | Clostridial | Clostridiac | Anaerobac | 46 | 0 | 0 |
| Bacteria | Bacteroido | Ignavibacte | SJA-28 | SJA-28 | SJA-28 | 84 | 13 | 0 |
| Bacteria | Chloroflex | SHA-26 | SHA-26 | SHA-26 | SHA-26 | 317 | 0 | 0 |
| Bacteria | PAUC34f | PAUC34f | PAUC34f | PAUC34f | PAUC34f | 0 | 54 | 0 |
| Bacteria | Armatimor | Armatimor | Armatimor | Armatimor | Armatimor | 18 | 13 | 0 |
| Bacteria | Firmicutes | Clostridia | Peptostrep | Peptostrep | Romboutsi | 17 | 0 | 0 |
| Bacteria | Bacteroido | Bacteroidia | Chitinopha | uncultured | uncultured | 0 | 0 | 0 |
| Bacteria | Proteobact | Gammapro | Xanthomo | Xanthomo | Pseudoxan | 0 | 0 | 0 |
| Bacteria | Acidobacte | Acidobacte | PAUC26f | PAUC26f | PAUC26f | 45 | 0 | 0 |
| Bacteria | Planctomy | Phycisphae | Pla1_linea | Pla1_linea | Pla1_linea | 78 | 11 | 0 |
| Bacteria | Myxococc | bacteriap2 | bacteriap2 | bacteriap2 | bacteriap2 | 56 | 0 | 0 |
| Bacteria | NB1-j | NB1-j | NB1-j | NB1-j | NB1-j | 62 | 9 | 0 |
| Bacteria | Acidobacte | Acidobacte | Elev-16S-1 | Elev-16S-1 | Elev-16S-1 | 0 | 0 | 0 |
| Bacteria | Acidobacte | Subgroup_ | Subgroup_ | Subgroup_ | Subgroup_ | 79 | 0 | 0 |
| Bacteria | Proteobact | Gammapro | HOC36 | HOC36 | HOC36 | 136 | 0 | 0 |
| Bacteria | Actinobact | Thermoleo | Solirubrob | Solirubrob | Solirubrob | 44 | 0 | 0 |
| Bacteria | Verrucomi | Verrucomi | Verrucomi | DEV007 | DEV007 | 0 | 44 | 0 |
| Bacteria | Firmicutes | Clostridia | Lachnospi | Lachnospi | Herbinix | 42 | 25 | 0 |
| Bacteria | Proteobact | Alphaprote | Rhodobact | Rhodobact | Defluviimo | 0 | 0 | 0 |
| Bacteria | Verrucomi | Lentisphae | Victivallal | Victivallal | Victivallal | 0 | 0 | 0 |
| Bacteria | Firmicutes | Clostridia | Christense | Christense | uncultured | 76 | 85 | 0 |
| Bacteria | Firmicutes | __ | __ | __ | __ | 27 | 0 | 0 |
| Bacteria | Bacteroido | Ignavibacte | Ignavibacte | UA-50 | UA-50 | 42 | 0 | 0 |

| | | | | | | | | |
|---|---|---|---|---|---|---|---|---|
| Bacteria | Verrucomi | Verrucomi | Pedosphae | Pedosphae | ADurb.Bin | 75 | 0 | 0 |
| Bacteria | Firmicutes | Desulfitob | Desulfitob | __ | __ | 42 | 0 | 0 |
| Bacteria | WS1 | WS1 | WS1 | WS1 | WS1 | 107 | 25 | 0 |
| Bacteria | Proteobact | Alphaprote | Sphingomo | Sphingomo | Plot4-2H12 | 42 | 0 | 0 |
| Bacteria | Fibrobacte | Fibrobacte | Fibrobacte | Fibrobacte | BBMC-4 | 99 | 0 | 0 |
| Bacteria | Patescibac | Parcubacte | Candidatus | Candidatus | Candidatus | 0 | 34 | 0 |
| Bacteria | Proteobact | Gammapro | Burkholde | Sutterellac | uncultured | 21 | 0 | 0 |
| Bacteria | Firmicutes | Bacilli | Paenibacil | Paenibacil | __ | 31 | 0 | 0 |
| Bacteria | Verrucomi | Verrucomi | Opitutales | Puniceicoc | __ | 41 | 0 | 0 |
| Bacteria | Firmicutes | Syntropho | Syntropho | Syntropho | Syntropho | 129 | 0 | 0 |
| Bacteria | Proteobact | Gammapro | Burkholde | SC-I-84 | SC-I-84 | 108 | 0 | 0 |
| Bacteria | Actinobact | Thermoleo | Solirubrob | Solirubrob | Patulibacte | 41 | 0 | 0 |
| Bacteria | Bacteroido | Bacteroidia | Flavobacte | Cryomorph | uncultured | 0 | 41 | 0 |
| Bacteria | Firmicutes | Bacilli | Paenibacil | Paenibacil | Cohnella | 0 | 0 | 0 |
| Bacteria | Chloroflex | __ | __ | __ | __ | 111 | 12 | 0 |
| Bacteria | Acidobacte | Acidobacte | GOUTB8 | GOUTB8 | GOUTB8 | 86 | 0 | 0 |
| Bacteria | Proteobact | Gammapro | Burkholde | Gallionella | __ | 56 | 0 | 0 |
| Bacteria | Verrucomi | Kiritimatie | WCHB1-4 | WCHB1-4 | WCHB1-4 | 44 | 100 | 0 |
| Bacteria | Proteobact | Alphaprote | Tistrellales | Geminicoc | Candidatus | 40 | 0 | 0 |
| Bacteria | Firmicutes | Clostridia | Clostridial | Clostridiac | Clostridiur | 112 | 0 | 0 |
| Bacteria | Actinobact | WCHB1-8 | WCHB1-8 | WCHB1-8 | WCHB1-8 | 72 | 13 | 0 |
| Bacteria | Firmicutes | Clostridia | Lachnospi | Lachnospi | __ | 142 | 16 | 0 |
| Bacteria | Chloroflex | Anaeroline | Anaeroline | Anaeroline | Anaeroline | 159 | 29 | 0 |
| Bacteria | Armatimor | __ | __ | __ | __ | 7 | 0 | 0 |
| Bacteria | Proteobact | Gammapro | Pseudomor | Moraxella | uncultured | 0 | 40 | 0 |
| Bacteria | Verrucomi | Verrucomi | Pedosphae | Pedosphae | uncultured | 50 | 0 | 0 |
| Bacteria | Actinobact | Thermoleo | Solirubrob | __ | __ | 0 | 0 | 0 |
| Bacteria | Proteobact | Gammapro | Burkholde | Comamona | Paucibacte | 0 | 0 | 0 |
| Bacteria | Myxococc | Myxococc | Myxococc | 27F-1492R | 27F-1492R | 39 | 4 | 0 |
| Bacteria | Patescibac | Parcubacte | Candidatus | Candidatus | Candidatus | 49 | 151 | 0 |
| Bacteria | Verrucomi | Chlamydia | Chlamydia | Parachlam | Parachlam | 0 | 0 | 0 |
| Bacteria | Firmicutes | Bacilli | Erysipelot | Erysipelot | Turicibacte | 0 | 0 | 0 |
| Bacteria | Firmicutes | Desulfitob | Desulfitob | Desulfitob | __ | 0 | 55 | 0 |
| Bacteria | Verrucomi | Verrucomi | Verrucomi | Verrucomi | Verrucomi | 0 | 0 | 0 |
| Bacteria | Planctomy | OM190 | OM190 | OM190 | OM190 | 18 | 5 | 0 |
| Bacteria | Bacteroido | __ | __ | __ | __ | 0 | 0 | 0 |
| Bacteria | Firmicutes | Clostridia | Oscillospi | Hungateicl | Ruminiclo | 31 | 20 | 0 |
| Bacteria | Actinobact | Actinobact | Kineospor | Kineospor | Angustibac | 22 | 0 | 0 |
| Bacteria | Armatimor | DG-56 | DG-56 | DG-56 | DG-56 | 37 | 0 | 0 |
| Bacteria | Myxococc | Polyangia | Polyangial | Polyangiac | Byssovora | 37 | 0 | 0 |
| Bacteria | Chloroflex | Dehalococ | S085 | S085 | S085 | 52 | 0 | 0 |
| Bacteria | Proteobact | Gammapro | Salinisphae | Solimonad | uncultured | 37 | 0 | 0 |
| Bacteria | Proteobact | Gammapro | Competiba | Competiba | Candidatus | 0 | 0 | 37 |
| Bacteria | Desulfobac | Desulfuror | Desulfuror | __ | __ | 0 | 37 | 0 |
| Bacteria | Planctomy | __ | __ | __ | __ | 0 | 44 | 0 |
| Bacteria | Proteobact | Gammapro | Gammapro | Unknown_ | Unknown_ | 0 | 43 | 0 |
| Bacteria | Patescibac | Gracilibact | Gracilibact | Gracilibact | Gracilibact | 39 | 22 | 0 |
| Bacteria | Proteobact | Gammapro | Gammapro | Unknown_ | Candidatus | 13 | 26 | 0 |
| Bacteria | Proteobact | Gammapro | Burkholde | Rhodocycl | __ | 0 | 5 | 0 |
| Bacteria | Proteobact | Gammapro | Burkholde | Rhodocycl | Sulfuritale | 0 | 46 | 0 |

| | | | | | | | | |
|---|---|---|---|---|---|---|---|---|
| Bacteria | Proteobact | Gammaprc | Burkholde | Rhodocycl | Sterolibact | 0 | 36 | 0 |
| Bacteria | Proteobact | Gammaprc | Legionella | Legionella | __ | 26 | 0 | 0 |
| Bacteria | Proteobact | Alphaprote | Rhizobiale | Beijerincki | 1174-901-1 | 0 | 0 | 0 |
| Bacteria | Actinobact | Actinobact | Propioniba | __ | __ | 0 | 0 | 0 |
| Bacteria | Proteobact | Alphaprote | Rhodobact | Rhodobact | Gemmobac | 0 | 0 | 0 |
| Bacteria | Proteobact | Alphaprote | Defluviico | Defluviico | Defluviico | 101 | 0 | 0 |
| Bacteria | Actinobact | Actinobact | Pseudonoc | Pseudonoc | Pseudonoc | 35 | 0 | 0 |
| Bacteria | Firmicutes | Clostridia | Clostridial | Oxobactera | Oxobacter | 35 | 0 | 0 |
| Bacteria | Proteobact | Alphaprote | Rhizobiale | Hyphomici | Pedomicro | 21 | 0 | 34 |
| Eukaryota | __ | __ | __ | __ | __ | 6 | 0 | 0 |
| Bacteria | Proteobact | Gammaprc | Xanthomo | __ | __ | 0 | 0 | 0 |
| Bacteria | Acidobacte | Acidobacte | Acidobacte | Acidobacte | uncultured | 34 | 0 | 0 |
| Bacteria | Firmicutes | Clostridia | __ | __ | __ | 34 | 17 | 0 |
| Bacteria | Firmicutes | Bacilli | Brevibacill | Brevibacill | Brevibacill | 9 | 0 | 0 |
| Bacteria | Firmicutes | Negativicu | Veillonella | Sporomusa | Pelosinus | 31 | 0 | 0 |
| Bacteria | Patescibac | Saccharim | Saccharim | WWH38 | WWH38 | 0 | 10 | 0 |
| Bacteria | Actinobact | Actinobact | Corynebac | Corynebac | Lawsonella | 0 | 0 | 0 |
| Bacteria | Planctomy | Planctomy | Isosphaera | Isosphaera | uncultured | 130 | 0 | 0 |
| Bacteria | Patescibac | CPR2 | CPR2 | CPR2 | CPR2 | 149 | 12 | 0 |
| Bacteria | Firmicutes | Clostridia | Oscillospir | Hungateicl | Acetivibric | 33 | 0 | 0 |
| Bacteria | Proteobact | Gammaprc | Diplorické | Diplorické | __ | 33 | 7 | 0 |
| Bacteria | Chloroflex | Dehalococ | Dehalococ | uncultured | uncultured | 137 | 0 | 0 |
| Bacteria | Firmicutes | Symbiobac | Symbiobac | Symbiobac | Symbiobac | 33 | 0 | 0 |
| Bacteria | Myxococc | Myxococc | Myxococca | Myxococca | KD3-10 | 47 | 0 | 0 |
| Bacteria | Desulfobac | Desulfuron | Geobactera | Geobactera | Geobacter | 80 | 33 | 0 |
| Bacteria | Proteobact | Alphaprote | Rhizobiale | Methylolig | uncultured | 0 | 0 | 0 |
| Bacteria | Firmicutes | Clostridia | Peptostrep | Family_XI | Finegoldia | 32 | 0 | 0 |
| Bacteria | Acidobacte | Acidobacte | Acidobacte | Koribacter | Candidatus | 32 | 0 | 0 |
| Bacteria | Verrucomi | Lentisphae | Oligosphae | Lenti-02 | Lenti-02 | 54 | 72 | 0 |
| Bacteria | Proteobact | Alphaprote | Rhizobiale | Beijerincki | Methylosin | 11 | 0 | 0 |
| Bacteria | Firmicutes | Clostridia | Peptostrep | Peptostrep | __ | 31 | 0 | 0 |
| Bacteria | TX1A-33 | TX1A-33 | TX1A-33 | TX1A-33 | TX1A-33 | 31 | 0 | 0 |
| Bacteria | Desulfobac | Syntrophia | Syntrophal | __ | __ | 106 | 0 | 0 |
| Bacteria | Desulfobac | Syntrophol | Syntrophol | Syntrophol | Syntrophol | 44 | 0 | 0 |
| Bacteria | Spirochaet | Spirochaet | Spirochaet | Spirochaet | Spirochaet | 70 | 0 | 0 |
| Bacteria | Patescibac | Saccharim | Saccharim | YM_S32_T | YM_S32_T | 0 | 31 | 0 |
| Bacteria | Firmicutes | Clostridia | Lachnospii | Lachnospii | Cellulosily | 21 | 0 | 0 |
| Bacteria | Proteobact | Alphaprote | Dongiales | Dongiacea | Dongia | 0 | 0 | 0 |
| Bacteria | Verrucomi | Omnitroph | Omnitroph | __ | __ | 37 | 4 | 0 |
| Bacteria | Proteobact | Alphaprote | Paracaedib | Paracaedib | Candidatus | 0 | 0 | 0 |
| Bacteria | Proteobact | Gammaprc | Burkholde | Sulfuricell | __ | 0 | 29 | 0 |
| Bacteria | Proteobact | Gammaprc | Burkholde | Comamona | Rhizobacte | 0 | 0 | 0 |
| Bacteria | Actinobact | Thermoleo | __ | __ | __ | 0 | 0 | 0 |
| Bacteria | Bacteroido | Bacteroidia | Bacteroida | Bacteroide | Bacteroide | 54 | 0 | 0 |
| Bacteria | Chloroflex | Anaeroline | Anaeroline | Anaeroline | RBG-16-58 | 34 | 18 | 0 |
| Bacteria | Acidobacte | Thermoana | Thermoana | Thermoana | TPD-58 | 0 | 54 | 0 |
| Bacteria | Actinobact | Thermoleo | Solirubrob | Solirubrob | JCM_1899 | 0 | 0 | 0 |
| Bacteria | Fusobacter | Fusobacter | Fusobacter | Fusobacter | Fusobacter | 0 | 0 | 0 |
| Bacteria | Firmicutes | Clostridia | Peptostrep | Family_XI | Peptoniphi | 0 | 0 | 0 |
| Bacteria | Verrucomi | Verrucomi | Pedosphae | Pedosphae | DEV114 | 0 | 0 | 0 |

| | | | | | | | | |
|---|---|---|---|---|---|---|---|---|
| Bacteria | Bacteroido | Bacteroidia | Chitinopha | Chitinopha | Flaviaestur | 0 | 0 | 0 |
| Bacteria | Proteobact | Alphaprote | Rhodobact | Rhodobact | Pseudorho | 28 | 0 | 0 |
| Bacteria | Sumerlaeo | Sumerlaeia | Sumerlaeal | Sumerlaea | Sumerlaea | 33 | 0 | 0 |
| Bacteria | Patescibac | Parcubacte | Candidatus | Candidatus | Candidatus | 74 | 39 | 0 |
| Bacteria | Chloroflex | Anaeroline | Anaeroline | Anaeroline | Pelolinea | 28 | 9 | 0 |
| Bacteria | Cyanobact | Cyanobact | Synechoco | Cyanobiac | Cyanobium | 0 | 52 | 0 |
| Bacteria | Proteobact | Alphaprote | Rhodospiri | Rhodospiri | uncultured | 0 | 51 | 0 |
| Bacteria | Proteobact | Alphaprote | Rhizobiale | Rhizobiale | Nordella | 0 | 0 | 0 |
| Bacteria | Dependent | Babeliae | Babeliales | Babeliales | Babeliales | 27 | 3 | 0 |
| Bacteria | Verrucomi | Verrucomi | Verrucomi | Verrucomi | Roseimicro | 0 | 0 | 0 |
| Bacteria | Proteobact | Gammapro | Burkholde | Sulfuricell | Ferritrophi | 0 | 11 | 0 |
| Bacteria | Deferrison | Defferrison | Defferrison | Defferrison | Deferrison | 69 | 0 | 0 |
| Bacteria | Patescibac | Parcubacte | Candidatus | Candidatus | Candidatus | 0 | 0 | 0 |
| Bacteria | Actinobact | Actinobact | uncultured | uncultured | uncultured | 26 | 0 | 0 |
| Bacteria | Firmicutes | Clostridia | Clostridial | Clostridiac | Clostridiac | 26 | 0 | 0 |
| Bacteria | Latescibac | Latescibac | Latescibac | Latescibac | Candidatus | 26 | 0 | 0 |
| Bacteria | Firmicutes | Clostridia | Clostridia | Gracilibact | Gracilibact | 40 | 0 | 0 |
| Bacteria | Firmicutes | Negativicu | Veillonella | Sporomusa | Sporomusa | 0 | 0 | 0 |
| Bacteria | Acidobacte | Blastocatel | __ | __ | __ | 0 | 0 | 0 |
| Bacteria | Firmicutes | Clostridia | Peptostrep | Family_XI | Anaerococ | 23 | 0 | 0 |
| Bacteria | Desulfobac | Desulfuron | PB19 | PB19 | PB19 | 0 | 0 | 0 |
| Bacteria | Cyanobact | Vampirivib | Obscuribac | Obscuribac | Obscuribac | 0 | 0 | 0 |
| Bacteria | Proteobact | Gammapro | Burkholde | Burkholde | Ralstonia | 0 | 0 | 0 |
| Bacteria | Verrucomi | Verrucomi | Pedosphae | Pedosphae | DEV008 | 0 | 0 | 0 |
| Bacteria | Myxococc | Myxococci | Myxococca | Myxococca | __ | 70 | 0 | 0 |
| Bacteria | Firmicutes | Bacilli | Paenibacill | Paenibacill | Paenibacill | 25 | 0 | 0 |
| Bacteria | Actinobact | Actinobact | Corynebac | Nocardiace | Nocardia | 25 | 0 | 0 |
| Bacteria | Planctomy | Phycisphae | MSBL9 | SG8-4 | SG8-4 | 134 | 0 | 0 |
| Bacteria | Acidobacte | Blastocatel | 24-11 | 24-11 | 24-11 | 25 | 0 | 0 |
| Bacteria | Proteobact | Alphaprote | Tistrellales | Geminicoc | __ | 0 | 0 | 25 |
| Bacteria | Cyanobact | Vampirivib | Gastranaer | Gastranaer | Gastranaer | 43 | 46 | 0 |
| Bacteria | Proteobact | Gammapro | R7C24 | R7C24 | R7C24 | 0 | 5 | 0 |
| Bacteria | Firmicutes | Negativicu | Veillonella | __ | __ | 0 | 0 | 0 |
| Bacteria | Chloroflex | JG30-KF-C | JG30-KF-C | JG30-KF-C | JG30-KF-C | 17 | 23 | 0 |
| Bacteria | Firmicutes | Symbiobac | Symbiobac | Symbiobac | uncultured | 0 | 0 | 0 |
| Bacteria | Actinobact | Actinobact | Micrococc | Microbacte | Aurantimic | 0 | 0 | 0 |
| Bacteria | Acidobacte | Acidobacte | __ | __ | __ | 47 | 0 | 0 |
| Bacteria | Actinobact | Actinobact | Micrococc | Cellulomo | __ | 24 | 20 | 0 |
| Bacteria | Patescibac | Kazania | Kazania | Kazania | Kazania | 0 | 40 | 0 |
| Bacteria | Planctomy | Planctomy | Gemmatale | Gemmatac | Gemmata | 0 | 24 | 0 |
| Bacteria | Proteobact | Alphaprote | Rhizobiale | Labraceae | Labrys | 19 | 0 | 0 |
| Bacteria | Firmicutes | Bacilli | Lactobacil | Lactobacil | Lactobacil | 0 | 0 | 14 |
| Bacteria | Desulfobac | Desulfovib | Desulfovib | Desulfovib | Desulfovib | 0 | 0 | 0 |
| Bacteria | Firmicutes | Clostridia | Peptostrep | Anaerovor | __ | 23 | 0 | 0 |
| Bacteria | Firmicutes | Negativicu | Veillonella | Sporomusa | Anaerosinu | 23 | 0 | 0 |
| Bacteria | Firmicutes | Desulfoton | Desulfoton | Desulfoton | Desulfofar | 23 | 0 | 0 |
| Bacteria | Acidobacte | __ | __ | __ | __ | 42 | 0 | 0 |
| Bacteria | Chloroflex | Anaeroline | Caldilineal | Caldilineac | Caldilineac | 23 | 0 | 0 |
| Bacteria | Acidobacte | Subgroup_ | Subgroup_ | Subgroup_ | Subgroup_ | 36 | 23 | 0 |
| Bacteria | Proteobact | Gammapro | 1013-28-C | 1013-28-C | 1013-28-C | 0 | 23 | 0 |

| | | | | | | | | |
|---|---|---|---|---|---|---|---|---|
| Bacteria | Dependent | Babeliae | Babeliales | UBA1240 | UBA1240 | 16 | 28 | 0 |
| Bacteria | Proteobact | Gammapro | Ga007753 | Ga007753 | Ga007753 | 22 | 0 | 0 |
| Bacteria | Elusimicro | Elusimicro | MVP-88 | MVP-88 | MVP-88 | 0 | 0 | 0 |
| Bacteria | Actinobact | Actinobact | Propioniba | Propioniba | Friedmann | 0 | 0 | 0 |
| Bacteria | Actinobact | Actinobact | Micrococc | Micrococc | Kocuria | 0 | 0 | 0 |
| Bacteria | Desulfoba | Desulfuron | Geobacter | __ | __ | 22 | 20 | 0 |
| Bacteria | Planctomy | Planctomy | Pirellulales | Pirellulace | uncultured | 22 | 0 | 0 |
| Bacteria | Verrucomi | Verrucomi | Verrucomi | Rubritalea | Luteolibac | 30 | 14 | 0 |
| Bacteria | Gemmatim | __ | __ | __ | __ | 22 | 0 | 0 |
| Bacteria | Firmicutes | Bacilli | Lactobacil | __ | __ | 38 | 0 | 0 |
| Bacteria | Acidobact | Acidobact | Subgroup_ | Subgroup_ | Subgroup_ | 22 | 0 | 0 |
| Bacteria | Proteobact | Gammapro | PLTA13 | PLTA13 | PLTA13 | 0 | 6 | 0 |
| Bacteria | Chloroflex | Dehalococ | 661239 | 661239 | 661239 | 0 | 3 | 0 |
| Bacteria | Armatimo | Chthonom | Chthonom | Chthonom | Chthonom | 0 | 0 | 0 |
| Bacteria | Actinobact | Actinobact | Streptomy | Streptomy | Kitasatosp | 0 | 0 | 0 |
| Bacteria | Firmicutes | Clostridia | Oscillospi | __ | __ | 10 | 0 | 0 |
| Eukaryota | Retaria | Foraminife | Rotaliida | Rotaliidae | Ammonia | 0 | 0 | 0 |
| Bacteria | Firmicutes | Desulfoton | Desulfoton | Pelotomac | Pelotomac | 21 | 0 | 0 |
| Bacteria | Patescibac | Microgeno | Candidatus | Candidatus | Candidatus | 21 | 0 | 0 |
| Bacteria | Proteobact | Gammapro | Pseudomo | __ | __ | 21 | 0 | 0 |
| Bacteria | Desulfoba | Syntrophol | Syntrophol | Syntrophol | __ | 35 | 0 | 0 |
| Bacteria | Chloroflex | Chloroflex | Chloroflex | Herpetosip | Herpetosip | 21 | 0 | 0 |
| Bacteria | Verrucomi | __ | __ | __ | __ | 0 | 28 | 0 |
| Bacteria | Proteobact | Gammapro | Burkholde | Burkholde | Lautropia | 0 | 0 | 0 |
| Bacteria | Planctomy | Pla3_linea | Pla3_linea | Pla3_linea | Pla3_linea | 20 | 0 | 0 |
| Bacteria | Bacteroido | Bacteroidi | Bacteroida | Prolixibact | BSV13 | 20 | 0 | 0 |
| Bacteria | Actinobact | Actinobact | Propioniba | Propioniba | __ | 20 | 0 | 0 |
| Bacteria | Chloroflex | Dehalococ | Sh765B-T | Sh765B-T | Sh765B-T | 20 | 0 | 0 |
| Bacteria | Verrucomi | Verrucomi | Chthoniob | Chthoniob | Candidatus | 23 | 0 | 0 |
| Bacteria | Proteobact | Gammapro | uncultured | uncultured | uncultured | 20 | 0 | 0 |
| Bacteria | Planctomy | Phycispha | Phycispha | AKAU356 | AKAU356 | 20 | 0 | 0 |
| Bacteria | Firmicutes | Clostridia | Peptostrep | Peptostrep | Sporacetig | 20 | 0 | 0 |
| Bacteria | Firmicutes | Clostridia | Clostridial | Clostridiac | Clostridiur | 0 | 0 | 0 |
| Bacteria | Elusimicro | Lineage_II | Lineage_II | Lineage_II | Lineage_II | 0 | 13 | 0 |
| Bacteria | Proteobact | Gammapro | Methyloco | Methyloco | Methylote | 0 | 0 | 0 |
| Eukaryota | Retaria | Foraminife | Rotaliida | __ | __ | 0 | 0 | 0 |
| Bacteria | Chloroflex | Anaeroline | Ardenticat | uncultured | uncultured | 19 | 14 | 0 |
| Bacteria | Chloroflex | Chloroflex | Kallotenua | AKIW781 | AKIW781 | 19 | 0 | 0 |
| Bacteria | Abditibact | Abditibact | Abditibact | Abditibact | Abditibact | 25 | 13 | 0 |
| Bacteria | Actinobact | Actinobact | Frankiales | Acidother | Acidother | 19 | 0 | 0 |
| Bacteria | Actinobact | Acidimicro | uncultured | uncultured | uncultured | 19 | 0 | 0 |
| Bacteria | Chloroflex | Anaeroline | Anaeroline | Anaeroline | Leptolinea | 33 | 28 | 0 |
| Bacteria | Patescibac | Parcubacte | Candidatus | Candidatus | Candidatus | 0 | 19 | 0 |
| Bacteria | LCP-89 | LCP-89 | LCP-89 | LCP-89 | LCP-89 | 0 | 19 | 0 |
| Bacteria | Firmicutes | Clostridia | Peptostrep | Thermotal | Anaerosoli | 0 | 11 | 0 |
| Bacteria | Proteobact | Gammapro | Burkholde | TRA3-20 | TRA3-20 | 12 | 5 | 0 |
| Bacteria | Desulfoba | Desulfuron | __ | __ | __ | 0 | 0 | 0 |
| Bacteria | Firmicutes | Clostridia | Lachnospi | Lachnospi | Mobilitale | 0 | 0 | 0 |
| Bacteria | Firmicutes | Clostridia | Clostridial | Clostridiac | Clostridiur | 0 | 0 | 0 |
| Bacteria | Proteobact | Gammapro | Burkholde | Comamona | Acidovora | 0 | 0 | 0 |

| | | | | | | | | |
|---|---|---|---|---|---|---|---|---|
| Bacteria | Myxococc | Myxococc | Myxococc | Myxococc | Corallococ | 18 | 0 | 0 |
| Bacteria | Actinobact | Actinobact | Micrococc | Cellulomo | Pseudactin | 18 | 0 | 0 |
| Bacteria | Planctomy | Phycisphae | Phycisphae | Phycisphae | Phycisphae | 18 | 0 | 0 |
| Bacteria | Chloroflex | Anaeroline | Anaeroline | Anaeroline | Bellilinea | 18 | 0 | 0 |
| Bacteria | Proteobact | Gammaprc | Pseudomo | Moraxella | Permianiba | 0 | 18 | 0 |
| Bacteria | Firmicutes | Clostridia | Clostridial | __ | __ | 0 | 0 | 0 |
| Bacteria | Proteobact | Alphaprote | Rhizobiale | Rhizobiale | __ | 0 | 0 | 0 |
| Bacteria | Bacteroido | Bacteroidia | Cytophaga | Spirosoma | Dyadobact | 0 | 0 | 0 |
| Bacteria | Firmicutes | Desulfitob | Desulfitob | Heliobacte | uncultured | 17 | 0 | 0 |
| Bacteria | Firmicutes | Bacilli | Bacillales | Bacillaceae | Caldibacill | 17 | 0 | 0 |
| Bacteria | Actinobact | Actinobact | Corynebac | Dietziacea | Dietzia | 17 | 0 | 0 |
| Bacteria | Actinobact | Actinobact | Streptomy | Streptomy | Streptomy | 17 | 0 | 0 |
| Bacteria | Proteobact | Gammaprc | Enterobact | Pasteurella | Haemophil | 17 | 0 | 0 |
| Bacteria | Firmicutes | Limnochor | Limnochor | Limnochor | Limnochor | 17 | 0 | 0 |
| Bacteria | Bacteroido | Bacteroidia | Bacteroida | __ | __ | 0 | 17 | 0 |
| Bacteria | Firmicutes | Clostridia | Clostridial | Clostridiac | Clostridiur | 8 | 0 | 0 |
| Bacteria | Firmicutes | Clostridia | Oscillospi | Hungateicl | Thermoclo | 0 | 0 | 0 |
| Bacteria | Actinobact | Coriobacte | __ | __ | __ | 0 | 0 | 0 |
| Bacteria | Bacteroido | Bacteroidia | Cytophaga | Amoeboph | Candidatus | 0 | 0 | 0 |
| Bacteria | Patescibac | Parcubacte | Candidatus | Candidatus | Candidatus | 0 | 26 | 0 |
| Bacteria | Firmicutes | Negativicu | Veillonella | Sporomusa | uncultured | 0 | 0 | 0 |
| Bacteria | Actinobact | Coriobacte | Coriobacte | Eggerthella | DNF00809 | 16 | 0 | 0 |
| Bacteria | Chloroflex | Chloroflex | Chloroflex | Roseiflexa | uncultured | 16 | 0 | 0 |
| Bacteria | Firmicutes | Clostridia | Peptostrep | Sedimentib | Sedimentib | 16 | 0 | 0 |
| Bacteria | Chloroflex | Anaeroline | Anaeroline | Anaeroline | Anaeroline | 16 | 0 | 0 |
| Bacteria | Planctomy | Phycisphae | MSBL9 | 4572-13 | 4572-13 | 16 | 0 | 0 |
| Bacteria | Bacteroido | Bacteroidia | Chitinopha | Saprospira | Phaeodacty | 0 | 0 | 0 |
| Bacteria | Fibrobacte | Fibrobacte | Fibrobacte | Candidatus | Candidatus | 15 | 0 | 0 |
| Bacteria | Proteobact | Alphaprote | Rhizobiale | Xanthobac | Rhodoplan | 15 | 4 | 0 |
| Bacteria | Chloroflex | Dehalococ | __ | __ | __ | 15 | 0 | 0 |
| Bacteria | Proteobact | Gammaprc | Burkholde | Rhodocycl | Ferribacter | 15 | 0 | 0 |
| Bacteria | Firmicutes | Clostridia | Peptostrep | Family_XI | Soehngenia | 15 | 0 | 0 |
| Bacteria | Proteobact | Alphaprote | Rhizobiale | Beijerinck | Methyloros | 15 | 0 | 0 |
| Bacteria | Actinobact | Actinobact | Bifidobact | Bifidobact | Bifidobact | 0 | 0 | 8 |
| Bacteria | Firmicutes | Syntrophor | Syntrophor | Syntrophor | __ | 0 | 0 | 0 |
| Bacteria | Spirochaet | Spirochaet | Spirochaet | Spirochaet | __ | 6 | 9 | 0 |
| Bacteria | Firmicutes | Bacilli | Alicycloba | Alicycloba | Alicycloba | 0 | 0 | 0 |
| Bacteria | Patescibac | Saccharim | Saccharim | Saccharim | TM7a | 0 | 0 | 0 |
| Bacteria | Firmicutes | Bacilli | Caldalkalib | Caldalkalib | Caldalkalib | 0 | 0 | 0 |
| Bacteria | Firmicutes | Clostridia | Oscillospi | Ruminococ | __ | 0 | 0 | 0 |
| Bacteria | Verrucomi | Lentisphae | Victivallal | PRD18C08 | PRD18C08 | 0 | 0 | 0 |
| Bacteria | Planctomy | Planctomy | Isosphaera | Isosphaera | Singulisph | 14 | 0 | 0 |
| Bacteria | Sumerlaeo | Sumerlaeia | uncultured | uncultured | uncultured | 14 | 0 | 0 |
| Bacteria | Planctomy | Phycisphae | mle1-8 | mle1-8 | mle1-8 | 27 | 0 | 0 |
| Bacteria | Actinobact | Actinobact | __ | __ | __ | 23 | 0 | 0 |
| Bacteria | Chloroflex | Ktedonoba | C0119 | C0119 | C0119 | 14 | 0 | 0 |
| Bacteria | Bacteroido | Ignavibact | Ignavibact | BSV40 | BSV40 | 14 | 0 | 0 |
| Bacteria | Patescibac | Microgeno | Candidatus | Candidatus | Candidatus | 3 | 25 | 0 |
| Bacteria | Proteobact | Alphaprote | Rhizobiale | Rhizobiace | Ensifer | 0 | 0 | 0 |
| Bacteria | Chloroflex | P2-11E | P2-11E | P2-11E | P2-11E | 0 | 0 | 0 |

| | | | | | | | | |
|---|---|---|---|---|---|---|---|---|
| Bacteria | Desulfobac | Desulfobul | Desulfobul | __ | __ | 0 | 0 | 0 |
| Bacteria | Firmicutes | Clostridia | Oscillospir | Hungateicl | Ercella | 13 | 0 | 0 |
| Bacteria | Desulfobac | Desulfobac | Desulfobac | Desulfosar | uncultured | 13 | 0 | 0 |
| Bacteria | Cloacimon | Cloacimon | Cloacimon | Cloacimon | Candidatus | 13 | 0 | 0 |
| Bacteria | Chloroflex | AD3 | AD3 | AD3 | AD3 | 13 | 0 | 0 |
| Bacteria | Latescibac | Latescibac | Latescibac | Latescibac | Latescibac | 13 | 0 | 0 |
| Bacteria | Firmicutes | Clostridia | Peptostrep | Family_XI | Tissierella | 13 | 0 | 0 |
| Bacteria | Chloroflex | Dehalococ | vadinBA26 | vadinBA26 | vadinBA26 | 0 | 0 | 0 |
| Bacteria | Proteobact | Gammapro | Steroidoba | Steroidoba | Steroidoba | 0 | 0 | 0 |
| Bacteria | Firmicutes | Bacilli | Thermoact | Thermoact | Novibacill | 0 | 0 | 0 |
| Bacteria | Firmicutes | Clostridia | Caldicopro | Caldicopro | Caldicopro | 0 | 0 | 0 |
| Bacteria | Bacteroido | Bacteroidia | Sphingoba | Sphingoba | Mucilagini | 0 | 8 | 0 |
| Bacteria | Actinobact | Actinobact | Kineospori | Kineospori | Kineospori | 12 | 0 | 0 |
| Bacteria | Spirochaet | Leptospira | Leptospira | Leptospira | Leptospira | 12 | 0 | 0 |
| Bacteria | Proteobact | Alphaprote | Rhizobiale | A0839 | A0839 | 0 | 12 | 0 |
| Bacteria | Chloroflex | Anaeroline | Caldilineal | Caldilinea | __ | 0 | 0 | 0 |
| Bacteria | Firmicutes | Clostridia | Lachnospi | Lachnospi | Anaerocolu | 0 | 0 | 0 |
| Bacteria | Actinobact | Actinobact | Pseudonoc | Pseudonoc | __ | 0 | 0 | 0 |
| Bacteria | Proteobact | Alphaprote | Rhizobiale | Rhizobiace | Aureimona | 8 | 0 | 0 |
| Bacteria | Proteobact | Gammapro | Chromatia | Chromatia | uncultured | 0 | 0 | 0 |
| Bacteria | Proteobact | Alphaprote | Rickettsial | Rickettsial | Candidatus | 10 | 0 | 0 |
| Bacteria | Verrucomi | Chlamydia | __ | __ | __ | 9 | 0 | 0 |
| Bacteria | Proteobact | Alphaprote | Caulobacte | Hyphomon | uncultured | 0 | 0 | 0 |
| Bacteria | Proteobact | Alphaprote | Sphingomo | Sphingomo | Sphingobiu | 0 | 5 | 0 |
| Bacteria | Firmicutes | Bacilli | Staphyloco | Staphyloco | Jeotgalicoc | 0 | 0 | 0 |
| Bacteria | Firmicutes | Bacilli | Bacillales | Bacillacea | Oceanobac | 0 | 0 | 0 |
| Bacteria | Planctomy | Phycisphae | MSBL9 | KCLunmb | KCLunmb | 11 | 0 | 0 |
| Bacteria | Elusimicro | Endomicro | Endomicro | Endomicro | Endomicro | 11 | 0 | 0 |
| Bacteria | Firmicutes | Clostridia | Oscillospir | Hungateicl | Pseudobac | 11 | 0 | 0 |
| Bacteria | Fibrobacte | Chitinivibr | uncultured | uncultured | uncultured | 11 | 8 | 0 |
| Bacteria | Proteobact | Gammapro | Burkholde | B1-7BS | B1-7BS | 11 | 0 | 0 |
| Bacteria | Firmicutes | Bacilli | Lactobacil | Enterococc | __ | 0 | 0 | 11 |
| Bacteria | Bacteroido | Bacteroidia | Bacteroida | Prolixibact | __ | 0 | 11 | 0 |
| Bacteria | Firmicutes | Desulfitob | Desulfitob | Desulfitob | Desulfitob | 0 | 0 | 0 |
| Bacteria | Firmicutes | Clostridia | Peptostrep | Family_XI | __ | 0 | 0 | 0 |
| Bacteria | Acidobacte | AT-s3-28 | AT-s3-28 | AT-s3-28 | AT-s3-28 | 0 | 0 | 0 |
| Bacteria | Firmicutes | Clostridia | Peptostrep | Peptostrep | Peptostrep | 0 | 0 | 0 |
| Bacteria | Patescibac | Saccharim | Saccharim | Saccharim | Saccharim | 0 | 0 | 0 |
| Bacteria | Actinobact | Actinobact | Micrococc | Micrococc | Micrococc | 0 | 0 | 0 |
| Bacteria | Proteobact | Alphaprote | Rhizobiale | Devosiacea | Arsenicital | 10 | 0 | 0 |
| Bacteria | Myxococc | Polyangia | Polyangial | Polyangiac | Sorangium | 10 | 0 | 0 |
| Bacteria | Actinobact | Actinobact | Frankiales | Sporichthy | Sporichthy | 10 | 0 | 0 |
| Bacteria | Proteobact | Alphaprote | Rhodospiri | Rhodospiri | Candidatus | 10 | 0 | 0 |
| Bacteria | SAR324_c | SAR324_c | SAR324_c | SAR324_c | SAR324_c | 10 | 0 | 0 |
| Bacteria | Desulfobac | Desulfobul | Desulfobul | Desulfobul | __ | 0 | 10 | 0 |
| Bacteria | Bacteroido | Rhodother | Rhodother | Rhodother | uncultured | 0 | 10 | 0 |
| Bacteria | Planctomy | Planctomy | Gemmatale | Gemmatac | __ | 0 | 17 | 0 |
| Bacteria | Bacteroido | Bacteroidia | Bacteroida | Prolixibact | uncultured | 0 | 10 | 0 |
| Bacteria | Firmicutes | Clostridia | Lachnospi | Lachnospi | Natranaero | 0 | 0 | 0 |
| Bacteria | Firmicutes | Bacilli | __ | __ | __ | 0 | 3 | 0 |

| | | | | | | | | |
|---|---|---|---|---|---|---|---|---|
| Bacteria | Firmicutes | Bacilli | Thermoact | Thermoact | Shimazuel | 0 | 0 | 0 |
| Bacteria | Acidobacte | Acidobacte | Acidobacte | __ | __ | 0 | 0 | 0 |
| Bacteria | Proteobact | Alphaprote | Elsterales | uncultured | uncultured | 6 | 0 | 0 |
| Bacteria | Firmicutes | Bacilli | Bacillales | Planococca | Rummeliib | 0 | 0 | 0 |
| Bacteria | Chloroflex | Dehalococ | FW22 | FW22 | FW22 | 0 | 0 | 0 |
| Bacteria | Desulfobac | Desulfobac | Desulfobac | Desulfobac | Desulfatife | 0 | 0 | 0 |
| Bacteria | Proteobact | Gammapro | Pseudomo | Spongiibac | BD1-7_cla | 0 | 0 | 0 |
| Bacteria | Bdellovibr | Bdellovibr | Bacteriovo | Bacteriovo | Bacteriovo | 9 | 0 | 0 |
| Bacteria | Myxococc | Polyangia | __ | __ | __ | 9 | 0 | 0 |
| Bacteria | Chloroflex | Ktedonoba | MVP-21 | MVP-21 | MVP-21 | 9 | 7 | 0 |
| Bacteria | Myxococc | Polyangia | Polyangial | Polyangiac | uncultured | 0 | 0 | 0 |
| Bacteria | Zixibacteri | Zixibacteri | Zixibacteri | Zixibacteri | Zixibacteri | 0 | 0 | 0 |
| Bacteria | Proteobact | Alphaprote | Parvibacul | Parvibacul | Parvibacul | 0 | 0 | 0 |
| Bacteria | Firmicutes | Bacilli | Bacillales | Planococca | Domibacil | 0 | 0 | 0 |
| Bacteria | Verrucomi | Verrucomi | Verrucomi | Verrucomi | uncultured | 0 | 0 | 0 |
| Bacteria | Proteobact | Gammapro | Gammapro | Unknown_ | Candidatus | 0 | 0 | 0 |
| Bacteria | Verrucomi | Verrucomi | Pedosphae | Pedosphae | Ellin517 | 0 | 0 | 0 |
| Bacteria | Firmicutes | Bacilli | Bacillales | Bacillacea | uncultured | 0 | 0 | 0 |
| Bacteria | Acidobacte | Blastocate | Pyrinomon | Pyrinomon | RB41 | 0 | 0 | 0 |
| Bacteria | Firmicutes | Negativicu | Veillonella | Sporomusa | __ | 8 | 0 | 0 |
| Bacteria | Acidobacte | Subgroup_ | Subgroup_ | Subgroup_ | Subgroup_ | 8 | 0 | 0 |
| Bacteria | Bacteroido | Bacteroidia | Chitinopha | __ | __ | 0 | 0 | 0 |
| Bacteria | Acidobacte | Subgroup_ | Subgroup_ | Subgroup_ | Subgroup_ | 0 | 8 | 0 |
| Bacteria | WOR-1 | WOR-1 | WOR-1 | WOR-1 | WOR-1 | 0 | 8 | 0 |
| Bacteria | Proteobact | Alphaprote | Rhizobiale | Beijerincki | Microvirga | 0 | 0 | 0 |
| Bacteria | Firmicutes | Clostridia | Peptococca | Peptococca | uncultured | 0 | 0 | 0 |
| Bacteria | Actinobact | Actinobact | Propioniba | Propioniba | Propionici | 0 | 0 | 0 |
| Bacteria | Actinobact | Actinobact | Propioniba | Propioniba | Aestuariim | 0 | 0 | 0 |
| Bacteria | Firmicutes | Clostridia | Oscillospir | Hungateicl | HN-HF010 | 0 | 0 | 0 |
| Bacteria | Proteobact | Gammapro | Burkholde | Nitrosomo | __ | 1 | 0 | 0 |
| Bacteria | Proteobact | Gammapro | Burkholde | Rhodocycl | Azoarcus | 0 | 0 | 0 |
| Bacteria | Proteobact | Alphaprote | Rhizobiale | Beijerincki | Roseiarcus | 0 | 0 | 0 |
| Bacteria | Proteobact | Gammapro | Burkholde | Gallionella | Sideroxyda | 7 | 0 | 0 |
| Bacteria | Verrucomi | Chlamydia | Chlamydia | Criblamyd | Criblamyd | 0 | 7 | 0 |
| Bacteria | Bacteroido | Bacteroidia | Flavobacte | NS9_marin | NS9_marin | 0 | 0 | 0 |
| Bacteria | Acidobacte | Subgroup_ | Subgroup_ | Subgroup_ | Subgroup_ | 0 | 0 | 0 |
| Bacteria | Chloroflex | Anaeroline | ADurb.Bin | ADurb.Bin | ADurb.Bin | 0 | 0 | 0 |
| Bacteria | Firmicutes | Clostridia | Peptostrep | Peptostrep | Alkaliphilu | 0 | 0 | 0 |
| Bacteria | Firmicutes | uncultured | uncultured | uncultured | uncultured | 4 | 0 | 0 |
| Bacteria | Proteobact | Gammapro | Methyloco | Methyloco | Candidatus | 0 | 0 | 0 |
| Bacteria | Firmicutes | Bacilli | Bacillales | Bacillacea | Geobacillu | 6 | 0 | 0 |
| Bacteria | Planctomy | Planctomy | Pirellulales | Pirellulace | Bythopirel | 6 | 0 | 0 |
| Bacteria | TA06 | TA06 | TA06 | TA06 | TA06 | 6 | 0 | 0 |
| Bacteria | Myxococc | Polyangia | Polyangial | __ | __ | 0 | 6 | 0 |
| Bacteria | Proteobact | Gammapro | Methyloco | Methylom | pLW-20 | 0 | 6 | 0 |
| Bacteria | Chloroflex | Dehalococ | Dehalococ | Dehalococ | Dehalogen | 0 | 0 | 0 |
| Bacteria | Proteobact | Gammapro | Enterobact | Shewanella | Shewanella | 0 | 0 | 0 |
| Bacteria | Proteobact | Gammapro | Chromatia | Chromatia | __ | 0 | 0 | 0 |
| Bacteria | Proteobact | Alphaprote | Paracaedib | Paracaedib | Candidatus | 0 | 0 | 0 |
| Bacteria | Proteobact | Alphaprote | Rhizobiale | Xanthobac | Xanthobac | 0 | 0 | 0 |

| | | | | | | | | |
|---|---|---|---|---|---|---|---|---|
| Bacteria | Firmicutes | Syntrophor | Syntrophor | Syntrophor | uncultured | 0 | 0 | 0 |
| Bacteria | Fibrobacte | Fibrobacte | Fibrobacte | __ | __ | 0 | 0 | 0 |
| Bacteria | Firmicutes | Clostridia | Therminco | Therminco | Therminco | 0 | 0 | 0 |
| Bacteria | Verrucomi | Verrucomi | Methylacid | Methylacid | uncultured | 0 | 0 | 0 |
| Bacteria | Actinobact | Actinobact | Actinomyc | Actinomyc | Actinomyc | 0 | 0 | 0 |
| Archaea | __ | __ | __ | __ | __ | 0 | 0 | 0 |
| Bacteria | Bacteroido | Bacteroidia | Bacteroida | Paludibact | Paludibact | 0 | 0 | 0 |
| Bacteria | Gemmatim | AKAU404 | AKAU404 | AKAU404 | AKAU404 | 0 | 0 | 0 |
| Bacteria | Firmicutes | Negativicu | Veillonella | Sporomusa | Anaerospo | 5 | 0 | 0 |
| Bacteria | Firmicutes | Clostridia | Lachnospi | Lachnospi | Epulopisci | 5 | 0 | 0 |
| Bacteria | Myxococc | __ | __ | __ | __ | 5 | 0 | 0 |
| Bacteria | Proteobact | Gammapro | Burkholde | Alcaligena | __ | 5 | 0 | 0 |
| Bacteria | Acidobacte | Thermoana | Thermoana | Thermoana | Thermoana | 5 | 0 | 0 |
| Bacteria | Firmicutes | Bacilli | Lactobacil | Streptococ | Streptococ | 0 | 0 | 5 |
| Bacteria | Proteobact | Alphaprote | Azospirilla | Azospirilla | Stella | 0 | 0 | 0 |
| Bacteria | Proteobact | Gammapro | Xanthomo | Xanthomo | uncultured | 0 | 0 | 0 |
| Bacteria | Planctomy | Phycisphae | __ | __ | __ | 0 | 0 | 0 |
| Bacteria | Proteobact | Gammapro | Gammapro | Unknown_ | Acidibacte | 0 | 0 | 0 |
| Bacteria | Bdellovibr | Oligoflexia | Oligoflexa | Oligoflexa | Oligoflexu | 0 | 0 | 0 |
| Bacteria | Firmicutes | Clostridia | Lachnospi | Lachnospi | Blautia | 0 | 0 | 0 |
| Bacteria | Fusobacter | Fusobacter | Fusobacter | Leptotrichi | Hypnocycl | 0 | 4 | 0 |
| Bacteria | Planctomy | Planctomy | Gemmatale | Gemmatac | Fimbriiglo | 0 | 0 | 0 |
| Bacteria | Planctomy | Phycisphae | Tepidispha | Tepidispha | Tepidispha | 0 | 0 | 0 |
| Bacteria | Methylomi | Methylomi | Rokubacte | Rokubacte | Rokubacte | 0 | 0 | 0 |
| Bacteria | Bacteroido | Bacteroidia | SM1A07 | SM1A07 | SM1A07 | 0 | 0 | 0 |
| Bacteria | Firmicutes | Bacilli | RF39 | RF39 | RF39 | 0 | 0 | 0 |
| Bacteria | Bacteroido | Bacteroidia | Bacteroida | Rikenellac | Alistipes | 0 | 0 | 0 |
| Eukaryota | Ascomyco | __ | __ | __ | __ | 0 | 0 | 0 |
| Bacteria | Firmicutes | Clostridia | Eubacteria | Garciellace | Irregularib | 0 | 0 | 0 |
| Bacteria | Verrucomi | Verrucomi | Chthoniob | Chthoniob | Chthoniob | 0 | 0 | 0 |
| Bacteria | Elusimicro | Elusimicro | FCPU453 | FCPU453 | FCPU453 | 0 | 0 | 0 |
| Bacteria | Planctomy | Phycisphae | Phycisphae | Phycisphae | AKYG587 | 0 | 0 | 0 |
| Bacteria | Bacteroido | Bacteroidia | Bacteroida | Muribacula | Muribacula | 0 | 0 | 0 |
| Bacteria | Firmicutes | Bacilli | Aneurinib | Aneurinib | Aneurinib | 0 | 0 | 0 |
| Eukaryota | Ciliophora | Intramacro | Conthreep | Colpodea | Bromelioth | 0 | 0 | 0 |
| Bacteria | Bacteroido | Bacteroidia | Chitinopha | Chitinopha | uncultured | 0 | 0 | 0 |
| Bacteria | Planctomy | Planctomy | Planctomy | Schlesneria | __ | 0 | 0 | 0 |
| Bacteria | Acidobacte | Blastocatel | Blastocatel | Blastocatel | Stenotroph | 0 | 0 | 0 |
| Bacteria | Bacteroido | Bacteroidia | Bacteroida | Rikenellac | Rikenellac | 0 | 0 | 0 |
| Bacteria | Firmicutes | Bacilli | Lactobacil | Lactobacil | Ligilactoba | 0 | 0 | 0 |
| Bacteria | Acidobacte | Holophaga | Holophaga | Holophaga | uncultured | 0 | 0 | 0 |
| Bacteria | Bacteroido | Bacteroidia | Sphingoba | Z4MB62 | Z4MB62 | 0 | 1 | 0 |

| 0.5%-$CH_4$ 2-µM-$Cu^{2+}$ | | | | 0.5%-$CH_4$ | | | | |
|---|---|---|---|---|---|---|---|---|
| **R20** | **R27** | **R34** | **CB** | **R18** | **R27** | **R39** | **R41** | **CB** |
| 18641 | 17162 | 15947 | 850 | 2378 | 15241 | 62042 | 54627 | 352 |
| 290 | 36 | 57 | 0 | 10257 | 5120 | 260 | 500 | 0 |
| 2165 | 3332 | 4925 | 159 | 3632 | 7946 | 13296 | 12125 | 65 |
| 0 | 0 | 0 | 54 | 0 | 0 | 0 | 0 | 38 |
| 0 | 0 | 0 | 20 | 136 | 45 | 0 | 0 | 0 |
| 143 | 7 | 6 | 0 | 2331 | 2136 | 239 | 458 | 0 |
| 94 | 24 | 63 | 54 | 250 | 142 | 9 | 195 | 0 |
| 3738 | 3843 | 963 | 250 | 171 | 337 | 181 | 210 | 102 |
| 118 | 75 | 54 | 0 | 17 | 16 | 0 | 10 | 8 |
| 35 | 5 | 0 | 0 | 0 | 0 | 0 | 0 | 0 |
| 107 | 36 | 26 | 0 | 1034 | 1241 | 235 | 385 | 128 |
| 0 | 0 | 0 | 668 | 6533 | 2717 | 935 | 880 | 65 |
| 1181 | 14 | 0 | 12 | 159 | 72 | 0 | 0 | 0 |
| 221 | 0 | 4 | 784 | 602 | 724 | 114 | 268 | 1286 |
| 242 | 58 | 0 | 0 | 0 | 0 | 0 | 0 | 155 |
| 163 | 158 | 224 | 229 | 255 | 1172 | 544 | 739 | 20 |
| 130 | 34 | 26 | 788 | 630 | 336 | 124 | 114 | 458 |
| 180 | 41 | 0 | 100 | 429 | 707 | 426 | 474 | 0 |
| 12 | 0 | 0 | 22 | 1502 | 100 | 0 | 0 | 0 |
| 373 | 118 | 154 | 991 | 729 | 1199 | 430 | 686 | 420 |
| 205 | 23 | 0 | 339 | 0 | 0 | 0 | 0 | 11 |
| 0 | 0 | 0 | 152 | 70 | 16 | 0 | 0 | 1288 |
| 241 | 128 | 102 | 209 | 1313 | 643 | 389 | 384 | 14 |
| 185 | 77 | 0 | 583 | 152 | 139 | 28 | 28 | 273 |
| 0 | 0 | 0 | 0 | 35 | 0 | 8 | 0 | 0 |
| 49 | 13 | 0 | 205 | 71 | 83 | 71 | 88 | 44 |
| 0 | 0 | 0 | 882 | 0 | 0 | 0 | 0 | 1556 |
| 101 | 97 | 56 | 176 | 0 | 149 | 11 | 51 | 373 |
| 0 | 0 | 0 | 0 | 22 | 65 | 36 | 43 | 0 |
| 141 | 34 | 81 | 374 | 288 | 163 | 0 | 24 | 305 |
| 0 | 0 | 0 | 2146 | 497 | 487 | 215 | 192 | 145 |
| 0 | 0 | 0 | 161 | 0 | 0 | 0 | 0 | 170 |
| 181 | 11 | 32 | 0 | 0 | 0 | 6 | 10 | 11 |
| 25 | 35 | 13 | 21 | 17 | 0 | 182 | 211 | 0 |
| 0 | 0 | 0 | 1857 | 0 | 0 | 0 | 8 | 326 |
| 0 | 0 | 0 | 461 | 0 | 0 | 0 | 0 | 1724 |
| 182 | 49 | 63 | 641 | 888 | 569 | 255 | 814 | 0 |
| 2 | 0 | 4 | 1 | 0 | 7 | 0 | 0 | 0 |
| 0 | 0 | 0 | 741 | 272 | 547 | 219 | 348 | 11 |
| 0 | 0 | 0 | 139 | 355 | 40 | 22 | 24 | 80 |
| 0 | 0 | 0 | 743 | 0 | 0 | 0 | 0 | 490 |
| 0 | 0 | 0 | 1879 | 26 | 55 | 32 | 44 | 185 |
| 0 | 0 | 0 | 0 | 0 | 0 | 0 | 0 | 0 |
| 0 | 0 | 0 | 1350 | 0 | 0 | 0 | 0 | 79 |
| 0 | 0 | 11 | 1274 | 15 | 245 | 64 | 17 | 639 |
| 432 | 128 | 141 | 203 | 157 | 173 | 79 | 177 | 0 |

| | | | | | | | | |
|---|---|---|---|---|---|---|---|---|
| 28 | 43 | 50 | 266 | 13 | 34 | 105 | 215 | 53 |
| 70 | 26 | 65 | 80 | 20 | 182 | 269 | 404 | 0 |
| 0 | 0 | 0 | 17 | 0 | 0 | 0 | 0 | 337 |
| 0 | 0 | 0 | 1240 | 563 | 340 | 164 | 154 | 31 |
| 0 | 0 | 0 | 2184 | 0 | 0 | 0 | 0 | 742 |
| 161 | 19 | 0 | 0 | 98 | 196 | 112 | 76 | 0 |
| 0 | 16 | 0 | 916 | 10 | 0 | 0 | 0 | 1186 |
| 0 | 0 | 0 | 165 | 56 | 906 | 757 | 764 | 0 |
| 0 | 0 | 0 | 11 | 652 | 292 | 66 | 56 | 0 |
| 25 | 0 | 0 | 0 | 62 | 29 | 0 | 0 | 0 |
| 0 | 0 | 0 | 523 | 154 | 68 | 0 | 15 | 0 |
| 69 | 39 | 15 | 15 | 0 | 0 | 0 | 0 | 0 |
| 174 | 71 | 91 | 658 | 43 | 27 | 14 | 44 | 270 |
| 211 | 58 | 23 | 140 | 916 | 458 | 0 | 56 | 28 |
| 42 | 3 | 28 | 0 | 60 | 927 | 619 | 716 | 0 |
| 119 | 19 | 16 | 258 | 79 | 84 | 0 | 52 | 123 |
| 0 | 0 | 0 | 397 | 16 | 0 | 0 | 0 | 348 |
| 0 | 0 | 0 | 31 | 719 | 237 | 57 | 77 | 0 |
| 0 | 0 | 0 | 0 | 0 | 0 | 0 | 0 | 0 |
| 114 | 29 | 12 | 303 | 30 | 122 | 0 | 38 | 651 |
| 0 | 0 | 0 | 1119 | 160 | 192 | 176 | 125 | 288 |
| 0 | 0 | 0 | 20 | 2 | 31 | 24 | 32 | 0 |
| 0 | 0 | 0 | 240 | 0 | 0 | 0 | 0 | 422 |
| 127 | 23 | 16 | 175 | 601 | 270 | 115 | 117 | 26 |
| 0 | 0 | 0 | 164 | 0 | 0 | 0 | 0 | 265 |
| 999 | 202 | 126 | 27 | 362 | 157 | 27 | 22 | 0 |
| 17 | 12 | 100 | 0 | 0 | 169 | 168 | 150 | 0 |
| 0 | 0 | 0 | 377 | 0 | 0 | 0 | 0 | 183 |
| 411 | 0 | 347 | 0 | 85 | 41 | 0 | 0 | 0 |
| 0 | 0 | 0 | 189 | 6 | 0 | 0 | 0 | 0 |
| 17 | 0 | 0 | 5 | 0 | 0 | 0 | 0 | 0 |
| 0 | 0 | 19 | 0 | 37 | 625 | 7 | 111 | 0 |
| 0 | 0 | 0 | 542 | 0 | 0 | 24 | 37 | 148 |
| 0 | 0 | 0 | 8 | 0 | 0 | 0 | 0 | 0 |
| 0 | 0 | 0 | 335 | 0 | 0 | 0 | 0 | 486 |
| 0 | 0 | 0 | 171 | 0 | 904 | 346 | 403 | 0 |
| 0 | 0 | 0 | 39 | 313 | 131 | 79 | 86 | 7 |
| 11 | 0 | 24 | 26 | 375 | 626 | 51 | 88 | 273 |
| 41 | 0 | 0 | 45 | 4 | 0 | 0 | 0 | 268 |
| 0 | 9 | 8 | 178 | 7 | 6 | 0 | 0 | 243 |
| 9 | 0 | 0 | 37 | 0 | 21 | 0 | 11 | 632 |
| 0 | 0 | 0 | 2 | 0 | 0 | 0 | 0 | 24 |
| 49 | 25 | 3 | 876 | 30 | 37 | 55 | 181 | 229 |
| 0 | 0 | 0 | 273 | 92 | 118 | 100 | 85 | 0 |
| 75 | 0 | 0 | 0 | 0 | 56 | 0 | 0 | 0 |
| 0 | 0 | 0 | 27 | 15 | 779 | 68 | 80 | 0 |
| 0 | 0 | 0 | 621 | 0 | 28 | 0 | 0 | 597 |
| 14 | 129 | 534 | 584 | 0 | 22 | 20 | 49 | 202 |
| 0 | 0 | 0 | 528 | 0 | 0 | 0 | 0 | 0 |
| 0 | 0 | 0 | 290 | 115 | 33 | 18 | 0 | 0 |

| | | | | | | | | |
|---|---|---|---|---|---|---|---|---|
| 0 | 0 | 0 | 359 | 0 | 0 | 0 | 0 | 39 |
| 0 | 0 | 0 | 105 | 150 | 667 | 175 | 219 | 29 |
| 0 | 0 | 0 | 182 | 0 | 180 | 155 | 226 | 0 |
| 0 | 0 | 0 | 845 | 9 | 35 | 37 | 62 | 140 |
| 0 | 0 | 0 | 899 | 25 | 50 | 35 | 40 | 200 |
| 0 | 0 | 0 | 0 | 0 | 0 | 0 | 0 | 0 |
| 54 | 3 | 52 | 190 | 0 | 0 | 300 | 230 | 75 |
| 0 | 0 | 6 | 245 | 23 | 220 | 186 | 139 | 0 |
| 0 | 0 | 0 | 0 | 0 | 148 | 89 | 281 | 0 |
| 0 | 0 | 11 | 27 | 147 | 374 | 43 | 81 | 0 |
| 10 | 351 | 164 | 522 | 197 | 114 | 0 | 44 | 165 |
| 17 | 6 | 0 | 137 | 58 | 291 | 101 | 128 | 0 |
| 0 | 0 | 0 | 199 | 179 | 207 | 129 | 180 | 100 |
| 0 | 0 | 0 | 277 | 202 | 112 | 31 | 26 | 0 |
| 0 | 0 | 0 | 99 | 0 | 0 | 0 | 0 | 0 |
| 0 | 0 | 0 | 191 | 53 | 78 | 53 | 56 | 119 |
| 0 | 0 | 0 | 326 | 8 | 0 | 0 | 0 | 6 |
| 0 | 0 | 0 | 0 | 166 | 38 | 237 | 184 | 0 |
| 0 | 0 | 0 | 184 | 0 | 0 | 0 | 0 | 0 |
| 0 | 0 | 0 | 121 | 31 | 64 | 16 | 15 | 0 |
| 6 | 0 | 146 | 196 | 161 | 143 | 100 | 66 | 83 |
| 0 | 0 | 0 | 258 | 0 | 0 | 0 | 0 | 27 |
| 33 | 0 | 0 | 13 | 108 | 226 | 61 | 127 | 0 |
| 0 | 0 | 0 | 0 | 46 | 31 | 0 | 0 | 0 |
| 0 | 16 | 0 | 147 | 0 | 0 | 0 | 0 | 0 |
| 0 | 0 | 0 | 475 | 0 | 0 | 0 | 4 | 0 |
| 0 | 0 | 0 | 7 | 0 | 28 | 190 | 218 | 0 |
| 0 | 0 | 0 | 501 | 14 | 5 | 0 | 0 | 665 |
| 0 | 0 | 0 | 0 | 0 | 0 | 0 | 0 | 0 |
| 0 | 0 | 0 | 157 | 0 | 0 | 0 | 0 | 168 |
| 0 | 0 | 0 | 111 | 0 | 0 | 0 | 0 | 105 |
| 0 | 0 | 0 | 608 | 16 | 32 | 0 | 5 | 262 |
| 9 | 0 | 209 | 244 | 63 | 11 | 0 | 31 | 0 |
| 0 | 0 | 0 | 10 | 16 | 82 | 20 | 42 | 0 |
| 0 | 0 | 0 | 0 | 0 | 0 | 0 | 0 | 0 |
| 0 | 0 | 0 | 117 | 0 | 0 | 0 | 0 | 159 |
| 0 | 0 | 0 | 174 | 0 | 0 | 0 | 0 | 274 |
| 0 | 0 | 0 | 389 | 4 | 0 | 0 | 0 | 28 |
| 304 | 148 | 24 | 996 | 9 | 25 | 0 | 26 | 124 |
| 9 | 0 | 0 | 75 | 57 | 15 | 0 | 11 | 0 |
| 0 | 0 | 0 | 72 | 1089 | 33 | 0 | 9 | 0 |
| 0 | 0 | 0 | 0 | 11 | 108 | 132 | 142 | 0 |
| 0 | 0 | 0 | 214 | 52 | 22 | 0 | 0 | 194 |
| 151 | 16 | 56 | 77 | 428 | 338 | 445 | 575 | 389 |
| 0 | 0 | 0 | 318 | 212 | 384 | 73 | 57 | 0 |
| 6 | 0 | 0 | 985 | 122 | 76 | 38 | 49 | 1956 |
| 11 | 0 | 34 | 110 | 0 | 0 | 0 | 0 | 255 |
| 0 | 0 | 0 | 80 | 0 | 0 | 0 | 0 | 539 |
| 0 | 0 | 0 | 224 | 0 | 0 | 0 | 0 | 0 |
| 0 | 0 | 0 | 0 | 0 | 0 | 0 | 0 | 178 |

| | | | | | | | | |
|---|---|---|---|---|---|---|---|---|
| 0 | 0 | 0 | 696 | 0 | 0 | 0 | 0 | 598 |
| 0 | 0 | 4 | 134 | 114 | 442 | 136 | 163 | 22 |
| 0 | 0 | 0 | 390 | 0 | 0 | 0 | 0 | 73 |
| 0 | 0 | 0 | 174 | 0 | 30 | 157 | 156 | 75 |
| 25 | 0 | 0 | 0 | 0 | 0 | 0 | 0 | 0 |
| 0 | 0 | 0 | 226 | 0 | 0 | 0 | 0 | 10 |
| 0 | 0 | 0 | 244 | 32 | 168 | 140 | 121 | 21 |
| 35 | 0 | 0 | 8 | 0 | 0 | 0 | 0 | 0 |
| 0 | 0 | 0 | 32 | 16 | 32 | 90 | 153 | 0 |
| 0 | 0 | 0 | 7 | 0 | 0 | 0 | 0 | 47 |
| 0 | 0 | 0 | 4 | 0 | 0 | 0 | 0 | 200 |
| 0 | 0 | 0 | 0 | 0 | 0 | 0 | 0 | 0 |
| 0 | 0 | 0 | 0 | 0 | 0 | 0 | 0 | 0 |
| 0 | 0 | 0 | 407 | 19 | 71 | 158 | 146 | 41 |
| 0 | 0 | 0 | 167 | 30 | 62 | 76 | 102 | 0 |
| 0 | 0 | 0 | 247 | 0 | 0 | 0 | 0 | 46 |
| 0 | 0 | 0 | 311 | 0 | 0 | 0 | 0 | 96 |
| 0 | 0 | 0 | 73 | 0 | 0 | 0 | 0 | 0 |
| 0 | 0 | 0 | 356 | 6 | 179 | 65 | 53 | 60 |
| 0 | 0 | 0 | 0 | 0 | 0 | 0 | 0 | 0 |
| 0 | 0 | 0 | 195 | 0 | 0 | 0 | 0 | 0 |
| 0 | 0 | 0 | 30 | 4 | 130 | 107 | 80 | 0 |
| 0 | 0 | 0 | 0 | 0 | 0 | 0 | 0 | 0 |
| 0 | 0 | 0 | 239 | 0 | 17 | 14 | 0 | 0 |
| 0 | 0 | 0 | 611 | 0 | 0 | 0 | 0 | 0 |
| 0 | 0 | 0 | 141 | 0 | 13 | 20 | 23 | 0 |
| 0 | 0 | 0 | 179 | 0 | 0 | 0 | 0 | 31 |
| 0 | 0 | 0 | 0 | 0 | 0 | 0 | 0 | 0 |
| 0 | 0 | 0 | 211 | 0 | 0 | 0 | 0 | 170 |
| 0 | 0 | 0 | 156 | 0 | 0 | 0 | 0 | 0 |
| 0 | 0 | 0 | 316 | 0 | 59 | 51 | 56 | 0 |
| 0 | 0 | 6 | 41 | 52 | 119 | 173 | 243 | 0 |
| 0 | 0 | 0 | 662 | 0 | 0 | 0 | 0 | 34 |
| 0 | 0 | 0 | 282 | 0 | 0 | 0 | 0 | 75 |
| 0 | 0 | 0 | 44 | 56 | 71 | 14 | 0 | 0 |
| 0 | 0 | 0 | 14 | 0 | 0 | 0 | 0 | 0 |
| 0 | 0 | 0 | 0 | 47 | 29 | 0 | 8 | 0 |
| 0 | 0 | 0 | 69 | 0 | 88 | 67 | 148 | 35 |
| 0 | 0 | 0 | 197 | 0 | 0 | 0 | 0 | 0 |
| 0 | 0 | 0 | 135 | 0 | 0 | 0 | 0 | 0 |
| 85 | 31 | 16 | 10 | 0 | 0 | 0 | 0 | 51 |
| 35 | 16 | 11 | 42 | 30 | 11 | 0 | 10 | 108 |
| 55 | 0 | 0 | 0 | 53 | 107 | 19 | 19 | 0 |
| 0 | 0 | 0 | 156 | 0 | 0 | 0 | 0 | 0 |
| 0 | 0 | 0 | 179 | 0 | 0 | 0 | 0 | 237 |
| 0 | 0 | 0 | 242 | 79 | 18 | 0 | 0 | 190 |
| 0 | 0 | 0 | 111 | 0 | 0 | 0 | 0 | 0 |
| 0 | 0 | 0 | 32 | 7 | 0 | 0 | 0 | 0 |
| 0 | 0 | 0 | 395 | 0 | 0 | 0 | 0 | 17 |
| 0 | 0 | 0 | 91 | 0 | 80 | 137 | 111 | 0 |

| | | | | | | | | |
|---|---|---|---|---|---|---|---|---|
| 0 | 0 | 0 | 119 | 0 | 20 | 9 | 36 | 0 |
| 0 | 0 | 0 | 0 | 0 | 0 | 0 | 0 | 0 |
| 23 | 3 | 0 | 73 | 0 | 0 | 0 | 9 | 12 |
| 0 | 0 | 0 | 338 | 0 | 0 | 0 | 0 | 16 |
| 0 | 0 | 0 | 55 | 0 | 0 | 0 | 0 | 12 |
| 0 | 0 | 0 | 0 | 0 | 0 | 0 | 0 | 0 |
| 43 | 99 | 0 | 121 | 0 | 0 | 0 | 0 | 0 |
| 0 | 0 | 0 | 103 | 0 | 0 | 0 | 0 | 0 |
| 0 | 0 | 0 | 0 | 0 | 67 | 72 | 70 | 0 |
| 0 | 6 | 16 | 8 | 24 | 115 | 140 | 126 | 0 |
| 0 | 0 | 0 | 116 | 0 | 8 | 0 | 0 | 0 |
| 13 | 32 | 41 | 304 | 34 | 10 | 0 | 0 | 37 |
| 0 | 0 | 0 | 320 | 0 | 0 | 0 | 0 | 0 |
| 0 | 0 | 0 | 143 | 0 | 0 | 0 | 0 | 0 |
| 0 | 0 | 0 | 0 | 0 | 70 | 0 | 0 | 0 |
| 0 | 0 | 0 | 13 | 0 | 0 | 0 | 0 | 0 |
| 0 | 0 | 0 | 157 | 0 | 0 | 14 | 0 | 23 |
| 0 | 0 | 0 | 109 | 0 | 0 | 0 | 0 | 0 |
| 0 | 0 | 0 | 70 | 0 | 0 | 0 | 0 | 139 |
| 0 | 0 | 0 | 0 | 5 | 0 | 0 | 0 | 292 |
| 0 | 0 | 0 | 0 | 0 | 0 | 0 | 0 | 0 |
| 0 | 0 | 0 | 0 | 0 | 0 | 0 | 0 | 0 |
| 0 | 0 | 0 | 317 | 0 | 0 | 0 | 0 | 186 |
| 0 | 0 | 0 | 0 | 0 | 0 | 0 | 0 | 0 |
| 0 | 0 | 0 | 0 | 0 | 0 | 0 | 0 | 0 |
| 0 | 0 | 0 | 17 | 0 | 0 | 0 | 0 | 0 |
| 0 | 0 | 0 | 0 | 0 | 0 | 0 | 0 | 0 |
| 131 | 29 | 34 | 0 | 0 | 0 | 0 | 0 | 0 |
| 28 | 0 | 0 | 0 | 0 | 0 | 0 | 0 | 0 |
| 0 | 0 | 0 | 0 | 17 | 91 | 45 | 30 | 0 |
| 0 | 0 | 0 | 8 | 0 | 7 | 12 | 31 | 57 |
| 0 | 0 | 0 | 89 | 117 | 91 | 49 | 41 | 4 |
| 0 | 0 | 0 | 80 | 0 | 0 | 0 | 0 | 0 |
| 0 | 0 | 0 | 49 | 0 | 0 | 0 | 0 | 0 |
| 121 | 0 | 0 | 53 | 0 | 0 | 0 | 0 | 0 |
| 0 | 0 | 0 | 24 | 0 | 0 | 0 | 0 | 0 |
| 0 | 0 | 0 | 92 | 0 | 0 | 0 | 0 | 0 |
| 0 | 0 | 0 | 212 | 0 | 0 | 0 | 0 | 0 |
| 0 | 0 | 0 | 0 | 56 | 31 | 8 | 6 | 0 |
| 0 | 0 | 0 | 79 | 86 | 73 | 6 | 0 | 0 |
| 0 | 0 | 0 | 215 | 0 | 0 | 0 | 0 | 0 |
| 0 | 0 | 0 | 39 | 0 | 0 | 0 | 0 | 0 |
| 49 | 0 | 0 | 0 | 0 | 0 | 0 | 0 | 0 |
| 0 | 0 | 0 | 17 | 6 | 0 | 0 | 0 | 12 |
| 0 | 0 | 0 | 13 | 0 | 0 | 80 | 80 | 0 |
| 0 | 0 | 0 | 292 | 0 | 0 | 0 | 0 | 12 |
| 0 | 0 | 0 | 146 | 0 | 0 | 0 | 0 | 56 |
| 0 | 0 | 0 | 209 | 0 | 0 | 0 | 0 | 0 |
| 0 | 0 | 0 | 50 | 0 | 0 | 0 | 0 | 0 |
| 0 | 0 | 0 | 60 | 0 | 0 | 0 | 0 | 27 |

| | | | | | | | | |
|---|---|---|---|---|---|---|---|---|
| 0 | 0 | 0 | 61 | 22 | 0 | 0 | 0 | 0 |
| 0 | 0 | 0 | 0 | 0 | 0 | 0 | 0 | 0 |
| 0 | 0 | 0 | 33 | 0 | 0 | 0 | 0 | 0 |
| 0 | 0 | 0 | 22 | 40 | 8 | 0 | 0 | 63 |
| 0 | 0 | 0 | 746 | 0 | 0 | 0 | 0 | 0 |
| 0 | 0 | 0 | 14 | 0 | 0 | 0 | 17 | 0 |
| 0 | 0 | 0 | 49 | 0 | 0 | 0 | 0 | 0 |
| 0 | 0 | 0 | 188 | 20 | 15 | 0 | 55 | 0 |
| 0 | 0 | 0 | 15 | 0 | 0 | 0 | 0 | 0 |
| 0 | 0 | 0 | 160 | 10 | 0 | 0 | 0 | 0 |
| 0 | 0 | 0 | 38 | 0 | 0 | 0 | 0 | 0 |
| 0 | 0 | 0 | 0 | 43 | 21 | 0 | 18 | 0 |
| 0 | 0 | 0 | 106 | 0 | 0 | 0 | 0 | 0 |
| 0 | 0 | 0 | 329 | 0 | 0 | 0 | 0 | 371 |
| 0 | 0 | 0 | 17 | 0 | 0 | 0 | 0 | 0 |
| 0 | 0 | 0 | 200 | 0 | 0 | 0 | 0 | 119 |
| 0 | 0 | 0 | 19 | 0 | 0 | 0 | 0 | 16 |
| 0 | 0 | 0 | 76 | 0 | 0 | 0 | 0 | 0 |
| 0 | 0 | 0 | 0 | 0 | 0 | 0 | 0 | 10 |
| 0 | 0 | 0 | 53 | 0 | 0 | 0 | 0 | 17 |
| 0 | 0 | 0 | 78 | 0 | 6 | 14 | 30 | 7 |
| 0 | 0 | 0 | 0 | 0 | 0 | 0 | 0 | 0 |
| 0 | 0 | 0 | 0 | 0 | 0 | 0 | 0 | 0 |
| 15 | 0 | 0 | 0 | 23 | 13 | 16 | 21 | 0 |
| 0 | 0 | 0 | 0 | 0 | 0 | 0 | 0 | 0 |
| 0 | 0 | 0 | 38 | 0 | 0 | 0 | 0 | 0 |
| 0 | 0 | 0 | 0 | 0 | 0 | 0 | 0 | 0 |
| 0 | 0 | 0 | 49 | 0 | 0 | 0 | 0 | 0 |
| 0 | 0 | 0 | 9 | 0 | 0 | 0 | 0 | 14 |
| 0 | 0 | 0 | 102 | 12 | 14 | 0 | 0 | 28 |
| 0 | 0 | 0 | 424 | 3 | 0 | 0 | 3 | 0 |
| 0 | 0 | 0 | 26 | 0 | 0 | 0 | 0 | 0 |
| 0 | 3 | 88 | 20 | 1 | 11 | 15 | 9 | 0 |
| 0 | 0 | 0 | 0 | 0 | 0 | 0 | 0 | 0 |
| 0 | 9 | 4 | 59 | 0 | 0 | 0 | 20 | 0 |
| 0 | 0 | 0 | 0 | 0 | 0 | 0 | 0 | 0 |
| 0 | 0 | 0 | 0 | 0 | 0 | 0 | 0 | 0 |
| 0 | 0 | 0 | 68 | 0 | 0 | 0 | 0 | 0 |
| 0 | 0 | 0 | 0 | 0 | 0 | 0 | 0 | 0 |
| 0 | 0 | 0 | 107 | 0 | 0 | 0 | 0 | 0 |
| 0 | 0 | 0 | 16 | 39 | 37 | 8 | 12 | 0 |
| 6 | 0 | 0 | 51 | 0 | 2 | 0 | 0 | 59 |
| 0 | 0 | 0 | 0 | 0 | 0 | 0 | 0 | 42 |
| 129 | 46 | 0 | 54 | 0 | 0 | 0 | 0 | 57 |
| 0 | 0 | 0 | 4 | 0 | 0 | 0 | 0 | 0 |
| 0 | 0 | 0 | 0 | 35 | 20 | 0 | 0 | 0 |
| 0 | 0 | 0 | 5 | 0 | 0 | 0 | 0 | 16 |
| 0 | 0 | 0 | 0 | 0 | 0 | 0 | 0 | 0 |
| 0 | 0 | 0 | 10 | 0 | 0 | 0 | 0 | 0 |
| 46 | 0 | 56 | 67 | 0 | 0 | 0 | 0 | 0 |

| | | | | | | | | |
|---|---|---|---|---|---|---|---|---|
| 0 | 0 | 0 | 289 | 21 | 8 | 0 | 0 | 62 |
| 0 | 0 | 0 | 43 | 0 | 0 | 0 | 0 | 0 |
| 0 | 0 | 0 | 55 | 0 | 0 | 0 | 0 | 9 |
| 0 | 0 | 0 | 99 | 0 | 0 | 0 | 0 | 0 |
| 0 | 0 | 0 | 16 | 0 | 0 | 0 | 0 | 0 |
| 0 | 0 | 0 | 0 | 0 | 0 | 0 | 0 | 14 |
| 0 | 0 | 0 | 0 | 0 | 0 | 0 | 0 | 0 |
| 0 | 0 | 0 | 0 | 0 | 0 | 0 | 0 | 0 |
| 0 | 0 | 0 | 8 | 0 | 0 | 0 | 0 | 106 |
| 0 | 0 | 0 | 97 | 0 | 0 | 0 | 0 | 6 |
| 0 | 0 | 0 | 56 | 0 | 0 | 0 | 5 | 0 |
| 0 | 0 | 0 | 0 | 41 | 54 | 19 | 38 | 0 |
| 0 | 0 | 0 | 339 | 0 | 0 | 0 | 0 | 0 |
| 0 | 0 | 0 | 0 | 0 | 0 | 0 | 0 | 0 |
| 0 | 0 | 0 | 160 | 26 | 0 | 0 | 0 | 0 |
| 0 | 0 | 0 | 0 | 0 | 0 | 0 | 0 | 35 |
| 0 | 0 | 0 | 50 | 0 | 0 | 0 | 0 | 0 |
| 0 | 0 | 0 | 68 | 0 | 0 | 0 | 0 | 35 |
| 0 | 0 | 0 | 141 | 0 | 0 | 0 | 0 | 26 |
| 0 | 0 | 0 | 134 | 0 | 0 | 0 | 0 | 0 |
| 0 | 0 | 0 | 0 | 0 | 0 | 0 | 0 | 0 |
| 0 | 0 | 0 | 58 | 0 | 0 | 0 | 0 | 0 |
| 0 | 0 | 0 | 19 | 0 | 0 | 0 | 0 | 0 |
| 0 | 0 | 0 | 22 | 0 | 0 | 0 | 0 | 0 |
| 0 | 0 | 0 | 0 | 0 | 0 | 0 | 0 | 0 |
| 0 | 0 | 0 | 0 | 0 | 0 | 0 | 0 | 0 |
| 0 | 0 | 0 | 0 | 0 | 0 | 0 | 0 | 0 |
| 0 | 0 | 0 | 0 | 0 | 0 | 0 | 0 | 0 |
| 0 | 0 | 0 | 14 | 0 | 0 | 0 | 0 | 0 |
| 0 | 0 | 0 | 59 | 4 | 0 | 0 | 0 | 0 |
| 0 | 0 | 0 | 0 | 0 | 0 | 0 | 0 | 0 |
| 0 | 0 | 0 | 0 | 0 | 0 | 0 | 0 | 0 |
| 0 | 0 | 0 | 21 | 0 | 0 | 0 | 0 | 17 |
| 0 | 0 | 0 | 95 | 101 | 15 | 3 | 6 | 7 |
| 0 | 0 | 0 | 200 | 0 | 0 | 0 | 0 | 0 |
| 0 | 0 | 0 | 6 | 0 | 0 | 0 | 0 | 3 |
| 0 | 0 | 0 | 34 | 0 | 0 | 0 | 0 | 32 |
| 0 | 0 | 7 | 92 | 0 | 2 | 7 | 0 | 11 |
| 0 | 0 | 0 | 16 | 0 | 0 | 0 | 0 | 5 |
| 0 | 0 | 0 | 117 | 0 | 0 | 0 | 0 | 11 |
| 0 | 0 | 0 | 0 | 0 | 0 | 0 | 0 | 0 |
| 0 | 0 | 0 | 35 | 0 | 0 | 0 | 0 | 0 |
| 0 | 0 | 0 | 51 | 0 | 0 | 0 | 0 | 0 |
| 0 | 0 | 0 | 61 | 10 | 0 | 0 | 0 | 0 |
| 0 | 0 | 0 | 0 | 40 | 42 | 0 | 0 | 0 |
| 0 | 0 | 0 | 0 | 4 | 0 | 0 | 9 | 0 |
| 0 | 0 | 0 | 54 | 0 | 0 | 0 | 0 | 13 |
| 0 | 0 | 0 | 0 | 0 | 0 | 0 | 0 | 25 |
| 0 | 0 | 0 | 95 | 13 | 8 | 4 | 24 | 0 |
| 0 | 0 | 0 | 0 | 0 | 0 | 0 | 0 | 0 |

| | | | | | | | | |
|---|---|---|---|---|---|---|---|---|
| 0 | 0 | 0 | 128 | 0 | 0 | 0 | 0 | 0 |
| 0 | 0 | 0 | 0 | 0 | 0 | 0 | 0 | 0 |
| 0 | 0 | 0 | 17 | 70 | 23 | 0 | 0 | 0 |
| 34 | 13 | 17 | 0 | 0 | 0 | 0 | 0 | 44 |
| 0 | 0 | 0 | 11 | 0 | 0 | 0 | 0 | 11 |
| 0 | 0 | 0 | 28 | 0 | 13 | 0 | 0 | 0 |
| 0 | 0 | 0 | 37 | 0 | 0 | 0 | 0 | 0 |
| 0 | 0 | 0 | 79 | 0 | 0 | 0 | 0 | 0 |
| 0 | 0 | 0 | 52 | 0 | 0 | 0 | 0 | 0 |
| 0 | 0 | 0 | 0 | 0 | 56 | 44 | 34 | 0 |
| 0 | 0 | 0 | 341 | 0 | 0 | 0 | 0 | 87 |
| 0 | 0 | 0 | 0 | 0 | 0 | 0 | 0 | 0 |
| 0 | 0 | 0 | 0 | 0 | 0 | 0 | 0 | 0 |
| 0 | 0 | 0 | 164 | 0 | 0 | 0 | 0 | 0 |
| 0 | 0 | 0 | 12 | 0 | 0 | 0 | 0 | 0 |
| 0 | 0 | 0 | 60 | 0 | 0 | 0 | 0 | 0 |
| 0 | 0 | 0 | 0 | 0 | 0 | 0 | 0 | 70 |
| 0 | 0 | 0 | 136 | 0 | 0 | 4 | 19 | 0 |
| 0 | 0 | 0 | 0 | 0 | 0 | 0 | 0 | 0 |
| 0 | 0 | 0 | 78 | 0 | 0 | 0 | 0 | 0 |
| 0 | 0 | 0 | 71 | 0 | 0 | 0 | 0 | 0 |
| 0 | 0 | 0 | 0 | 0 | 0 | 0 | 0 | 0 |
| 0 | 0 | 8 | 0 | 0 | 7 | 0 | 15 | 0 |
| 0 | 0 | 0 | 0 | 0 | 17 | 0 | 0 | 0 |
| 0 | 0 | 0 | 0 | 0 | 0 | 0 | 0 | 0 |
| 0 | 0 | 0 | 37 | 0 | 0 | 0 | 0 | 30 |
| 0 | 0 | 0 | 6 | 65 | 10 | 10 | 1 | 0 |
| 0 | 0 | 0 | 0 | 5 | 0 | 0 | 0 | 0 |
| 0 | 0 | 0 | 33 | 0 | 0 | 0 | 0 | 0 |
| 0 | 0 | 0 | 56 | 0 | 0 | 0 | 0 | 0 |
| 0 | 0 | 0 | 24 | 0 | 0 | 0 | 0 | 0 |
| 0 | 0 | 0 | 17 | 0 | 0 | 0 | 0 | 29 |
| 0 | 0 | 0 | 149 | 0 | 0 | 0 | 0 | 0 |
| 0 | 0 | 0 | 60 | 0 | 0 | 0 | 0 | 0 |
| 0 | 0 | 0 | 0 | 0 | 0 | 0 | 0 | 0 |
| 0 | 0 | 0 | 0 | 0 | 0 | 0 | 0 | 0 |
| 0 | 0 | 0 | 38 | 0 | 0 | 0 | 0 | 0 |
| 0 | 0 | 0 | 74 | 0 | 0 | 0 | 0 | 0 |
| 0 | 0 | 0 | 14 | 0 | 0 | 0 | 19 | 11 |
| 0 | 0 | 0 | 0 | 0 | 0 | 0 | 0 | 0 |
| 0 | 0 | 0 | 13 | 0 | 0 | 0 | 0 | 0 |
| 0 | 0 | 0 | 0 | 0 | 0 | 0 | 0 | 11 |
| 0 | 0 | 0 | 0 | 0 | 0 | 0 | 0 | 0 |
| 0 | 0 | 0 | 0 | 0 | 0 | 0 | 0 | 0 |
| 0 | 0 | 0 | 0 | 0 | 0 | 0 | 0 | 0 |
| 0 | 0 | 0 | 14 | 0 | 0 | 0 | 0 | 0 |
| 0 | 0 | 0 | 70 | 25 | 14 | 0 | 0 | 0 |
| 0 | 0 | 0 | 25 | 0 | 0 | 0 | 0 | 0 |
| 0 | 0 | 0 | 15 | 0 | 0 | 0 | 0 | 14 |
| 0 | 0 | 0 | 44 | 0 | 25 | 0 | 0 | 0 |

| | | | | | | | | |
|---|---|---|---|---|---|---|---|---|
| 5 | 0 | 0 | 0 | 0 | 0 | 0 | 0 | 0 |
| 0 | 0 | 9 | 0 | 0 | 0 | 0 | 0 | 0 |
| 0 | 0 | 0 | 0 | 0 | 0 | 0 | 0 | 0 |
| 0 | 0 | 0 | 0 | 0 | 0 | 0 | 0 | 0 |
| 0 | 0 | 0 | 0 | 0 | 0 | 0 | 0 | 0 |
| 0 | 0 | 0 | 23 | 0 | 0 | 0 | 0 | 0 |
| 0 | 0 | 0 | 0 | 0 | 0 | 0 | 0 | 0 |
| 0 | 0 | 0 | 26 | 41 | 22 | 0 | 0 | 0 |
| 0 | 0 | 0 | 18 | 0 | 7 | 39 | 50 | 0 |
| 0 | 0 | 0 | 75 | 0 | 0 | 0 | 0 | 0 |
| 0 | 0 | 0 | 80 | 0 | 0 | 0 | 0 | 65 |
| 0 | 0 | 0 | 45 | 0 | 0 | 0 | 0 | 0 |
| 0 | 0 | 0 | 12 | 0 | 0 | 0 | 0 | 0 |
| 0 | 0 | 0 | 38 | 0 | 0 | 0 | 0 | 0 |
| 0 | 0 | 0 | 31 | 0 | 0 | 0 | 0 | 0 |
| 0 | 0 | 0 | 57 | 0 | 0 | 0 | 0 | 0 |
| 0 | 0 | 0 | 0 | 0 | 0 | 0 | 0 | 0 |
| 0 | 0 | 0 | 0 | 0 | 0 | 0 | 0 | 28 |
| 0 | 0 | 0 | 79 | 0 | 0 | 0 | 0 | 10 |
| 0 | 0 | 0 | 24 | 0 | 0 | 0 | 0 | 0 |
| 0 | 0 | 0 | 69 | 6 | 0 | 0 | 0 | 0 |
| 0 | 0 | 0 | 0 | 0 | 0 | 0 | 0 | 0 |
| 0 | 0 | 0 | 0 | 0 | 0 | 0 | 0 | 0 |
| 0 | 0 | 0 | 61 | 0 | 0 | 0 | 0 | 0 |
| 0 | 0 | 0 | 0 | 0 | 0 | 0 | 0 | 0 |
| 0 | 0 | 0 | 0 | 0 | 0 | 0 | 0 | 0 |
| 0 | 0 | 0 | 0 | 0 | 0 | 0 | 0 | 58 |
| 0 | 0 | 0 | 0 | 0 | 0 | 0 | 0 | 0 |
| 0 | 0 | 0 | 0 | 0 | 0 | 0 | 0 | 0 |
| 0 | 0 | 0 | 0 | 0 | 0 | 0 | 0 | 57 |
| 0 | 0 | 0 | 73 | 0 | 0 | 0 | 0 | 0 |
| 0 | 0 | 0 | 0 | 0 | 0 | 0 | 0 | 0 |
| 0 | 0 | 0 | 56 | 0 | 0 | 0 | 0 | 0 |
| 0 | 0 | 0 | 0 | 0 | 0 | 0 | 0 | 94 |
| 0 | 0 | 0 | 0 | 0 | 0 | 0 | 0 | 0 |
| 0 | 0 | 0 | 0 | 0 | 0 | 0 | 0 | 0 |
| 0 | 0 | 0 | 0 | 0 | 0 | 0 | 0 | 0 |
| 0 | 0 | 0 | 0 | 0 | 0 | 0 | 0 | 0 |
| 0 | 0 | 0 | 0 | 0 | 0 | 0 | 0 | 0 |
| 0 | 0 | 0 | 0 | 0 | 0 | 0 | 0 | 0 |
| 0 | 0 | 0 | 0 | 0 | 0 | 0 | 0 | 0 |
| 0 | 0 | 0 | 39 | 0 | 0 | 0 | 0 | 0 |
| 0 | 0 | 0 | 0 | 0 | 0 | 0 | 0 | 0 |
| 0 | 0 | 0 | 0 | 0 | 0 | 0 | 0 | 0 |
| 0 | 0 | 0 | 58 | 0 | 0 | 0 | 0 | 0 |
| 53 | 11 | 38 | 0 | 0 | 0 | 0 | 0 | 0 |
| 0 | 0 | 0 | 0 | 0 | 0 | 0 | 0 | 0 |
| 0 | 0 | 0 | 0 | 0 | 0 | 0 | 0 | 0 |
| 0 | 0 | 0 | 36 | 0 | 0 | 0 | 0 | 0 |
| 0 | 0 | 0 | 0 | 0 | 0 | 0 | 0 | 0 |

| | | | | | | | | |
|---|---|---|---|---|---|---|---|---|
| 0 | 0 | 0 | 0 | 0 | 0 | 0 | 0 | 0 |
| 0 | 0 | 0 | 24 | 0 | 0 | 0 | 0 | 0 |
| 0 | 0 | 0 | 0 | 0 | 0 | 0 | 16 | 0 |
| 0 | 0 | 0 | 49 | 10 | 16 | 9 | 6 | 0 |
| 0 | 0 | 0 | 44 | 0 | 0 | 0 | 0 | 0 |
| 0 | 0 | 0 | 32 | 0 | 0 | 0 | 0 | 0 |
| 0 | 0 | 0 | 0 | 0 | 0 | 0 | 0 | 0 |
| 0 | 0 | 0 | 24 | 0 | 0 | 0 | 0 | 0 |
| 0 | 0 | 0 | 24 | 0 | 0 | 0 | 0 | 0 |
| 0 | 0 | 0 | 12 | 0 | 0 | 0 | 0 | 0 |
| 0 | 0 | 0 | 0 | 0 | 0 | 0 | 0 | 0 |
| 0 | 0 | 0 | 6 | 0 | 0 | 0 | 0 | 0 |
| 0 | 0 | 0 | 0 | 0 | 0 | 0 | 0 | 0 |
| 0 | 0 | 0 | 19 | 0 | 0 | 0 | 0 | 0 |
| 0 | 0 | 0 | 0 | 0 | 0 | 0 | 0 | 0 |
| 0 | 0 | 0 | 26 | 0 | 0 | 0 | 0 | 0 |
| 0 | 0 | 0 | 49 | 0 | 0 | 0 | 0 | 0 |
| 0 | 0 | 0 | 0 | 0 | 0 | 0 | 0 | 0 |
| 0 | 0 | 0 | 0 | 0 | 0 | 0 | 0 | 0 |
| 0 | 0 | 0 | 0 | 0 | 0 | 0 | 0 | 0 |
| 0 | 0 | 0 | 20 | 0 | 0 | 0 | 0 | 0 |
| 0 | 0 | 0 | 48 | 0 | 0 | 0 | 0 | 0 |
| 0 | 0 | 0 | 12 | 0 | 0 | 0 | 0 | 0 |
| 0 | 0 | 0 | 23 | 47 | 6 | 0 | 0 | 0 |
| 0 | 0 | 0 | 0 | 0 | 0 | 0 | 0 | 0 |
| 0 | 0 | 0 | 47 | 0 | 0 | 0 | 0 | 0 |
| 0 | 0 | 0 | 86 | 0 | 0 | 0 | 0 | 0 |
| 0 | 0 | 0 | 0 | 0 | 0 | 0 | 0 | 0 |
| 0 | 0 | 0 | 42 | 0 | 6 | 0 | 0 | 0 |
| 0 | 0 | 0 | 35 | 0 | 0 | 0 | 0 | 0 |
| 0 | 0 | 0 | 0 | 0 | 0 | 0 | 0 | 0 |
| 0 | 0 | 0 | 33 | 0 | 0 | 0 | 0 | 0 |
| 0 | 0 | 0 | 46 | 0 | 0 | 0 | 0 | 0 |
| 0 | 0 | 0 | 24 | 0 | 0 | 0 | 0 | 0 |
| 0 | 0 | 0 | 0 | 0 | 0 | 0 | 0 | 0 |
| 0 | 0 | 0 | 0 | 0 | 0 | 0 | 0 | 0 |
| 0 | 0 | 0 | 19 | 0 | 0 | 0 | 0 | 0 |
| 0 | 0 | 0 | 0 | 0 | 0 | 0 | 0 | 0 |
| 0 | 0 | 0 | 13 | 0 | 0 | 0 | 0 | 0 |
| 0 | 0 | 0 | 0 | 0 | 24 | 5 | 21 | 0 |
| 0 | 0 | 0 | 33 | 0 | 0 | 0 | 0 | 0 |
| 0 | 0 | 0 | 4 | 0 | 0 | 0 | 0 | 0 |
| 0 | 0 | 0 | 0 | 0 | 0 | 0 | 0 | 0 |
| 0 | 0 | 0 | 5 | 0 | 0 | 0 | 0 | 0 |
| 0 | 0 | 0 | 31 | 0 | 0 | 0 | 0 | 0 |
| 0 | 0 | 0 | 0 | 0 | 0 | 0 | 0 | 0 |
| 0 | 0 | 0 | 6 | 0 | 0 | 0 | 0 | 0 |
| 0 | 0 | 0 | 139 | 0 | 0 | 0 | 0 | 9 |
| 0 | 0 | 0 | 24 | 0 | 0 | 0 | 0 | 0 |
| 0 | 0 | 0 | 0 | 0 | 0 | 0 | 0 | 0 |

| | | | | | | | | |
|---|---|---|---|---|---|---|---|---|
| 0 | 0 | 0 | 38 | 0 | 0 | 0 | 0 | 0 |
| 0 | 0 | 0 | 5 | 0 | 0 | 0 | 0 | 0 |
| 0 | 0 | 0 | 7 | 0 | 0 | 0 | 0 | 0 |
| 0 | 0 | 0 | 0 | 0 | 0 | 13 | 0 | 0 |
| 0 | 0 | 0 | 44 | 0 | 0 | 0 | 0 | 0 |
| 0 | 0 | 0 | 0 | 0 | 0 | 0 | 0 | 8 |
| 0 | 0 | 0 | 0 | 0 | 0 | 0 | 0 | 0 |
| 0 | 0 | 0 | 51 | 0 | 5 | 0 | 0 | 22 |
| 0 | 0 | 0 | 0 | 0 | 0 | 0 | 0 | 0 |
| 0 | 0 | 0 | 0 | 0 | 0 | 0 | 0 | 0 |
| 0 | 0 | 0 | 61 | 0 | 11 | 0 | 0 | 0 |
| 0 | 0 | 0 | 0 | 0 | 0 | 0 | 0 | 0 |
| 0 | 0 | 0 | 7 | 0 | 0 | 0 | 0 | 0 |
| 0 | 0 | 0 | 27 | 0 | 0 | 0 | 0 | 0 |
| 0 | 0 | 0 | 58 | 0 | 0 | 0 | 0 | 0 |
| 0 | 0 | 0 | 20 | 0 | 0 | 0 | 0 | 0 |
| 0 | 0 | 0 | 0 | 0 | 0 | 0 | 0 | 0 |
| 0 | 0 | 0 | 23 | 0 | 0 | 0 | 0 | 18 |
| 0 | 0 | 0 | 0 | 0 | 0 | 0 | 0 | 0 |
| 0 | 0 | 0 | 0 | 0 | 0 | 0 | 0 | 0 |
| 0 | 0 | 0 | 0 | 0 | 0 | 0 | 0 | 15 |
| 0 | 0 | 0 | 200 | 0 | 0 | 0 | 0 | 0 |
| 0 | 0 | 0 | 17 | 0 | 0 | 0 | 0 | 0 |
| 0 | 0 | 0 | 33 | 0 | 0 | 0 | 0 | 0 |
| 0 | 0 | 0 | 0 | 0 | 0 | 0 | 0 | 0 |
| 0 | 0 | 0 | 0 | 0 | 10 | 10 | 51 | 0 |
| 0 | 0 | 0 | 56 | 0 | 0 | 0 | 0 | 23 |
| 0 | 0 | 0 | 0 | 0 | 0 | 0 | 0 | 0 |
| 0 | 0 | 0 | 9 | 0 | 0 | 0 | 0 | 0 |
| 0 | 0 | 0 | 0 | 0 | 0 | 0 | 0 | 18 |
| 0 | 0 | 0 | 0 | 0 | 0 | 39 | 15 | 0 |
| 0 | 0 | 0 | 24 | 0 | 0 | 0 | 0 | 0 |
| 0 | 0 | 0 | 0 | 0 | 0 | 0 | 0 | 0 |
| 0 | 0 | 0 | 0 | 0 | 0 | 0 | 38 | 0 |
| 0 | 0 | 0 | 78 | 0 | 13 | 0 | 0 | 0 |
| 0 | 0 | 0 | 38 | 0 | 0 | 0 | 0 | 0 |
| 0 | 0 | 0 | 116 | 0 | 0 | 0 | 0 | 5 |
| 0 | 0 | 0 | 18 | 0 | 0 | 0 | 0 | 0 |
| 0 | 0 | 0 | 0 | 0 | 0 | 0 | 0 | 0 |
| 0 | 0 | 0 | 0 | 0 | 0 | 0 | 0 | 0 |
| 0 | 0 | 0 | 21 | 0 | 0 | 0 | 0 | 0 |
| 0 | 13 | 0 | 0 | 0 | 0 | 0 | 0 | 0 |
| 0 | 0 | 0 | 0 | 0 | 0 | 0 | 0 | 0 |
| 0 | 0 | 0 | 0 | 0 | 0 | 0 | 0 | 0 |
| 0 | 0 | 0 | 10 | 7 | 17 | 0 | 0 | 14 |
| 0 | 0 | 0 | 56 | 0 | 0 | 0 | 0 | 0 |
| 0 | 0 | 0 | 37 | 0 | 0 | 0 | 0 | 0 |
| 0 | 0 | 0 | 37 | 14 | 0 | 8 | 14 | 0 |
| 0 | 0 | 0 | 61 | 0 | 0 | 0 | 0 | 0 |
| 0 | 0 | 0 | 0 | 0 | 0 | 0 | 0 | 0 |

| | | | | | | | | |
|---|---|---|---|---|---|---|---|---|
| 0 | 0 | 0 | 0 | 0 | 0 | 0 | 0 | 0 |
| 0 | 0 | 0 | 89 | 0 | 0 | 0 | 0 | 0 |
| 0 | 0 | 0 | 0 | 0 | 0 | 0 | 0 | 36 |
| 4 | 0 | 0 | 0 | 0 | 0 | 0 | 0 | 0 |
| 0 | 0 | 0 | 0 | 0 | 0 | 0 | 0 | 0 |
| 0 | 0 | 0 | 0 | 0 | 0 | 0 | 0 | 0 |
| 0 | 0 | 0 | 0 | 0 | 0 | 0 | 0 | 0 |
| 0 | 0 | 0 | 18 | 0 | 0 | 0 | 0 | 0 |
| 22 | 0 | 0 | 0 | 17 | 5 | 17 | 18 | 0 |
| 16 | 12 | 0 | 4 | 12 | 2 | 21 | 33 | 13 |
| 0 | 0 | 0 | 35 | 0 | 0 | 0 | 0 | 0 |
| 0 | 0 | 0 | 0 | 0 | 0 | 0 | 0 | 0 |
| 0 | 0 | 0 | 60 | 0 | 0 | 0 | 0 | 9 |
| 0 | 0 | 0 | 49 | 0 | 0 | 0 | 0 | 0 |
| 0 | 0 | 0 | 45 | 0 | 0 | 0 | 0 | 0 |
| 0 | 0 | 0 | 45 | 0 | 0 | 0 | 0 | 47 |
| 0 | 0 | 0 | 0 | 0 | 0 | 0 | 0 | 34 |
| 0 | 0 | 0 | 34 | 7 | 0 | 0 | 6 | 34 |
| 0 | 0 | 0 | 12 | 0 | 0 | 0 | 0 | 27 |
| 0 | 0 | 0 | 0 | 0 | 0 | 0 | 0 | 0 |
| 0 | 0 | 0 | 0 | 0 | 0 | 0 | 0 | 0 |
| 0 | 0 | 0 | 81 | 0 | 0 | 0 | 0 | 0 |
| 0 | 0 | 0 | 0 | 0 | 0 | 0 | 0 | 0 |
| 0 | 0 | 0 | 11 | 0 | 0 | 0 | 0 | 0 |
| 0 | 0 | 0 | 0 | 0 | 0 | 0 | 0 | 0 |
| 0 | 0 | 0 | 52 | 0 | 0 | 0 | 0 | 0 |
| 0 | 0 | 0 | 14 | 0 | 0 | 0 | 0 | 0 |
| 0 | 0 | 0 | 3 | 0 | 0 | 0 | 0 | 0 |
| 0 | 0 | 0 | 0 | 0 | 0 | 0 | 0 | 0 |
| 0 | 0 | 0 | 10 | 0 | 0 | 0 | 0 | 0 |
| 0 | 0 | 0 | 0 | 0 | 0 | 0 | 0 | 0 |
| 0 | 0 | 0 | 5 | 0 | 0 | 0 | 0 | 0 |
| 0 | 0 | 0 | 29 | 0 | 0 | 0 | 0 | 0 |
| 0 | 0 | 0 | 27 | 0 | 0 | 0 | 0 | 0 |
| 0 | 0 | 0 | 0 | 0 | 0 | 0 | 0 | 0 |
| 0 | 0 | 0 | 0 | 0 | 0 | 0 | 0 | 0 |
| 0 | 0 | 0 | 31 | 0 | 0 | 0 | 0 | 0 |
| 0 | 0 | 0 | 0 | 0 | 0 | 0 | 0 | 0 |
| 0 | 0 | 0 | 67 | 0 | 0 | 0 | 0 | 0 |
| 7 | 0 | 30 | 0 | 0 | 0 | 0 | 0 | 0 |
| 0 | 0 | 0 | 47 | 0 | 0 | 0 | 0 | 0 |
| 0 | 0 | 0 | 0 | 0 | 0 | 0 | 0 | 0 |
| 0 | 0 | 0 | 4 | 17 | 0 | 0 | 0 | 30 |
| 0 | 0 | 0 | 16 | 0 | 0 | 0 | 0 | 0 |
| 0 | 0 | 0 | 6 | 0 | 0 | 0 | 0 | 0 |
| 0 | 0 | 0 | 0 | 0 | 0 | 0 | 0 | 0 |
| 0 | 0 | 0 | 0 | 0 | 17 | 0 | 12 | 0 |
| 0 | 0 | 0 | 0 | 0 | 0 | 0 | 0 | 29 |
| 0 | 0 | 0 | 0 | 0 | 0 | 0 | 0 | 0 |
| 0 | 0 | 0 | 14 | 0 | 0 | 0 | 0 | 12 |

| | | | | | | | | |
|---|---|---|---|---|---|---|---|---|
| 5 | 0 | 0 | 0 | 0 | 0 | 0 | 0 | 0 |
| 0 | 0 | 0 | 0 | 0 | 0 | 0 | 0 | 0 |
| 0 | 0 | 0 | 0 | 0 | 6 | 0 | 0 | 0 |
| 0 | 0 | 0 | 8 | 0 | 0 | 0 | 0 | 0 |
| 0 | 0 | 0 | 0 | 0 | 0 | 0 | 0 | 0 |
| 0 | 0 | 0 | 0 | 0 | 0 | 0 | 0 | 9 |
| 0 | 0 | 0 | 35 | 0 | 0 | 0 | 0 | 0 |
| 0 | 0 | 0 | 0 | 0 | 0 | 38 | 20 | 0 |
| 0 | 0 | 0 | 0 | 0 | 0 | 0 | 0 | 0 |
| 0 | 0 | 0 | 0 | 0 | 0 | 10 | 12 | 0 |
| 0 | 0 | 0 | 27 | 0 | 0 | 0 | 0 | 0 |
| 0 | 0 | 0 | 27 | 0 | 0 | 0 | 0 | 0 |
| 0 | 0 | 0 | 19 | 0 | 0 | 0 | 0 | 0 |
| 0 | 0 | 0 | 0 | 0 | 0 | 0 | 0 | 0 |
| 0 | 0 | 0 | 0 | 0 | 0 | 0 | 0 | 0 |
| 0 | 0 | 0 | 0 | 0 | 0 | 0 | 0 | 0 |
| 0 | 0 | 0 | 9 | 0 | 0 | 0 | 0 | 0 |
| 0 | 0 | 0 | 26 | 0 | 0 | 0 | 0 | 0 |
| 0 | 0 | 0 | 28 | 0 | 0 | 0 | 0 | 0 |
| 0 | 0 | 0 | 26 | 0 | 0 | 0 | 0 | 0 |
| 0 | 0 | 0 | 0 | 0 | 0 | 0 | 0 | 0 |
| 0 | 0 | 0 | 6 | 0 | 0 | 0 | 0 | 0 |
| 0 | 0 | 0 | 0 | 0 | 0 | 0 | 0 | 26 |
| 0 | 0 | 0 | 0 | 0 | 22 | 0 | 0 | 0 |
| 0 | 0 | 0 | 12 | 0 | 0 | 0 | 0 | 0 |
| 0 | 0 | 0 | 7 | 0 | 0 | 0 | 0 | 0 |
| 0 | 0 | 0 | 0 | 0 | 0 | 0 | 0 | 0 |
| 0 | 0 | 0 | 21 | 0 | 0 | 0 | 0 | 0 |
| 0 | 0 | 0 | 5 | 0 | 0 | 0 | 0 | 0 |
| 0 | 0 | 0 | 0 | 0 | 0 | 0 | 0 | 0 |
| 0 | 0 | 0 | 35 | 0 | 0 | 0 | 0 | 0 |
| 0 | 0 | 0 | 25 | 0 | 0 | 0 | 0 | 0 |
| 0 | 0 | 0 | 25 | 0 | 0 | 0 | 0 | 0 |
| 0 | 0 | 0 | 57 | 0 | 0 | 0 | 0 | 0 |
| 0 | 0 | 0 | 25 | 0 | 0 | 0 | 0 | 0 |
| 0 | 0 | 0 | 0 | 0 | 0 | 0 | 0 | 25 |
| 0 | 0 | 0 | 0 | 0 | 0 | 0 | 0 | 0 |
| 0 | 0 | 0 | 0 | 0 | 0 | 0 | 0 | 0 |
| 0 | 0 | 0 | 7 | 0 | 0 | 0 | 0 | 0 |
| 0 | 0 | 0 | 31 | 0 | 0 | 0 | 0 | 0 |
| 0 | 0 | 0 | 24 | 0 | 0 | 0 | 0 | 0 |
| 0 | 0 | 0 | 0 | 0 | 0 | 0 | 0 | 24 |
| 0 | 0 | 0 | 0 | 0 | 0 | 0 | 0 | 0 |
| 0 | 0 | 0 | 23 | 0 | 0 | 0 | 0 | 0 |
| 0 | 0 | 0 | 0 | 0 | 0 | 0 | 0 | 0 |
| 0 | 0 | 0 | 0 | 0 | 0 | 0 | 0 | 0 |
| 0 | 0 | 0 | 0 | 0 | 0 | 0 | 0 | 0 |
| 0 | 0 | 0 | 0 | 0 | 0 | 0 | 0 | 0 |
| 0 | 0 | 0 | 10 | 0 | 0 | 0 | 0 | 0 |
| 0 | 0 | 0 | 0 | 0 | 0 | 0 | 0 | 0 |

| | | | | | | | | |
|---|---|---|---|---|---|---|---|---|
| 0 | 0 | 0 | 14 | 0 | 0 | 0 | 0 | 0 |
| 0 | 0 | 0 | 0 | 13 | 10 | 0 | 0 | 0 |
| 0 | 0 | 0 | 23 | 0 | 0 | 0 | 0 | 0 |
| 0 | 0 | 0 | 0 | 0 | 0 | 0 | 0 | 0 |
| 0 | 0 | 0 | 0 | 0 | 0 | 0 | 0 | 0 |
| 0 | 0 | 0 | 0 | 0 | 0 | 0 | 0 | 0 |
| 0 | 0 | 0 | 4 | 0 | 0 | 0 | 0 | 0 |
| 0 | 0 | 0 | 0 | 0 | 0 | 0 | 0 | 0 |
| 0 | 0 | 0 | 0 | 0 | 0 | 0 | 0 | 16 |
| 0 | 0 | 0 | 8 | 0 | 0 | 0 | 0 | 9 |
| 0 | 0 | 0 | 0 | 0 | 0 | 0 | 0 | 0 |
| 0 | 0 | 0 | 18 | 0 | 0 | 0 | 0 | 0 |
| 0 | 0 | 0 | 22 | 0 | 0 | 0 | 0 | 0 |
| 0 | 0 | 0 | 22 | 0 | 0 | 0 | 0 | 0 |
| 0 | 0 | 0 | 22 | 0 | 0 | 0 | 0 | 0 |
| 0 | 0 | 0 | 2 | 0 | 0 | 0 | 0 | 22 |
| 0 | 0 | 0 | 0 | 0 | 0 | 0 | 0 | 0 |
| 0 | 0 | 0 | 6 | 0 | 0 | 0 | 0 | 0 |
| 0 | 0 | 0 | 0 | 0 | 0 | 0 | 0 | 12 |
| 0 | 0 | 0 | 33 | 0 | 0 | 0 | 0 | 0 |
| 0 | 0 | 0 | 9 | 0 | 0 | 0 | 0 | 0 |
| 0 | 0 | 0 | 6 | 0 | 0 | 0 | 0 | 0 |
| 0 | 0 | 0 | 6 | 0 | 0 | 0 | 0 | 0 |
| 0 | 0 | 0 | 0 | 0 | 0 | 0 | 0 | 0 |
| 0 | 0 | 0 | 0 | 0 | 0 | 0 | 0 | 0 |
| 0 | 0 | 0 | 3 | 0 | 0 | 0 | 0 | 0 |
| 0 | 0 | 0 | 23 | 0 | 0 | 0 | 0 | 0 |
| 0 | 0 | 0 | 11 | 0 | 0 | 0 | 0 | 0 |
| 0 | 0 | 0 | 0 | 0 | 0 | 0 | 0 | 0 |
| 0 | 0 | 0 | 0 | 0 | 0 | 0 | 0 | 0 |
| 0 | 0 | 0 | 8 | 0 | 0 | 0 | 0 | 0 |
| 0 | 0 | 0 | 0 | 0 | 0 | 0 | 0 | 0 |
| 0 | 0 | 0 | 20 | 0 | 0 | 0 | 0 | 0 |
| 0 | 0 | 0 | 26 | 0 | 0 | 0 | 0 | 3 |
| 0 | 0 | 0 | 0 | 0 | 0 | 0 | 0 | 0 |
| 0 | 0 | 0 | 0 | 0 | 0 | 0 | 0 | 0 |
| 0 | 0 | 0 | 0 | 0 | 0 | 0 | 0 | 0 |
| 0 | 0 | 0 | 0 | 0 | 0 | 0 | 0 | 0 |
| 0 | 0 | 0 | 17 | 0 | 0 | 0 | 0 | 15 |
| 0 | 0 | 0 | 0 | 0 | 0 | 0 | 0 | 0 |
| 0 | 0 | 0 | 0 | 0 | 0 | 0 | 0 | 0 |
| 0 | 0 | 0 | 8 | 0 | 0 | 0 | 0 | 5 |
| 0 | 0 | 0 | 15 | 0 | 0 | 0 | 0 | 0 |
| 0 | 0 | 0 | 0 | 0 | 0 | 0 | 0 | 0 |
| 0 | 0 | 0 | 0 | 0 | 0 | 0 | 0 | 0 |
| 0 | 0 | 0 | 9 | 0 | 0 | 0 | 0 | 14 |
| 0 | 0 | 0 | 19 | 0 | 0 | 0 | 0 | 0 |
| 0 | 0 | 0 | 24 | 0 | 0 | 0 | 0 | 0 |
| 0 | 0 | 0 | 19 | 0 | 0 | 0 | 0 | 0 |
| 0 | 0 | 0 | 0 | 0 | 0 | 0 | 0 | 0 |

| | | | | | | | | |
|---|---|---|---|---|---|---|---|---|
| 0 | 0 | 0 | 0 | 0 | 0 | 0 | 0 | 0 |
| 0 | 0 | 0 | 16 | 0 | 0 | 0 | 0 | 0 |
| 0 | 0 | 0 | 0 | 0 | 0 | 0 | 0 | 0 |
| 0 | 0 | 0 | 0 | 0 | 0 | 0 | 0 | 0 |
| 0 | 0 | 0 | 0 | 0 | 0 | 0 | 0 | 0 |
| 0 | 0 | 0 | 21 | 0 | 0 | 0 | 0 | 0 |
| 0 | 0 | 0 | 18 | 0 | 0 | 0 | 0 | 0 |
| 0 | 0 | 0 | 0 | 0 | 0 | 0 | 0 | 0 |
| 0 | 0 | 0 | 0 | 0 | 0 | 0 | 0 | 0 |
| 0 | 0 | 0 | 0 | 0 | 0 | 0 | 0 | 0 |
| 0 | 0 | 0 | 0 | 0 | 0 | 0 | 0 | 0 |
| 0 | 0 | 0 | 34 | 0 | 0 | 0 | 0 | 0 |
| 0 | 0 | 0 | 0 | 0 | 0 | 0 | 0 | 0 |
| 0 | 0 | 0 | 9 | 0 | 0 | 0 | 0 | 0 |
| 0 | 0 | 0 | 0 | 0 | 0 | 0 | 0 | 7 |
| 0 | 0 | 0 | 23 | 0 | 0 | 0 | 0 | 0 |
| 0 | 0 | 0 | 17 | 0 | 0 | 0 | 0 | 0 |
| 0 | 0 | 0 | 17 | 0 | 0 | 0 | 0 | 0 |
| 0 | 0 | 0 | 17 | 0 | 0 | 0 | 0 | 0 |
| 0 | 0 | 0 | 0 | 0 | 0 | 0 | 0 | 0 |
| 0 | 0 | 0 | 4 | 0 | 0 | 0 | 0 | 0 |
| 0 | 0 | 0 | 0 | 0 | 0 | 0 | 0 | 0 |
| 0 | 0 | 0 | 0 | 0 | 17 | 0 | 0 | 0 |
| 0 | 0 | 0 | 9 | 0 | 0 | 0 | 0 | 0 |
| 0 | 0 | 0 | 0 | 0 | 0 | 0 | 0 | 0 |
| 0 | 0 | 0 | 0 | 0 | 0 | 0 | 0 | 0 |
| 13 | 7 | 16 | 0 | 0 | 0 | 0 | 0 | 0 |
| 0 | 0 | 0 | 0 | 0 | 0 | 0 | 0 | 0 |
| 0 | 0 | 0 | 0 | 0 | 0 | 0 | 0 | 23 |
| 0 | 0 | 0 | 8 | 0 | 0 | 0 | 0 | 0 |
| 0 | 0 | 0 | 0 | 0 | 0 | 0 | 0 | 0 |
| 0 | 0 | 0 | 0 | 0 | 0 | 0 | 0 | 0 |
| 0 | 0 | 0 | 0 | 0 | 0 | 0 | 0 | 0 |
| 0 | 0 | 15 | 0 | 0 | 0 | 0 | 0 | 0 |
| 0 | 0 | 0 | 22 | 0 | 0 | 0 | 0 | 0 |
| 0 | 0 | 0 | 15 | 0 | 0 | 0 | 0 | 0 |
| 0 | 0 | 0 | 15 | 0 | 0 | 0 | 0 | 0 |
| 0 | 0 | 0 | 15 | 0 | 0 | 0 | 0 | 0 |
| 0 | 0 | 0 | 15 | 0 | 0 | 0 | 0 | 0 |
| 0 | 0 | 0 | 15 | 0 | 0 | 0 | 0 | 0 |
| 0 | 0 | 0 | 15 | 0 | 0 | 0 | 0 | 0 |
| 0 | 0 | 0 | 12 | 0 | 0 | 0 | 0 | 0 |
| 0 | 0 | 0 | 0 | 0 | 0 | 0 | 0 | 0 |
| 0 | 0 | 0 | 0 | 0 | 0 | 0 | 0 | 0 |
| 0 | 10 | 0 | 0 | 3 | 0 | 0 | 0 | 0 |
| 0 | 0 | 0 | 12 | 0 | 0 | 0 | 0 | 0 |
| 0 | 0 | 0 | 4 | 3 | 0 | 0 | 0 | 0 |
| 0 | 0 | 0 | 0 | 0 | 0 | 0 | 0 | 0 |
| 0 | 0 | 0 | 0 | 14 | 2 | 0 | 0 | 0 |
| 0 | 0 | 0 | 20 | 0 | 0 | 0 | 0 | 0 |

| | | | | | | | | |
|---|---|---|---|---|---|---|---|---|
| 0 | 0 | 0 | 14 | 0 | 0 | 0 | 0 | 0 |
| 0 | 0 | 0 | 0 | 0 | 0 | 0 | 0 | 0 |
| 0 | 0 | 0 | 7 | 0 | 0 | 0 | 0 | 0 |
| 0 | 0 | 0 | 0 | 0 | 0 | 0 | 0 | 0 |
| 0 | 0 | 0 | 6 | 0 | 0 | 0 | 0 | 0 |
| 0 | 0 | 0 | 0 | 0 | 0 | 0 | 0 | 0 |
| 0 | 0 | 0 | 0 | 0 | 0 | 0 | 0 | 0 |
| 0 | 0 | 0 | 13 | 0 | 0 | 0 | 0 | 0 |
| 0 | 0 | 0 | 13 | 0 | 0 | 0 | 0 | 0 |
| 0 | 0 | 0 | 13 | 0 | 0 | 0 | 0 | 0 |
| 0 | 0 | 0 | 22 | 0 | 0 | 0 | 0 | 0 |
| 0 | 0 | 0 | 0 | 0 | 0 | 0 | 0 | 0 |
| 0 | 0 | 0 | 0 | 0 | 0 | 0 | 0 | 0 |
| 0 | 0 | 0 | 4 | 0 | 0 | 0 | 0 | 0 |
| 0 | 0 | 0 | 16 | 0 | 0 | 0 | 0 | 0 |
| 0 | 0 | 0 | 0 | 12 | 0 | 0 | 0 | 0 |
| 0 | 0 | 0 | 12 | 0 | 0 | 0 | 0 | 0 |
| 0 | 0 | 0 | 12 | 0 | 0 | 0 | 0 | 0 |
| 0 | 0 | 0 | 12 | 0 | 0 | 0 | 0 | 0 |
| 0 | 0 | 0 | 12 | 0 | 0 | 0 | 0 | 0 |
| 0 | 0 | 0 | 12 | 0 | 0 | 0 | 0 | 0 |
| 0 | 0 | 0 | 12 | 0 | 0 | 0 | 0 | 0 |
| 0 | 0 | 0 | 0 | 0 | 0 | 0 | 0 | 0 |
| 0 | 0 | 0 | 0 | 0 | 0 | 0 | 0 | 0 |
| 0 | 0 | 0 | 0 | 0 | 0 | 0 | 0 | 12 |
| 0 | 0 | 0 | 0 | 0 | 0 | 0 | 0 | 0 |
| 0 | 0 | 0 | 0 | 0 | 0 | 0 | 0 | 0 |
| 0 | 0 | 0 | 4 | 0 | 0 | 0 | 0 | 0 |
| 0 | 0 | 0 | 0 | 0 | 0 | 0 | 0 | 0 |
| 0 | 0 | 0 | 0 | 0 | 0 | 0 | 0 | 0 |
| 0 | 0 | 0 | 0 | 0 | 0 | 0 | 0 | 0 |
| 0 | 0 | 0 | 0 | 0 | 0 | 0 | 0 | 0 |
| 0 | 0 | 0 | 0 | 0 | 0 | 0 | 0 | 0 |
| 0 | 0 | 0 | 11 | 0 | 0 | 0 | 0 | 0 |
| 0 | 0 | 0 | 11 | 0 | 0 | 0 | 0 | 0 |
| 0 | 0 | 0 | 11 | 0 | 0 | 0 | 0 | 0 |
| 0 | 0 | 0 | 11 | 0 | 0 | 0 | 0 | 0 |
| 0 | 0 | 0 | 0 | 0 | 0 | 0 | 0 | 11 |
| 0 | 0 | 0 | 0 | 0 | 0 | 0 | 0 | 11 |
| 0 | 0 | 0 | 0 | 0 | 0 | 0 | 0 | 0 |
| 0 | 0 | 0 | 0 | 0 | 0 | 0 | 0 | 0 |
| 0 | 0 | 0 | 0 | 0 | 0 | 0 | 0 | 0 |
| 0 | 0 | 0 | 0 | 0 | 0 | 0 | 0 | 0 |
| 0 | 0 | 0 | 0 | 0 | 0 | 0 | 0 | 0 |
| 0 | 0 | 0 | 0 | 0 | 0 | 0 | 0 | 0 |
| 0 | 0 | 0 | 0 | 0 | 0 | 0 | 0 | 0 |
| 0 | 0 | 0 | 0 | 0 | 0 | 0 | 0 | 0 |
| 0 | 0 | 0 | 0 | 0 | 0 | 0 | 0 | 0 |
| 0 | 0 | 0 | 10 | 0 | 0 | 0 | 0 | 0 |
| 0 | 0 | 0 | 19 | 0 | 0 | 0 | 0 | 0 |

| | | | | | | | | |
|---|---|---|---|---|---|---|---|---|
| 0 | 0 | 0 | 10 | 0 | 0 | 0 | 0 | 0 |
| 0 | 0 | 0 | 10 | 0 | 0 | 0 | 0 | 0 |
| 0 | 0 | 0 | 10 | 0 | 0 | 0 | 0 | 0 |
| 0 | 0 | 0 | 10 | 0 | 0 | 0 | 0 | 0 |
| 0 | 0 | 0 | 10 | 0 | 0 | 0 | 0 | 0 |
| 0 | 0 | 0 | 0 | 0 | 0 | 0 | 0 | 10 |
| 0 | 0 | 0 | 0 | 0 | 0 | 0 | 0 | 10 |
| 0 | 0 | 0 | 8 | 0 | 0 | 0 | 0 | 0 |
| 0 | 0 | 0 | 0 | 0 | 0 | 0 | 0 | 0 |
| 0 | 0 | 0 | 0 | 0 | 0 | 0 | 0 | 0 |
| 0 | 0 | 0 | 0 | 0 | 8 | 0 | 9 | 0 |
| 0 | 0 | 0 | 9 | 0 | 0 | 0 | 0 | 0 |
| 0 | 0 | 0 | 9 | 0 | 0 | 0 | 0 | 0 |
| 0 | 0 | 0 | 9 | 0 | 0 | 0 | 0 | 0 |
| 0 | 0 | 0 | 9 | 0 | 0 | 0 | 0 | 0 |
| 0 | 0 | 0 | 0 | 0 | 0 | 0 | 0 | 9 |
| 0 | 0 | 0 | 0 | 0 | 0 | 0 | 0 | 0 |
| 0 | 0 | 0 | 0 | 0 | 0 | 0 | 0 | 0 |
| 0 | 0 | 0 | 0 | 0 | 0 | 0 | 0 | 0 |
| 0 | 0 | 0 | 0 | 0 | 0 | 0 | 0 | 0 |
| 0 | 0 | 0 | 0 | 0 | 0 | 0 | 0 | 0 |
| 0 | 8 | 0 | 0 | 0 | 0 | 0 | 0 | 0 |
| 0 | 0 | 0 | 0 | 0 | 0 | 0 | 0 | 0 |
| 0 | 0 | 0 | 0 | 0 | 0 | 0 | 0 | 0 |
| 0 | 0 | 0 | 0 | 0 | 8 | 0 | 0 | 0 |
| 0 | 0 | 0 | 8 | 0 | 0 | 0 | 0 | 0 |
| 0 | 0 | 0 | 8 | 0 | 0 | 0 | 0 | 0 |
| 0 | 0 | 0 | 8 | 0 | 0 | 0 | 0 | 0 |
| 0 | 0 | 0 | 8 | 0 | 0 | 0 | 0 | 0 |
| 0 | 0 | 0 | 14 | 0 | 0 | 0 | 0 | 8 |
| 0 | 0 | 0 | 0 | 0 | 0 | 0 | 0 | 0 |
| 0 | 0 | 0 | 0 | 0 | 0 | 0 | 0 | 0 |
| 0 | 0 | 0 | 0 | 0 | 0 | 0 | 0 | 0 |
| 0 | 0 | 0 | 0 | 0 | 0 | 0 | 0 | 0 |
| 0 | 0 | 0 | 0 | 0 | 7 | 0 | 0 | 0 |
| 0 | 0 | 0 | 7 | 0 | 0 | 0 | 0 | 0 |
| 0 | 0 | 0 | 7 | 0 | 0 | 0 | 0 | 0 |
| 0 | 0 | 0 | 7 | 0 | 0 | 0 | 0 | 0 |
| 0 | 0 | 0 | 0 | 0 | 0 | 0 | 0 | 0 |
| 0 | 0 | 0 | 0 | 0 | 0 | 0 | 0 | 0 |
| 0 | 0 | 0 | 0 | 0 | 0 | 0 | 0 | 0 |
| 0 | 0 | 0 | 0 | 0 | 0 | 0 | 0 | 0 |
| 0 | 0 | 0 | 0 | 0 | 0 | 0 | 0 | 0 |
| 0 | 0 | 0 | 0 | 0 | 0 | 0 | 0 | 0 |
| 0 | 0 | 0 | 0 | 0 | 0 | 0 | 0 | 0 |
| 0 | 0 | 0 | 6 | 0 | 0 | 0 | 0 | 0 |
| 0 | 0 | 0 | 6 | 0 | 0 | 0 | 0 | 0 |
| 0 | 0 | 0 | 6 | 0 | 0 | 0 | 0 | 0 |
| 0 | 0 | 0 | 6 | 0 | 0 | 0 | 0 | 0 |
| 0 | 0 | 0 | 6 | 0 | 0 | 0 | 0 | 0 |

| | | | | | | | | |
|---|---|---|---|---|---|---|---|---|
| 0 | 0 | 0 | 6 | 0 | 0 | 0 | 0 | 0 |
| 0 | 0 | 0 | 6 | 0 | 0 | 0 | 0 | 0 |
| 0 | 0 | 0 | 6 | 0 | 0 | 0 | 0 | 0 |
| 0 | 0 | 0 | 0 | 0 | 0 | 0 | 0 | 0 |
| 0 | 0 | 0 | 0 | 0 | 0 | 0 | 0 | 0 |
| 0 | 0 | 0 | 0 | 0 | 0 | 0 | 0 | 0 |
| 0 | 0 | 0 | 0 | 0 | 0 | 0 | 0 | 0 |
| 0 | 0 | 0 | 0 | 0 | 0 | 0 | 0 | 0 |
| 0 | 0 | 0 | 0 | 0 | 0 | 0 | 0 | 0 |
| 0 | 0 | 0 | 0 | 0 | 0 | 0 | 0 | 0 |
| 0 | 0 | 0 | 0 | 0 | 0 | 0 | 0 | 0 |
| 0 | 0 | 0 | 0 | 0 | 0 | 0 | 0 | 0 |
| 0 | 0 | 0 | 0 | 0 | 0 | 0 | 0 | 0 |
| 2 | 0 | 0 | 0 | 0 | 0 | 0 | 0 | 0 |
| 0 | 0 | 0 | 0 | 0 | 0 | 0 | 5 | 0 |
| 0 | 0 | 0 | 5 | 0 | 0 | 0 | 0 | 0 |
| 0 | 0 | 0 | 5 | 0 | 0 | 0 | 0 | 0 |
| 0 | 0 | 0 | 5 | 0 | 0 | 0 | 0 | 0 |
| 0 | 0 | 0 | 5 | 0 | 0 | 0 | 0 | 0 |
| 0 | 0 | 4 | 0 | 0 | 0 | 0 | 0 | 0 |
| 0 | 0 | 0 | 0 | 0 | 0 | 0 | 0 | 0 |
| 0 | 0 | 0 | 0 | 4 | 0 | 0 | 0 | 0 |
| 0 | 0 | 0 | 0 | 0 | 0 | 0 | 4 | 0 |
| 0 | 0 | 0 | 4 | 0 | 0 | 0 | 0 | 0 |
| 0 | 0 | 0 | 4 | 0 | 0 | 0 | 0 | 0 |
| 0 | 0 | 0 | 4 | 0 | 0 | 0 | 0 | 0 |
| 0 | 0 | 0 | 0 | 0 | 0 | 0 | 0 | 0 |
| 0 | 0 | 0 | 0 | 0 | 0 | 0 | 0 | 0 |
| 0 | 0 | 0 | 0 | 0 | 0 | 0 | 0 | 0 |
| 0 | 0 | 0 | 3 | 0 | 0 | 0 | 0 | 0 |
| 0 | 0 | 0 | 3 | 0 | 0 | 0 | 0 | 0 |
| 0 | 0 | 0 | 3 | 0 | 0 | 0 | 0 | 0 |
| 0 | 0 | 0 | 0 | 0 | 0 | 0 | 0 | 0 |
| 0 | 0 | 0 | 0 | 0 | 0 | 0 | 0 | 0 |
| 0 | 0 | 0 | 0 | 0 | 0 | 0 | 0 | 0 |
| 0 | 2 | 0 | 0 | 0 | 0 | 0 | 0 | 0 |
| 0 | 0 | 0 | 0 | 0 | 2 | 0 | 0 | 0 |
| 0 | 0 | 0 | 2 | 0 | 0 | 0 | 0 | 0 |
| 0 | 0 | 0 | 0 | 0 | 0 | 0 | 0 | 0 |
| 0 | 0 | 0 | 0 | 0 | 0 | 0 | 0 | 0 |
| 0 | 0 | 0 | 0 | 0 | 0 | 0 | 0 | 0 |
| 0 | 0 | 0 | 0 | 0 | 0 | 0 | 0 | 0 |

| 10-µM-$Cu^{2+}$ | | | | 10%-$CH_4$ Cu-free | | | | 10%-$CH_4$ |
|---|---|---|---|---|---|---|---|---|
| R13 | R19 | R30 | R43 | CB | R06 | R08 | R12 | CB |
| 7044 | 18493 | 21570 | 17351 | 407 | 46 | 146 | 159 | 2599 |
| 257 | 0 | 0 | 0 | 918 | 43876 | 40976 | 4183 | 32 |
| 460 | 3165 | 2064 | 875 | 484 | 0 | 10 | 10 | 183 |
| 317 | 0 | 0 | 0 | 271 | 0 | 321 | 21148 | 0 |
| 0 | 0 | 0 | 0 | 65 | 0 | 0 | 0 | 347 |
| 1517 | 1031 | 452 | 165 | 0 | 628 | 587 | 283 | 0 |
| 0 | 0 | 0 | 0 | 77 | 0 | 0 | 0 | 412 |
| 74 | 0 | 16 | 46 | 283 | 0 | 0 | 1 | 1023 |
| 142 | 69 | 8 | 0 | 0 | 0 | 0 | 0 | 0 |
| 0 | 39 | 157 | 15 | 5 | 1540 | 2386 | 589 | 62 |
| 53 | 90 | 1396 | 587 | 0 | 58 | 242 | 323 | 115 |
| 0 | 0 | 0 | 0 | 0 | 0 | 0 | 0 | 6 |
| 665 | 444 | 257 | 330 | 68 | 296 | 851 | 656 | 0 |
| 790 | 138 | 191 | 111 | 212 | 534 | 1278 | 2333 | 2763 |
| 0 | 983 | 2231 | 1270 | 0 | 0 | 53 | 28 | 20 |
| 702 | 0 | 1072 | 3799 | 211 | 5 | 7 | 0 | 177 |
| 27 | 0 | 195 | 556 | 184 | 82 | 75 | 0 | 109 |
| 386 | 252 | 535 | 261 | 45 | 0 | 0 | 62 | 11 |
| 0 | 0 | 0 | 0 | 106 | 0 | 0 | 0 | 140 |
| 1301 | 7 | 35 | 0 | 283 | 84 | 330 | 1177 | 831 |
| 498 | 558 | 379 | 47 | 257 | 123 | 797 | 1740 | 1096 |
| 0 | 0 | 0 | 0 | 628 | 0 | 0 | 0 | 1983 |
| 0 | 0 | 0 | 30 | 0 | 0 | 0 | 0 | 101 |
| 77 | 0 | 1138 | 1578 | 101 | 0 | 25 | 148 | 48 |
| 0 | 0 | 0 | 0 | 0 | 11 | 30 | 234 | 0 |
| 3055 | 58 | 261 | 0 | 0 | 124 | 482 | 464 | 240 |
| 0 | 0 | 0 | 0 | 280 | 0 | 0 | 0 | 647 |
| 2758 | 15 | 352 | 262 | 0 | 11 | 24 | 393 | 180 |
| 57 | 781 | 217 | 87 | 55 | 93 | 353 | 757 | 0 |
| 63 | 0 | 0 | 0 | 107 | 121 | 659 | 1856 | 566 |
| 0 | 0 | 14 | 0 | 862 | 0 | 0 | 33 | 1581 |
| 0 | 0 | 0 | 0 | 304 | 0 | 0 | 0 | 1505 |
| 319 | 877 | 800 | 270 | 0 | 50 | 287 | 221 | 0 |
| 39 | 203 | 825 | 761 | 0 | 0 | 0 | 0 | 0 |
| 0 | 0 | 0 | 0 | 0 | 0 | 0 | 0 | 13 |
| 0 | 0 | 0 | 6 | 339 | 0 | 0 | 0 | 1395 |
| 312 | 0 | 0 | 0 | 438 | 0 | 0 | 0 | 81 |
| 1606 | 0 | 17 | 105 | 0 | 0 | 0 | 0 | 0 |
| 0 | 0 | 0 | 0 | 1061 | 0 | 0 | 0 | 190 |
| 1795 | 11 | 8 | 54 | 0 | 0 | 0 | 84 | 167 |
| 0 | 0 | 0 | 0 | 63 | 0 | 0 | 0 | 29 |
| 0 | 0 | 0 | 0 | 1677 | 0 | 0 | 0 | 475 |
| 37 | 0 | 0 | 0 | 439 | 37 | 111 | 77 | 156 |
| 0 | 0 | 0 | 0 | 0 | 0 | 0 | 0 | 142 |
| 15 | 0 | 0 | 0 | 120 | 0 | 0 | 0 | 5289 |
| 59 | 0 | 45 | 203 | 178 | 0 | 0 | 0 | 62 |

| | | | | | | | | |
|---|---|---|---|---|---|---|---|---|
| 0 | 85 | 546 | 87 | 36 | 9 | 20 | 7 | 9 |
| 0 | 0 | 0 | 88 | 0 | 0 | 0 | 0 | 0 |
| 0 | 0 | 0 | 0 | 0 | 0 | 0 | 0 | 1395 |
| 0 | 0 | 0 | 0 | 0 | 0 | 0 | 0 | 0 |
| 0 | 0 | 0 | 0 | 0 | 0 | 0 | 0 | 156 |
| 1319 | 329 | 82 | 19 | 0 | 64 | 16 | 0 | 17 |
| 0 | 0 | 0 | 0 | 127 | 0 | 0 | 0 | 192 |
| 0 | 0 | 0 | 0 | 0 | 0 | 0 | 0 | 26 |
| 0 | 176 | 35 | 43 | 0 | 397 | 183 | 78 | 0 |
| 23 | 0 | 0 | 0 | 0 | 4 | 0 | 0 | 0 |
| 0 | 0 | 0 | 0 | 169 | 0 | 0 | 0 | 4 |
| 16 | 29 | 41 | 24 | 0 | 23 | 0 | 138 | 0 |
| 0 | 285 | 166 | 43 | 7 | 0 | 9 | 165 | 164 |
| 144 | 0 | 0 | 0 | 0 | 0 | 0 | 0 | 104 |
| 32 | 0 | 0 | 0 | 74 | 0 | 0 | 0 | 6 |
| 30 | 103 | 50 | 6 | 112 | 65 | 266 | 603 | 65 |
| 0 | 0 | 0 | 0 | 0 | 0 | 0 | 0 | 237 |
| 0 | 0 | 0 | 0 | 0 | 0 | 0 | 0 | 9 |
| 0 | 0 | 0 | 0 | 0 | 0 | 0 | 0 | 0 |
| 113 | 0 | 83 | 103 | 0 | 0 | 0 | 0 | 85 |
| 9 | 0 | 0 | 0 | 0 | 0 | 0 | 0 | 75 |
| 990 | 0 | 0 | 0 | 0 | 0 | 0 | 0 | 43 |
| 0 | 0 | 0 | 0 | 834 | 0 | 0 | 0 | 536 |
| 341 | 0 | 0 | 0 | 51 | 0 | 0 | 0 | 120 |
| 0 | 0 | 0 | 0 | 85 | 0 | 0 | 0 | 3201 |
| 509 | 301 | 269 | 235 | 119 | 0 | 0 | 0 | 0 |
| 0 | 0 | 0 | 0 | 0 | 0 | 0 | 0 | 0 |
| 0 | 0 | 0 | 0 | 61 | 0 | 0 | 0 | 0 |
| 245 | 221 | 42 | 0 | 0 | 350 | 135 | 258 | 0 |
| 70 | 40 | 0 | 0 | 173 | 0 | 0 | 0 | 37 |
| 0 | 0 | 0 | 0 | 0 | 0 | 64 | 33 | 0 |
| 0 | 4 | 63 | 0 | 0 | 0 | 0 | 0 | 0 |
| 0 | 0 | 0 | 0 | 83 | 0 | 0 | 0 | 168 |
| 754 | 0 | 0 | 0 | 0 | 0 | 0 | 0 | 0 |
| 243 | 0 | 0 | 0 | 0 | 0 | 0 | 0 | 97 |
| 0 | 0 | 0 | 0 | 0 | 0 | 0 | 0 | 4 |
| 117 | 225 | 106 | 48 | 0 | 0 | 0 | 0 | 0 |
| 118 | 0 | 0 | 0 | 748 | 0 | 0 | 0 | 445 |
| 0 | 0 | 0 | 0 | 597 | 0 | 0 | 0 | 12 |
| 121 | 180 | 205 | 106 | 0 | 0 | 0 | 0 | 140 |
| 14 | 41 | 115 | 45 | 0 | 0 | 0 | 0 | 229 |
| 11 | 0 | 0 | 0 | 0 | 0 | 0 | 0 | 0 |
| 555 | 14 | 6 | 9 | 118 | 9 | 78 | 169 | 619 |
| 0 | 0 | 0 | 0 | 0 | 0 | 0 | 0 | 32 |
| 32 | 0 | 0 | 308 | 0 | 0 | 0 | 0 | 0 |
| 0 | 0 | 0 | 0 | 23 | 0 | 0 | 0 | 0 |
| 0 | 0 | 0 | 0 | 776 | 0 | 0 | 0 | 405 |
| 15 | 0 | 0 | 0 | 26 | 0 | 0 | 0 | 37 |
| 0 | 0 | 0 | 0 | 164 | 0 | 0 | 0 | 0 |
| 0 | 0 | 0 | 0 | 144 | 23 | 0 | 0 | 0 |

| | | | | | | | | |
|---|---|---|---|---|---|---|---|---|
| 0 | 0 | 0 | 0 | 80 | 0 | 0 | 0 | 0 |
| 0 | 0 | 0 | 0 | 36 | 0 | 0 | 0 | 64 |
| 0 | 0 | 0 | 0 | 0 | 0 | 0 | 0 | 0 |
| 0 | 0 | 0 | 0 | 240 | 0 | 0 | 0 | 98 |
| 0 | 0 | 0 | 0 | 227 | 0 | 0 | 0 | 150 |
| 0 | 0 | 0 | 0 | 0 | 0 | 0 | 0 | 0 |
| 172 | 0 | 0 | 0 | 0 | 0 | 0 | 0 | 49 |
| 0 | 0 | 0 | 0 | 86 | 0 | 0 | 0 | 0 |
| 0 | 0 | 0 | 0 | 0 | 0 | 0 | 0 | 0 |
| 7 | 0 | 26 | 34 | 0 | 0 | 0 | 9 | 0 |
| 0 | 0 | 0 | 0 | 131 | 0 | 0 | 0 | 0 |
| 0 | 0 | 0 | 0 | 50 | 0 | 0 | 0 | 0 |
| 0 | 92 | 53 | 7 | 274 | 0 | 0 | 0 | 113 |
| 0 | 0 | 0 | 0 | 60 | 0 | 0 | 0 | 0 |
| 0 | 0 | 0 | 0 | 176 | 0 | 0 | 0 | 138 |
| 33 | 117 | 181 | 131 | 0 | 0 | 0 | 0 | 0 |
| 0 | 0 | 0 | 0 | 271 | 0 | 0 | 0 | 120 |
| 0 | 0 | 0 | 0 | 0 | 0 | 0 | 0 | 0 |
| 0 | 0 | 0 | 0 | 0 | 0 | 0 | 0 | 29 |
| 0 | 0 | 0 | 0 | 43 | 0 | 0 | 0 | 19 |
| 9 | 0 | 0 | 0 | 0 | 0 | 0 | 4 | 0 |
| 0 | 0 | 0 | 0 | 0 | 0 | 0 | 0 | 10 |
| 0 | 12 | 50 | 22 | 30 | 0 | 0 | 0 | 0 |
| 50 | 327 | 117 | 0 | 32 | 0 | 0 | 0 | 0 |
| 0 | 0 | 0 | 0 | 17 | 38 | 0 | 0 | 0 |
| 0 | 0 | 0 | 0 | 0 | 0 | 0 | 0 | 0 |
| 0 | 0 | 0 | 0 | 0 | 0 | 0 | 0 | 0 |
| 0 | 0 | 0 | 0 | 122 | 0 | 0 | 0 | 78 |
| 0 | 0 | 0 | 0 | 0 | 0 | 0 | 0 | 49 |
| 0 | 0 | 0 | 0 | 49 | 0 | 0 | 0 | 342 |
| 0 | 0 | 0 | 0 | 150 | 0 | 0 | 0 | 22 |
| 0 | 0 | 0 | 0 | 132 | 0 | 0 | 0 | 114 |
| 0 | 98 | 30 | 0 | 50 | 118 | 62 | 39 | 1 |
| 9 | 41 | 122 | 64 | 0 | 0 | 5 | 16 | 0 |
| 476 | 0 | 0 | 0 | 0 | 0 | 0 | 0 | 0 |
| 0 | 62 | 0 | 0 | 374 | 0 | 0 | 0 | 0 |
| 0 | 0 | 0 | 0 | 0 | 0 | 0 | 0 | 45 |
| 0 | 0 | 0 | 0 | 251 | 0 | 0 | 0 | 8 |
| 24 | 0 | 0 | 0 | 73 | 0 | 0 | 0 | 96 |
| 0 | 21 | 0 | 0 | 12 | 61 | 52 | 23 | 16 |
| 0 | 0 | 0 | 0 | 97 | 0 | 0 | 0 | 71 |
| 0 | 0 | 0 | 0 | 0 | 0 | 0 | 0 | 0 |
| 0 | 185 | 14 | 0 | 70 | 0 | 0 | 0 | 160 |
| 0 | 20 | 28 | 50 | 388 | 89 | 95 | 613 | 204 |
| 53 | 0 | 0 | 0 | 67 | 0 | 0 | 0 | 6 |
| 0 | 0 | 0 | 0 | 265 | 0 | 0 | 0 | 82 |
| 32 | 60 | 79 | 114 | 0 | 0 | 0 | 46 | 70 |
| 0 | 0 | 0 | 0 | 0 | 0 | 0 | 0 | 9 |
| 0 | 0 | 0 | 0 | 0 | 0 | 0 | 0 | 0 |
| 8 | 0 | 0 | 0 | 0 | 0 | 0 | 0 | 18 |

| | | | | | | | | |
|---|---|---|---|---|---|---|---|---|
| 0 | 0 | 0 | 0 | 226 | 0 | 0 | 0 | 0 |
| 0 | 0 | 0 | 0 | 128 | 0 | 0 | 0 | 95 |
| 0 | 0 | 0 | 0 | 0 | 0 | 0 | 0 | 0 |
| 0 | 0 | 0 | 0 | 0 | 0 | 0 | 0 | 32 |
| 51 | 7 | 0 | 0 | 0 | 5 | 43 | 215 | 0 |
| 0 | 0 | 0 | 0 | 153 | 0 | 0 | 0 | 47 |
| 0 | 0 | 0 | 0 | 118 | 0 | 0 | 0 | 0 |
| 0 | 0 | 0 | 0 | 138 | 0 | 0 | 16 | 47 |
| 0 | 0 | 0 | 0 | 166 | 0 | 0 | 0 | 70 |
| 0 | 0 | 0 | 0 | 74 | 0 | 0 | 0 | 3 |
| 0 | 0 | 0 | 0 | 48 | 0 | 0 | 0 | 243 |
| 313 | 74 | 14 | 0 | 0 | 0 | 0 | 0 | 0 |
| 379 | 0 | 0 | 0 | 0 | 0 | 0 | 0 | 0 |
| 0 | 0 | 0 | 0 | 139 | 0 | 0 | 0 | 9 |
| 0 | 0 | 0 | 0 | 34 | 0 | 0 | 0 | 24 |
| 0 | 0 | 0 | 0 | 0 | 0 | 0 | 0 | 25 |
| 0 | 0 | 0 | 0 | 0 | 0 | 0 | 0 | 13 |
| 0 | 0 | 0 | 0 | 0 | 0 | 0 | 0 | 0 |
| 10 | 0 | 0 | 0 | 159 | 0 | 0 | 0 | 23 |
| 0 | 0 | 0 | 0 | 0 | 0 | 0 | 0 | 0 |
| 0 | 0 | 0 | 0 | 0 | 0 | 0 | 0 | 36 |
| 0 | 0 | 0 | 0 | 0 | 0 | 0 | 0 | 0 |
| 295 | 0 | 0 | 0 | 0 | 0 | 0 | 1 | 0 |
| 0 | 0 | 0 | 0 | 25 | 0 | 0 | 0 | 0 |
| 0 | 0 | 0 | 0 | 49 | 0 | 0 | 0 | 27 |
| 0 | 0 | 0 | 0 | 0 | 0 | 0 | 0 | 0 |
| 0 | 0 | 0 | 0 | 8 | 0 | 0 | 0 | 25 |
| 0 | 0 | 0 | 0 | 0 | 0 | 62 | 215 | 0 |
| 0 | 0 | 0 | 0 | 175 | 0 | 0 | 0 | 42 |
| 0 | 0 | 0 | 0 | 0 | 0 | 0 | 0 | 0 |
| 0 | 0 | 0 | 0 | 0 | 0 | 0 | 0 | 0 |
| 0 | 0 | 0 | 0 | 0 | 0 | 0 | 0 | 0 |
| 0 | 0 | 0 | 0 | 147 | 0 | 0 | 0 | 0 |
| 0 | 0 | 0 | 0 | 56 | 0 | 0 | 0 | 0 |
| 0 | 0 | 0 | 0 | 0 | 0 | 0 | 0 | 0 |
| 0 | 0 | 0 | 0 | 0 | 0 | 0 | 0 | 0 |
| 500 | 0 | 0 | 0 | 0 | 0 | 0 | 0 | 0 |
| 0 | 0 | 0 | 0 | 0 | 0 | 0 | 0 | 11 |
| 0 | 0 | 0 | 0 | 0 | 0 | 0 | 0 | 0 |
| 0 | 0 | 0 | 0 | 0 | 0 | 0 | 0 | 0 |
| 0 | 0 | 0 | 0 | 0 | 0 | 0 | 0 | 0 |
| 27 | 0 | 0 | 0 | 47 | 0 | 0 | 0 | 103 |
| 165 | 0 | 0 | 0 | 0 | 65 | 59 | 22 | 0 |
| 234 | 0 | 0 | 0 | 0 | 0 | 0 | 0 | 0 |
| 0 | 0 | 0 | 0 | 83 | 0 | 0 | 0 | 11 |
| 0 | 0 | 0 | 0 | 32 | 0 | 0 | 0 | 7 |
| 0 | 0 | 0 | 0 | 0 | 0 | 0 | 0 | 0 |
| 0 | 0 | 0 | 0 | 93 | 0 | 0 | 0 | 0 |
| 0 | 0 | 0 | 0 | 0 | 0 | 0 | 0 | 0 |
| 0 | 0 | 0 | 0 | 10 | 0 | 0 | 0 | 0 |

| | | | | | | | | |
|---|---|---|---|---|---|---|---|---|
| 0 | 0 | 0 | 0 | 0 | 0 | 0 | 0 | 0 |
| 0 | 0 | 0 | 0 | 202 | 0 | 0 | 0 | 0 |
| 133 | 0 | 0 | 11 | 0 | 25 | 11 | 0 | 0 |
| 0 | 0 | 0 | 0 | 0 | 0 | 0 | 0 | 0 |
| 0 | 0 | 0 | 0 | 0 | 0 | 0 | 0 | 12 |
| 0 | 0 | 0 | 0 | 0 | 0 | 0 | 0 | 0 |
| 0 | 0 | 0 | 0 | 0 | 0 | 0 | 0 | 63 |
| 0 | 0 | 0 | 0 | 48 | 0 | 0 | 0 | 19 |
| 0 | 0 | 0 | 0 | 84 | 0 | 0 | 0 | 0 |
| 6 | 0 | 0 | 0 | 55 | 0 | 0 | 0 | 0 |
| 0 | 0 | 0 | 0 | 0 | 0 | 0 | 0 | 0 |
| 6 | 0 | 0 | 0 | 17 | 0 | 0 | 0 | 29 |
| 0 | 0 | 0 | 0 | 0 | 0 | 0 | 0 | 0 |
| 0 | 0 | 0 | 0 | 0 | 0 | 0 | 0 | 65 |
| 0 | 0 | 0 | 0 | 0 | 0 | 0 | 0 | 0 |
| 0 | 0 | 68 | 135 | 0 | 0 | 0 | 0 | 0 |
| 0 | 0 | 0 | 0 | 134 | 0 | 0 | 0 | 69 |
| 0 | 0 | 0 | 0 | 0 | 0 | 0 | 0 | 0 |
| 0 | 0 | 21 | 0 | 198 | 0 | 0 | 0 | 0 |
| 0 | 0 | 0 | 0 | 0 | 0 | 0 | 0 | 0 |
| 0 | 0 | 0 | 0 | 18 | 0 | 0 | 0 | 0 |
| 0 | 0 | 0 | 0 | 0 | 0 | 0 | 0 | 0 |
| 0 | 0 | 0 | 0 | 161 | 0 | 0 | 0 | 321 |
| 0 | 0 | 0 | 0 | 0 | 0 | 0 | 0 | 0 |
| 0 | 0 | 0 | 0 | 0 | 0 | 0 | 0 | 0 |
| 196 | 0 | 0 | 0 | 0 | 0 | 0 | 0 | 17 |
| 185 | 0 | 0 | 0 | 0 | 0 | 0 | 0 | 0 |
| 0 | 0 | 0 | 0 | 0 | 0 | 0 | 0 | 0 |
| 219 | 0 | 0 | 0 | 0 | 0 | 0 | 0 | 0 |
| 0 | 0 | 0 | 0 | 0 | 0 | 0 | 0 | 0 |
| 0 | 0 | 0 | 0 | 0 | 0 | 0 | 0 | 0 |
| 0 | 0 | 0 | 0 | 0 | 0 | 0 | 0 | 20 |
| 0 | 0 | 0 | 0 | 0 | 0 | 0 | 0 | 0 |
| 0 | 0 | 0 | 0 | 141 | 0 | 0 | 0 | 0 |
| 8 | 0 | 30 | 88 | 0 | 0 | 0 | 0 | 0 |
| 0 | 0 | 0 | 0 | 0 | 0 | 0 | 0 | 0 |
| 0 | 0 | 0 | 0 | 0 | 0 | 0 | 0 | 0 |
| 0 | 0 | 0 | 0 | 0 | 0 | 0 | 0 | 0 |
| 37 | 0 | 0 | 0 | 0 | 0 | 0 | 0 | 0 |
| 0 | 0 | 0 | 0 | 0 | 0 | 0 | 0 | 14 |
| 0 | 0 | 0 | 0 | 0 | 0 | 0 | 0 | 0 |
| 0 | 0 | 0 | 0 | 0 | 0 | 0 | 0 | 0 |
| 0 | 112 | 0 | 0 | 0 | 0 | 0 | 0 | 0 |
| 0 | 0 | 0 | 0 | 0 | 14 | 12 | 26 | 0 |
| 0 | 0 | 0 | 0 | 0 | 0 | 0 | 0 | 0 |
| 0 | 0 | 0 | 0 | 52 | 0 | 0 | 0 | 0 |
| 0 | 0 | 0 | 0 | 0 | 0 | 0 | 0 | 0 |
| 0 | 0 | 0 | 0 | 25 | 0 | 0 | 0 | 0 |
| 0 | 0 | 0 | 0 | 0 | 0 | 0 | 0 | 0 |
| 0 | 0 | 0 | 0 | 0 | 0 | 0 | 0 | 12 |

| | | | | | | | | |
|---|---|---|---|---|---|---|---|---|
| 0 | 0 | 0 | 0 | 0 | 115 | 75 | 28 | 0 |
| 0 | 0 | 0 | 0 | 0 | 0 | 0 | 0 | 0 |
| 0 | 0 | 0 | 0 | 50 | 0 | 0 | 0 | 0 |
| 21 | 0 | 0 | 0 | 0 | 0 | 0 | 0 | 8 |
| 0 | 0 | 0 | 0 | 0 | 0 | 0 | 0 | 55 |
| 0 | 0 | 0 | 0 | 0 | 0 | 0 | 0 | 0 |
| 0 | 0 | 0 | 0 | 0 | 0 | 0 | 0 | 0 |
| 0 | 0 | 0 | 0 | 0 | 0 | 0 | 0 | 0 |
| 0 | 0 | 0 | 0 | 140 | 0 | 0 | 0 | 28 |
| 0 | 0 | 0 | 0 | 68 | 0 | 0 | 0 | 0 |
| 0 | 0 | 0 | 0 | 0 | 0 | 0 | 0 | 0 |
| 0 | 0 | 0 | 0 | 0 | 0 | 0 | 7 | 0 |
| 0 | 0 | 0 | 0 | 0 | 0 | 0 | 0 | 0 |
| 0 | 0 | 0 | 0 | 24 | 0 | 0 | 0 | 71 |
| 0 | 0 | 0 | 0 | 0 | 0 | 0 | 0 | 0 |
| 0 | 0 | 0 | 0 | 0 | 0 | 0 | 0 | 11 |
| 0 | 0 | 0 | 0 | 152 | 0 | 0 | 1 | 0 |
| 0 | 0 | 0 | 0 | 0 | 0 | 0 | 0 | 0 |
| 0 | 0 | 0 | 0 | 7 | 0 | 0 | 0 | 0 |
| 0 | 0 | 0 | 0 | 59 | 0 | 0 | 0 | 27 |
| 0 | 0 | 0 | 0 | 5 | 0 | 0 | 0 | 30 |
| 0 | 0 | 0 | 0 | 0 | 0 | 0 | 0 | 0 |
| 0 | 0 | 0 | 0 | 0 | 0 | 0 | 0 | 0 |
| 167 | 0 | 0 | 0 | 0 | 0 | 0 | 0 | 0 |
| 262 | 0 | 0 | 0 | 0 | 0 | 0 | 0 | 0 |
| 0 | 0 | 0 | 0 | 0 | 0 | 0 | 0 | 79 |
| 0 | 0 | 0 | 0 | 93 | 0 | 0 | 0 | 0 |
| 0 | 0 | 0 | 0 | 0 | 0 | 0 | 0 | 0 |
| 0 | 0 | 0 | 0 | 0 | 0 | 0 | 0 | 0 |
| 0 | 0 | 0 | 0 | 0 | 0 | 0 | 0 | 0 |
| 0 | 0 | 0 | 0 | 0 | 0 | 0 | 0 | 5 |
| 0 | 0 | 0 | 0 | 0 | 0 | 0 | 0 | 0 |
| 43 | 0 | 13 | 0 | 0 | 0 | 16 | 112 | 0 |
| 0 | 0 | 0 | 0 | 0 | 0 | 0 | 0 | 0 |
| 0 | 0 | 0 | 0 | 0 | 0 | 0 | 0 | 0 |
| 0 | 0 | 0 | 0 | 0 | 0 | 0 | 0 | 0 |
| 0 | 0 | 0 | 0 | 0 | 0 | 0 | 0 | 0 |
| 0 | 0 | 0 | 0 | 41 | 0 | 0 | 0 | 0 |
| 0 | 0 | 0 | 0 | 0 | 0 | 0 | 0 | 0 |
| 0 | 0 | 0 | 0 | 53 | 0 | 0 | 0 | 0 |
| 0 | 0 | 0 | 0 | 0 | 0 | 0 | 0 | 0 |
| 0 | 0 | 0 | 0 | 0 | 0 | 0 | 6 | 0 |
| 0 | 56 | 35 | 14 | 0 | 0 | 0 | 0 | 47 |
| 62 | 9 | 52 | 21 | 12 | 0 | 0 | 0 | 0 |
| 0 | 0 | 0 | 0 | 0 | 0 | 0 | 0 | 0 |
| 0 | 0 | 0 | 0 | 0 | 16 | 0 | 0 | 0 |
| 0 | 0 | 0 | 0 | 0 | 0 | 0 | 0 | 0 |
| 103 | 0 | 0 | 0 | 0 | 0 | 0 | 0 | 0 |
| 0 | 0 | 0 | 0 | 103 | 0 | 0 | 0 | 0 |
| 0 | 0 | 0 | 0 | 41 | 0 | 0 | 0 | 80 |

| | | | | | | | | |
|---|---|---|---|---|---|---|---|---|
| 0 | 0 | 0 | 0 | 0 | 0 | 0 | 0 | 7 |
| 0 | 0 | 0 | 0 | 0 | 0 | 0 | 0 | 0 |
| 0 | 0 | 0 | 0 | 0 | 0 | 0 | 0 | 11 |
| 0 | 0 | 0 | 0 | 0 | 0 | 0 | 0 | 0 |
| 0 | 0 | 0 | 0 | 98 | 0 | 0 | 0 | 0 |
| 0 | 0 | 0 | 0 | 110 | 0 | 0 | 0 | 0 |
| 0 | 0 | 0 | 0 | 0 | 0 | 0 | 0 | 0 |
| 0 | 0 | 0 | 0 | 0 | 0 | 0 | 0 | 0 |
| 0 | 0 | 0 | 0 | 0 | 0 | 0 | 0 | 0 |
| 0 | 0 | 0 | 0 | 0 | 0 | 0 | 0 | 0 |
| 0 | 0 | 0 | 0 | 26 | 0 | 0 | 0 | 20 |
| 0 | 0 | 0 | 0 | 45 | 0 | 0 | 0 | 0 |
| 0 | 0 | 0 | 0 | 0 | 0 | 0 | 0 | 0 |
| 0 | 0 | 0 | 0 | 75 | 0 | 0 | 0 | 0 |
| 0 | 0 | 0 | 0 | 0 | 0 | 0 | 0 | 0 |
| 0 | 0 | 0 | 0 | 0 | 0 | 0 | 0 | 0 |
| 0 | 0 | 0 | 0 | 0 | 0 | 0 | 0 | 7 |
| 0 | 0 | 0 | 0 | 0 | 0 | 0 | 0 | 3 |
| 2 | 0 | 0 | 0 | 52 | 0 | 0 | 0 | 6 |
| 13 | 0 | 0 | 48 | 0 | 0 | 0 | 0 | 0 |
| 0 | 0 | 0 | 0 | 0 | 0 | 0 | 0 | 0 |
| 0 | 0 | 0 | 0 | 0 | 0 | 0 | 0 | 0 |
| 0 | 0 | 0 | 0 | 0 | 0 | 0 | 0 | 0 |
| 0 | 0 | 0 | 0 | 0 | 0 | 0 | 0 | 0 |
| 0 | 0 | 0 | 0 | 0 | 0 | 0 | 0 | 0 |
| 90 | 0 | 0 | 0 | 0 | 0 | 0 | 0 | 0 |
| 0 | 10 | 71 | 8 | 0 | 0 | 0 | 0 | 0 |
| 0 | 0 | 0 | 0 | 0 | 0 | 0 | 0 | 0 |
| 0 | 0 | 0 | 0 | 36 | 0 | 0 | 0 | 0 |
| 0 | 0 | 0 | 0 | 19 | 0 | 0 | 0 | 0 |
| 0 | 0 | 0 | 0 | 52 | 0 | 0 | 0 | 9 |
| 0 | 0 | 0 | 0 | 65 | 0 | 0 | 171 | 0 |
| 0 | 0 | 0 | 0 | 0 | 0 | 0 | 0 | 0 |
| 0 | 0 | 0 | 0 | 0 | 0 | 0 | 0 | 0 |
| 0 | 0 | 0 | 0 | 0 | 0 | 0 | 0 | 0 |
| 0 | 0 | 0 | 0 | 0 | 0 | 0 | 0 | 0 |
| 0 | 0 | 0 | 0 | 0 | 0 | 0 | 0 | 71 |
| 12 | 0 | 0 | 0 | 0 | 0 | 0 | 0 | 0 |
| 0 | 0 | 0 | 0 | 0 | 0 | 0 | 0 | 0 |
| 0 | 0 | 0 | 0 | 0 | 0 | 0 | 0 | 0 |
| 0 | 0 | 0 | 0 | 0 | 0 | 0 | 0 | 0 |
| 0 | 0 | 0 | 0 | 0 | 0 | 0 | 0 | 0 |
| 0 | 0 | 0 | 0 | 0 | 0 | 0 | 0 | 0 |
| 0 | 0 | 0 | 0 | 0 | 0 | 0 | 0 | 0 |
| 0 | 0 | 0 | 0 | 0 | 0 | 0 | 0 | 0 |
| 0 | 0 | 0 | 0 | 0 | 0 | 0 | 10 | 0 |
| 0 | 0 | 0 | 0 | 0 | 0 | 0 | 0 | 0 |
| 0 | 0 | 0 | 0 | 0 | 0 | 0 | 0 | 0 |
| 0 | 0 | 0 | 0 | 23 | 0 | 0 | 0 | 6 |
| 0 | 0 | 0 | 0 | 79 | 0 | 0 | 0 | 0 |

| | | | | | | | | |
|---|---|---|---|---|---|---|---|---|
| 0 | 0 | 0 | 0 | 0 | 0 | 0 | 0 | 0 |
| 0 | 0 | 0 | 0 | 0 | 0 | 0 | 0 | 20 |
| 0 | 0 | 0 | 0 | 0 | 0 | 8 | 0 | 0 |
| 0 | 0 | 0 | 0 | 0 | 0 | 0 | 0 | 20 |
| 0 | 0 | 0 | 0 | 0 | 0 | 0 | 0 | 0 |
| 0 | 0 | 0 | 0 | 0 | 0 | 0 | 0 | 0 |
| 0 | 0 | 0 | 0 | 0 | 0 | 0 | 0 | 0 |
| 0 | 0 | 0 | 0 | 0 | 0 | 0 | 0 | 0 |
| 0 | 0 | 0 | 0 | 0 | 0 | 0 | 0 | 0 |
| 0 | 0 | 0 | 0 | 0 | 0 | 0 | 0 | 28 |
| 0 | 0 | 0 | 0 | 0 | 0 | 0 | 0 | 10 |
| 75 | 0 | 0 | 0 | 0 | 0 | 0 | 0 | 0 |
| 0 | 0 | 0 | 0 | 74 | 0 | 0 | 0 | 0 |
| 0 | 0 | 0 | 0 | 62 | 0 | 0 | 0 | 0 |
| 0 | 0 | 0 | 0 | 0 | 0 | 0 | 0 | 0 |
| 0 | 0 | 0 | 0 | 0 | 0 | 0 | 0 | 0 |
| 0 | 0 | 0 | 0 | 0 | 0 | 0 | 0 | 0 |
| 0 | 0 | 0 | 0 | 15 | 0 | 0 | 0 | 0 |
| 0 | 0 | 0 | 0 | 0 | 0 | 0 | 0 | 0 |
| 0 | 0 | 0 | 0 | 0 | 0 | 0 | 0 | 0 |
| 0 | 0 | 0 | 0 | 0 | 0 | 0 | 0 | 0 |
| 0 | 0 | 0 | 0 | 0 | 0 | 0 | 0 | 0 |
| 0 | 24 | 16 | 0 | 0 | 0 | 0 | 0 | 0 |
| 0 | 0 | 0 | 0 | 70 | 0 | 0 | 0 | 0 |
| 0 | 0 | 0 | 0 | 0 | 0 | 0 | 0 | 0 |
| 0 | 0 | 0 | 0 | 0 | 0 | 0 | 0 | 0 |
| 0 | 0 | 0 | 0 | 0 | 0 | 0 | 0 | 0 |
| 0 | 0 | 0 | 0 | 0 | 0 | 0 | 0 | 0 |
| 0 | 0 | 0 | 0 | 0 | 0 | 0 | 0 | 0 |
| 0 | 0 | 0 | 0 | 0 | 0 | 0 | 0 | 0 |
| 0 | 0 | 0 | 0 | 0 | 0 | 0 | 0 | 0 |
| 0 | 0 | 0 | 0 | 0 | 0 | 0 | 0 | 0 |
| 0 | 0 | 0 | 0 | 0 | 0 | 0 | 0 | 0 |
| 0 | 0 | 0 | 0 | 0 | 0 | 0 | 0 | 0 |
| 0 | 0 | 0 | 0 | 0 | 0 | 0 | 0 | 0 |
| 0 | 0 | 67 | 0 | 0 | 0 | 0 | 0 | 0 |
| 0 | 0 | 0 | 0 | 0 | 0 | 0 | 0 | 0 |
| 0 | 0 | 0 | 0 | 0 | 0 | 0 | 0 | 0 |
| 0 | 0 | 0 | 0 | 41 | 0 | 0 | 0 | 0 |
| 0 | 0 | 0 | 0 | 0 | 0 | 0 | 0 | 0 |
| 0 | 0 | 0 | 0 | 0 | 0 | 0 | 0 | 0 |
| 0 | 0 | 0 | 0 | 0 | 0 | 0 | 0 | 0 |
| 0 | 0 | 0 | 0 | 0 | 0 | 0 | 0 | 0 |
| 0 | 0 | 0 | 0 | 0 | 0 | 0 | 0 | 0 |
| 0 | 0 | 0 | 0 | 0 | 0 | 0 | 0 | 0 |
| 0 | 0 | 0 | 0 | 0 | 0 | 0 | 0 | 0 |
| 6 | 0 | 0 | 0 | 0 | 0 | 0 | 0 | 0 |
| 0 | 0 | 0 | 0 | 0 | 0 | 0 | 0 | 0 |
| 0 | 0 | 10 | 17 | 0 | 0 | 0 | 0 | 0 |
| 0 | 0 | 0 | 0 | 0 | 0 | 0 | 0 | 0 |

| | | | | | | | | |
|---|---|---|---|---|---|---|---|---|
| 64 | 0 | 0 | 0 | 0 | 0 | 0 | 0 | 0 |
| 44 | 30 | 0 | 0 | 0 | 0 | 0 | 34 | 0 |
| 0 | 0 | 0 | 0 | 64 | 0 | 0 | 35 | 0 |
| 0 | 0 | 0 | 0 | 0 | 0 | 0 | 0 | 0 |
| 0 | 0 | 0 | 0 | 0 | 0 | 0 | 0 | 0 |
| 0 | 0 | 0 | 0 | 0 | 0 | 0 | 0 | 0 |
| 0 | 0 | 0 | 0 | 0 | 0 | 0 | 0 | 0 |
| 0 | 0 | 0 | 0 | 0 | 0 | 0 | 0 | 0 |
| 0 | 0 | 0 | 0 | 22 | 0 | 0 | 0 | 0 |
| 0 | 0 | 0 | 0 | 0 | 0 | 0 | 0 | 0 |
| 0 | 0 | 0 | 0 | 0 | 0 | 0 | 0 | 0 |
| 0 | 0 | 0 | 0 | 0 | 0 | 0 | 0 | 0 |
| 0 | 0 | 0 | 0 | 0 | 0 | 0 | 0 | 0 |
| 0 | 0 | 0 | 0 | 0 | 0 | 0 | 0 | 0 |
| 0 | 0 | 0 | 0 | 0 | 0 | 0 | 0 | 0 |
| 62 | 0 | 0 | 0 | 44 | 0 | 0 | 0 | 71 |
| 0 | 0 | 0 | 0 | 62 | 0 | 0 | 0 | 0 |
| 0 | 0 | 0 | 0 | 0 | 0 | 0 | 0 | 56 |
| 0 | 0 | 0 | 0 | 0 | 0 | 0 | 0 | 0 |
| 0 | 0 | 0 | 0 | 0 | 0 | 0 | 0 | 0 |
| 0 | 0 | 0 | 0 | 0 | 0 | 0 | 0 | 0 |
| 59 | 0 | 0 | 0 | 0 | 0 | 0 | 0 | 0 |
| 0 | 0 | 0 | 0 | 0 | 0 | 0 | 0 | 0 |
| 0 | 0 | 0 | 0 | 0 | 0 | 0 | 0 | 0 |
| 0 | 0 | 0 | 0 | 0 | 0 | 0 | 0 | 0 |
| 0 | 0 | 26 | 32 | 0 | 0 | 0 | 0 | 0 |
| 0 | 0 | 0 | 0 | 0 | 0 | 0 | 0 | 0 |
| 0 | 0 | 0 | 0 | 0 | 0 | 0 | 0 | 0 |
| 0 | 0 | 0 | 0 | 0 | 0 | 0 | 0 | 0 |
| 0 | 0 | 0 | 0 | 0 | 0 | 0 | 15 | 0 |
| 0 | 0 | 0 | 0 | 0 | 0 | 0 | 0 | 0 |
| 0 | 0 | 0 | 0 | 0 | 0 | 0 | 0 | 0 |
| 0 | 0 | 0 | 0 | 0 | 0 | 0 | 0 | 0 |
| 0 | 0 | 0 | 0 | 0 | 0 | 0 | 0 | 0 |
| 0 | 0 | 0 | 0 | 0 | 0 | 0 | 0 | 0 |
| 0 | 0 | 0 | 0 | 0 | 0 | 0 | 0 | 0 |
| 0 | 0 | 0 | 0 | 55 | 0 | 0 | 0 | 0 |
| 0 | 0 | 0 | 0 | 55 | 0 | 0 | 0 | 0 |
| 0 | 0 | 0 | 0 | 55 | 0 | 0 | 0 | 0 |
| 54 | 0 | 0 | 0 | 0 | 0 | 0 | 0 | 0 |
| 0 | 0 | 0 | 0 | 54 | 0 | 0 | 0 | 0 |
| 0 | 0 | 0 | 0 | 54 | 0 | 0 | 0 | 0 |
| 0 | 4 | 0 | 0 | 0 | 0 | 0 | 0 | 54 |
| 0 | 0 | 0 | 0 | 0 | 0 | 0 | 0 | 0 |
| 0 | 0 | 0 | 0 | 0 | 0 | 0 | 0 | 0 |
| 0 | 0 | 0 | 0 | 0 | 0 | 0 | 0 | 0 |
| 0 | 0 | 0 | 0 | 0 | 0 | 0 | 0 | 0 |
| 51 | 0 | 0 | 0 | 0 | 0 | 0 | 0 | 0 |
| 0 | 0 | 0 | 0 | 0 | 0 | 0 | 0 | 0 |
| 0 | 0 | 0 | 0 | 53 | 0 | 0 | 0 | 0 |

| | | | | | | | | |
|---|---|---|---|---|---|---|---|---|
| 0 | 0 | 0 | 0 | 0 | 0 | 0 | 0 | 0 |
| 0 | 0 | 0 | 0 | 0 | 0 | 0 | 0 | 0 |
| 0 | 0 | 0 | 0 | 0 | 0 | 0 | 0 | 0 |
| 0 | 0 | 0 | 0 | 0 | 0 | 0 | 0 | 0 |
| 0 | 0 | 0 | 0 | 0 | 0 | 0 | 0 | 0 |
| 0 | 0 | 0 | 0 | 0 | 0 | 0 | 0 | 0 |
| 0 | 0 | 0 | 0 | 0 | 0 | 0 | 0 | 0 |
| 0 | 0 | 0 | 0 | 0 | 0 | 0 | 0 | 0 |
| 0 | 0 | 0 | 0 | 0 | 0 | 0 | 0 | 0 |
| 0 | 0 | 0 | 0 | 0 | 0 | 0 | 0 | 0 |
| 0 | 0 | 0 | 0 | 0 | 0 | 0 | 0 | 0 |
| 0 | 0 | 0 | 0 | 0 | 0 | 0 | 0 | 0 |
| 0 | 0 | 0 | 0 | 0 | 0 | 0 | 0 | 0 |
| 0 | 0 | 0 | 0 | 0 | 0 | 0 | 0 | 0 |
| 0 | 0 | 0 | 0 | 0 | 0 | 0 | 0 | 0 |
| 0 | 0 | 0 | 0 | 0 | 0 | 0 | 0 | 0 |
| 0 | 0 | 0 | 0 | 0 | 0 | 0 | 0 | 0 |
| 0 | 0 | 0 | 0 | 0 | 0 | 0 | 0 | 0 |
| 0 | 0 | 0 | 0 | 0 | 0 | 0 | 0 | 0 |
| 0 | 0 | 0 | 0 | 0 | 0 | 0 | 0 | 0 |
| 0 | 0 | 0 | 0 | 0 | 0 | 0 | 0 | 0 |
| 0 | 0 | 0 | 0 | 0 | 0 | 0 | 0 | 0 |
| 0 | 0 | 0 | 0 | 0 | 0 | 0 | 0 | 0 |
| 0 | 0 | 0 | 0 | 0 | 0 | 0 | 0 | 0 |
| 0 | 0 | 0 | 0 | 0 | 0 | 0 | 0 | 0 |
| 0 | 0 | 0 | 0 | 0 | 0 | 0 | 0 | 0 |
| 0 | 0 | 0 | 0 | 0 | 0 | 0 | 0 | 0 |
| 0 | 0 | 0 | 0 | 0 | 0 | 0 | 0 | 0 |
| 0 | 0 | 0 | 0 | 0 | 0 | 0 | 0 | 0 |
| 0 | 0 | 0 | 0 | 0 | 0 | 0 | 0 | 0 |
| 0 | 0 | 0 | 0 | 0 | 0 | 0 | 0 | 0 |
| 0 | 0 | 0 | 0 | 0 | 0 | 0 | 0 | 0 |
| 0 | 0 | 0 | 0 | 0 | 0 | 0 | 0 | 0 |
| 0 | 0 | 0 | 0 | 0 | 0 | 0 | 0 | 7 |
| 46 | 0 | 0 | 0 | 0 | 0 | 0 | 0 | 0 |
| 0 | 0 | 0 | 0 | 0 | 0 | 0 | 0 | 0 |
| 0 | 0 | 0 | 0 | 0 | 0 | 0 | 0 | 0 |
| 0 | 0 | 0 | 0 | 0 | 0 | 0 | 0 | 0 |
| 0 | 0 | 0 | 0 | 0 | 0 | 0 | 0 | 0 |
| 0 | 0 | 0 | 0 | 0 | 0 | 0 | 0 | 0 |
| 0 | 0 | 0 | 0 | 0 | 0 | 0 | 0 | 0 |
| 0 | 0 | 0 | 0 | 0 | 0 | 0 | 0 | 0 |
| 0 | 0 | 0 | 0 | 0 | 0 | 0 | 0 | 0 |
| 0 | 0 | 0 | 0 | 0 | 0 | 0 | 0 | 0 |
| 0 | 0 | 0 | 0 | 0 | 0 | 0 | 0 | 0 |
| 0 | 0 | 0 | 0 | 0 | 0 | 0 | 0 | 0 |
| 0 | 0 | 0 | 0 | 0 | 0 | 0 | 0 | 0 |
| 0 | 0 | 0 | 0 | 0 | 0 | 0 | 0 | 4 |
| 0 | 0 | 0 | 0 | 0 | 0 | 0 | 0 | 0 |
| 0 | 0 | 0 | 0 | 0 | 0 | 0 | 0 | 0 |

| | | | | | | | | |
|---|---|---|---|---|---|---|---|---|
| 0 | 0 | 0 | 0 | 0 | 0 | 0 | 0 | 0 |
| 0 | 0 | 0 | 0 | 0 | 0 | 0 | 0 | 0 |
| 0 | 0 | 0 | 0 | 0 | 0 | 0 | 0 | 0 |
| 0 | 0 | 0 | 0 | 0 | 0 | 0 | 0 | 0 |
| 0 | 0 | 0 | 0 | 0 | 0 | 0 | 0 | 0 |
| 0 | 0 | 0 | 0 | 0 | 0 | 0 | 0 | 7 |
| 0 | 0 | 0 | 0 | 42 | 0 | 0 | 0 | 0 |
| 0 | 0 | 0 | 0 | 42 | 0 | 0 | 0 | 7 |
| 0 | 0 | 0 | 0 | 0 | 0 | 0 | 0 | 0 |
| 0 | 0 | 0 | 0 | 0 | 0 | 0 | 0 | 0 |
| 0 | 0 | 0 | 0 | 12 | 0 | 0 | 0 | 0 |
| 0 | 0 | 0 | 0 | 0 | 0 | 0 | 0 | 0 |
| 0 | 0 | 0 | 0 | 0 | 0 | 0 | 0 | 0 |
| 4 | 0 | 0 | 0 | 0 | 0 | 0 | 0 | 0 |
| 0 | 0 | 0 | 0 | 0 | 0 | 0 | 0 | 9 |
| 0 | 0 | 0 | 0 | 0 | 0 | 0 | 0 | 0 |
| 0 | 0 | 0 | 0 | 0 | 0 | 0 | 0 | 0 |
| 0 | 0 | 0 | 0 | 0 | 0 | 0 | 0 | 0 |
| 0 | 0 | 0 | 0 | 0 | 0 | 0 | 0 | 0 |
| 0 | 0 | 0 | 0 | 0 | 0 | 0 | 0 | 0 |
| 0 | 0 | 0 | 0 | 0 | 0 | 0 | 0 | 0 |
| 0 | 0 | 0 | 0 | 0 | 0 | 0 | 0 | 0 |
| 0 | 0 | 0 | 0 | 29 | 0 | 0 | 0 | 0 |
| 0 | 0 | 0 | 0 | 0 | 0 | 0 | 0 | 0 |
| 0 | 0 | 0 | 0 | 0 | 0 | 0 | 0 | 0 |
| 0 | 0 | 0 | 0 | 0 | 0 | 0 | 0 | 0 |
| 0 | 0 | 0 | 0 | 0 | 0 | 0 | 0 | 0 |
| 0 | 0 | 0 | 0 | 0 | 0 | 11 | 29 | 0 |
| 0 | 0 | 0 | 0 | 0 | 0 | 0 | 0 | 0 |
| 0 | 0 | 0 | 0 | 0 | 0 | 0 | 0 | 4 |
| 0 | 0 | 0 | 0 | 0 | 0 | 0 | 0 | 0 |
| 0 | 0 | 0 | 0 | 0 | 0 | 0 | 0 | 15 |
| 0 | 0 | 0 | 0 | 0 | 0 | 0 | 0 | 0 |
| 0 | 0 | 0 | 0 | 0 | 0 | 0 | 0 | 0 |
| 0 | 0 | 0 | 0 | 0 | 0 | 0 | 0 | 0 |
| 0 | 0 | 0 | 0 | 0 | 0 | 0 | 0 | 0 |
| 0 | 0 | 0 | 0 | 0 | 0 | 0 | 0 | 0 |
| 0 | 0 | 0 | 0 | 0 | 0 | 0 | 0 | 0 |
| 0 | 0 | 0 | 0 | 0 | 0 | 0 | 0 | 0 |
| 0 | 0 | 0 | 0 | 0 | 0 | 0 | 0 | 0 |
| 0 | 0 | 0 | 0 | 0 | 0 | 0 | 0 | 0 |
| 0 | 0 | 0 | 0 | 0 | 0 | 0 | 0 | 0 |
| 0 | 0 | 0 | 0 | 0 | 0 | 0 | 0 | 0 |
| 0 | 0 | 0 | 0 | 0 | 0 | 0 | 0 | 0 |
| 0 | 0 | 0 | 0 | 0 | 0 | 0 | 0 | 0 |
| 0 | 0 | 0 | 0 | 0 | 0 | 0 | 0 | 0 |
| 0 | 0 | 0 | 0 | 0 | 0 | 0 | 0 | 0 |
| 0 | 0 | 0 | 0 | 0 | 0 | 0 | 0 | 0 |
| 0 | 0 | 0 | 0 | 0 | 0 | 10 | 12 | 0 |
| 0 | 0 | 0 | 0 | 0 | 0 | 0 | 0 | 0 |

| | | | | | | | | |
|---|---|---|---|---|---|---|---|---|
| 0 | 0 | 0 | 0 | 0 | 0 | 0 | 0 | 0 |
| 0 | 0 | 0 | 0 | 0 | 0 | 0 | 0 | 0 |
| 0 | 0 | 0 | 0 | 0 | 0 | 0 | 0 | 0 |
| 0 | 0 | 0 | 0 | 23 | 0 | 0 | 0 | 0 |
| 0 | 0 | 0 | 0 | 0 | 0 | 0 | 0 | 0 |
| 0 | 0 | 0 | 0 | 0 | 0 | 0 | 0 | 0 |
| 0 | 0 | 0 | 0 | 8 | 0 | 0 | 0 | 0 |
| 0 | 0 | 0 | 0 | 0 | 0 | 0 | 0 | 0 |
| 17 | 0 | 0 | 0 | 0 | 0 | 0 | 0 | 0 |
| 0 | 0 | 0 | 0 | 0 | 0 | 0 | 0 | 11 |
| 0 | 0 | 0 | 0 | 0 | 0 | 0 | 0 | 0 |
| 0 | 0 | 0 | 0 | 0 | 0 | 0 | 0 | 0 |
| 0 | 0 | 0 | 0 | 0 | 0 | 0 | 0 | 0 |
| 0 | 0 | 0 | 0 | 0 | 0 | 0 | 0 | 0 |
| 0 | 0 | 0 | 0 | 0 | 0 | 0 | 0 | 3 |
| 0 | 0 | 0 | 0 | 0 | 0 | 0 | 0 | 0 |
| 0 | 0 | 0 | 0 | 0 | 0 | 0 | 0 | 0 |
| 0 | 0 | 0 | 0 | 0 | 0 | 0 | 0 | 0 |
| 0 | 0 | 0 | 0 | 0 | 0 | 0 | 0 | 0 |
| 0 | 0 | 0 | 0 | 0 | 0 | 0 | 0 | 0 |
| 0 | 0 | 0 | 0 | 0 | 0 | 0 | 0 | 0 |
| 0 | 0 | 0 | 0 | 0 | 0 | 0 | 0 | 0 |
| 0 | 0 | 0 | 0 | 0 | 0 | 0 | 0 | 0 |
| 0 | 0 | 0 | 0 | 0 | 0 | 0 | 0 | 0 |
| 0 | 0 | 0 | 0 | 0 | 0 | 0 | 0 | 0 |
| 0 | 0 | 0 | 0 | 0 | 0 | 0 | 0 | 0 |
| 0 | 0 | 0 | 0 | 0 | 0 | 0 | 0 | 0 |
| 0 | 0 | 0 | 0 | 0 | 0 | 0 | 0 | 0 |
| 0 | 0 | 0 | 0 | 0 | 0 | 0 | 0 | 0 |
| 0 | 0 | 0 | 0 | 0 | 0 | 0 | 0 | 32 |
| 0 | 0 | 0 | 0 | 0 | 0 | 0 | 0 | 0 |
| 0 | 0 | 0 | 0 | 0 | 0 | 0 | 0 | 0 |
| 0 | 0 | 0 | 0 | 0 | 0 | 0 | 0 | 0 |
| 0 | 0 | 0 | 0 | 0 | 0 | 0 | 0 | 0 |
| 0 | 0 | 0 | 0 | 0 | 0 | 0 | 0 | 0 |
| 0 | 0 | 0 | 0 | 0 | 0 | 0 | 0 | 0 |
| 0 | 0 | 0 | 0 | 0 | 0 | 0 | 0 | 0 |
| 0 | 0 | 0 | 0 | 0 | 0 | 0 | 0 | 0 |
| 0 | 0 | 0 | 0 | 0 | 0 | 0 | 0 | 0 |
| 0 | 0 | 0 | 0 | 0 | 0 | 0 | 0 | 0 |
| 0 | 0 | 0 | 0 | 0 | 0 | 0 | 0 | 0 |
| 30 | 0 | 0 | 0 | 0 | 0 | 0 | 0 | 0 |
| 0 | 0 | 0 | 0 | 0 | 0 | 0 | 0 | 0 |
| 0 | 0 | 0 | 0 | 0 | 0 | 0 | 0 | 0 |
| 0 | 0 | 0 | 0 | 0 | 0 | 0 | 0 | 0 |
| 0 | 0 | 0 | 0 | 0 | 0 | 0 | 0 | 0 |
| 0 | 0 | 0 | 0 | 0 | 0 | 0 | 0 | 0 |
| 0 | 21 | 0 | 0 | 0 | 0 | 0 | 0 | 0 |
| 0 | 0 | 0 | 0 | 29 | 0 | 0 | 0 | 0 |
| 0 | 0 | 0 | 0 | 0 | 0 | 0 | 0 | 10 |

| | | | | | | | | |
|---|---|---|---|---|---|---|---|---|
| 0 | 0 | 0 | 0 | 0 | 0 | 0 | 0 | 29 |
| 0 | 0 | 0 | 0 | 0 | 0 | 0 | 0 | 0 |
| 0 | 0 | 0 | 0 | 0 | 0 | 0 | 0 | 0 |
| 0 | 0 | 0 | 0 | 0 | 0 | 0 | 0 | 0 |
| 0 | 0 | 0 | 0 | 0 | 0 | 0 | 0 | 0 |
| 0 | 0 | 0 | 0 | 0 | 0 | 0 | 0 | 0 |
| 0 | 12 | 14 | 3 | 0 | 0 | 0 | 0 | 0 |
| 0 | 0 | 0 | 0 | 0 | 0 | 0 | 0 | 0 |
| 0 | 0 | 0 | 0 | 0 | 0 | 0 | 0 | 0 |
| 0 | 0 | 0 | 0 | 0 | 0 | 0 | 0 | 0 |
| 0 | 0 | 0 | 0 | 0 | 0 | 0 | 0 | 0 |
| 0 | 0 | 0 | 0 | 0 | 0 | 0 | 0 | 0 |
| 0 | 0 | 0 | 0 | 0 | 0 | 0 | 0 | 0 |
| 0 | 0 | 0 | 0 | 0 | 0 | 0 | 0 | 0 |
| 0 | 0 | 0 | 0 | 0 | 0 | 0 | 0 | 0 |
| 0 | 0 | 0 | 0 | 0 | 0 | 0 | 0 | 0 |
| 0 | 0 | 0 | 0 | 0 | 0 | 0 | 0 | 0 |
| 0 | 0 | 0 | 0 | 0 | 0 | 0 | 0 | 0 |
| 0 | 0 | 0 | 0 | 0 | 0 | 0 | 0 | 0 |
| 0 | 0 | 0 | 0 | 0 | 0 | 0 | 0 | 4 |
| 26 | 0 | 0 | 0 | 0 | 0 | 0 | 0 | 0 |
| 0 | 0 | 0 | 34 | 0 | 0 | 0 | 0 | 0 |
| 0 | 0 | 0 | 0 | 0 | 0 | 0 | 0 | 0 |
| 0 | 0 | 0 | 0 | 26 | 0 | 0 | 0 | 0 |
| 0 | 0 | 0 | 0 | 0 | 0 | 0 | 0 | 0 |
| 0 | 0 | 0 | 0 | 0 | 0 | 0 | 0 | 0 |
| 0 | 0 | 0 | 0 | 0 | 0 | 0 | 0 | 0 |
| 0 | 0 | 0 | 0 | 0 | 0 | 0 | 0 | 0 |
| 0 | 0 | 0 | 0 | 0 | 0 | 0 | 0 | 8 |
| 0 | 0 | 0 | 0 | 0 | 0 | 0 | 0 | 0 |
| 0 | 0 | 0 | 0 | 0 | 0 | 0 | 0 | 0 |
| 0 | 0 | 0 | 0 | 0 | 0 | 0 | 0 | 0 |
| 0 | 0 | 0 | 0 | 0 | 0 | 0 | 0 | 0 |
| 0 | 0 | 0 | 0 | 0 | 0 | 0 | 0 | 0 |
| 0 | 0 | 0 | 0 | 0 | 0 | 0 | 0 | 0 |
| 0 | 0 | 0 | 0 | 0 | 0 | 0 | 0 | 0 |
| 0 | 0 | 0 | 0 | 0 | 0 | 0 | 0 | 0 |
| 0 | 0 | 0 | 0 | 0 | 0 | 0 | 0 | 0 |
| 0 | 0 | 0 | 0 | 0 | 0 | 0 | 0 | 0 |
| 0 | 0 | 0 | 0 | 0 | 0 | 0 | 0 | 0 |
| 0 | 0 | 0 | 0 | 0 | 0 | 0 | 0 | 0 |
| 0 | 0 | 0 | 0 | 0 | 0 | 0 | 0 | 0 |
| 0 | 0 | 0 | 0 | 0 | 0 | 0 | 0 | 0 |
| 0 | 0 | 0 | 0 | 0 | 0 | 0 | 0 | 0 |
| 0 | 0 | 0 | 0 | 0 | 0 | 0 | 0 | 0 |
| 0 | 0 | 0 | 0 | 0 | 0 | 0 | 0 | 0 |
| 0 | 0 | 0 | 0 | 0 | 0 | 0 | 0 | 0 |
| 0 | 0 | 0 | 0 | 0 | 0 | 0 | 0 | 0 |
| 0 | 0 | 0 | 0 | 0 | 0 | 0 | 0 | 0 |
| 0 | 0 | 0 | 0 | 0 | 0 | 0 | 0 | 0 |

| | | | | | | | | |
|---|---|---|---|---|---|---|---|---|
| 0 | 0 | 0 | 0 | 0 | 0 | 0 | 0 | 6 |
| 0 | 0 | 0 | 0 | 0 | 0 | 0 | 0 | 0 |
| 0 | 0 | 0 | 0 | 0 | 0 | 0 | 0 | 0 |
| 0 | 0 | 0 | 0 | 23 | 0 | 0 | 0 | 0 |
| 0 | 0 | 0 | 0 | 0 | 0 | 0 | 0 | 0 |
| 0 | 0 | 0 | 0 | 0 | 0 | 0 | 0 | 0 |
| 0 | 0 | 0 | 0 | 0 | 0 | 0 | 0 | 0 |
| 0 | 0 | 0 | 0 | 0 | 0 | 0 | 0 | 9 |
| 0 | 0 | 0 | 0 | 0 | 0 | 0 | 0 | 0 |
| 0 | 0 | 0 | 0 | 0 | 0 | 0 | 0 | 0 |
| 0 | 0 | 0 | 0 | 0 | 0 | 0 | 0 | 0 |
| 0 | 0 | 0 | 0 | 0 | 0 | 0 | 0 | 16 |
| 0 | 0 | 0 | 0 | 0 | 0 | 0 | 0 | 0 |
| 0 | 0 | 0 | 0 | 0 | 0 | 0 | 0 | 0 |
| 0 | 0 | 0 | 0 | 0 | 0 | 0 | 0 | 0 |
| 0 | 0 | 0 | 0 | 0 | 0 | 0 | 0 | 0 |
| 0 | 0 | 0 | 0 | 0 | 0 | 0 | 0 | 0 |
| 0 | 0 | 0 | 0 | 0 | 0 | 0 | 0 | 0 |
| 0 | 0 | 0 | 0 | 0 | 0 | 0 | 0 | 0 |
| 0 | 0 | 0 | 0 | 0 | 0 | 0 | 0 | 0 |
| 0 | 0 | 0 | 0 | 0 | 0 | 0 | 0 | 0 |
| 0 | 0 | 0 | 0 | 0 | 0 | 0 | 0 | 0 |
| 0 | 0 | 0 | 0 | 0 | 0 | 0 | 0 | 0 |
| 21 | 0 | 0 | 0 | 0 | 0 | 0 | 0 | 0 |
| 0 | 0 | 0 | 0 | 0 | 0 | 0 | 0 | 0 |
| 0 | 0 | 0 | 0 | 0 | 0 | 0 | 0 | 0 |
| 0 | 0 | 0 | 0 | 0 | 0 | 0 | 0 | 0 |
| 0 | 0 | 0 | 0 | 0 | 0 | 0 | 0 | 0 |
| 0 | 0 | 0 | 0 | 0 | 0 | 0 | 0 | 7 |
| 0 | 0 | 0 | 0 | 0 | 0 | 0 | 0 | 0 |
| 0 | 0 | 0 | 0 | 0 | 0 | 0 | 0 | 0 |
| 0 | 0 | 0 | 0 | 0 | 0 | 0 | 0 | 0 |
| 0 | 0 | 0 | 0 | 0 | 0 | 0 | 0 | 0 |
| 0 | 0 | 0 | 0 | 0 | 0 | 0 | 0 | 0 |
| 0 | 0 | 0 | 0 | 0 | 0 | 0 | 0 | 0 |
| 0 | 0 | 0 | 0 | 0 | 0 | 0 | 0 | 0 |
| 0 | 0 | 0 | 0 | 0 | 0 | 0 | 0 | 0 |
| 0 | 0 | 0 | 0 | 0 | 0 | 0 | 0 | 0 |
| 0 | 0 | 0 | 0 | 0 | 0 | 0 | 0 | 0 |
| 0 | 0 | 0 | 0 | 0 | 0 | 0 | 0 | 0 |
| 0 | 0 | 0 | 0 | 0 | 0 | 0 | 0 | 0 |
| 0 | 0 | 0 | 0 | 0 | 0 | 0 | 0 | 0 |
| 0 | 0 | 0 | 0 | 0 | 0 | 0 | 0 | 0 |
| 0 | 0 | 0 | 0 | 0 | 0 | 0 | 0 | 0 |
| 0 | 0 | 0 | 0 | 0 | 0 | 0 | 0 | 0 |
| 0 | 0 | 0 | 0 | 0 | 0 | 0 | 0 | 0 |
| 0 | 0 | 0 | 0 | 0 | 0 | 0 | 0 | 0 |
| 0 | 0 | 0 | 0 | 0 | 0 | 0 | 0 | 0 |
| 0 | 0 | 0 | 0 | 0 | 0 | 0 | 0 | 0 |
| 19 | 0 | 0 | 0 | 0 | 0 | 0 | 0 | 0 |

| | | | | | | | | |
|---|---|---|---|---|---|---|---|---|
| 0 | 0 | 0 | 0 | 0 | 0 | 0 | 0 | 0 |
| 0 | 0 | 0 | 0 | 0 | 0 | 0 | 0 | 0 |
| 0 | 0 | 0 | 0 | 0 | 0 | 0 | 0 | 0 |
| 0 | 0 | 0 | 0 | 0 | 0 | 0 | 0 | 0 |
| 0 | 0 | 0 | 0 | 0 | 0 | 0 | 0 | 0 |
| 0 | 0 | 0 | 0 | 0 | 0 | 0 | 0 | 0 |
| 0 | 0 | 0 | 0 | 0 | 0 | 0 | 0 | 0 |
| 0 | 15 | 18 | 0 | 0 | 0 | 0 | 0 | 0 |
| 0 | 0 | 0 | 0 | 0 | 0 | 0 | 0 | 0 |
| 0 | 0 | 0 | 0 | 0 | 0 | 0 | 0 | 3 |
| 0 | 0 | 0 | 0 | 0 | 0 | 0 | 0 | 0 |
| 0 | 0 | 0 | 0 | 0 | 0 | 0 | 0 | 0 |
| 0 | 0 | 0 | 0 | 0 | 0 | 0 | 0 | 0 |
| 0 | 0 | 0 | 0 | 0 | 0 | 0 | 0 | 0 |
| 0 | 0 | 0 | 0 | 0 | 0 | 0 | 0 | 0 |
| 0 | 0 | 0 | 0 | 0 | 0 | 0 | 0 | 0 |
| 0 | 0 | 0 | 0 | 0 | 0 | 0 | 0 | 0 |
| 0 | 0 | 0 | 0 | 0 | 0 | 0 | 0 | 0 |
| 0 | 0 | 0 | 0 | 0 | 0 | 0 | 0 | 0 |
| 0 | 0 | 0 | 0 | 12 | 0 | 0 | 0 | 21 |
| 0 | 0 | 0 | 0 | 0 | 0 | 0 | 0 | 0 |
| 0 | 0 | 0 | 0 | 0 | 0 | 0 | 0 | 0 |
| 0 | 0 | 0 | 0 | 0 | 0 | 0 | 0 | 0 |
| 0 | 0 | 0 | 0 | 0 | 0 | 0 | 0 | 0 |
| 0 | 0 | 0 | 0 | 0 | 0 | 0 | 0 | 0 |
| 0 | 0 | 0 | 0 | 0 | 0 | 0 | 0 | 0 |
| 0 | 0 | 0 | 0 | 0 | 0 | 0 | 0 | 0 |
| 0 | 0 | 0 | 0 | 0 | 0 | 0 | 0 | 0 |
| 0 | 0 | 0 | 0 | 0 | 0 | 0 | 0 | 0 |
| 0 | 0 | 0 | 0 | 0 | 0 | 0 | 0 | 0 |
| 0 | 0 | 0 | 0 | 0 | 0 | 0 | 0 | 0 |
| 0 | 0 | 0 | 0 | 0 | 0 | 0 | 0 | 0 |
| 0 | 0 | 0 | 0 | 0 | 0 | 0 | 0 | 0 |
| 0 | 4 | 0 | 0 | 0 | 0 | 0 | 0 | 0 |
| 0 | 0 | 0 | 0 | 0 | 0 | 0 | 0 | 0 |
| 0 | 0 | 0 | 0 | 0 | 0 | 0 | 0 | 0 |
| 0 | 0 | 0 | 0 | 0 | 0 | 0 | 0 | 0 |
| 0 | 0 | 0 | 0 | 0 | 0 | 0 | 0 | 0 |
| 0 | 0 | 0 | 0 | 0 | 0 | 0 | 0 | 0 |
| 0 | 0 | 0 | 0 | 0 | 0 | 0 | 0 | 0 |
| 0 | 0 | 0 | 0 | 0 | 0 | 0 | 0 | 0 |
| 0 | 0 | 0 | 0 | 0 | 0 | 0 | 0 | 0 |
| 0 | 0 | 0 | 0 | 0 | 0 | 0 | 0 | 0 |
| 0 | 0 | 0 | 0 | 0 | 0 | 0 | 0 | 0 |
| 0 | 0 | 0 | 0 | 0 | 0 | 0 | 0 | 0 |
| 0 | 0 | 0 | 0 | 0 | 0 | 0 | 0 | 0 |
| 0 | 0 | 0 | 0 | 0 | 0 | 0 | 0 | 0 |
| 0 | 0 | 0 | 0 | 0 | 0 | 0 | 0 | 0 |
| 0 | 0 | 0 | 0 | 0 | 0 | 0 | 0 | 0 |
| 0 | 0 | 0 | 0 | 0 | 0 | 0 | 0 | 0 |

| | | | | | | | | |
|---|---|---|---|---|---|---|---|---|
| 0 | 0 | 0 | 0 | 0 | 0 | 0 | 0 | 0 |
| 0 | 0 | 0 | 0 | 0 | 0 | 0 | 0 | 0 |
| 0 | 0 | 0 | 0 | 0 | 0 | 0 | 0 | 0 |
| 0 | 0 | 0 | 0 | 0 | 0 | 0 | 0 | 0 |
| 0 | 0 | 0 | 0 | 0 | 0 | 0 | 0 | 0 |
| 0 | 0 | 0 | 0 | 0 | 0 | 0 | 0 | 0 |
| 0 | 0 | 0 | 0 | 0 | 0 | 0 | 0 | 0 |
| 0 | 0 | 0 | 0 | 0 | 0 | 0 | 0 | 0 |
| 0 | 0 | 0 | 0 | 0 | 0 | 3 | 0 | 0 |
| 0 | 0 | 0 | 0 | 0 | 0 | 0 | 0 | 0 |
| 0 | 0 | 0 | 0 | 0 | 0 | 0 | 0 | 0 |
| 0 | 0 | 0 | 0 | 0 | 0 | 0 | 0 | 0 |
| 0 | 0 | 0 | 0 | 0 | 0 | 0 | 0 | 0 |
| 0 | 0 | 0 | 0 | 0 | 0 | 0 | 0 | 0 |
| 0 | 0 | 0 | 0 | 0 | 0 | 0 | 0 | 0 |
| 0 | 0 | 0 | 0 | 0 | 0 | 0 | 0 | 0 |
| 0 | 0 | 0 | 0 | 0 | 0 | 0 | 0 | 0 |
| 0 | 0 | 0 | 0 | 0 | 0 | 0 | 0 | 0 |
| 0 | 0 | 0 | 0 | 0 | 0 | 0 | 0 | 0 |
| 0 | 0 | 0 | 0 | 0 | 0 | 0 | 0 | 0 |
| 0 | 0 | 0 | 0 | 0 | 0 | 0 | 0 | 0 |
| 0 | 0 | 0 | 0 | 0 | 0 | 0 | 0 | 0 |
| 0 | 12 | 0 | 0 | 0 | 0 | 0 | 0 | 0 |
| 0 | 0 | 12 | 0 | 0 | 0 | 0 | 0 | 0 |
| 0 | 0 | 0 | 0 | 0 | 0 | 0 | 0 | 0 |
| 0 | 0 | 0 | 0 | 0 | 0 | 0 | 0 | 0 |
| 0 | 0 | 0 | 0 | 0 | 0 | 0 | 0 | 0 |
| 0 | 0 | 0 | 0 | 0 | 0 | 0 | 0 | 0 |
| 0 | 0 | 0 | 0 | 0 | 0 | 0 | 0 | 0 |
| 0 | 0 | 0 | 0 | 0 | 0 | 0 | 0 | 0 |
| 0 | 0 | 0 | 0 | 0 | 0 | 0 | 0 | 0 |
| 0 | 0 | 0 | 0 | 0 | 0 | 0 | 0 | 0 |
| 0 | 0 | 0 | 0 | 0 | 0 | 0 | 0 | 0 |
| 0 | 0 | 0 | 0 | 0 | 0 | 0 | 0 | 0 |
| 0 | 0 | 0 | 0 | 0 | 0 | 0 | 0 | 0 |
| 0 | 0 | 0 | 0 | 0 | 0 | 0 | 0 | 0 |
| 0 | 0 | 0 | 0 | 0 | 0 | 0 | 0 | 0 |
| 0 | 0 | 0 | 0 | 0 | 0 | 0 | 0 | 0 |
| 0 | 0 | 0 | 0 | 0 | 0 | 0 | 0 | 0 |
| 0 | 0 | 0 | 0 | 0 | 0 | 0 | 0 | 0 |
| 0 | 0 | 0 | 0 | 0 | 0 | 0 | 0 | 0 |
| 0 | 0 | 0 | 0 | 0 | 0 | 0 | 0 | 0 |
| 0 | 0 | 0 | 0 | 0 | 0 | 0 | 0 | 0 |
| 0 | 0 | 0 | 0 | 0 | 0 | 0 | 0 | 0 |
| 0 | 0 | 0 | 0 | 0 | 0 | 0 | 0 | 0 |
| 0 | 0 | 0 | 0 | 0 | 0 | 0 | 0 | 0 |
| 0 | 0 | 0 | 0 | 0 | 0 | 0 | 0 | 0 |
| 0 | 0 | 0 | 0 | 0 | 0 | 0 | 0 | 0 |
| 0 | 0 | 0 | 0 | 0 | 0 | 0 | 0 | 0 |
| 0 | 0 | 0 | 0 | 0 | 0 | 0 | 0 | 0 |

| | | | | | | | | |
|---|---|---|---|---|---|---|---|---|
| 0 | 0 | 0 | 0 | 0 | 0 | 0 | 0 | 0 |
| 0 | 0 | 0 | 0 | 0 | 0 | 0 | 0 | 0 |
| 0 | 0 | 0 | 0 | 0 | 0 | 0 | 0 | 0 |
| 0 | 0 | 0 | 0 | 0 | 0 | 0 | 0 | 0 |
| 0 | 0 | 0 | 0 | 0 | 0 | 0 | 0 | 0 |
| 0 | 0 | 0 | 0 | 0 | 0 | 0 | 0 | 0 |
| 0 | 0 | 0 | 0 | 0 | 0 | 0 | 0 | 0 |
| 0 | 0 | 0 | 0 | 0 | 0 | 0 | 0 | 0 |
| 0 | 0 | 0 | 0 | 0 | 0 | 0 | 0 | 0 |
| 0 | 0 | 0 | 0 | 0 | 0 | 0 | 0 | 0 |
| 0 | 0 | 0 | 0 | 0 | 0 | 0 | 0 | 0 |
| 0 | 0 | 0 | 0 | 0 | 0 | 0 | 0 | 0 |
| 0 | 0 | 0 | 0 | 0 | 0 | 0 | 0 | 0 |
| 0 | 0 | 0 | 0 | 0 | 0 | 0 | 0 | 0 |
| 0 | 0 | 0 | 0 | 0 | 0 | 0 | 0 | 0 |
| 0 | 0 | 0 | 0 | 0 | 0 | 0 | 0 | 0 |
| 0 | 0 | 0 | 0 | 0 | 0 | 0 | 0 | 9 |
| 0 | 0 | 0 | 0 | 0 | 0 | 0 | 0 | 0 |
| 0 | 0 | 0 | 0 | 0 | 0 | 0 | 0 | 0 |
| 0 | 0 | 0 | 0 | 0 | 0 | 0 | 0 | 0 |
| 0 | 0 | 0 | 0 | 0 | 0 | 0 | 0 | 0 |
| 0 | 0 | 0 | 0 | 0 | 0 | 0 | 0 | 0 |
| 0 | 0 | 0 | 0 | 0 | 0 | 0 | 0 | 0 |
| 0 | 0 | 0 | 0 | 0 | 0 | 0 | 0 | 0 |
| 0 | 0 | 0 | 0 | 0 | 0 | 0 | 0 | 0 |
| 0 | 0 | 0 | 0 | 0 | 0 | 0 | 0 | 0 |
| 0 | 0 | 0 | 0 | 0 | 0 | 0 | 0 | 0 |
| 0 | 0 | 0 | 0 | 0 | 0 | 0 | 0 | 0 |
| 0 | 0 | 0 | 0 | 0 | 0 | 0 | 0 | 0 |
| 0 | 0 | 0 | 0 | 0 | 0 | 0 | 0 | 4 |
| 0 | 0 | 0 | 0 | 6 | 0 | 0 | 0 | 0 |
| 0 | 0 | 0 | 0 | 0 | 0 | 0 | 0 | 0 |
| 0 | 0 | 0 | 0 | 0 | 0 | 0 | 0 | 0 |
| 0 | 0 | 0 | 0 | 0 | 0 | 0 | 0 | 0 |
| 0 | 0 | 0 | 0 | 0 | 0 | 0 | 0 | 0 |
| 0 | 0 | 0 | 0 | 0 | 0 | 0 | 0 | 0 |
| 0 | 0 | 0 | 0 | 0 | 0 | 0 | 0 | 0 |
| 0 | 0 | 0 | 0 | 0 | 0 | 0 | 0 | 0 |
| 0 | 0 | 0 | 0 | 7 | 0 | 0 | 0 | 0 |
| 0 | 0 | 0 | 0 | 0 | 0 | 0 | 0 | 0 |
| 0 | 0 | 0 | 0 | 0 | 0 | 0 | 0 | 0 |
| 0 | 0 | 0 | 0 | 0 | 0 | 0 | 0 | 0 |
| 0 | 0 | 0 | 0 | 0 | 0 | 0 | 0 | 0 |
| 0 | 0 | 0 | 0 | 0 | 0 | 0 | 0 | 0 |
| 0 | 0 | 0 | 0 | 0 | 0 | 0 | 0 | 0 |
| 0 | 0 | 0 | 0 | 0 | 0 | 0 | 0 | 0 |
| 0 | 0 | 0 | 0 | 0 | 0 | 0 | 0 | 0 |
| 0 | 0 | 0 | 0 | 0 | 0 | 0 | 0 | 0 |
| 0 | 0 | 0 | 0 | 0 | 0 | 0 | 0 | 0 |
| 0 | 0 | 0 | 0 | 0 | 0 | 0 | 0 | 0 |

| | | | | | | | | |
|---|---|---|---|---|---|---|---|---|
| 0 | 0 | 0 | 0 | 0 | 0 | 0 | 0 | 0 |
| 0 | 0 | 0 | 0 | 0 | 0 | 0 | 0 | 0 |
| 0 | 0 | 0 | 0 | 0 | 0 | 0 | 0 | 0 |
| 0 | 0 | 0 | 0 | 6 | 0 | 0 | 0 | 0 |
| 0 | 0 | 0 | 0 | 6 | 0 | 0 | 0 | 0 |
| 0 | 0 | 0 | 0 | 0 | 0 | 0 | 0 | 0 |
| 0 | 0 | 0 | 0 | 0 | 0 | 0 | 0 | 0 |
| 0 | 0 | 0 | 0 | 0 | 0 | 0 | 0 | 0 |
| 0 | 0 | 0 | 0 | 0 | 0 | 0 | 0 | 0 |
| 0 | 0 | 0 | 0 | 0 | 0 | 0 | 0 | 0 |
| 0 | 0 | 0 | 0 | 0 | 0 | 0 | 0 | 0 |
| 0 | 0 | 0 | 0 | 0 | 0 | 0 | 0 | 0 |
| 0 | 0 | 0 | 0 | 0 | 0 | 0 | 0 | 0 |
| 0 | 0 | 0 | 0 | 0 | 0 | 0 | 0 | 0 |
| 0 | 0 | 0 | 0 | 0 | 0 | 0 | 0 | 0 |
| 0 | 0 | 0 | 0 | 0 | 0 | 0 | 0 | 0 |
| 0 | 0 | 0 | 0 | 0 | 0 | 0 | 0 | 0 |
| 0 | 0 | 0 | 0 | 0 | 0 | 0 | 0 | 3 |
| 0 | 0 | 0 | 0 | 0 | 0 | 0 | 0 | 0 |
| 0 | 0 | 0 | 0 | 0 | 0 | 0 | 0 | 0 |
| 0 | 0 | 0 | 0 | 0 | 0 | 0 | 0 | 0 |
| 0 | 0 | 0 | 0 | 0 | 0 | 0 | 0 | 0 |
| 0 | 0 | 0 | 0 | 0 | 0 | 0 | 0 | 0 |
| 0 | 0 | 0 | 0 | 0 | 0 | 0 | 0 | 0 |
| 0 | 0 | 0 | 0 | 0 | 0 | 0 | 0 | 0 |
| 0 | 0 | 0 | 0 | 0 | 0 | 0 | 0 | 0 |
| 0 | 0 | 0 | 0 | 4 | 0 | 0 | 0 | 0 |
| 0 | 0 | 0 | 0 | 0 | 0 | 0 | 0 | 4 |
| 0 | 0 | 0 | 0 | 0 | 0 | 0 | 0 | 4 |
| 0 | 0 | 0 | 0 | 0 | 0 | 0 | 0 | 0 |
| 0 | 0 | 0 | 0 | 0 | 0 | 0 | 0 | 0 |
| 0 | 0 | 0 | 0 | 0 | 0 | 0 | 0 | 0 |
| 0 | 2 | 0 | 0 | 0 | 0 | 0 | 0 | 0 |
| 0 | 0 | 0 | 0 | 0 | 0 | 0 | 0 | 0 |
| 0 | 0 | 0 | 0 | 0 | 0 | 0 | 0 | 0 |
| 0 | 0 | 0 | 0 | 0 | 0 | 0 | 0 | 0 |
| 0 | 0 | 0 | 0 | 0 | 0 | 0 | 0 | 0 |
| 0 | 0 | 0 | 0 | 0 | 0 | 0 | 0 | 0 |
| 0 | 2 | 0 | 0 | 0 | 0 | 0 | 0 | 0 |
| 0 | 0 | 0 | 0 | 0 | 0 | 0 | 0 | 0 |
| 0 | 0 | 0 | 0 | 0 | 0 | 0 | 0 | 0 |
| 0 | 0 | 0 | 0 | 0 | 0 | 0 | 0 | 0 |

| 2-µM-$Cu^{2+}$ | | | 10%-$CH_4$ 10-µM-$Cu^{2+}$ | | | |
|---|---|---|---|---|---|---|
| **R06** | **R10** | **R15** | **CB** | **R07** | **R10** | **R14** |
| 3137 | 712 | 269 | 566 | 14985 | 25850 | 420 |
| 19664 | 7706 | 3925 | 233 | 2801 | 878 | 271 |
| 110 | 95 | 96 | 375 | 1239 | 534 | 366 |
| 488 | 12143 | 130 | 291 | 48 | 7 | 0 |
| 0 | 0 | 18 | 232 | 3612 | 6263 | 11122 |
| 1223 | 951 | 861 | 10 | 3303 | 1179 | 287 |
| 302 | 534 | 9266 | 0 | 0 | 0 | 0 |
| 43 | 80 | 117 | 98 | 0 | 60 | 110 |
| 0 | 0 | 0 | 1989 | 1578 | 3582 | 8247 |
| 9526 | 1074 | 1022 | 0 | 0 | 0 | 0 |
| 550 | 1764 | 2832 | 217 | 64 | 132 | 206 |
| 0 | 0 | 0 | 74 | 0 | 0 | 0 |
| 1077 | 1377 | 1463 | 0 | 510 | 224 | 0 |
| 1782 | 2385 | 2073 | 1539 | 211 | 118 | 50 |
| 60 | 51 | 150 | 59 | 554 | 33 | 0 |
| 134 | 110 | 472 | 213 | 31 | 133 | 54 |
| 13 | 113 | 118 | 405 | 0 | 17 | 38 |
| 213 | 352 | 439 | 345 | 165 | 199 | 178 |
| 82 | 48 | 56 | 0 | 1588 | 611 | 189 |
| 532 | 407 | 369 | 1780 | 643 | 666 | 217 |
| 0 | 0 | 0 | 392 | 0 | 0 | 0 |
| 5 | 0 | 0 | 25 | 0 | 0 | 0 |
| 151 | 503 | 722 | 423 | 213 | 189 | 232 |
| 0 | 34 | 307 | 26 | 0 | 0 | 0 |
| 695 | 859 | 1052 | 71 | 160 | 78 | 0 |
| 62 | 276 | 490 | 204 | 106 | 74 | 454 |
| 0 | 0 | 0 | 89 | 0 | 0 | 0 |
| 124 | 402 | 1008 | 1583 | 5 | 0 | 21 |
| 0 | 0 | 0 | 0 | 0 | 0 | 0 |
| 119 | 199 | 128 | 269 | 0 | 0 | 0 |
| 112 | 61 | 117 | 1190 | 538 | 151 | 0 |
| 0 | 0 | 0 | 0 | 0 | 0 | 0 |
| 10 | 0 | 11 | 0 | 0 | 20 | 13 |
| 0 | 0 | 0 | 7 | 0 | 0 | 40 |
| 0 | 0 | 0 | 80 | 0 | 0 | 0 |
| 4 | 0 | 8 | 131 | 0 | 0 | 0 |
| 0 | 0 | 0 | 293 | 299 | 386 | 209 |
| 0 | 0 | 32 | 2 | 0 | 0 | 0 |
| 20 | 12 | 0 | 2725 | 159 | 111 | 188 |
| 289 | 45 | 32 | 326 | 133 | 72 | 218 |
| 0 | 0 | 0 | 64 | 0 | 0 | 0 |
| 0 | 0 | 0 | 581 | 21 | 5 | 0 |
| 61 | 13 | 0 | 0 | 229 | 669 | 327 |
| 0 | 0 | 0 | 211 | 0 | 0 | 0 |
| 25 | 0 | 0 | 1044 | 0 | 0 | 0 |
| 236 | 548 | 766 | 252 | 85 | 26 | 11 |

| | | | | | | |
|---|---|---|---|---|---|---|
| 40 | 0 | 0 | 30 | 15 | 6 | 7 |
| 0 | 0 | 69 | 0 | 0 | 59 | 78 |
| 0 | 0 | 0 | 9 | 0 | 0 | 0 |
| 0 | 0 | 0 | 210 | 7 | 0 | 13 |
| 0 | 0 | 0 | 105 | 0 | 0 | 0 |
| 0 | 0 | 0 | 0 | 190 | 245 | 42 |
| 0 | 0 | 0 | 54 | 0 | 0 | 0 |
| 0 | 0 | 0 | 185 | 0 | 0 | 15 |
| 11 | 0 | 0 | 37 | 126 | 79 | 32 |
| 227 | 330 | 390 | 0 | 0 | 0 | 0 |
| 0 | 0 | 0 | 242 | 367 | 84 | 0 |
| 184 | 252 | 267 | 76 | 272 | 110 | 46 |
| 5 | 180 | 332 | 140 | 11 | 10 | 184 |
| 0 | 0 | 91 | 150 | 39 | 38 | 315 |
| 0 | 89 | 98 | 232 | 13 | 0 | 27 |
| 339 | 589 | 761 | 397 | 181 | 73 | 39 |
| 0 | 0 | 0 | 140 | 0 | 0 | 0 |
| 0 | 8 | 71 | 10 | 0 | 0 | 0 |
| 0 | 0 | 0 | 0 | 0 | 0 | 0 |
| 0 | 0 | 71 | 23 | 0 | 0 | 89 |
| 0 | 0 | 8 | 195 | 0 | 0 | 0 |
| 0 | 0 | 0 | 65 | 0 | 0 | 0 |
| 4 | 0 | 0 | 0 | 0 | 0 | 0 |
| 93 | 220 | 313 | 155 | 59 | 71 | 378 |
| 4 | 0 | 0 | 28 | 0 | 0 | 0 |
| 69 | 37 | 46 | 307 | 68 | 139 | 383 |
| 0 | 26 | 128 | 0 | 0 | 16 | 236 |
| 0 | 0 | 0 | 0 | 0 | 0 | 0 |
| 163 | 21 | 0 | 0 | 630 | 111 | 51 |
| 0 | 7 | 15 | 152 | 123 | 40 | 0 |
| 37 | 358 | 250 | 0 | 0 | 0 | 0 |
| 0 | 0 | 0 | 30 | 0 | 0 | 0 |
| 0 | 0 | 0 | 72 | 0 | 0 | 0 |
| 0 | 0 | 0 | 4 | 0 | 0 | 0 |
| 0 | 0 | 0 | 0 | 0 | 0 | 0 |
| 0 | 0 | 0 | 5 | 0 | 0 | 0 |
| 0 | 126 | 136 | 34 | 0 | 0 | 0 |
| 67 | 0 | 10 | 0 | 0 | 392 | 581 |
| 7 | 0 | 0 | 98 | 4 | 0 | 0 |
| 0 | 0 | 0 | 94 | 0 | 0 | 49 |
| 17 | 0 | 0 | 46 | 0 | 0 | 0 |
| 0 | 0 | 0 | 0 | 0 | 0 | 0 |
| 205 | 180 | 117 | 410 | 112 | 79 | 73 |
| 0 | 0 | 0 | 186 | 0 | 0 | 0 |
| 0 | 0 | 0 | 0 | 0 | 0 | 0 |
| 0 | 0 | 0 | 70 | 0 | 0 | 0 |
| 0 | 0 | 0 | 225 | 109 | 114 | 0 |
| 0 | 0 | 0 | 263 | 0 | 0 | 0 |
| 0 | 0 | 0 | 210 | 0 | 0 | 0 |
| 27 | 0 | 0 | 117 | 268 | 17 | 0 |

| | | | | | | |
|---|---|---|---|---|---|---|
| 0 | 0 | 0 | 108 | 0 | 0 | 0 |
| 8 | 0 | 3 | 36 | 0 | 16 | 0 |
| 0 | 0 | 0 | 200 | 0 | 0 | 0 |
| 0 | 0 | 0 | 533 | 0 | 0 | 0 |
| 0 | 0 | 0 | 606 | 0 | 0 | 0 |
| 0 | 0 | 97 | 0 | 0 | 138 | 347 |
| 0 | 0 | 0 | 216 | 0 | 0 | 0 |
| 0 | 0 | 23 | 79 | 5 | 0 | 0 |
| 0 | 0 | 0 | 0 | 0 | 0 | 0 |
| 0 | 0 | 33 | 57 | 0 | 0 | 58 |
| 0 | 0 | 0 | 353 | 18 | 0 | 0 |
| 0 | 0 | 0 | 69 | 5 | 0 | 0 |
| 0 | 0 | 0 | 132 | 40 | 31 | 23 |
| 4 | 0 | 0 | 131 | 48 | 10 | 0 |
| 0 | 0 | 0 | 278 | 12 | 0 | 0 |
| 0 | 0 | 0 | 75 | 14 | 10 | 0 |
| 0 | 0 | 0 | 52 | 0 | 0 | 0 |
| 0 | 0 | 169 | 8 | 0 | 0 | 0 |
| 0 | 0 | 0 | 160 | 0 | 16 | 61 |
| 117 | 56 | 92 | 306 | 77 | 27 | 0 |
| 0 | 0 | 0 | 9 | 2 | 0 | 0 |
| 0 | 0 | 0 | 0 | 0 | 0 | 0 |
| 0 | 0 | 0 | 91 | 199 | 50 | 15 |
| 0 | 0 | 0 | 0 | 12 | 0 | 0 |
| 341 | 0 | 0 | 0 | 25 | 0 | 0 |
| 0 | 0 | 0 | 117 | 0 | 0 | 0 |
| 0 | 0 | 0 | 0 | 0 | 0 | 0 |
| 0 | 0 | 0 | 72 | 0 | 0 | 0 |
| 24 | 0 | 0 | 444 | 0 | 0 | 0 |
| 0 | 0 | 0 | 8 | 0 | 0 | 0 |
| 0 | 0 | 0 | 0 | 0 | 0 | 0 |
| 0 | 0 | 0 | 419 | 0 | 0 | 0 |
| 17 | 0 | 0 | 78 | 0 | 0 | 0 |
| 0 | 0 | 0 | 0 | 9 | 23 | 4 |
| 0 | 0 | 0 | 0 | 0 | 0 | 0 |
| 0 | 0 | 0 | 0 | 0 | 0 | 0 |
| 0 | 0 | 0 | 25 | 0 | 0 | 0 |
| 0 | 0 | 0 | 240 | 0 | 0 | 0 |
| 0 | 0 | 6 | 228 | 0 | 0 | 0 |
| 0 | 0 | 0 | 16 | 105 | 18 | 0 |
| 0 | 0 | 0 | 293 | 0 | 0 | 0 |
| 0 | 0 | 0 | 0 | 0 | 0 | 0 |
| 0 | 0 | 0 | 72 | 34 | 25 | 0 |
| 262 | 127 | 349 | 928 | 713 | 568 | 913 |
| 0 | 0 | 0 | 98 | 0 | 0 | 0 |
| 0 | 0 | 0 | 149 | 0 | 0 | 0 |
| 27 | 74 | 45 | 43 | 20 | 0 | 0 |
| 0 | 0 | 0 | 0 | 0 | 0 | 0 |
| 0 | 0 | 0 | 89 | 0 | 0 | 0 |
| 0 | 0 | 0 | 0 | 0 | 0 | 0 |

| | | | | | | |
|---|---|---|---|---|---|---|
| 0 | 0 | 0 | 122 | 0 | 0 | 0 |
| 7 | 66 | 147 | 176 | 0 | 0 | 0 |
| 0 | 0 | 0 | 79 | 0 | 0 | 0 |
| 0 | 0 | 0 | 5 | 0 | 0 | 0 |
| 70 | 0 | 0 | 0 | 0 | 0 | 0 |
| 0 | 0 | 0 | 230 | 0 | 0 | 0 |
| 0 | 0 | 0 | 0 | 0 | 0 | 0 |
| 32 | 0 | 0 | 222 | 114 | 11 | 0 |
| 0 | 6 | 408 | 0 | 0 | 0 | 0 |
| 0 | 0 | 0 | 19 | 0 | 0 | 0 |
| 0 | 0 | 0 | 16 | 0 | 0 | 0 |
| 0 | 0 | 0 | 0 | 0 | 0 | 0 |
| 0 | 0 | 0 | 0 | 0 | 0 | 0 |
| 0 | 0 | 0 | 160 | 0 | 0 | 0 |
| 8 | 0 | 0 | 239 | 31 | 0 | 0 |
| 0 | 0 | 0 | 0 | 0 | 0 | 0 |
| 0 | 0 | 0 | 21 | 0 | 0 | 0 |
| 0 | 0 | 0 | 80 | 0 | 0 | 0 |
| 0 | 0 | 5 | 323 | 0 | 0 | 9 |
| 0 | 0 | 0 | 116 | 172 | 0 | 0 |
| 0 | 0 | 0 | 72 | 0 | 0 | 0 |
| 0 | 30 | 108 | 0 | 0 | 0 | 0 |
| 0 | 0 | 0 | 0 | 0 | 0 | 0 |
| 0 | 0 | 0 | 114 | 0 | 0 | 0 |
| 0 | 0 | 0 | 58 | 0 | 0 | 0 |
| 0 | 0 | 0 | 46 | 0 | 0 | 0 |
| 0 | 0 | 0 | 266 | 0 | 0 | 0 |
| 0 | 9 | 9 | 0 | 0 | 0 | 0 |
| 0 | 0 | 0 | 0 | 0 | 0 | 0 |
| 0 | 0 | 0 | 12 | 0 | 0 | 0 |
| 0 | 0 | 0 | 63 | 0 | 0 | 0 |
| 0 | 0 | 0 | 10 | 0 | 0 | 0 |
| 0 | 0 | 0 | 20 | 0 | 0 | 0 |
| 0 | 0 | 0 | 170 | 0 | 0 | 0 |
| 12 | 82 | 83 | 21 | 13 | 24 | 36 |
| 0 | 0 | 0 | 273 | 0 | 0 | 0 |
| 0 | 37 | 32 | 4 | 95 | 9 | 0 |
| 0 | 0 | 0 | 50 | 0 | 0 | 0 |
| 0 | 0 | 0 | 8 | 0 | 0 | 0 |
| 0 | 0 | 0 | 16 | 0 | 0 | 0 |
| 0 | 0 | 0 | 10 | 0 | 0 | 0 |
| 0 | 0 | 0 | 25 | 0 | 0 | 0 |
| 0 | 0 | 0 | 0 | 0 | 0 | 0 |
| 0 | 0 | 0 | 7 | 0 | 0 | 0 |
| 0 | 0 | 0 | 57 | 0 | 0 | 0 |
| 0 | 3 | 4 | 193 | 0 | 0 | 0 |
| 0 | 0 | 0 | 117 | 0 | 0 | 0 |
| 0 | 0 | 0 | 77 | 0 | 0 | 0 |
| 0 | 0 | 0 | 48 | 0 | 0 | 0 |
| 0 | 0 | 0 | 6 | 0 | 0 | 0 |

| | | | | | | |
|---|---|---|---|---|---|---|
| 0 | 0 | 0 | 41 | 0 | 0 | 0 |
| 0 | 0 | 0 | 0 | 0 | 0 | 0 |
| 0 | 0 | 0 | 0 | 0 | 0 | 0 |
| 0 | 0 | 0 | 23 | 0 | 0 | 0 |
| 0 | 0 | 0 | 18 | 0 | 0 | 0 |
| 0 | 0 | 0 | 98 | 114 | 0 | 0 |
| 0 | 0 | 0 | 47 | 0 | 0 | 0 |
| 0 | 0 | 0 | 0 | 0 | 0 | 0 |
| 0 | 0 | 0 | 0 | 0 | 0 | 0 |
| 0 | 0 | 0 | 0 | 0 | 0 | 0 |
| 0 | 0 | 0 | 0 | 0 | 0 | 0 |
| 0 | 0 | 0 | 21 | 0 | 0 | 0 |
| 0 | 0 | 0 | 34 | 0 | 0 | 0 |
| 0 | 0 | 0 | 99 | 20 | 22 | 0 |
| 0 | 17 | 116 | 0 | 0 | 0 | 0 |
| 0 | 0 | 0 | 7 | 0 | 0 | 0 |
| 0 | 0 | 0 | 132 | 0 | 0 | 0 |
| 0 | 0 | 0 | 0 | 0 | 0 | 0 |
| 0 | 0 | 0 | 16 | 0 | 0 | 0 |
| 0 | 0 | 0 | 0 | 0 | 0 | 0 |
| 0 | 0 | 0 | 0 | 0 | 0 | 0 |
| 0 | 0 | 0 | 192 | 0 | 0 | 0 |
| 0 | 0 | 0 | 76 | 0 | 0 | 0 |
| 0 | 0 | 0 | 0 | 0 | 0 | 0 |
| 0 | 0 | 0 | 0 | 0 | 0 | 0 |
| 0 | 0 | 0 | 0 | 0 | 0 | 0 |
| 0 | 0 | 0 | 0 | 0 | 0 | 0 |
| 0 | 0 | 24 | 0 | 0 | 0 | 0 |
| 0 | 0 | 0 | 0 | 0 | 0 | 112 |
| 0 | 0 | 0 | 0 | 0 | 0 | 0 |
| 0 | 0 | 0 | 191 | 0 | 0 | 0 |
| 0 | 0 | 0 | 75 | 6 | 0 | 11 |
| 0 | 0 | 0 | 65 | 0 | 0 | 0 |
| 0 | 0 | 0 | 0 | 0 | 0 | 0 |
| 0 | 0 | 0 | 0 | 0 | 0 | 0 |
| 0 | 0 | 0 | 0 | 0 | 0 | 0 |
| 0 | 0 | 0 | 23 | 0 | 0 | 0 |
| 0 | 0 | 0 | 14 | 0 | 0 | 0 |
| 0 | 0 | 0 | 0 | 0 | 0 | 166 |
| 0 | 0 | 0 | 46 | 0 | 0 | 0 |
| 0 | 0 | 0 | 0 | 0 | 0 | 0 |
| 0 | 0 | 0 | 14 | 0 | 0 | 0 |
| 0 | 0 | 0 | 0 | 0 | 0 | 0 |
| 54 | 11 | 0 | 16 | 48 | 16 | 12 |
| 0 | 0 | 0 | 0 | 0 | 0 | 0 |
| 0 | 0 | 0 | 62 | 0 | 0 | 0 |
| 0 | 0 | 0 | 6 | 0 | 0 | 0 |
| 0 | 0 | 0 | 264 | 0 | 0 | 0 |
| 0 | 0 | 0 | 0 | 0 | 0 | 0 |
| 0 | 0 | 0 | 0 | 0 | 0 | 0 |

| | | | | | | |
|---|---|---|---|---|---|---|
| 85 | 36 | 15 | 8 | 234 | 21 | 0 |
| 0 | 0 | 0 | 150 | 0 | 0 | 0 |
| 0 | 0 | 0 | 9 | 0 | 0 | 0 |
| 0 | 36 | 12 | 6 | 0 | 0 | 0 |
| 0 | 0 | 0 | 23 | 0 | 0 | 0 |
| 0 | 0 | 0 | 29 | 0 | 0 | 0 |
| 0 | 0 | 0 | 0 | 0 | 0 | 0 |
| 0 | 0 | 0 | 70 | 0 | 0 | 0 |
| 18 | 0 | 0 | 0 | 0 | 0 | 0 |
| 0 | 0 | 0 | 84 | 0 | 0 | 0 |
| 0 | 0 | 0 | 12 | 0 | 0 | 0 |
| 0 | 0 | 0 | 0 | 0 | 0 | 0 |
| 0 | 0 | 0 | 0 | 0 | 0 | 0 |
| 0 | 0 | 0 | 110 | 0 | 0 | 0 |
| 0 | 0 | 0 | 117 | 0 | 0 | 0 |
| 0 | 0 | 0 | 33 | 0 | 0 | 0 |
| 0 | 0 | 0 | 48 | 0 | 0 | 0 |
| 0 | 0 | 0 | 0 | 0 | 0 | 0 |
| 0 | 0 | 0 | 0 | 0 | 0 | 0 |
| 0 | 0 | 0 | 0 | 0 | 0 | 0 |
| 0 | 0 | 0 | 0 | 0 | 0 | 0 |
| 0 | 0 | 0 | 0 | 0 | 0 | 2 |
| 0 | 0 | 0 | 73 | 0 | 0 | 0 |
| 0 | 0 | 0 | 0 | 0 | 0 | 0 |
| 0 | 0 | 0 | 0 | 0 | 0 | 0 |
| 0 | 0 | 0 | 0 | 0 | 0 | 0 |
| 0 | 0 | 0 | 0 | 0 | 0 | 0 |
| 0 | 0 | 0 | 0 | 0 | 0 | 0 |
| 0 | 0 | 0 | 0 | 0 | 0 | 0 |
| 0 | 0 | 0 | 63 | 0 | 0 | 0 |
| 0 | 0 | 0 | 178 | 0 | 0 | 0 |
| 0 | 0 | 0 | 0 | 0 | 0 | 0 |
| 0 | 0 | 1 | 92 | 4 | 15 | 0 |
| 0 | 0 | 0 | 0 | 21 | 0 | 0 |
| 0 | 0 | 0 | 91 | 0 | 0 | 5 |
| 0 | 0 | 0 | 0 | 0 | 0 | 0 |
| 0 | 0 | 0 | 0 | 0 | 62 | 48 |
| 0 | 0 | 0 | 14 | 0 | 0 | 0 |
| 0 | 0 | 0 | 195 | 0 | 0 | 0 |
| 0 | 0 | 0 | 32 | 0 | 0 | 0 |
| 0 | 0 | 29 | 0 | 0 | 0 | 25 |
| 18 | 0 | 0 | 14 | 0 | 0 | 0 |
| 0 | 0 | 0 | 0 | 0 | 0 | 0 |
| 0 | 0 | 14 | 0 | 0 | 0 | 0 |
| 0 | 0 | 0 | 0 | 0 | 0 | 0 |
| 0 | 0 | 0 | 0 | 0 | 0 | 0 |
| 0 | 0 | 0 | 0 | 0 | 0 | 0 |
| 0 | 0 | 0 | 0 | 0 | 0 | 0 |
| 0 | 0 | 0 | 0 | 0 | 0 | 0 |
| 0 | 0 | 0 | 57 | 0 | 0 | 0 |

| | | | | | | |
|---|---|---|---|---|---|---|
| 0 | 0 | 0 | 0 | 0 | 0 | 0 |
| 0 | 0 | 0 | 32 | 0 | 0 | 0 |
| 0 | 0 | 0 | 34 | 0 | 0 | 0 |
| 0 | 0 | 0 | 6 | 0 | 0 | 0 |
| 0 | 0 | 0 | 0 | 0 | 0 | 0 |
| 0 | 0 | 0 | 0 | 0 | 0 | 0 |
| 0 | 0 | 0 | 0 | 0 | 0 | 0 |
| 0 | 0 | 0 | 0 | 0 | 0 | 0 |
| 0 | 0 | 0 | 12 | 0 | 0 | 0 |
| 0 | 0 | 4 | 120 | 0 | 0 | 0 |
| 0 | 0 | 0 | 9 | 0 | 3 | 0 |
| 0 | 0 | 0 | 0 | 0 | 0 | 0 |
| 0 | 0 | 0 | 20 | 0 | 0 | 0 |
| 0 | 0 | 0 | 0 | 0 | 0 | 0 |
| 0 | 0 | 0 | 17 | 0 | 0 | 0 |
| 0 | 0 | 0 | 0 | 0 | 0 | 0 |
| 0 | 0 | 0 | 0 | 0 | 0 | 0 |
| 0 | 0 | 0 | 0 | 0 | 0 | 0 |
| 0 | 0 | 0 | 15 | 0 | 0 | 0 |
| 0 | 0 | 0 | 0 | 0 | 0 | 0 |
| 0 | 0 | 0 | 0 | 0 | 0 | 0 |
| 0 | 0 | 0 | 0 | 0 | 0 | 0 |
| 0 | 0 | 0 | 0 | 0 | 0 | 0 |
| 0 | 0 | 0 | 0 | 0 | 0 | 0 |
| 0 | 0 | 0 | 0 | 0 | 19 | 0 |
| 0 | 0 | 0 | 0 | 0 | 0 | 0 |
| 0 | 0 | 0 | 0 | 0 | 0 | 0 |
| 0 | 0 | 0 | 0 | 0 | 0 | 0 |
| 0 | 0 | 0 | 5 | 0 | 3 | 0 |
| 0 | 0 | 0 | 0 | 0 | 0 | 0 |
| 0 | 0 | 0 | 0 | 0 | 0 | 0 |
| 0 | 0 | 14 | 18 | 0 | 0 | 36 |
| 0 | 0 | 0 | 23 | 0 | 0 | 0 |
| 0 | 0 | 0 | 0 | 0 | 0 | 0 |
| 0 | 0 | 0 | 20 | 0 | 0 | 0 |
| 0 | 0 | 0 | 0 | 0 | 0 | 0 |
| 0 | 0 | 0 | 68 | 0 | 0 | 0 |
| 0 | 0 | 0 | 16 | 0 | 0 | 0 |
| 0 | 0 | 0 | 0 | 0 | 0 | 0 |
| 0 | 0 | 0 | 0 | 0 | 0 | 0 |
| 0 | 0 | 0 | 0 | 0 | 0 | 0 |
| 0 | 0 | 0 | 22 | 0 | 0 | 0 |
| 0 | 0 | 0 | 0 | 0 | 0 | 0 |
| 0 | 0 | 0 | 42 | 0 | 0 | 0 |
| 0 | 0 | 0 | 21 | 0 | 0 | 0 |
| 82 | 0 | 0 | 0 | 0 | 0 | 0 |
| 0 | 0 | 0 | 5 | 0 | 0 | 0 |
| 0 | 0 | 0 | 0 | 0 | 0 | 0 |
| 0 | 0 | 0 | 34 | 0 | 0 | 0 |
| 0 | 0 | 0 | 0 | 0 | 0 | 0 |

| | | | | | | |
|---|---|---|---|---|---|---|
| 0 | 0 | 0 | 10 | 0 | 0 | 0 |
| 0 | 0 | 0 | 0 | 0 | 0 | 0 |
| 0 | 0 | 0 | 0 | 0 | 0 | 0 |
| 0 | 0 | 0 | 0 | 0 | 0 | 0 |
| 0 | 0 | 0 | 0 | 0 | 0 | 0 |
| 0 | 0 | 0 | 0 | 23 | 0 | 0 |
| 0 | 0 | 0 | 42 | 0 | 0 | 0 |
| 0 | 0 | 0 | 13 | 0 | 0 | 0 |
| 0 | 0 | 0 | 0 | 0 | 0 | 0 |
| 0 | 0 | 0 | 0 | 0 | 0 | 0 |
| 0 | 0 | 0 | 9 | 0 | 0 | 0 |
| 0 | 0 | 0 | 0 | 0 | 0 | 0 |
| 0 | 0 | 0 | 0 | 0 | 0 | 0 |
| 0 | 0 | 0 | 18 | 0 | 0 | 0 |
| 0 | 0 | 0 | 0 | 0 | 0 | 0 |
| 0 | 0 | 0 | 0 | 0 | 0 | 0 |
| 0 | 0 | 0 | 0 | 0 | 0 | 0 |
| 0 | 0 | 0 | 15 | 0 | 0 | 0 |
| 0 | 0 | 0 | 72 | 0 | 0 | 0 |
| 0 | 0 | 12 | 19 | 0 | 0 | 0 |
| 0 | 0 | 0 | 0 | 0 | 0 | 0 |
| 0 | 0 | 0 | 0 | 41 | 30 | 0 |
| 0 | 0 | 0 | 0 | 0 | 0 | 0 |
| 0 | 0 | 0 | 8 | 0 | 0 | 0 |
| 0 | 0 | 0 | 0 | 0 | 0 | 0 |
| 0 | 0 | 0 | 0 | 0 | 0 | 0 |
| 0 | 0 | 0 | 0 | 0 | 0 | 0 |
| 0 | 0 | 0 | 0 | 0 | 0 | 0 |
| 0 | 0 | 0 | 0 | 0 | 0 | 0 |
| 0 | 0 | 0 | 28 | 0 | 0 | 0 |
| 0 | 0 | 0 | 11 | 0 | 0 | 0 |
| 0 | 0 | 0 | 0 | 0 | 0 | 0 |
| 0 | 0 | 0 | 0 | 0 | 0 | 0 |
| 0 | 0 | 0 | 0 | 0 | 0 | 0 |
| 0 | 0 | 0 | 0 | 0 | 0 | 0 |
| 0 | 0 | 0 | 0 | 0 | 0 | 0 |
| 0 | 0 | 0 | 0 | 0 | 0 | 0 |
| 0 | 0 | 0 | 0 | 0 | 0 | 0 |
| 0 | 0 | 0 | 20 | 0 | 0 | 0 |
| 0 | 0 | 0 | 72 | 0 | 0 | 0 |
| 0 | 0 | 0 | 0 | 0 | 0 | 0 |
| 0 | 0 | 0 | 0 | 0 | 0 | 66 |
| 0 | 0 | 0 | 0 | 0 | 0 | 0 |
| 0 | 0 | 0 | 0 | 0 | 0 | 0 |
| 0 | 0 | 0 | 0 | 0 | 0 | 0 |
| 0 | 0 | 0 | 67 | 0 | 0 | 0 |
| 0 | 0 | 0 | 26 | 0 | 0 | 18 |
| 0 | 0 | 0 | 0 | 0 | 0 | 0 |
| 0 | 0 | 0 | 0 | 0 | 0 | 0 |
| 0 | 0 | 0 | 20 | 0 | 0 | 0 |

| | | | | | | |
|---|---|---|---|---|---|---|
| 0 | 0 | 0 | 0 | 0 | 0 | 0 |
| 0 | 0 | 0 | 0 | 0 | 0 | 0 |
| 0 | 0 | 0 | 0 | 0 | 0 | 0 |
| 0 | 30 | 34 | 0 | 0 | 0 | 0 |
| 0 | 0 | 0 | 0 | 0 | 0 | 0 |
| 0 | 0 | 0 | 15 | 0 | 0 | 0 |
| 0 | 0 | 0 | 0 | 0 | 0 | 0 |
| 0 | 0 | 0 | 26 | 43 | 0 | 0 |
| 0 | 0 | 0 | 0 | 0 | 0 | 0 |
| 0 | 0 | 0 | 0 | 0 | 0 | 0 |
| 0 | 0 | 0 | 6 | 0 | 0 | 0 |
| 0 | 0 | 0 | 0 | 0 | 0 | 0 |
| 0 | 0 | 0 | 16 | 0 | 0 | 0 |
| 0 | 0 | 0 | 19 | 0 | 0 | 0 |
| 0 | 0 | 0 | 31 | 0 | 0 | 0 |
| 0 | 0 | 0 | 6 | 0 | 0 | 0 |
| 0 | 0 | 0 | 0 | 0 | 0 | 0 |
| 0 | 0 | 4 | 0 | 0 | 0 | 0 |
| 0 | 0 | 0 | 0 | 0 | 0 | 0 |
| 0 | 0 | 0 | 0 | 0 | 0 | 0 |
| 0 | 0 | 0 | 8 | 0 | 0 | 0 |
| 0 | 0 | 0 | 0 | 0 | 0 | 0 |
| 0 | 0 | 0 | 59 | 0 | 0 | 0 |
| 0 | 0 | 0 | 13 | 0 | 0 | 0 |
| 0 | 0 | 0 | 0 | 0 | 0 | 0 |
| 0 | 0 | 0 | 0 | 0 | 0 | 0 |
| 0 | 0 | 0 | 0 | 0 | 0 | 0 |
| 0 | 0 | 0 | 0 | 0 | 0 | 0 |
| 0 | 0 | 0 | 0 | 0 | 0 | 0 |
| 0 | 0 | 0 | 0 | 0 | 0 | 0 |
| 0 | 0 | 0 | 12 | 0 | 0 | 0 |
| 0 | 0 | 0 | 0 | 0 | 0 | 0 |
| 0 | 0 | 0 | 0 | 0 | 0 | 0 |
| 0 | 0 | 0 | 0 | 0 | 0 | 0 |
| 0 | 0 | 0 | 9 | 0 | 0 | 0 |
| 0 | 0 | 0 | 0 | 0 | 0 | 0 |
| 0 | 0 | 0 | 0 | 0 | 0 | 0 |
| 0 | 0 | 6 | 0 | 0 | 0 | 0 |
| 0 | 0 | 0 | 0 | 0 | 0 | 0 |
| 10 | 6 | 28 | 0 | 0 | 0 | 0 |
| 0 | 0 | 0 | 0 | 0 | 0 | 0 |
| 0 | 0 | 0 | 0 | 0 | 0 | 0 |
| 9 | 0 | 0 | 0 | 0 | 0 | 0 |
| 0 | 0 | 0 | 12 | 0 | 0 | 0 |
| 0 | 0 | 0 | 0 | 0 | 0 | 0 |
| 0 | 0 | 0 | 0 | 0 | 0 | 0 |
| 0 | 0 | 0 | 0 | 0 | 0 | 0 |
| 0 | 0 | 0 | 0 | 0 | 0 | 0 |
| 0 | 0 | 0 | 0 | 0 | 0 | 0 |
| 0 | 0 | 0 | 0 | 0 | 0 | 0 |

| | | | | | | |
|---|---|---|---|---|---|---|
| 0 | 0 | 0 | 0 | 0 | 0 | 0 |
| 0 | 0 | 0 | 8 | 0 | 0 | 0 |
| 0 | 0 | 0 | 16 | 0 | 0 | 0 |
| 0 | 0 | 0 | 28 | 0 | 0 | 0 |
| 0 | 0 | 0 | 0 | 0 | 0 | 0 |
| 0 | 0 | 0 | 11 | 0 | 0 | 0 |
| 0 | 0 | 0 | 0 | 0 | 0 | 0 |
| 0 | 0 | 0 | 0 | 0 | 0 | 0 |
| 0 | 0 | 0 | 0 | 0 | 0 | 0 |
| 0 | 0 | 0 | 0 | 0 | 0 | 0 |
| 0 | 0 | 0 | 0 | 0 | 0 | 0 |
| 0 | 0 | 0 | 0 | 0 | 0 | 0 |
| 0 | 0 | 0 | 0 | 0 | 0 | 0 |
| 0 | 0 | 0 | 0 | 0 | 0 | 0 |
| 0 | 0 | 0 | 7 | 0 | 0 | 0 |
| 0 | 0 | 0 | 12 | 0 | 0 | 0 |
| 0 | 0 | 0 | 0 | 0 | 0 | 0 |
| 0 | 0 | 0 | 84 | 0 | 0 | 0 |
| 0 | 0 | 0 | 0 | 0 | 0 | 0 |
| 0 | 0 | 0 | 0 | 0 | 0 | 0 |
| 0 | 0 | 0 | 0 | 0 | 0 | 0 |
| 0 | 0 | 0 | 16 | 0 | 0 | 0 |
| 0 | 0 | 0 | 14 | 0 | 0 | 0 |
| 0 | 0 | 0 | 0 | 0 | 0 | 0 |
| 0 | 0 | 0 | 0 | 0 | 0 | 0 |
| 0 | 0 | 0 | 9 | 0 | 0 | 0 |
| 0 | 0 | 0 | 0 | 0 | 0 | 0 |
| 0 | 0 | 0 | 0 | 0 | 0 | 0 |
| 0 | 0 | 0 | 0 | 0 | 0 | 0 |
| 0 | 0 | 0 | 2 | 0 | 0 | 0 |
| 0 | 0 | 0 | 0 | 0 | 0 | 0 |
| 0 | 0 | 0 | 0 | 0 | 0 | 0 |
| 0 | 0 | 0 | 0 | 0 | 0 | 0 |
| 0 | 0 | 0 | 30 | 0 | 0 | 0 |
| 0 | 0 | 0 | 0 | 0 | 0 | 0 |
| 0 | 0 | 0 | 0 | 0 | 0 | 0 |
| 0 | 0 | 0 | 8 | 0 | 0 | 0 |
| 0 | 0 | 0 | 0 | 0 | 0 | 0 |
| 0 | 0 | 0 | 0 | 0 | 0 | 0 |
| 0 | 0 | 0 | 0 | 0 | 0 | 0 |
| 0 | 0 | 0 | 5 | 0 | 0 | 0 |
| 0 | 0 | 0 | 0 | 0 | 0 | 0 |
| 0 | 0 | 0 | 0 | 0 | 0 | 0 |
| 0 | 0 | 0 | 0 | 0 | 0 | 0 |
| 0 | 0 | 0 | 19 | 0 | 0 | 0 |
| 0 | 23 | 21 | 0 | 0 | 0 | 0 |
| 0 | 0 | 0 | 44 | 0 | 0 | 0 |
| 0 | 0 | 0 | 11 | 0 | 0 | 0 |
| 0 | 0 | 0 | 43 | 0 | 0 | 0 |
| 0 | 0 | 0 | 0 | 0 | 0 | 0 |

| | | | | | | |
|---|---|---|---|---|---|---|
| 0 | 0 | 0 | 0 | 0 | 0 | 0 |
| 0 | 0 | 0 | 0 | 0 | 0 | 0 |
| 0 | 0 | 0 | 0 | 0 | 0 | 0 |
| 0 | 0 | 0 | 0 | 0 | 0 | 0 |
| 0 | 0 | 0 | 12 | 0 | 0 | 0 |
| 0 | 0 | 0 | 0 | 0 | 0 | 0 |
| 0 | 0 | 0 | 0 | 0 | 0 | 0 |
| 0 | 0 | 0 | 11 | 0 | 0 | 0 |
| 0 | 0 | 0 | 0 | 0 | 0 | 0 |
| 0 | 0 | 0 | 12 | 0 | 0 | 0 |
| 0 | 0 | 0 | 0 | 0 | 0 | 0 |
| 0 | 0 | 0 | 0 | 0 | 0 | 0 |
| 0 | 0 | 0 | 0 | 0 | 0 | 0 |
| 0 | 0 | 0 | 0 | 0 | 0 | 41 |
| 0 | 0 | 0 | 4 | 0 | 0 | 0 |
| 0 | 0 | 0 | 0 | 0 | 0 | 0 |
| 0 | 0 | 0 | 0 | 0 | 0 | 0 |
| 0 | 0 | 0 | 9 | 0 | 0 | 0 |
| 0 | 0 | 0 | 4 | 0 | 0 | 0 |
| 0 | 0 | 0 | 24 | 0 | 0 | 0 |
| 0 | 0 | 0 | 0 | 0 | 0 | 0 |
| 0 | 0 | 0 | 29 | 0 | 0 | 0 |
| 0 | 0 | 0 | 0 | 0 | 0 | 0 |
| 0 | 0 | 0 | 0 | 0 | 0 | 0 |
| 0 | 0 | 0 | 0 | 0 | 0 | 0 |
| 0 | 0 | 0 | 0 | 0 | 0 | 0 |
| 0 | 0 | 0 | 0 | 0 | 0 | 0 |
| 0 | 0 | 0 | 0 | 0 | 0 | 0 |
| 0 | 0 | 0 | 0 | 0 | 0 | 0 |
| 0 | 0 | 0 | 0 | 0 | 0 | 0 |
| 0 | 0 | 0 | 0 | 0 | 0 | 0 |
| 0 | 0 | 0 | 0 | 0 | 0 | 0 |
| 0 | 0 | 0 | 0 | 0 | 0 | 0 |
| 0 | 0 | 0 | 0 | 0 | 0 | 0 |
| 0 | 0 | 0 | 15 | 0 | 0 | 0 |
| 0 | 0 | 0 | 0 | 0 | 0 | 0 |
| 0 | 0 | 0 | 13 | 0 | 0 | 0 |
| 0 | 0 | 0 | 20 | 0 | 0 | 0 |
| 0 | 0 | 0 | 0 | 0 | 0 | 0 |
| 0 | 0 | 0 | 0 | 0 | 0 | 0 |
| 0 | 0 | 0 | 8 | 0 | 0 | 0 |
| 0 | 0 | 0 | 0 | 0 | 0 | 0 |
| 0 | 0 | 0 | 0 | 0 | 0 | 0 |
| 0 | 0 | 0 | 0 | 0 | 0 | 0 |
| 0 | 0 | 0 | 0 | 0 | 0 | 0 |
| 0 | 0 | 0 | 0 | 0 | 0 | 0 |
| 0 | 0 | 0 | 0 | 0 | 0 | 0 |
| 0 | 0 | 0 | 0 | 0 | 0 | 0 |
| 0 | 0 | 0 | 26 | 0 | 0 | 2 |
| 0 | 0 | 0 | 0 | 0 | 0 | 0 |

| | | | | | | |
|---|---|---|---|---|---|---|
| 0 | 0 | 0 | 0 | 0 | 0 | 0 |
| 0 | 0 | 0 | 0 | 0 | 0 | 0 |
| 0 | 0 | 0 | 0 | 0 | 0 | 0 |
| 0 | 0 | 0 | 13 | 0 | 0 | 0 |
| 0 | 0 | 0 | 0 | 0 | 36 | 0 |
| 0 | 0 | 0 | 0 | 0 | 0 | 0 |
| 0 | 0 | 0 | 0 | 0 | 0 | 0 |
| 0 | 0 | 0 | 0 | 0 | 0 | 0 |
| 0 | 0 | 0 | 0 | 0 | 0 | 0 |
| 0 | 0 | 0 | 45 | 5 | 2 | 18 |
| 0 | 0 | 0 | 32 | 0 | 0 | 0 |
| 0 | 0 | 0 | 13 | 0 | 0 | 0 |
| 0 | 0 | 0 | 6 | 0 | 3 | 0 |
| 0 | 0 | 0 | 0 | 0 | 0 | 0 |
| 0 | 0 | 0 | 18 | 0 | 0 | 0 |
| 0 | 0 | 0 | 0 | 0 | 0 | 0 |
| 0 | 0 | 0 | 15 | 0 | 0 | 0 |
| 0 | 0 | 0 | 11 | 0 | 0 | 0 |
| 0 | 0 | 0 | 8 | 0 | 0 | 0 |
| 0 | 0 | 0 | 0 | 0 | 0 | 0 |
| 0 | 0 | 0 | 0 | 0 | 0 | 0 |
| 0 | 0 | 0 | 15 | 0 | 0 | 0 |
| 0 | 0 | 0 | 8 | 0 | 0 | 0 |
| 0 | 0 | 0 | 0 | 0 | 0 | 0 |
| 0 | 0 | 0 | 0 | 0 | 0 | 0 |
| 0 | 0 | 0 | 0 | 0 | 0 | 0 |
| 0 | 0 | 0 | 0 | 0 | 0 | 0 |
| 0 | 0 | 0 | 0 | 0 | 0 | 0 |
| 0 | 0 | 0 | 17 | 0 | 0 | 0 |
| 0 | 0 | 0 | 0 | 0 | 0 | 0 |
| 0 | 0 | 0 | 0 | 0 | 0 | 0 |
| 0 | 0 | 0 | 0 | 0 | 0 | 0 |
| 0 | 0 | 0 | 0 | 0 | 0 | 0 |
| 0 | 0 | 0 | 0 | 0 | 0 | 0 |
| 0 | 0 | 0 | 0 | 0 | 0 | 0 |
| 0 | 0 | 0 | 0 | 0 | 0 | 0 |
| 0 | 0 | 0 | 0 | 0 | 0 | 0 |
| 0 | 0 | 0 | 45 | 0 | 0 | 31 |
| 0 | 0 | 0 | 0 | 0 | 0 | 0 |
| 0 | 0 | 0 | 0 | 0 | 0 | 0 |
| 0 | 0 | 0 | 19 | 0 | 0 | 0 |
| 0 | 0 | 0 | 22 | 0 | 0 | 0 |
| 0 | 0 | 0 | 0 | 0 | 0 | 0 |
| 0 | 0 | 0 | 8 | 0 | 0 | 0 |
| 0 | 0 | 0 | 0 | 0 | 0 | 0 |
| 0 | 0 | 0 | 0 | 0 | 0 | 0 |
| 0 | 0 | 0 | 0 | 0 | 0 | 0 |
| 0 | 0 | 0 | 0 | 0 | 0 | 0 |
| 0 | 0 | 0 | 0 | 0 | 0 | 0 |
| 0 | 0 | 19 | 0 | 0 | 0 | 0 |

| | | | | | | |
|---|---|---|---|---|---|---|
| 0 | 0 | 0 | 0 | 0 | 0 | 0 |
| 0 | 0 | 0 | 0 | 0 | 0 | 0 |
| 0 | 0 | 0 | 0 | 0 | 0 | 0 |
| 0 | 0 | 0 | 7 | 0 | 0 | 0 |
| 0 | 0 | 0 | 0 | 0 | 0 | 0 |
| 0 | 0 | 0 | 0 | 0 | 0 | 0 |
| 0 | 0 | 0 | 6 | 0 | 0 | 0 |
| 0 | 0 | 0 | 0 | 0 | 0 | 0 |
| 0 | 0 | 0 | 0 | 0 | 0 | 0 |
| 0 | 0 | 5 | 0 | 0 | 0 | 0 |
| 0 | 0 | 0 | 0 | 0 | 0 | 0 |
| 0 | 0 | 0 | 0 | 0 | 0 | 0 |
| 0 | 0 | 0 | 8 | 0 | 0 | 0 |
| 0 | 0 | 0 | 0 | 0 | 0 | 0 |
| 0 | 0 | 0 | 0 | 0 | 0 | 0 |
| 0 | 0 | 0 | 0 | 0 | 0 | 0 |
| 0 | 0 | 0 | 0 | 0 | 0 | 0 |
| 0 | 0 | 0 | 0 | 0 | 0 | 0 |
| 0 | 0 | 0 | 0 | 0 | 0 | 0 |
| 0 | 0 | 0 | 0 | 0 | 0 | 0 |
| 0 | 0 | 0 | 0 | 0 | 0 | 0 |
| 0 | 0 | 0 | 0 | 0 | 0 | 0 |
| 0 | 0 | 0 | 0 | 0 | 0 | 0 |
| 0 | 0 | 0 | 0 | 0 | 0 | 0 |
| 0 | 0 | 0 | 18 | 0 | 0 | 0 |
| 0 | 0 | 0 | 0 | 0 | 0 | 14 |
| 0 | 0 | 0 | 0 | 0 | 0 | 0 |
| 0 | 0 | 0 | 0 | 0 | 0 | 0 |
| 0 | 0 | 0 | 0 | 0 | 0 | 0 |
| 0 | 0 | 0 | 0 | 0 | 0 | 0 |
| 0 | 0 | 0 | 0 | 0 | 0 | 0 |
| 0 | 0 | 0 | 0 | 0 | 0 | 0 |
| 0 | 0 | 0 | 5 | 0 | 0 | 0 |
| 0 | 0 | 0 | 8 | 0 | 0 | 0 |
| 0 | 0 | 0 | 0 | 0 | 0 | 0 |
| 0 | 0 | 0 | 0 | 0 | 0 | 0 |
| 0 | 0 | 0 | 0 | 0 | 0 | 0 |
| 0 | 0 | 0 | 0 | 0 | 0 | 0 |
| 0 | 0 | 0 | 0 | 0 | 0 | 0 |
| 0 | 0 | 0 | 6 | 0 | 0 | 0 |
| 0 | 0 | 0 | 0 | 0 | 0 | 0 |
| 0 | 0 | 0 | 0 | 0 | 0 | 0 |
| 0 | 0 | 0 | 24 | 0 | 0 | 0 |
| 0 | 0 | 0 | 0 | 0 | 0 | 0 |
| 0 | 0 | 0 | 0 | 0 | 0 | 0 |
| 0 | 0 | 0 | 0 | 0 | 0 | 0 |
| 0 | 0 | 0 | 0 | 0 | 0 | 0 |
| 0 | 0 | 0 | 0 | 0 | 0 | 0 |
| 0 | 0 | 0 | 0 | 0 | 0 | 0 |
| 0 | 0 | 0 | 0 | 0 | 0 | 0 |

| | | | | | | |
|---|---|---|---|---|---|---|
| 0 | 0 | 0 | 0 | 0 | 0 | 0 |
| 0 | 0 | 0 | 0 | 0 | 0 | 0 |
| 0 | 0 | 0 | 0 | 0 | 0 | 0 |
| 0 | 0 | 0 | 0 | 0 | 0 | 0 |
| 0 | 0 | 0 | 0 | 23 | 0 | 0 |
| 0 | 0 | 0 | 0 | 0 | 0 | 0 |
| 0 | 0 | 0 | 0 | 0 | 0 | 0 |
| 0 | 0 | 0 | 0 | 0 | 0 | 0 |
| 0 | 0 | 0 | 0 | 0 | 0 | 0 |
| 0 | 0 | 0 | 0 | 0 | 0 | 0 |
| 0 | 0 | 0 | 0 | 0 | 0 | 0 |
| 0 | 0 | 0 | 0 | 0 | 0 | 0 |
| 0 | 0 | 0 | 0 | 0 | 0 | 0 |
| 0 | 0 | 0 | 0 | 0 | 0 | 0 |
| 0 | 0 | 0 | 0 | 0 | 0 | 0 |
| 0 | 0 | 0 | 0 | 0 | 2 | 0 |
| 0 | 0 | 0 | 0 | 0 | 0 | 22 |
| 0 | 0 | 0 | 0 | 0 | 0 | 0 |
| 0 | 0 | 0 | 0 | 0 | 0 | 0 |
| 0 | 0 | 0 | 0 | 0 | 0 | 0 |
| 0 | 0 | 0 | 0 | 0 | 0 | 0 |
| 0 | 0 | 0 | 0 | 0 | 0 | 0 |
| 0 | 0 | 0 | 0 | 0 | 0 | 0 |
| 0 | 0 | 0 | 0 | 0 | 0 | 0 |
| 0 | 0 | 0 | 0 | 0 | 0 | 0 |
| 0 | 0 | 0 | 0 | 0 | 0 | 0 |
| 0 | 0 | 0 | 12 | 0 | 0 | 0 |
| 0 | 0 | 0 | 0 | 0 | 0 | 0 |
| 0 | 0 | 0 | 0 | 0 | 0 | 0 |
| 0 | 0 | 0 | 0 | 0 | 0 | 0 |
| 0 | 0 | 0 | 0 | 0 | 0 | 0 |
| 0 | 0 | 0 | 0 | 0 | 0 | 0 |
| 0 | 0 | 0 | 0 | 0 | 0 | 0 |
| 0 | 0 | 0 | 0 | 0 | 0 | 0 |
| 0 | 0 | 0 | 0 | 0 | 20 | 0 |
| 0 | 0 | 0 | 0 | 0 | 0 | 25 |
| 0 | 0 | 0 | 0 | 0 | 0 | 0 |
| 0 | 0 | 0 | 0 | 0 | 0 | 0 |
| 0 | 0 | 0 | 5 | 0 | 0 | 0 |
| 0 | 0 | 0 | 0 | 0 | 0 | 0 |
| 0 | 0 | 0 | 0 | 0 | 0 | 0 |
| 0 | 0 | 0 | 10 | 0 | 0 | 0 |
| 0 | 0 | 0 | 0 | 0 | 0 | 0 |
| 0 | 0 | 0 | 0 | 0 | 0 | 0 |
| 0 | 0 | 0 | 8 | 0 | 0 | 0 |
| 0 | 0 | 0 | 0 | 0 | 0 | 0 |
| 0 | 0 | 0 | 0 | 0 | 0 | 0 |
| 0 | 0 | 0 | 0 | 0 | 0 | 0 |
| 0 | 0 | 0 | 0 | 0 | 0 | 0 |
| 0 | 0 | 0 | 0 | 0 | 0 | 0 |

| | | | | | | |
|---|---|---|---|---|---|---|
| 0 | 0 | 0 | 0 | 0 | 0 | 0 |
| 0 | 0 | 0 | 0 | 0 | 0 | 0 |
| 0 | 0 | 0 | 0 | 0 | 0 | 0 |
| 0 | 0 | 0 | 0 | 0 | 0 | 0 |
| 0 | 0 | 0 | 0 | 0 | 0 | 0 |
| 0 | 0 | 0 | 0 | 0 | 0 | 0 |
| 0 | 0 | 0 | 0 | 0 | 0 | 0 |
| 0 | 0 | 0 | 0 | 0 | 0 | 0 |
| 0 | 0 | 0 | 0 | 0 | 0 | 0 |
| 0 | 0 | 0 | 0 | 0 | 0 | 0 |
| 0 | 0 | 0 | 0 | 0 | 0 | 0 |
| 0 | 0 | 0 | 0 | 0 | 0 | 0 |
| 0 | 0 | 0 | 0 | 0 | 0 | 0 |
| 0 | 0 | 0 | 0 | 0 | 0 | 0 |
| 0 | 0 | 0 | 6 | 0 | 0 | 0 |
| 0 | 0 | 0 | 0 | 0 | 0 | 0 |
| 0 | 0 | 0 | 0 | 0 | 0 | 0 |
| 0 | 0 | 0 | 0 | 0 | 0 | 0 |
| 0 | 0 | 0 | 0 | 0 | 0 | 0 |
| 0 | 0 | 0 | 0 | 0 | 0 | 0 |
| 0 | 0 | 0 | 17 | 0 | 0 | 0 |
| 0 | 0 | 0 | 0 | 0 | 0 | 0 |
| 0 | 0 | 0 | 0 | 0 | 0 | 0 |
| 0 | 0 | 0 | 0 | 0 | 0 | 0 |
| 0 | 0 | 0 | 0 | 0 | 0 | 0 |
| 0 | 0 | 0 | 0 | 0 | 0 | 0 |
| 0 | 0 | 0 | 0 | 0 | 0 | 0 |
| 0 | 0 | 0 | 0 | 0 | 0 | 0 |
| 0 | 0 | 0 | 0 | 0 | 0 | 0 |
| 0 | 0 | 0 | 0 | 0 | 0 | 0 |
| 0 | 0 | 0 | 12 | 0 | 0 | 0 |
| 0 | 0 | 0 | 0 | 0 | 0 | 0 |
| 0 | 0 | 0 | 0 | 0 | 0 | 0 |
| 0 | 0 | 0 | 0 | 0 | 4 | 0 |
| 0 | 0 | 0 | 0 | 0 | 0 | 0 |
| 0 | 0 | 0 | 0 | 0 | 0 | 0 |
| 0 | 0 | 0 | 0 | 0 | 0 | 0 |
| 0 | 0 | 0 | 0 | 0 | 0 | 0 |
| 0 | 0 | 0 | 0 | 0 | 0 | 0 |
| 0 | 0 | 0 | 8 | 0 | 0 | 0 |
| 0 | 0 | 0 | 0 | 0 | 0 | 0 |
| 0 | 0 | 0 | 0 | 0 | 0 | 0 |
| 0 | 0 | 0 | 0 | 0 | 0 | 0 |
| 0 | 0 | 0 | 0 | 0 | 0 | 0 |
| 0 | 0 | 0 | 0 | 0 | 0 | 0 |
| 0 | 0 | 0 | 0 | 0 | 0 | 0 |
| 0 | 0 | 0 | 7 | 0 | 0 | 0 |
| 0 | 0 | 0 | 7 | 0 | 0 | 0 |
| 0 | 0 | 0 | 0 | 0 | 0 | 0 |
| 0 | 0 | 0 | 0 | 0 | 0 | 0 |

| | | | | | | |
|---|---|---|---|---|---|---|
| 0 | 0 | 0 | 0 | 0 | 0 | 0 |
| 0 | 0 | 0 | 0 | 0 | 0 | 0 |
| 0 | 0 | 0 | 0 | 0 | 0 | 0 |
| 0 | 0 | 0 | 0 | 0 | 0 | 0 |
| 0 | 0 | 0 | 0 | 0 | 0 | 0 |
| 0 | 0 | 0 | 0 | 0 | 0 | 0 |
| 0 | 0 | 0 | 0 | 0 | 0 | 0 |
| 0 | 0 | 0 | 0 | 0 | 0 | 0 |
| 0 | 0 | 0 | 0 | 0 | 0 | 0 |
| 0 | 0 | 0 | 0 | 0 | 0 | 0 |
| 0 | 0 | 0 | 0 | 0 | 0 | 0 |
| 0 | 0 | 0 | 5 | 0 | 0 | 13 |
| 0 | 0 | 0 | 0 | 0 | 0 | 0 |
| 0 | 0 | 0 | 0 | 0 | 0 | 0 |
| 0 | 0 | 0 | 0 | 0 | 0 | 0 |
| 0 | 0 | 0 | 0 | 0 | 0 | 0 |
| 0 | 0 | 0 | 0 | 0 | 0 | 0 |
| 0 | 0 | 0 | 0 | 0 | 0 | 0 |
| 0 | 0 | 0 | 0 | 0 | 0 | 0 |
| 0 | 0 | 0 | 0 | 0 | 0 | 0 |
| 0 | 0 | 0 | 0 | 0 | 0 | 0 |
| 0 | 0 | 0 | 0 | 0 | 0 | 0 |
| 0 | 0 | 0 | 0 | 0 | 0 | 0 |
| 0 | 0 | 0 | 0 | 0 | 0 | 0 |
| 0 | 0 | 0 | 0 | 0 | 0 | 0 |
| 0 | 0 | 0 | 0 | 12 | 0 | 0 |
| 0 | 0 | 0 | 0 | 0 | 0 | 0 |
| 0 | 0 | 0 | 0 | 0 | 0 | 0 |
| 0 | 0 | 0 | 0 | 0 | 0 | 0 |
| 0 | 0 | 0 | 0 | 0 | 0 | 0 |
| 0 | 0 | 0 | 0 | 0 | 0 | 0 |
| 0 | 0 | 0 | 0 | 0 | 0 | 0 |
| 0 | 0 | 0 | 0 | 0 | 0 | 0 |
| 0 | 0 | 0 | 0 | 0 | 0 | 0 |
| 0 | 0 | 0 | 0 | 0 | 0 | 0 |
| 0 | 0 | 0 | 0 | 0 | 0 | 0 |
| 0 | 0 | 0 | 0 | 0 | 0 | 0 |
| 0 | 0 | 0 | 0 | 0 | 0 | 0 |
| 0 | 0 | 0 | 0 | 0 | 0 | 0 |
| 0 | 0 | 0 | 0 | 0 | 0 | 0 |
| 0 | 0 | 0 | 0 | 0 | 0 | 0 |
| 0 | 0 | 0 | 0 | 0 | 0 | 0 |
| 0 | 0 | 0 | 0 | 0 | 0 | 0 |
| 0 | 0 | 0 | 0 | 0 | 0 | 0 |
| 0 | 0 | 0 | 0 | 0 | 0 | 0 |
| 0 | 0 | 0 | 0 | 0 | 0 | 0 |
| 0 | 0 | 0 | 6 | 0 | 0 | 0 |
| 0 | 0 | 0 | 0 | 0 | 0 | 0 |
| 0 | 0 | 0 | 0 | 0 | 0 | 0 |
| 0 | 0 | 0 | 0 | 0 | 0 | 0 |

| | | | | | | |
|---|---|---|---|---|---|---|
| 0 | 0 | 0 | 0 | 0 | 0 | 0 |
| 0 | 0 | 0 | 0 | 0 | 0 | 0 |
| 0 | 0 | 0 | 0 | 0 | 0 | 0 |
| 0 | 0 | 0 | 0 | 0 | 0 | 0 |
| 0 | 0 | 0 | 0 | 0 | 0 | 0 |
| 0 | 0 | 0 | 0 | 0 | 0 | 0 |
| 0 | 0 | 0 | 0 | 0 | 0 | 0 |
| 0 | 0 | 0 | 0 | 0 | 0 | 0 |
| 0 | 0 | 0 | 0 | 0 | 0 | 0 |
| 0 | 0 | 0 | 0 | 0 | 0 | 0 |
| 0 | 0 | 0 | 0 | 0 | 0 | 0 |
| 0 | 0 | 0 | 0 | 0 | 0 | 0 |
| 0 | 0 | 0 | 0 | 0 | 0 | 0 |
| 0 | 0 | 0 | 0 | 0 | 0 | 0 |
| 0 | 0 | 0 | 0 | 0 | 0 | 0 |
| 0 | 0 | 0 | 0 | 0 | 0 | 0 |
| 0 | 0 | 0 | 0 | 0 | 0 | 0 |
| 0 | 0 | 0 | 9 | 0 | 0 | 0 |
| 0 | 0 | 0 | 9 | 0 | 0 | 0 |
| 0 | 0 | 0 | 0 | 0 | 0 | 0 |
| 0 | 0 | 0 | 0 | 0 | 0 | 0 |
| 0 | 0 | 0 | 0 | 0 | 0 | 0 |
| 0 | 0 | 0 | 0 | 0 | 0 | 0 |
| 0 | 0 | 0 | 0 | 0 | 0 | 0 |
| 0 | 0 | 0 | 0 | 0 | 0 | 0 |
| 0 | 0 | 0 | 0 | 0 | 0 | 0 |
| 0 | 0 | 0 | 0 | 0 | 0 | 0 |
| 0 | 0 | 0 | 0 | 0 | 0 | 0 |
| 0 | 0 | 0 | 0 | 0 | 0 | 0 |
| 0 | 0 | 0 | 0 | 0 | 0 | 0 |
| 8 | 0 | 0 | 0 | 0 | 0 | 0 |
| 0 | 0 | 0 | 8 | 0 | 0 | 0 |
| 0 | 0 | 0 | 0 | 0 | 0 | 0 |
| 0 | 0 | 0 | 0 | 0 | 0 | 0 |
| 0 | 0 | 0 | 0 | 0 | 0 | 0 |
| 0 | 0 | 0 | 0 | 0 | 0 | 0 |
| 0 | 0 | 0 | 0 | 0 | 0 | 0 |
| 0 | 0 | 0 | 0 | 0 | 0 | 0 |
| 0 | 0 | 0 | 0 | 0 | 0 | 0 |
| 0 | 0 | 0 | 7 | 0 | 0 | 0 |
| 0 | 0 | 0 | 0 | 0 | 0 | 0 |
| 0 | 0 | 0 | 0 | 0 | 0 | 0 |
| 0 | 0 | 0 | 0 | 0 | 0 | 0 |
| 0 | 0 | 0 | 0 | 0 | 0 | 0 |
| 0 | 0 | 0 | 0 | 0 | 0 | 0 |
| 0 | 0 | 0 | 0 | 0 | 0 | 0 |
| 0 | 0 | 0 | 0 | 0 | 0 | 0 |
| 0 | 0 | 0 | 0 | 0 | 0 | 0 |
| 0 | 0 | 0 | 0 | 0 | 0 | 0 |
| 0 | 0 | 0 | 0 | 0 | 0 | 0 |

| | | | | | | |
|---|---|---|---|---|---|---|
| 0 | 0 | 0 | 0 | 0 | 0 | 0 |
| 0 | 0 | 0 | 0 | 0 | 0 | 0 |
| 0 | 0 | 0 | 0 | 0 | 0 | 0 |
| 0 | 0 | 0 | 0 | 0 | 0 | 0 |
| 0 | 0 | 0 | 0 | 0 | 0 | 0 |
| 0 | 0 | 6 | 0 | 0 | 0 | 0 |
| 0 | 0 | 0 | 6 | 0 | 0 | 0 |
| 0 | 0 | 0 | 6 | 0 | 0 | 0 |
| 0 | 0 | 0 | 0 | 0 | 0 | 0 |
| 0 | 0 | 0 | 0 | 0 | 0 | 0 |
| 0 | 0 | 0 | 0 | 0 | 0 | 0 |
| 0 | 0 | 0 | 0 | 0 | 0 | 0 |
| 0 | 0 | 0 | 0 | 0 | 0 | 0 |
| 0 | 0 | 0 | 0 | 0 | 0 | 0 |
| 0 | 0 | 0 | 0 | 0 | 0 | 0 |
| 0 | 0 | 0 | 0 | 0 | 0 | 0 |
| 0 | 0 | 0 | 0 | 0 | 0 | 0 |
| 0 | 0 | 0 | 0 | 0 | 0 | 0 |
| 0 | 0 | 0 | 0 | 0 | 0 | 0 |
| 0 | 0 | 0 | 0 | 0 | 0 | 0 |
| 0 | 0 | 0 | 0 | 0 | 0 | 0 |
| 0 | 0 | 0 | 0 | 0 | 0 | 0 |
| 0 | 0 | 0 | 0 | 0 | 0 | 0 |
| 0 | 0 | 0 | 0 | 0 | 0 | 0 |
| 0 | 0 | 0 | 0 | 0 | 0 | 0 |
| 0 | 0 | 0 | 0 | 0 | 0 | 0 |
| 0 | 0 | 0 | 0 | 0 | 0 | 0 |
| 0 | 0 | 0 | 0 | 0 | 0 | 0 |
| 0 | 0 | 0 | 0 | 0 | 0 | 0 |
| 0 | 0 | 0 | 0 | 0 | 0 | 0 |
| 0 | 0 | 0 | 0 | 0 | 0 | 0 |
| 0 | 0 | 0 | 0 | 0 | 0 | 0 |
| 0 | 0 | 0 | 0 | 3 | 0 | 0 |
| 0 | 0 | 0 | 0 | 3 | 0 | 0 |
| 0 | 0 | 0 | 0 | 0 | 3 | 0 |
| 0 | 0 | 0 | 0 | 0 | 0 | 0 |
| 0 | 0 | 0 | 0 | 0 | 0 | 0 |
| 0 | 0 | 0 | 0 | 0 | 0 | 0 |
| 0 | 0 | 0 | 0 | 0 | 0 | 0 |
| 0 | 0 | 0 | 0 | 2 | 0 | 0 |
| 0 | 0 | 0 | 2 | 0 | 0 | 0 |
| 0 | 0 | 0 | 0 | 0 | 0 | 0 |
